# Viral piracy of host RNA phosphatase DUSP11 by avipoxviruses

**DOI:** 10.1101/2024.08.06.606876

**Authors:** Kayla H. Szymanik, Dustin C. Hancks, Christopher S. Sullivan

## Abstract

Proper recognition of viral pathogens is an essential part of the innate immune response. A common viral replicative intermediate and chemical signal that cells use to identify pathogens is the presence of a triphosphorylated 5’ end (5’ppp) RNA, which activates the cytosolic RNA sensor RIG-I and initiates downstream antiviral signaling. While 5’pppRNA generated by viral RNA-dependent RNA polymerases (RdRps) can be a potent activator of the immune response, endogenous RNA polymerase III (RNAPIII) transcripts can retain the 5’pppRNA generated during transcription and induce a RIG-I-mediated immune response. We have previously shown that host RNA triphosphatase dual-specificity phosphatase 11 (DUSP11) can act on both host and viral RNAs, altering their levels and reducing their ability to induce RIG-I activation. Our previous work explored how artificially altered DUSP11 can impact immune activation, prompting further exploration into natural contexts of altered DUSP11. Here, we have identified viral DUSP11 homologs (vDUSP11s) present in some avipoxviruses. Consistent with the known functions of endogenous DUSP11, we have shown that expression of vDUSP11s: 1) reduces levels of endogenous RNAPIII transcripts, 2) reduces a cell’s sensitivity to 5’pppRNA-mediated immune activation, and 3) restores virus infection defects seen in the absence of DUSP11. Our results identify a virus-relevant context where DUSP11 activity has been co-opted to alter RNA metabolism and influence the outcome of infection.

## Introduction

The interface between host defenses and viral antagonists is an evolutionary “arms race,” where both groups are under constant pressure to gain the advantage (1,2). Host cells are armed with specialized pattern recognition receptors (PRRs) to sense conserved pathogen signatures, known as pathogen-associated molecular patterns (PAMPs), that initiate downstream antiviral signaling and protect against infection (3). Viruses encode proteins to combat these pathways, and while they are limited by their coding capacity, they have a unique advantage due to their shorter generation times and faster mutation rates. This leaves host cells chasing a moving target, challenging both the specificity and selectivity of interactions.

RIG-I is a cytosolic PRR best characterized to detect structured or double-stranded RNA (dsRNA) bearing a di/tri-phosphate on the 5’-end, a common viral replicative intermediate generated by viral RNA-dependent RNA polymerases (RdRPs) (4–6). Cellular RNAs are initially generated with a 5’-triphosphate end, and transcription via RNAP III occurs without a co-transcriptional capping mechanism (7). In addition, recent studies have shown that RNAs generated by RNAP III can function as endogenous damage-associated molecular patterns (DAMPs) and activate RIG-I during virus infection (8–10). This necessitates proper regulation of these signals to prevent immune activation in the absence of a pathogen.

DUSP11 is an RNA 5’-triphosphatase that converts 5’-di or triphosphorylated RNAs into their monophosphorylated form and can act on both host and viral RNAs (11–15). DUSP11 activity on RNAs reduces their immunogenicity but also renders them more susceptible to nuclease decay (13,14). Current literature details varying relationships between DUSP11 and virus infection. In the case of Hepatitis C virus (HCV), 5’-triphosphorylated transcripts are converted to monophosphates by DUSP11, rendering them susceptible to attack by endogenous exoribonucleases such as XRNs (13,16). For bovine leukemia virus (BLV) and adenoviruses, viral miRNA processing is influenced by the presence of DUSP11, featuring a shift in the ratio of 5p:3p miRNA abundance in the absence of DUSP11 (12). DUSP11 has been shown to modulate numerous host RNAP III transcripts, such as Y RNAs, SINEs, and vault RNAs (12). We have previously demonstrated that DUSP11 activity renders cells and mice less active in inducing a RIG-I-mediated immune response (14). These studies demonstrate that DUSP11 activity can be either pro-viral or anti-viral depending on the context, yet so far, there are few natural contexts identified in which DUSP11 activity levels are altered during infection (8,9).

Viruses encode diverse proteins targeting components of the host’s immune response to escape immune detection. Some viruses incorporate host genes into their genomes throughout their evolutionary history, and a large fraction of these "pirated" genes are involved in evading host antiviral defenses (3,17). Here, we have identified the presence of DUSP11 homologs (vDUSP11) in some avipoxviruses (APVs) and unclassified avipox-adjacent poxviruses (AdjPVs). Previous studies have shown that while poxviruses are DNA viruses, their infection can activate RIG-I (3,18). Due to the established role of DUSP11 in modulating the innate immune response, we hypothesize that these viruses have acquired vDUSP11 with conserved catalytic activity to aid in immune evasion.

## Results

### DUSP11 (vDUSP11) genes are encoded by a subset of avipox and unclassified avipox-adjacent poxvirus genomes

DUSP11 is a member of the dual specificity phosphatase subgroup of type-I based cysteine-based protein tyrosine phosphatases (PTPs). Most members of the DUSP subgroup are active on protein substrates, but DUSP11 is atypical in that it is more active on RNA substrates (11). Human DUSP11 (hDUSP11) (Chr2P13.1, O75319) has three isoforms, the major isoform being 330 amino acids (AA) in length. All DUSPs share a highly conserved phosphate-binding loop (P-loop) containing the consensus phosphatase sequence (HCXXXXXR; AA: 151-158 in human) (19) (Fig 1A). In addition, DUSP11 contains an R residue (R192 in hDUSP11), which has been previously suggested to be involved in the β-phosphatase activity, unique to RNA 5’-triphosphatases and lacking in mRNA capping enzymes (20). Backed by recent insight into the immunomodulatory abilities of DUSP11, we hypothesized viruses might co-opt this activity. To this end, we set out to determine whether vertebrate virus genomes contain genes with similar features to DUSP11.

**Figure 1:**
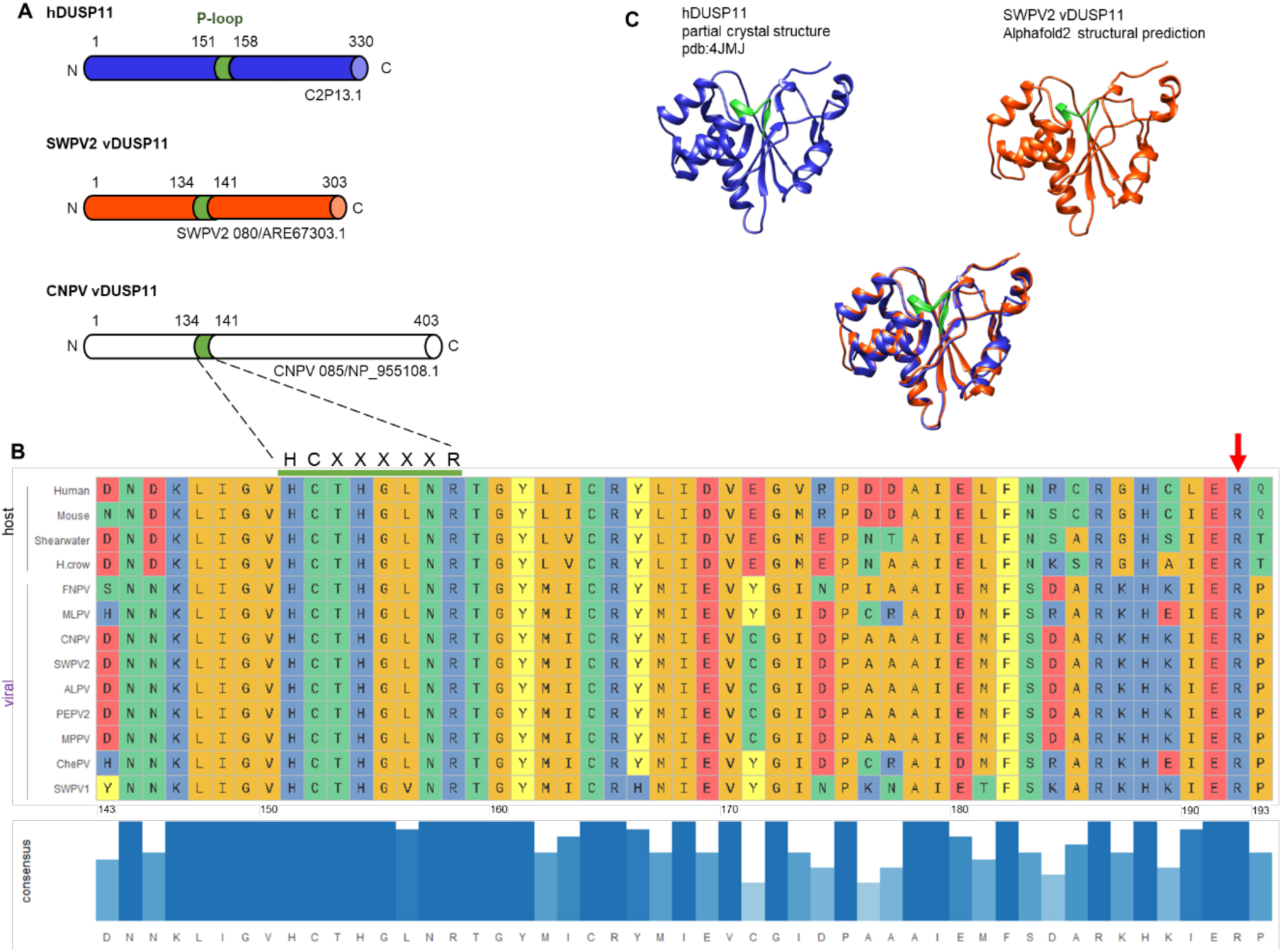
Avipoxviruses and related unclassified avipox-adjacent poxviruses encode putative homolog of DUSP11. (A). Schematic of human *DUSP11* (hDUSP11) and select putative viral *DUSP11s* with predicted domains indicated. Colored domain indicates hDUSP11 P-loop with corresponding identical sequences present in putative viral DUSP11 homologs (vDUSP11). Putative vDUSP11 include shearwaterpox virus 2 vDUSP11 (SWPV2 vDUSP11) and canarypox virus vDUSP11 (CNPV vDUSP11). (B). Key catalytic residues are conserved between hDUSP11 and putative avipox vDUSP11. Multiple sequence comparison by log-expectation (Clustal) amino acid (AA) sequence alignment of human DUSP11, mouse DUSP11, *C. borealis* (Shearwater), and putative vDUSP11s from shearwaterpox virus 1 (SWPV1), finchpox virus (FNPV), mudlarkpox virus (MLPV), cheloniidpox virus (ChePV), canarypox virus (CNPV), penguinpox virus 2 (PEPV2), albatrosspox virus (ALPV), magpiepox virus (MPPV), and shearwaterpox virus 2 (SWPV2). A subset of AAs surrounding the hDUSP11 P-loop are shown. Amino acids are colored based on side-chain chemistry. Bar graph indicating relative sequence conservation with consensus amino acid sequence determined from listed sequences. Taller bar height and darker blue color correspond to relative correlation to consensus sequence. The numbering below the consensus sequence corresponds to hDUSP11 amino acid numbering. hDUSP11 residue R192 is indicated by a red arrow. (C). Partial crystal structure of hDUSP11 (pdb:4JMJ) (blue) and alphafold2 structural prediction of corresponding region from putative shearwaterpox virus 2 vDUSP11 (orange) with structural overlay. Key catalytic residues (P-loop) present in hDUSP11 are indicated in green, and corresponding residues are also indicated on the putative SWPV2 vDUSP11 structural prediction.

Using BLASTp analysis (21), we identified several APVs/AdjPVs predicted to encode proteins featuring high sequence identity (∼44-60%) to human (hDUSP11) (Table S2). Of those viruses, we initially sought to evaluate the putative viral homolog from one of the better-studied APVs, canarypox virus (CNPV), which encodes a 403 AA predicted homolog to DUSP11 (CNPV085/NP_955108.1) [44% (96/217) amino acid identity, 64% positives (140/217)] (Fig 1A). Canarypox viruses have been studied largely for use as a recombinant vaccine vector (22,23). However, due to initial difficulties cloning the putative CNPV viral homolog, we pursued the closely related virus shearwaterpox virus 2 (SWPV2) as the model putative viral DUSP11 homolog. SWPV2 genome contains a 303 AA predicted ORF (SWPV080/ARE67303.1) featuring high sequence identity to hDUSP11 [50% (94/188) amino acid identity, 67% positives (121/188)] (Fig 1A), hereafter vDUSP11. Notably, this protein includes the corresponding R192 residue in SWPV2 vDUSP11 (AA: 161) (Fig 1B), suggesting a distinct role from existing mRNA capping enzymes. Using AlphaFold2 (24), we generated a structural prediction of the SWPV2 putative vDUSP11 homolog and aligned it to a partial crystal structure of hDUSP11(PDB: 4JMJ; AA: 28-208) (25) (Fig 1C). This structural analysis highlights that hDUSP11 and SWPV2 vDUSP11 are nearly superimposable, excluding the C-terminal extension of the viral protein, and that key catalytic residues (shown in green) are structurally conserved.

Using the SWPV2 vDUSP11 AA sequence as a query returns numerous host DUSP11 sequences but not other phosphatases from related viruses, such as Vaccinia VH1 (Table S3). This reciprocal BLASTp analysis indicates that the progenitor of SWPV2 vDUSP11 was likely acquired by horizontal gene transfer (HGT) from a host copy of DUSP11. Many of the avipoxviruses that encode vDUSP11 infect hosts within the order of *Passeriformes*, making them a potential source of DUSP11. Further searches detected a total of 11 DUSP11 ORFs encoded by other APV/AdjPVs, 2 of which are notably smaller (Fig S1 & Table S4). These findings show that multiple APV/AdjPVs encode proteins that resemble DUSP11.

To further examine the acquisition of the putative vDUSP11 in an evolutionary context, we generated an inferred phylogenetic tree using host DUSP AA sequences, the putative avipox viral DUSP11 sequences, and the poxviral VH1-like DUSPs from Vaccinia virus (VACV) and corresponding avipoxvirus homologs (Table S5). The putative avipox vDUSP11 cluster with related host DUSP11, and not with the other poxviral VH1-like DUSPs (Fig 2). These further support the avipox vDUSP11 arose from host DUSP11 and not other viral or host DUSPs.

**Figure 2:**
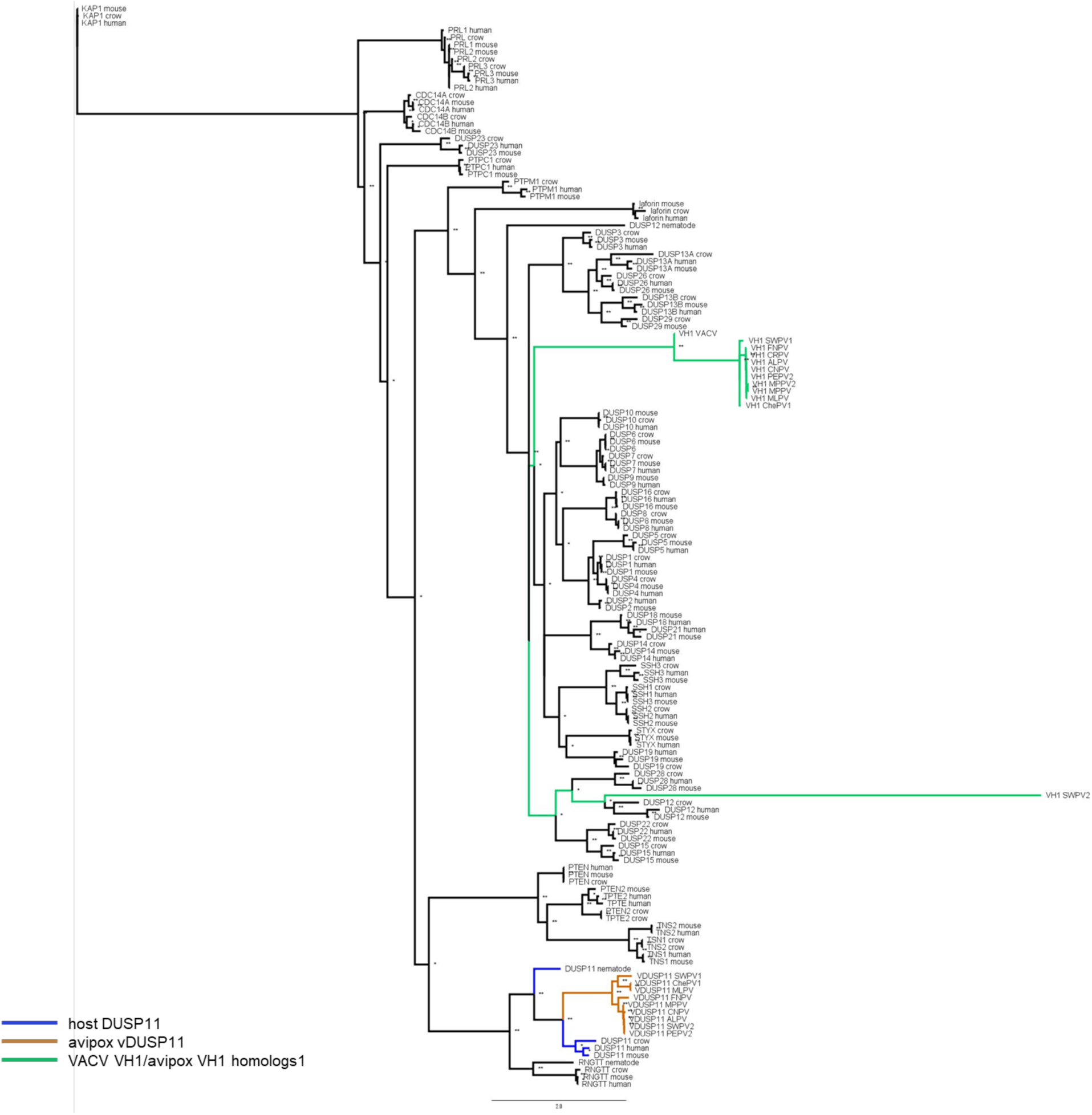
Phylogenetic analysis supports putative vDUSP11 acquisition from host DUSP11 sequence. An inferred tree built using 150 DUSP amino acid (AA) sequences by maximum-likelihood analysis phylogenetic tree using PhyML (38). AA sequences for host DUSPs, poxviral putative vDUSP11s, and Vaccinia virus DUSP *H1L* protein phosphatase were aligned using Clustal Omega (Fig S3) (49). Clustal alignment was used to run PhyML analysis with LG +R model selected by SMS (50). Sequences were retrieved from the NCBI sequence database (37) and Uniprot (www.uniprot.org/) (Table S5). Putative APV/AdjPV vDUSP11s (orange) cluster with host DUSP11s (blue) and not with other related host protein DUSPs or Vaccinia VH1 or avipox homolog protein DUSP (green). 100 bootstrap replicates were performed; branch support >50% (*) or >70% (**) are indicated.

### vDUSP11s are enzymatically active

Based on AA sequence and predicted structural similarity, we hypothesized that these vDUSP11s possess similar catalytic activities as hDUSP11. hDUSP11 has been demonstrated to be active on 5’-triphosphate RNA, sequentially liberating both the γ- and β-phosphates, leaving a 5’-monophosphate (26). This modification renders RNA susceptible to decay via monophosphate-dependent 5’-3’ exoribonuclease XRN1. We hypothesized that vDUSP11s are catalytically active RNA 5’-triphosphatases capable of performing similar functions as hDUSP11.

To assess this hypothesis, we applied a previously established assay to measure the ability of vDUSP11 to sensitize *in vitro* transcribed RNAs to decay via XRN1 (27,28). 5’-RNA phosphatase treatment followed by XRN treatment of a hepatitis C virus (HCV) 5’-UTR RNA results in the generation of a stable quantifiable smaller fragment (13,27,28). We generated *in vitro* transcribed and translated flag-tagged full-length proteins for hDUSP11, negative control hDUSP11 catalytically inactive mutant (D11-CM) (C152S), the SWPV2 vDUSP11, and the predicted SWPV2 vDUSP11 catalytically inactive mutant (SWPV-CM) (C135S). We also included an additional negative control of untagged luciferase. Protein expression was confirmed via immunoblot analysis (Fig 3A). We treated *in vitro* transcribed HCV 5’ UTR RNA with the *in vitro* transcribed/translated constructs. Following phosphatase treatment, we purified the RNA and then treated ± recombinant XRN1. We evaluated the abundance and length of RNA by using a urea-PAGE gel stained with Ethidium Bromide (EtBr) to visualize RNA. All conditions using treatment with XRN1 resulted in a small degree of generation of a shorter RNA fragment, suggesting low levels of susceptible monophosphate RNA present in the substrate population (Fig 3B). However, treatment with catalytically active hDUSP11 or the SWPV2 vDUSP11, but not either of the catalytic mutants (CM), resulted in a substantial reduction in the full-length HCV 5’ UTR fragment.

**Figure 3:**
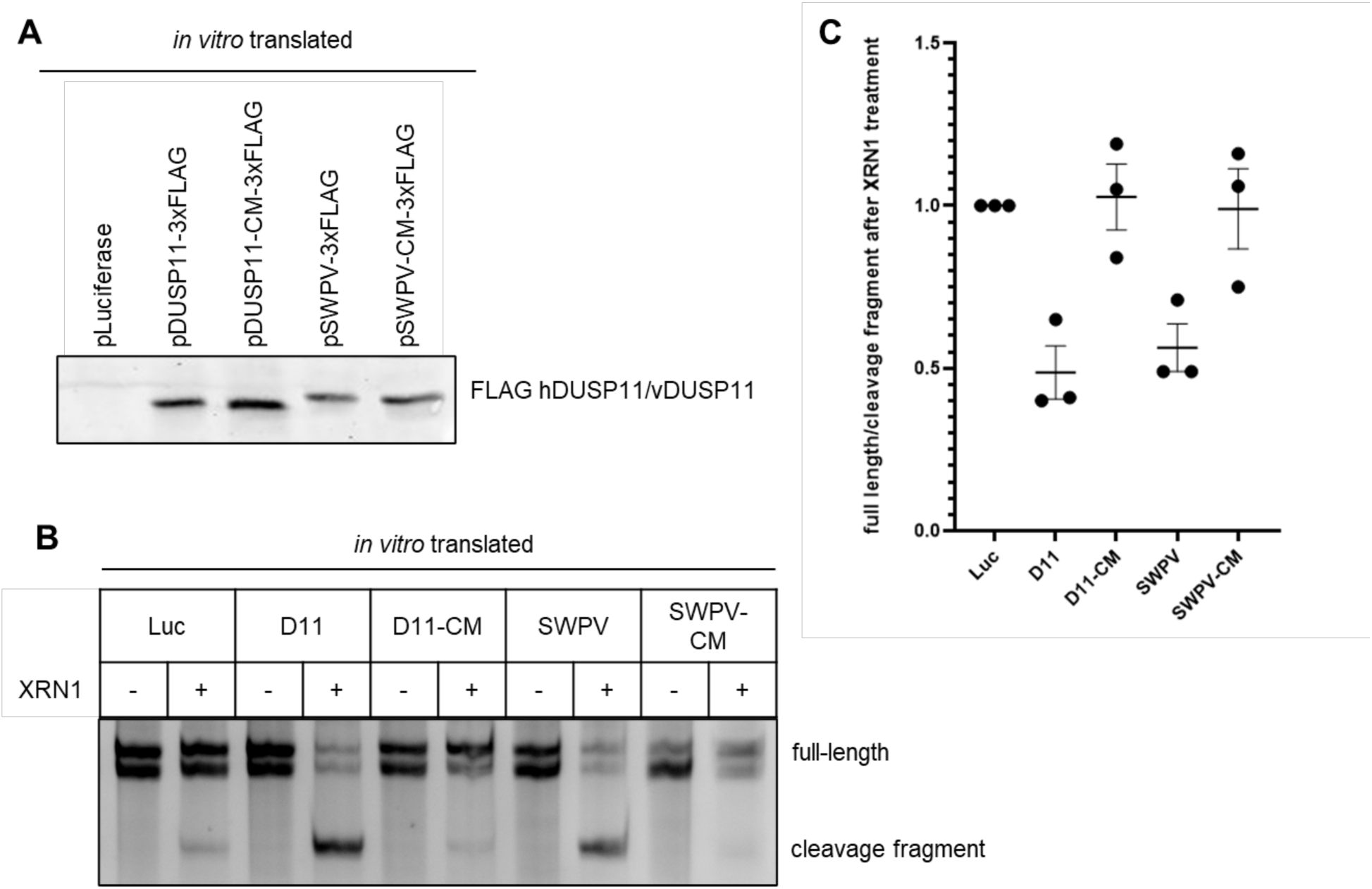
vDUSP11 sensitizes HCV 5’ UTR RNA to XRN-mediated degradation. (A). Confirmation of *in vitro* translated constructs via immunoblot analysis. Membrane was probed for FLAG-tagged proteins. Plasmid encoding luciferase was used for negative control reactions. (B). *In vitro* XRN susceptibility assay. *In vitro* transcribed HCV 5’ UTR RNA was incubated with *in vitro* translated products from (A) (luciferase (Luc), hDUSP11 (D11), hDUSP11 catalytic mutant (D11-CM), shearwaterpox virus 2 vDUSP11 (SWPV) or shearwaterpox virus 2 vDUSP11 catalytic mutant (SWPV-CM)) and RNA was purified. Purified RNA was then subjected to treatment ± recombinant XRN1. Products were separated using urea PAGE and stained with EtBr. Migration of full-length HCV 5’ UTR is indicated, which for unknown reasons migrates as a doublet, the position of the faster-migrating cleavage fragment is indicated. (C). Graphical representation of (B), displaying relative band intensity of HCV 5’ UTR full-length to cleavage fragment following treatment with recombinant XRN1 (+XRN1/-XRN1). Values are normalized to the luciferase negative control. Data are derived from n = 3 independent replicates. In all panels, data are represented as mean ± SEM.

Quantification indicated a ∼30-50% increase in the ratio of degradation fragment to full-length RNA seen following XRN1 treatment (Fig 3B-C). This result is what is expected for the known catalytic activity of DUSP11 to act on 5’-triphosphorylated RNA, sensitizing them to decay via XRN1. Thus, SWPV2 vDUSP11 is a catalytically active 5’-RNA triphosphatase and this activity depends on cysteine in the catalytic pocket, as previously demonstrated for hDUSP11.

### vDUSP11s modulate sensitivity to immune activation triggered by 5’-triphosphate RNAs

We next determined if vDUSP11 shares additional functions with hDUSP11. RIG-I is a key PRR that recognizes structured or double-stranded 5’ di-/tri-phosphorylated RNAs, which, when activated, leads to the induction of antiviral signaling, including transcripts such as ISG15 and IFN-β (4–6). We hypothesized that APVs/AdjPVs have acquired vDUSP11 to mimic host DUSP11’s immunomodulatory properties, which alter the sensitivity of RIG-I activation and modulate RNAP III transcripts upregulated in the context of various virus infections (8,9,14).

To assay the functions of the vDUSP11s in cell culture, we transfected immunostimulatory RNAs (Fig 4A) into previously described A549 DUSP11 knockout (KO) cells (29) stably expressing various DUSP11s and measured RIG-I activity via IFN-stimulated gene (ISG) expression. Included in this analysis is a second full-length vDUSP11 from cheloniidpox virus 1 (ChePV) [44% (88/201) amino acid identity, 66% positives (133/201) to hDUSP11]. We generated cells stably reconstituted with a control empty-vector (EV) or the following N-terminally flag-tagged proteins: 1) wild-type hDUSP11 (D11), 2) hDUSP11 catalytic-mutant (C152S, D11-CM), 3) SWPV2 vDUSP11 (SWPV2), 4) SWPV2 catalytic-mutant vDUSP11 (C135S, SWPV2-CM), 5) ChePV vDUSP11 (ChePV), 6) ChePV catalytic-mutant of ChePV (C135S, ChePV-CM), or 7) phosphatase DUSP12 (D12) as a negative control protein. Protein expression for each was confirmed via immunoblot analysis (Fig 4B).

**Figure 4:**
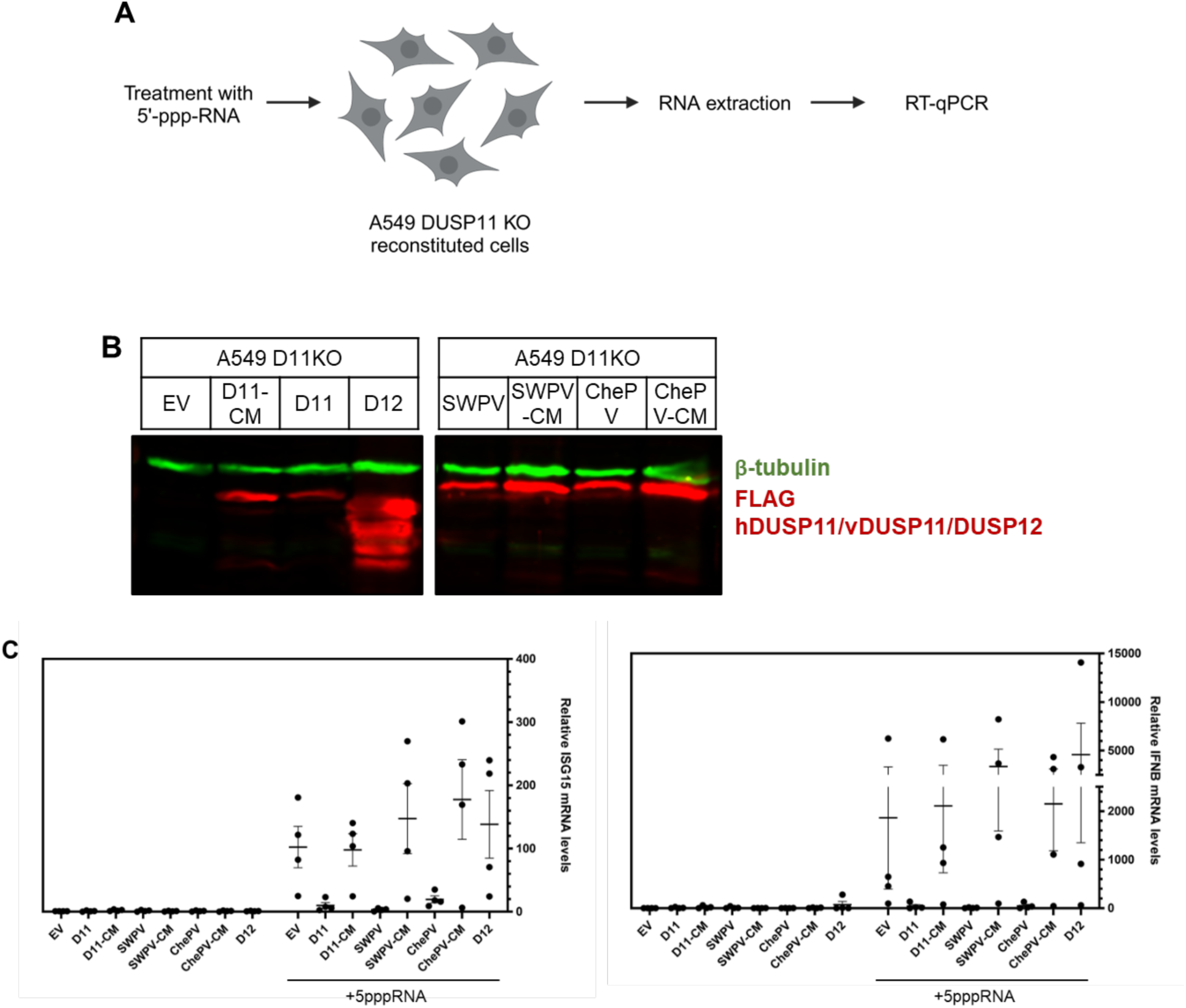
vDUSP11 modulates immune activation in response to liposomal 5’-ppp-RNAs. (A). Schematic diagram of the 5’-triphosphate RNA (5’-ppp-RNA) transfection assay. A549 DUSP11 knockout (KO) reconstituted cells (12-well) were transfected with 5-10 ng of *in vitro* transcribed 5’-ppp-RNA for 18 hours followed by RT-qPCR to assay induction of ISGs. (B). Immunoblot analysis of A549 DUSP11 knockout (KO) cells transduced with pLenti empty vector (EV), pLenti hDUSP11-3xFLAG (D11), pLenti hDUSP11 catalytic mutant-3xFLAG (D11-CM), pLenti shearwaterpox virus 2 vDUSP11-3xFLAG (SWPV), pLenti shearwaterpox virus 2 vDUSP11 catalytic mutant-3xFLAG (SWPV-CM), pLenti cheloniidpox virus 1 vDUSP11-3xFLAG (ChePV), pLenti 3xFLAG-cheloniidpox virus 1 vDUSP11 catalytic mutant-3xFLAG (ChePV-CM), or negative control pLenti DUSP12-3xFLAG (D12). (C). RT-qPCR analysis of *ISG15* and *IFNB1* mRNA normalized to *GAPDH* mRNA in 5’-ppp-RNA transfected in A549 DUSP11 KO reconstituted cells. Results are represented relative to those of empty vector-expressing cells. Data are derived from n = 4 independent replicates in and are represented as mean ± SEM.

Upon transfection of 5’-triphosphate RNA, cells expressing hDUSP11, SWPV vDUSP11, or ChePV vDUSP11, but not the empty-vector nor any catalytic mutants/negative controls, displayed a reduction in both induced *ISG15* and *IFN-β* ISG transcript levels (Fig 4C and Fig S5). Compared to empty vector cell lines, those containing either hDUSP11 or the SWPV and ChePV vDUSP11s showed on average a ∼40-50-, ∼40-90-, and ∼30-50-percent reduction in induced *ISG15* mRNA levels, respectively, and all showed a greater than a ∼70-90-percent average reduction in induced *IFNB1* mRNA levels. These results indicate that both vDUSP11s can modulate immune activation, as previously shown for hDUSP11.

### vDUSP11 catalytic activity promotes VSV virus replication

The role of DUSP11 in virus infection is context-dependent, however, previous work from our lab has shown that some viruses (e.g. +ssRNA sindbis virus (SINV) and -RNA vesicular stomatitis virus (VSV)) benefit from DUSP11 activity during infection (14). Both SINV and VSV have been shown to produce 5’-triphosphorylated RNA as part of their replication cycle. Additionally, host RNAs have also been implicated in activating RIG-I upon infection with some RNA viruses (9). We hypothesized that vDUSP11s can enhance virus infection in a manner similar to DUSP11. To investigate this, we infected A549 DUSP11 KO cells reconstituted with the various viral and human DUSP11s with M51R mutant VSV for 48 hours at an MOI of 0.01 PFU/cell. VSV M51R lacks the ability to fully block host gene expression and, as a result, stimulates a stronger interferon response (30).

Cells expressing either hDUSP11, SWPV vDUSP11, or ChePV vDUSP11, but not catalytic mutants or negative controls (D12/EV) enhanced VSV infection (Fig 5A-C). Compared to cells expressing only the empty vector, those reconstituted with hDUSP11 had, on average, a ∼7-fold increase in the total virus they produced, consistent with previously published findings (14). Cells expressing the SWPV vDUSP11 had, on average, a ∼20-fold increase in viral titer, and those expressing the ChePV vDUSP11 had a ∼3-fold increase in viral titer. These findings lend further support to evolutionarily conserved shared functionality between host and viral DUSP11s.

**Figure 5:**
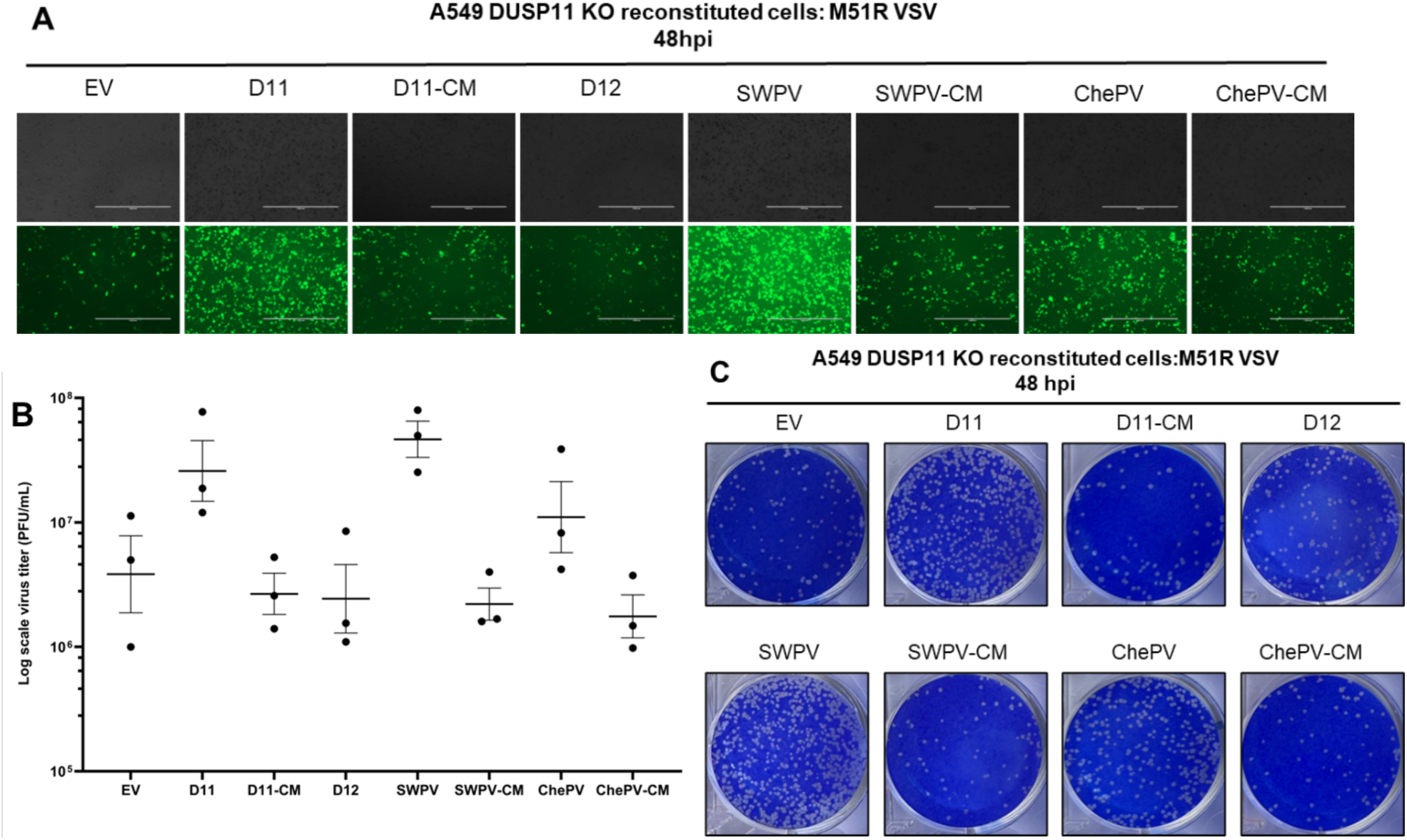
vDUSP11 catalytic activity promotes VSV virus replication. (A). Representative GFP images of A549 DUSP11 knockout (KO) cells stably reconstituted with either empty vector (EV) plasmid, hDUSP11 (D11), hDUSP11 catalytic mutant (D11-CM), shearwaterpox virus 2 vDUSP11 (SWPV), shearwaterpox virus 2 vDUSP11 catalytic mutant (SWPV-CM), cheloniidpox virus vDUSP11 (ChePV), cheloniidpox virus vDUSP11 catalytic mutant (ChePV-CM), or protein phosphatase DUSP12 (D12) infected with GFP-M51R VSV at 48 hours post-infection (hpi). Cells were infected with GFP-M51R VSV at an MOI of 0.01 PFU/cell. (B). VSV viral titer of indicated A549 DUSP11 KO reconstituted cells at 48 hours post-infection (hpi). Cells were infected with GFP-M51R VSV at an MOI of 0.01 PFU/cell and virus supernatant was collected for plaque assay analysis. (C). Representative images of plaque assay analysis of virus supernatant at 48 hours post-infection. Data are derived from n = 3 independent replicates. Data are presented as mean ± SEM.

### vDUSP11s alter steady-state levels of select RNA polymerase III RNAs

Previous work from our lab demonstrated that DUSP11 modulates the 5’-triphosphate status and steady-state levels of several RNAP III transcripts, including vault RNAs (vtRNAs), Y RNAs, and RMRP (12). This activity could be advantageous to viruses as RNAP III transcripts can activate host PRRs such as RIG-I during virus infection (8,9). Based on this, we sought to determine if the presence of vDUSP11 can impact the steady-state levels of endogenous RNAP III transcripts.

To evaluate if vDUSP11 catalytic activity can alter the abundance of endogenous RNAPIII transcripts, we harvested total RNA from A549 DUSP11 KO reconstituted cells and conducted Northern blot analysis (Fig 6A). We probed for vtRNA1-1 and vtRNA2-1, using the 5’ monophosphate tRNA cysteine (tRNA-Cys) as a loading control. In the presence of either hDUSP11 or either vDUSP11, we observed ∼30-70% reduced levels of vtRNA1-1 and vtRNA2-1 when compared to the empty vector negative control (Fig 6B-C and Fig S6). These data suggest that vDUSP11s can alter vtRNA levels comparable to hDUSP11.

**Figure 6:**
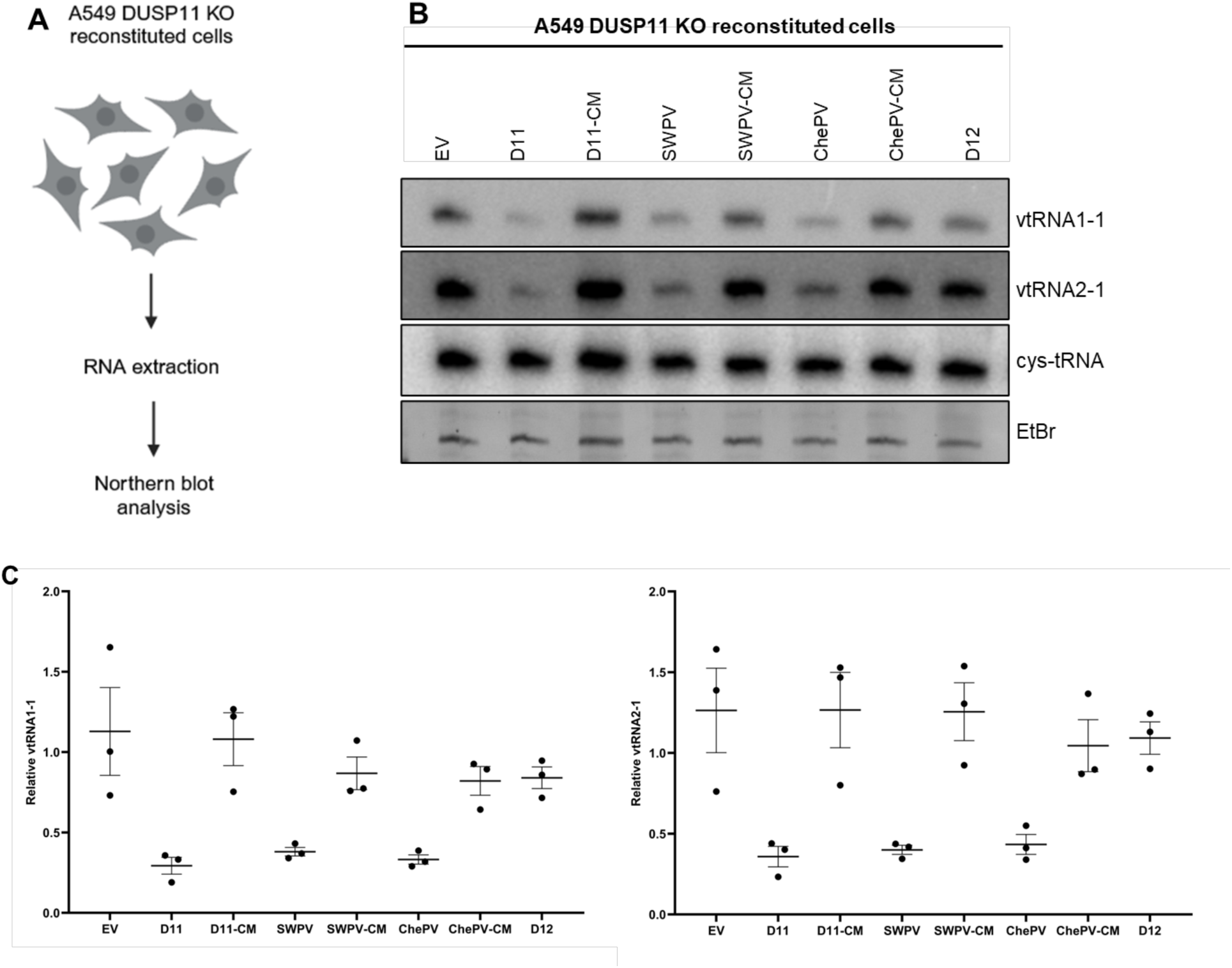
vDUSP11 modulates steady-state RNA levels of endogenous RNAP III transcripts. (A) Schematic diagram of vtRNA Northern blot. RNA was collected from resting A549 DUSP11 KO cells stably reconstituted with empty vector (EV) plasmid, hDUSP11 (D11), hDUSP11 catalytic mutant (D11-CM), shearwaterpox virus 2 vDUSP11 (SWPV), shearwaterpox virus 2 vDUSP11 catalytic mutant (SWPV-CM), cheloniidpox virus vDUSP11 (ChePV), cheloniidpox virus vDUSP11 catalytic mutant (ChePV-CM), or negative control protein phosphatase DUSP12 (D12). Purified RNA was then subjected to Northern blot analysis. (B) Northern blot analysis of vtRNA1-1 and vtRNA2-1 using RNA from A549 DUSP11 KO reconstituted cells. (C) Graphical representation of relative band intensity of vtRNA1-1 and vtRNA2-1 normalized to the relative band intensity of the 5’ monophosphate control cysteine-tRNA. Data are derived from n =3 independent replicates. In all panels, data are represented as mean ± SEM.

### vDUSP11s exhibit pan-cellular localization

RIG-I is predominantly localized to the cytoplasm (31), but RNAP III RNAs are typically distributed in both the cytoplasm and nucleus. In resting cells, hDUSP11 has been shown to predominantly reside in the nucleus (32). However, in infected cells, DUSP11 must also be active in the cytoplasm, as both HCV and VSV transcripts are direct substrates that are restricted to the cytosol (13,14). Poxviruses are DNA viruses, but unlike the majority of DNA viruses, they replicate in the cytoplasm. We sought to determine how vDUSP11 localization compares to hDUSP11.

We performed confocal immunofluorescence microscopy using the various hDUSP11 and vDUSP11 reconstituted A549 DUSP11 KO cell lines. As expected, resting cells expressing hDUSP11 and hDUSP11-CM displayed predominantly nuclear localization (Fig 7). In contrast, both vDUSP11s and their catalytic mutants appear to be pan-cellular with substantial abundance in the cytoplasm (Fig 7). The localization of these vDUSP11s is consistent with overlapping the site of poxviral replication and replicative intermediates.

**Figure 7:**
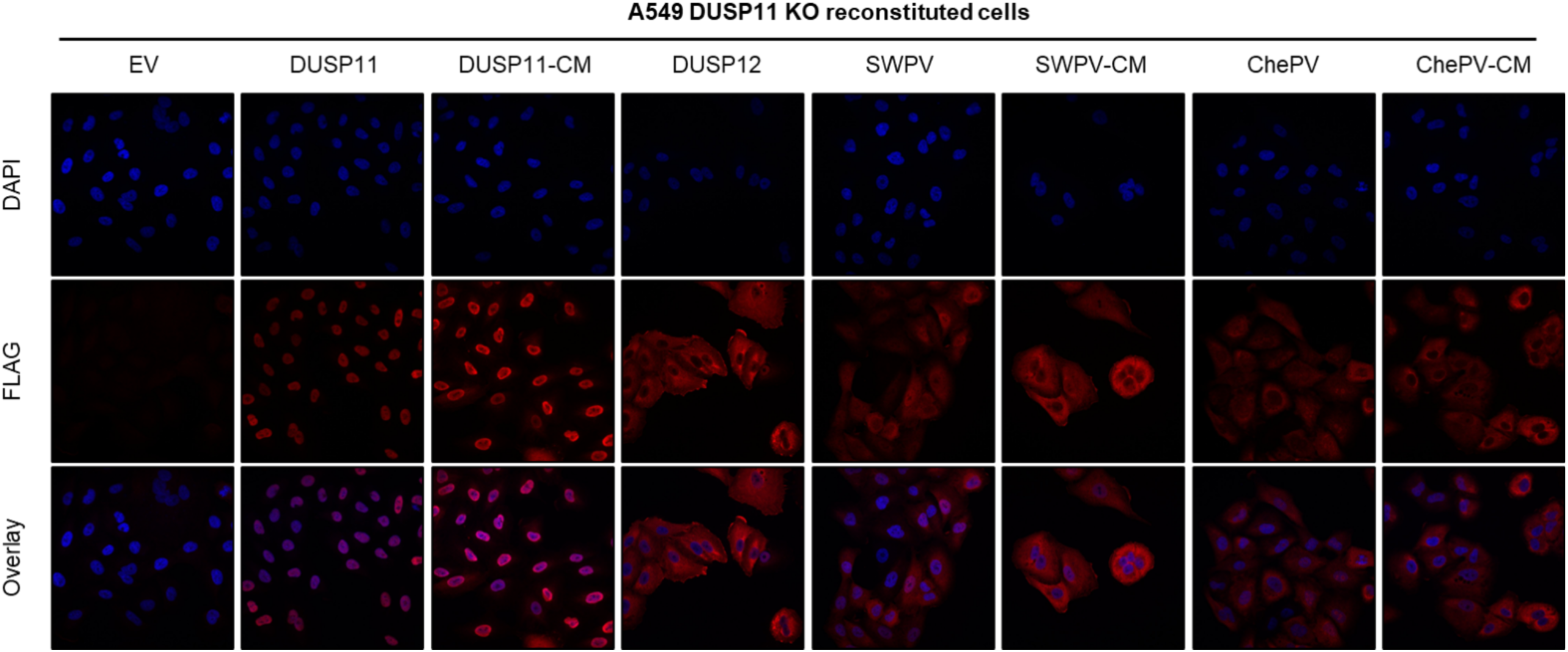
vDUSP11s differ in subcellular localization compared to DUSP11. Representative confocal images of A549 DUSP11 KO cells stably expressing either an empty vector (EV) plasmid or individual 3xFLAG tagged proteins including hDUSP11 (D11), hDUSP11 catalytic mutant (D11-CM), SWPV2 vDUSP11 (SWPV), SWPV2-CM (SWPV-CM), ChePV1 vDUSP11 (ChePV), ChePV1 vDUSP11-CM (ChePV-CM), or negative control protein phosphatase DUSP12 (D12). Prolong™ Gold Antifade Mountant (Thermo Fisher Scientific) was used in slide preparation to visualize nuclei.

### Synteny analysis supports a single avipox vDUSP11 acquisition event from an ancestral host and is consistent with rapid evolutionary pressure

Viral piracy of host proteins often occurs when genes have important immunological functions (2). Due to their role at the host-pathogen interface, pirated genes are often subject to rapid evolution, which can include frequent duplications and loss (2,33). For poxviruses in particular, genes that inhibit host defense often cluster together within the genome and often appear in regions towards the ends of the genome (34–36). Comparing the genomic localization of these vDUSP11s can also inform the evolutionary history of vDUSP11 including whether these genes were acquired from the host multiple times or through a single event.

To assess the acquisition of vDUSP11 into the various APV/AdjPV genomes, we analyzed the genomic location and adjacent genes. Using reference genome sequences in the NCBI database (37) for each APV/AdjPV virus containing a vDUSP11, we analyzed genes upstream and downstream of the vDUSP11 coding region (Fig 8 & Table S6). We found that while vDUSP11 is not encoded in the inverted terminal repeats (ITRs), in many cases, it is surrounded by other genes predicted to be involved in modulating the host response to infection (Fig 8 & Table S6). Due to the similarity in the order of flanking genes, these data suggest that vDUSP11s are orthologous and likely acquired by a single acquisition event.

**Figure 8:**
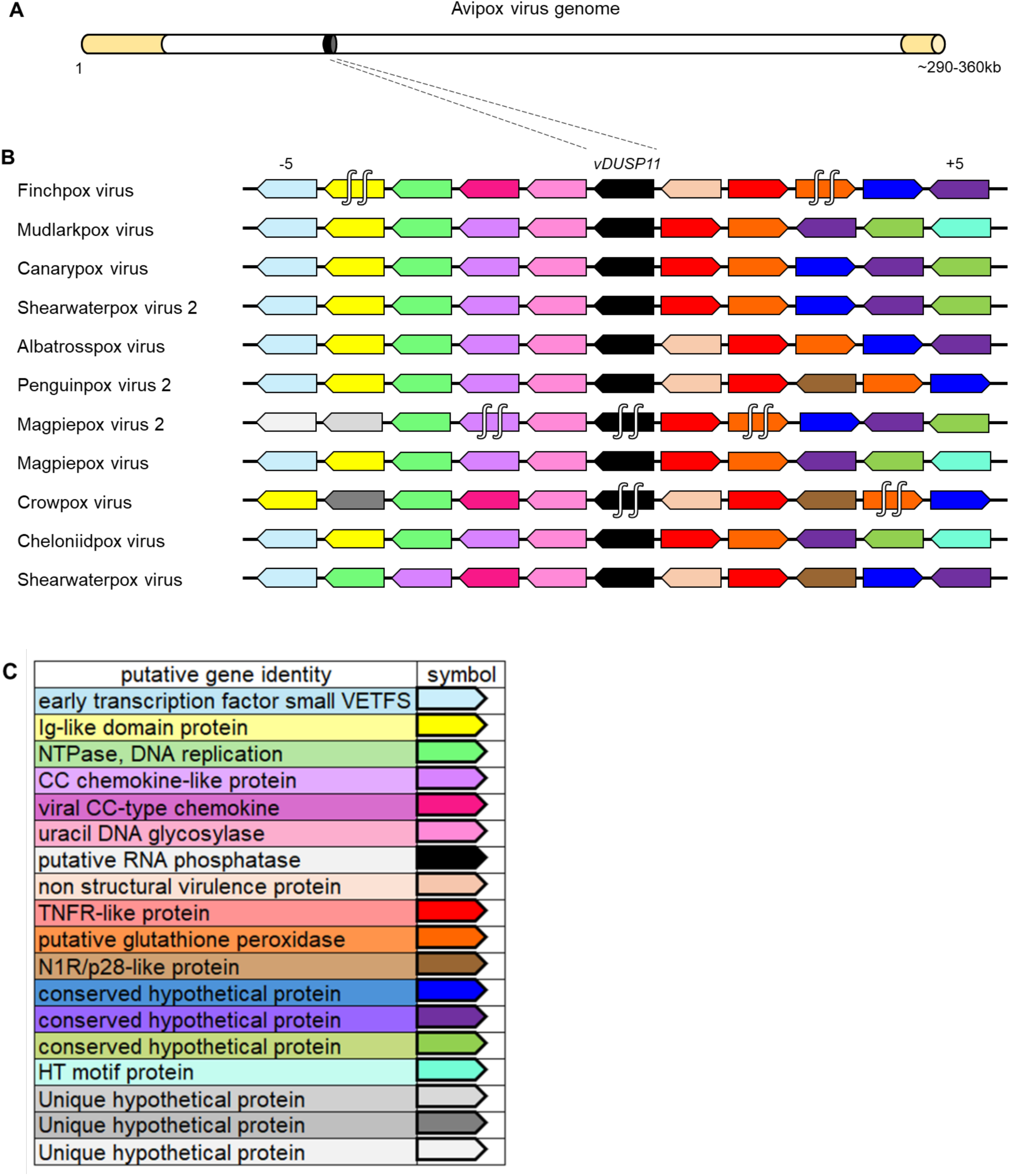
A single acquisition event for vDUSP11 is indicated by synteny analysis. (A) Graphical representation of avipoxvirus genome using published information from the canarypox virus genome as a model (58). Inverted terminal repeats (ITRs) are indicated in yellow, relative genome position of *vDUSP11* indicated in black. (B) Analysis of upstream and downstream flanking genes indicated *vDUSP11*s (black) reside at similar genomic locations across poxvirus genomes. Protein homologs were determined using reciprocal BLAST hits for the 5 genes upstream (left of *vDUSP11*) and 5 genes downstream (right of *vDUSP11*) of *vDUSP11*. Homologous genes are indicated by shared color. Genes oriented to the right indicated ORFs on the top strand while genes oriented to the left indicated genes on the bottom strand. (C) Corresponding identity for genes indicated in (B). Putative gene identity was determined using NCBI database genome annotations (37). For genes lacking descriptive annotations, more detailed annotations based on identified homologous genes were used (Table S6). 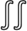 indicates *vDUSP11* ORFs disrupted by stop codons, resulting in multiple separate coding regions.

During this analysis, we noted that the vDUSP11 loci for crowpox virus and magpiepox virus 2 are substantially shorter. The coding region for the magpiepox virus 2 vDUSP11 encodes a predicted 60 AA protein retaining key residues essential for catalytic activity. However, it is unclear if this is sufficient for functionality. In the case of crowpox virus, comparing the genomic location to other closely related viruses showed a mutated start codon resulting in a protein not predicted to retain catalytic activity. The presence of these truncated forms is consistent with this locus being a site of rapid evolutionary pressure. Thus, these evolutionary profiles are consistent with what would be expected if vDUSP11 activity is involved in modulating host defenses.

Our findings support the model of a single acquisition event of a host DUSP11 in an ancestor virus that gave rise to the modern lineage of avipox and avipox-adjacent viruses with vDUSP11s. To decipher a potential source of acquisition, we performed reciprocal BLASTp analysis for each viral homolog to identify the most similar host DUSP11. Based on both BLASTp E-value and percent identity, we identified the most similar host DUSP11 to be from *Corvus hawaiiensis* (Hawaiian crow) to the penguinpox 2 vDUSP11 [53% (94/176) amino acid identity, 71% positives (126/176)] (Table S7). Furthermore, this host is recalled for all the full-length vDUSP11, except for mudlarkpox virus for which *Varanus komodoensis*, (Komodo dragon) was recalled [51% (96/189) amino acid identity, 69% positives (131/189)] (Table S8). This further supports the likelihood that the vDUSP11s were acquired from an avian species and potentially one closely related to *Corvus hawaiiensis*.

To further evaluate the acquisition source of APV/AdjPV vDUSP11, we performed phylogenetic analysis on the avipox vDUSP11 and selected host DUSP11 AA sequences (Table S10). Using PhyML (38) to generate an inferred phylogenetic tree, we found that the avipox vDUSP11 clusters with avian DUSP11 (Fig S2). These data lend additional support for an avian origin for the various avipox vDUSP11. Combined, the above data demonstrate that APV/AdjPVs have acquired a viral homolog of host RNA phosphatase DUSP11, which is enzymatically active and features characteristics of a protein involved in the modulation of the host immune response.

## Discussion

Historically, unraveling the mechanisms of how viruses replicate has allowed for a greater understanding of not only viruses but also of how cells function. In addition, viral piracy of host genes can implicate that a host factor and/or its activity influences infection outcomes (2). A detailed understanding and characterization of viral replication strategies and immune pathways is essential to developing new therapies and applications. Our studies of vDUSP11s not only shed light on how APVs/AdjPVs deal with barriers to viral replication, but it also supports the role of DUSP11 in modulating the antiviral response.

As previously published, knockout of cellular hDUSP11 leads to increased sensitivity to RIG-I-related immune activation (14). Herein, we also find that that SWPV2- and ChePV-encoded vDUSP11s can modulate immune activation in response to liposomal 5’-triphosphate RNAs (Fig 4). The functional relevance of vDUSP11 activity is illustrated by these factors being pro-viral in the context of VSV infection, as previously seen for hDUSP11 (Fig 5). These observations demonstrate similar functionalities of host and viral DUSP11 and are consistent with a possible role of vDUSP11 in modulating RIG-I during poxvirus infection. However, due to the current lack of a facile model system for these APVs/AdjPVs, this model has not been tested under native infection.

It is established that for some viruses, viral RNA produced during infection can have immunostimulatory properties, in part, by activating host PRRs (4,6). However, there is a growing pool of evidence demonstrating that endogenous RNAs may also be relevant activators of RIG-I during virus infection (8,9). Here, we show that vDUSP11 can modulate steady-state levels of endogenous RNAP III transcripts (Fig 6). It remains a possibility that the ability of vDUSP11 to modulate endogenous RNAs is not merely a side effect but instead an intentional consequence to prevent these RNAs from acting as DAMPs/PAMPs during APV/AdjPV infection.

5’-triphosphate RNA can be a replicative intermediate of RNA virus replication, but the mechanism for how DNA virus infection enhances levels of total (host and viral) 5’-triphosphate RNA is less clear. Endogenous RNAP III transcripts are a potential RIG-I agonist (8–10), which could explain the source for APV/AdjPV infection. Another proposed model for the source of 5’-triphosphate RNA during DNA virus infection is RNAP III translocation to the cytoplasm, which then generates immunostimulatory transcripts from foreign DNA templates (18,39). These models are not mutually exclusive, and both suggest a role for host RNAP III in activating the immune response during poxvirus infection.

During virus infection, the site within the cell where viral replication occurs dictates the availability of cellular resources and which host defense factors are present. DNA viruses commonly replicate in the nucleus to allow access to host replication machinery. Atypically, poxviruses are large DNA viruses that replicate in the cytoplasm, as they have large genomes that encode their own replication machinery (40). It follows that via immunofluorescence, we observed vDUSP11 to be pan-cellular with substantial presence in the cytosol, having different localization compared to endogenous DUSP11 (Fig 7). Despite this altered cellular distribution, vDUSP11 and host DUSP11 share a comparable ability to modulate the abundance of host RNAPIII transcripts. This further supports a model where endogenous RNAs can function to activate host PRRs.

Noncoding RNA has been shown in well-studied poxviruses, such as Vaccinia virus, to be at the interface of host-pathogen conflict (e.g., SAMD9/9L) (41). Accordingly, Vaccinia virus is known to activate RIG-I (42). Vaccinia virus encodes a protein E3 (encoded by the gene *E3L*) that, among other functions, is involved in reducing RIG-I activation. APV/AdjPV do not encode E3, and Vaccinia and other closely related poxviruses do not encode vDUSP11. Given its essential functions, the absence of E3 in APVs likely necessitates the presence of additional proteins involved in immune modulation, such as vDUSP11.

In summary, we have identified a viral homolog of the host protein DUSP11 that has been co-opted by APVs/AdjPVs. Future comparative studies with vDUSP11 may inform the role of cellular DUSP11 in shaping antiviral responses including functionally relevant amino acid residues. While the function of this vDUSP11 during native APV/AdjPV infection has yet to be elucidated, vDUSP11s have conserved functions in modulating the immune response, as seen with host DUSP11. This, along with the propensity for immune-related proteins to be pirated, suggests the role of vDUSP11 in modulating the innate immune response.

## Materials and Methods

### Identification of avipoxvirus vDUSP11

Viral homologs to DUSP11 were identified using BLASTp (43) with the hDUSP11 sequence as query (Table S2). Sequences were selected based on returned BLAST parameters (total score, e-value, and percent identity). Sequence alignment was generated using FASTA AA sequences acquired from NCBI database (37) and MUSCLE (44) was used for multisequence alignment. Sequence alignment was plotted using the R package ggmsa (45).

### Plasmids

The pcDNA3.1 puro and pLenti DUSP11-3xFLAG and DUSP11-3xFLAG-CM expression vectors were previously described (12). The PISK-T7-HCV5’UTR vector was previously described (13). vDUSP11 reference AA sequences were inputted into the IDT (Integrated DNA technologies) codon optimization tool to generate codon-optimized DNA sequences for expression in human cells. DNA sequences for SWPV2 vDUSP11, ChePV vDUSP11 and DUSP12 were synthesized as gBlocks (IDT) containing XhoI/XbaI restriction enzyme sites. The gBlocks were cloned into pcDNA3.1+ (puro) and plenti-EF1α (pLenti) backbones. Around-the-horn PCR(46) was utilized to add amino-terminal epitope (3xFLAG) tags to pLenti vDUSP11 and DUSP12 constructs. vDUSP11 catalytic mutant (CM) constructs were generated using overlap PCR(47) combined with restriction enzyme digest. Tagged pcDNA3.1+ (puro) 3xFLAG-tagged vDUSP11/vDUSP11-CM constructs were subcloned into pcDNA3.1+ behind a T7 promoter using restriction enzyme digest. Clones were confirmed using whole plasmid sequencing (Plasmidsaurus).

### Cell lines and viruses

A549, BHK-21 and Vero cells were obtained from ATCC. A549 DUSP11 CRISPR–Cas9 targeted cell lines were previously described (12). Cells were maintained in DMEM supplemented with 10% (v/v) fetal bovine serum (Corning) and 1% (v/v) penicillin-streptomycin (Corning). Generation of lentiviral particles, transduction and selection with blasticidin were performed as previously described (48). A549 DUSP11 KO reconstituted cell lines were generated via transduction of lentiviral particles generated from the following lentiviral vectors: pLenti empty-vector (EV), pLenti DUSP11-3xFLAG, DUSP11 CM-3xFLAG, pLenti SWPV2 vDUSP11-3xFLAG, pLenti SWPV2 vDUSP11 CM-3xFLAG, pLenti ChePV vDUSP11-3xFLAG, pLenti ChePV vDUSP11 CM-3xFLAG, or pLenti DUSP12-3xFLAG as a negative control. Mutant VSV (rM51R-M-GFP) was provided by Douglas Lyles (Wake Forest School of Medicine, NC) and has been described previously (30). Stocks were grown in BHK-21 cells and infectious virus concentration was determined using Vero cells.

### Phylogenetic analysis

DUSP11 AA sequences and related information were retrieved from Uniprot.org (ID: O75319). Accession numbers and related information for other dual-specificity phosphatases for human, and the homologs in mouse, and hawaiian crow were determined using Uniprot.org and BLASTp (43) for confirmation (Table S5). AA sequences were retrieved from NCBI (37). A multiple sequence alignment (MSA) of AA sequences were performed using Clustal Omega standard parameters (Fig S3) (49). Phylogenetic analysis was performed using PhyML, with bootstrap replicates set to 100 (38). Model selection was performed by Smart Model Selection (SMS) integrated into PhyML (50). FigTree v1.4.4 (http://tree.bio.ed.ac.uk/software/figtree/) was used for visualization. Additional phylogenetic analysis was performed using the same parameters for host and avipox viral DUSP11 sequences (Table S10). MSA of AA sequences was performed using Clustal Omega standard parameters (Fig S4) (49).

### Protein modeling

A partial crystal structure of DUSP11 was obtained from wwPDB.org (PDB: 4JMJ) (25). Structural predictions of the SWPV2 vDUSP11 were generated for the corresponding amino acids (9–190) using AlphaFold2 using standard parameters and the rank 1 structure was used for analysis (24). UCSF chimera was used for visualization mapping, and analysis (51).

### Synteny analysis

NCBI was used to identify genomic locations for viral DUSP11 sequences within the viral genomes (37). Accession numbers for adjacent coding portions corresponding to the 5 genes upstream and downstream from each viral DUSP11 were recorded (Table S6). Reciprocal BLAST hits (RBH) were performed for coding sequences of vDUSP11-adjacent genes across virus species to identify homologs (Table S6) (43). Genomic database annotations and the NCBI database were used for gene identities and homology (37). CoreGenes 5.0 was used to further verify the presence of homologous genes between selected poxvirus genomes (52).

### Virus infections and plaque assays

A549 DUSP11 KO cells reconstituted with various DUSP11/vDUSP11s were plated at a density of 0.1 x 10^6^ cells per well in a 12-well plate for virus infection. After 1 hour of virus adsorption with gentle shaking at 15-minute intervals, virus inoculum was aspirated from the wells and washed gently with PBS before adding growth medium. GFP images were taken at indicated time points, one representative image is shown with replicates shown in Figure S10. 100 μL of media was collected at select time points post infection and stored in −80°C to determine the yield of infectious virus. Virus concentration was quantified by standard plaque assay on Vero cells in 6-well plate format, serially diluted in a countable range (5-250 plaques per well). 72 hours post-carboxymethyl cellulose (2.5%) overlay, plates were fixed and stained with 0.25% Coomassie blue in 10% acetic acid, 45% methanol or 0.5% methylene blue in 50% methanol.

### Northern blot analysis

Northern blot analysis was performed as previously described (53). In brief, total RNA was harvested from cells using TRIzol reagent (Thermo Fisher) or PIG-B (53,54) and fractionated on 10-12% polyacrylamide gel electrophoresis (PAGE)-urea gel and transferred to Amersham Hybond-N+ membrane (GE Healthcare). Transferred membranes were crosslinked 2x with a UV crosslinker (UVP) with the RNA side of the membrane facing the UV source at 1,200 μJ/m^2^. Crosslinked membranes were prehybridized for ∼1 hour at 55C in hybridization solution (Takara). DNA oligos (Table S1) were radiolabeled with [γ-^32^P] ATP (Perkin Elmer) by T4 polynucleotide kinase (New England Biolabs) for ∼1 hour at 37C. Radiolabeled probes were added to membranes following prehybridization and hybridized while rotating in UVP hybridization oven (HL-2000 HybriLinker UVP) overnight at 38C. Probed membranes were exposed to a phosphor screen and visualized using Typhoon Biomolecular Imager (GE Healthcare). Membranes were stripped by incubating membrane with boiled 0.1% SDS with agitation for 15 minutes and repeated three times. Replicate northern blots shown in Figure S7.

### In vitro transcription

To generate 5’-triphosphorylated HCV 5’ UTR RNA used for liposomal delivery and the *in vitro* phosphatase assays, *in vitro* transcription was performed with the AmpliScribe T7-Flash Transcription Kit (Bioresearch™ Technologies) according to manufacturer’s instructions. DNA templates for the *in vitro* transcription reaction were previously described (13). RNA was then purified using MicroSpin G-25 columns [GE Healthcare].

### In vitro phosphatase assays

*In vitro* translated DUSP11, DUSP11-CM, SWPV2 vDUSP11, SWPV2 vDUSP11-CM lysates were generated using the TnT quick-coupled transcription/translation system (Promega) and the pcDNA3.1 DUSP11-3xFLAG, pcDNA3.1 DUSP11-CM-3xFLAG, pcDNA3.1 SWPV2-vDUSP11-3xFLAG, and pcDNA3.1 SWPV2-vDUSP11-CM-3xFLAG, plasmids as templates. Five micrograms of HCV 5’ UTR RNA was incubated in a 100-μL reaction [50mM Tris, 10mM KCl, 5mM DTT, 50mM EDTA, 40 units SUPERaseIn (Thermo Fisher Scientific)] with 6 μL of in vitro translated products (DUSP11-3xFLAG, DUSP11-CM-3xFLAG, SWPV2-vDUSP11-3xFLAG, SWPV2-vDUSP11-CM-3xFLAG, or a luciferase negative control) for 10 min at 37C. EDTA was omitted from CIP control reactions as it is inhibitory of its enzymatic activity. RNA was purified using PIG-B (54). Purified RNA from each phosphatase or control reaction was then used to set up two reactions 20 μL reactions containing 1 μg of treated RNA in NEBuffer 3.1 (New England Biolabs) with or without the addition of 1 μL (1U) recombinant XRN1 (New England Biolabs) and was then incubated for 60 minutes at 37C as previously described (13). The reactions were fractionated by 7.5% urea/PAGE and were then stained using ethidium bromide (∼1 μg/mL) for 3-5 minutes while rocking. The Bio-Rad Molecular Imager® GelDoc XR™ was then used to visualize gel via exposure to UV. Gel images were processed using Fiji (ImageJ variant) (55,56). Replicate EtBr-stained urea gels shown in Figure S8.

### 5pppRNA transfections

A549 DUSP11 KO reconstituted cells were seeded at a density of 0.1 × 10^6^ cells per well in a 12-well format. 24 hours following plating, cells were transfected with 5-10 ng of *in vitro* transcribed 5’-triphosphorylated HCV 5’ UTR RNA, or with Lipofectamine 2000 (Invitrogen) alone control (3 µL). Transfection was terminated after 18 hours by the addition of either TRIzol reagent (Invitrogen) or PIG-B, followed by RNA extraction and subsequent RT-qPCR analysis.

### Real-time PCR (RT-qPCR)

Total RNA was extracted from cells using TRIzol reagent or PIG-B (54). Extracted RNA was treated with DNase I (Thermo Fisher) followed by ethanol precipitation (100% ethanol, 3M sodium acetate). Precipitated RNA was resuspended in nuclease-free water (Thermo Fisher) and was quantified using the NanoDrop® ND-1000 Spectrophotometer. RNA purity was assessed using the measured OD at 260/280 nm, with RNA considered acceptable having values within ∼1.7-2.1. Complementary DNA (cDNA) was synthesized using SuperScript III reverse transcriptase (Invitrogen) using equal amounts of RNA. All RT-qPCR quantification and measurements were performed on a StepOnePlus Real-Time PCR system (Applied BioSystems). A standard 3-step PCR cycling protocol (95°C for 15 sec, 60°C for 15 sec, 68°C for 30 sec, 40 cycles) using PerfeCTa SYBR Green FastMix (Quantabio) was used according to the manufacturer’s instructions. At least two technical replicates per sample/condition. Gene expression of human IFNB1 and ISG15 were normalized to the expression of GAPDH (Table S1). GAPDH values for A549 DUSP11 KO + EV cells transfected with L2K alone were used when defining the experiment negative control within the Applied BioSystems program (14).

### Confocal Images

A549 DUSP11 KO reconstituted lines were plated at 0.5 × 10^5^ cells per well in a 24-well plate containing a 12-mm glass coverslip. The next day cells were washed three times with 1X PBS before fixation with 2% paraformaldehyde for 20 minutes. Paraformaldehyde was removed and cells were washed three times with 1X PBS. Following washes, cells were permeabilized with 0.5% Triton X-100 in 1X PBS 3% BSA for 2 minutes at room temperature. Next, the permeabilization buffer was removed and cells were incubated in blocking buffer (0.2% Triton X-100, 3% BSA, 1X PBS) while rocking for 1 hour at room temperature. Following incubation, M2 FLAG antibody (Sigma Aldrich #F1804) was diluted to 1:1000 in blocking buffer and incubated overnight at 4C. The next day cells were subjected to three 5-minute washes in blocking buffer. Secondary antibodies (Table S1) were diluted 1:1000 in blocking buffer and added to cells to incubate for 30 minutes at room temperature in the dark. Cells were washed three times in blocking buffer for 5 minutes each. ProLong Diamond Antifade Mountant (Invitrogen #P36961) was used when mounting coverslips on slides. Cells were imaged on ZEISS confocal using the 40x objective (1 representative image shown). Images were processed to final form using ZEISS image analysis software and Image J.

### Immunoblot analysis

Protein was extracted from cells using either RIPA lysis buffer (50 mM Tris-HCl, 150 mM NaCl, 0.25% sodium-deoxycholate, 1% NP-40, 0.1% SDS with protease inhibitor) supplemented with Protease and Phosphatase Inhibitor Cocktail (Abcam, ab201119) or SDS page sample buffer (1X) (57). Cell lysate was fractionated on an 8% SDS-PAGE and transferred to Amersham Protran 0.45 μm nitrocellulose membrane (GE Healthcare). Blots were incubated in blocking buffer (Intercept® PBS blocking buffer, LI-COR)) then probed with primary antibodies for M2 FLAG (monoclonal, 1:2000 dilution, Sigma Aldrich, F1804) and beta-Tubulin (polyclonal, 1:2000 dilution, Cell Signaling, 2146S) in blocking buffer overnight at 4C. Blots were washed with ∼10 mL 1X PBST three times for 5 minutes prior to incubation with secondary antibodies. IRDye 800CW (1:10,000 dilution, LI-COR, 926-32213) and IRDye 680LT (1:10,000 dilution, LI-COR, 926-68022) were used as secondary antibodies, diluted in blocking buffer. Blots were washed with ∼10mL 1X PBST three times for 5 minutes, then imaged with an Odyssey CLx infrared imaging system (LI-COR). Uncropped western blots shown in Figure S9.

### Statistical analysis

GraphPad Prism software was used for statistical analyses. Error bars were presented as mean ± standard error of the mean.

Schematic figures were Created with BioRender.com

## Author contributions

Conceptualization, KHS, CSS, DCH;

Data curation, KHS;

Formal analysis, KHS;

Funding Acquisition, CSS, DCH;

Investigation, KHS;

Methodology, KHS, CSS, DCH;

Project administration, CSS, DCH;

Supervision, CSS, DCH;

Visualization, KHS;

Writing – Original Draft, KHS;

Writing – Review & Editing, KHS, CSS, DCH.

## Acknowledgements

Justin Lau provided technical expertise including independent validation of experiments in Figures 4, 6 and 8. (shown in Supplemental Figure S5 and S6). Don Gammon, UT Southwestern Medical Center, provided helpful discussions regarding this project.

---

**Figure S1:**
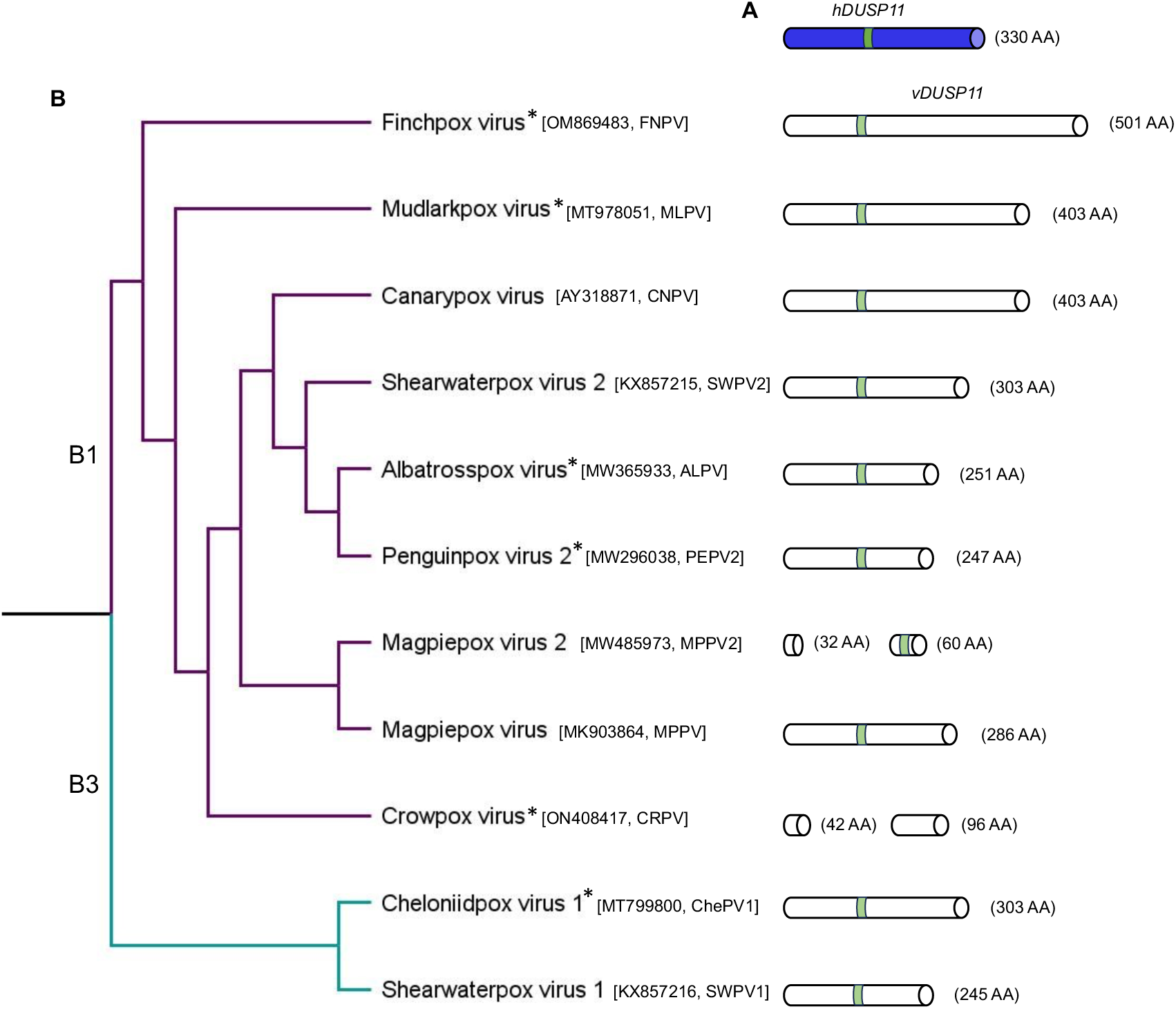
A broad phylogenetic distribution of avipox and related avipox-adjacent poxviruses containing vDUSP11. (A) A graphical representation of *hDUSP11* with AA sequence length. (B) A cladogram built from published phylogenetic data (59), focusing on a subset of poxviruses encoding vDUSP11. Sub-clades (B1 and B3) are designated according to Gyuranecz et al. (2013) (60). Graphical representations of *vDUSP11* are to the right of each virus, demonstrating variations in vDUSP11 AA sequence length between viruses. Magpiepox virus 2 and crowpox virus encode truncated vDUSP11 as indicated. The presence of * indicates unclassified poxviruses. vDUSP11 p-loop indicated in green.

**Figure S2.**
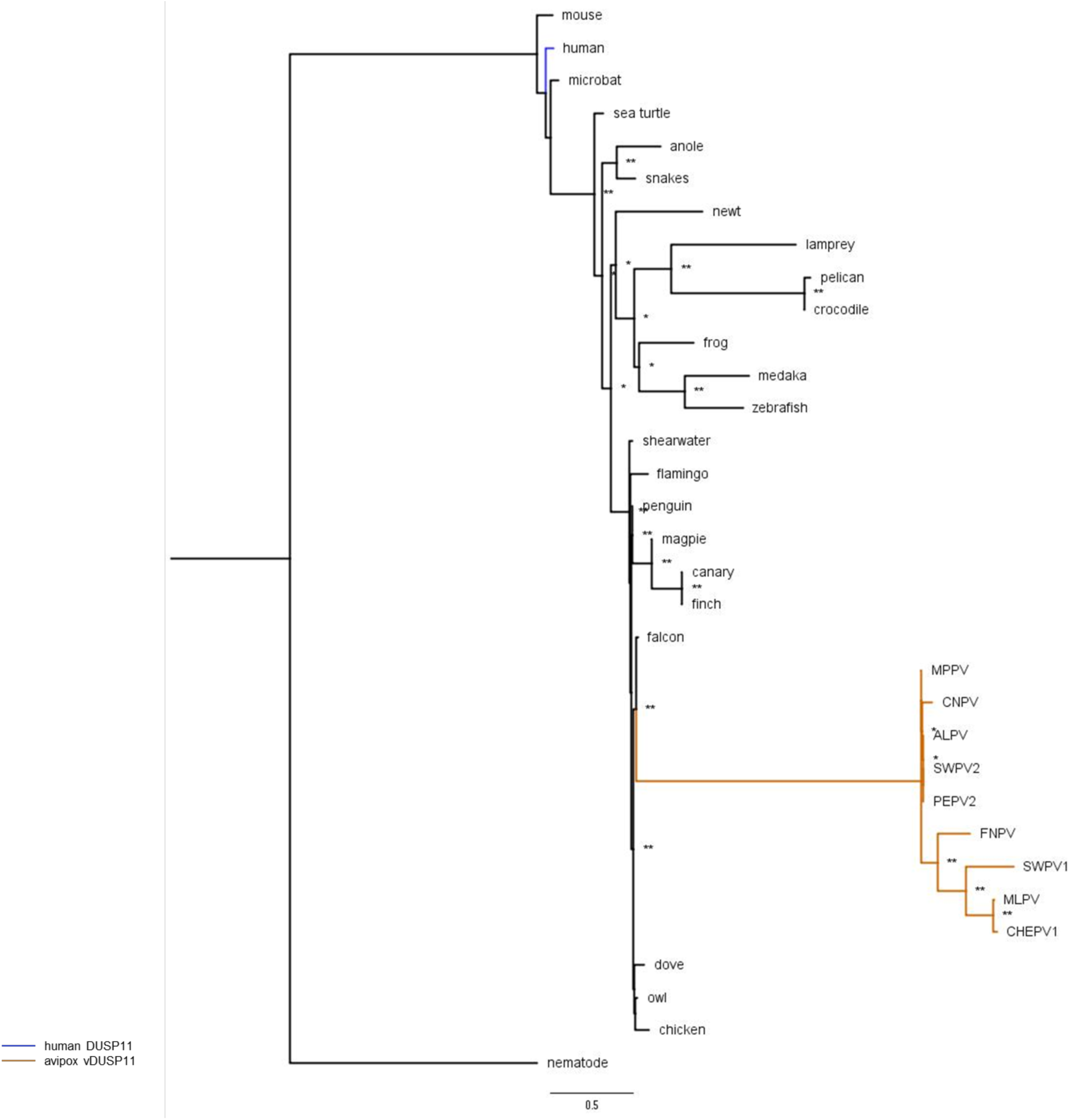
An inferred phylogenetic tree highlighting likely avian origin of APV/AdjPV vDUSP11s. An inferred tree built using 33 host and viral DUSP11 amino acid (AA) sequences by maximum-likelihood analysis phylogenetic tree using PhyML (38). AA sequences for host DUSP11s and APV/AdjPV putative vDUSP11s were aligned using Clustal Omega (Fig S4) (49). Clustal alignment was used to run PhyML analysis with the Q.plant +G+I model selected by SMS (50). Sequences were retrieved from the NCBI sequence database (37) and Uniprot (www.uniprot.org/) (Table S10). Putative APV/AdjPV vDUSP11s (orange) cluster with avian host DUSP11s. 100 bootstrap replicates were performed; branch support >50% (*) or >70% (**) are indicated.

**Figure S3.**
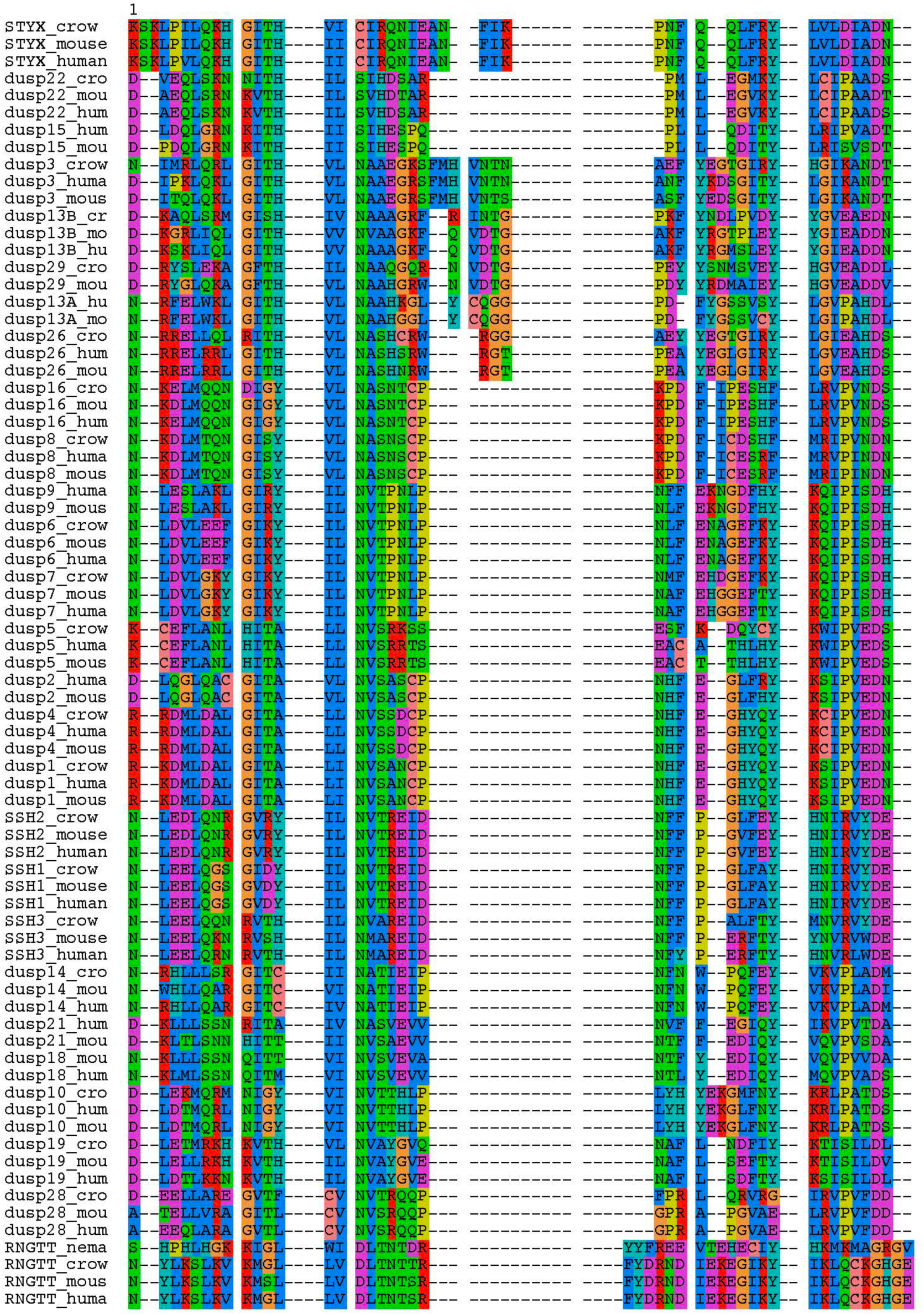
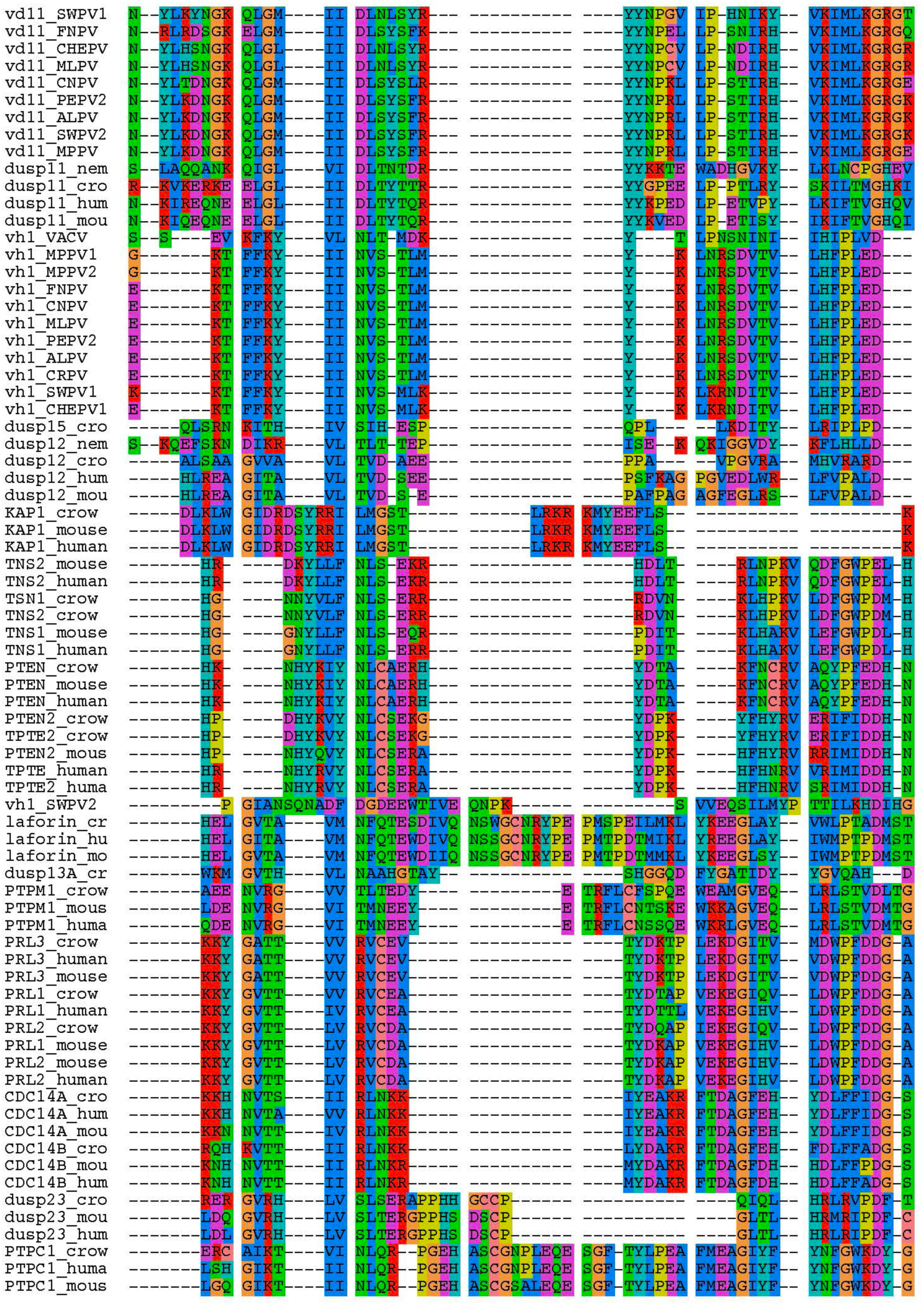
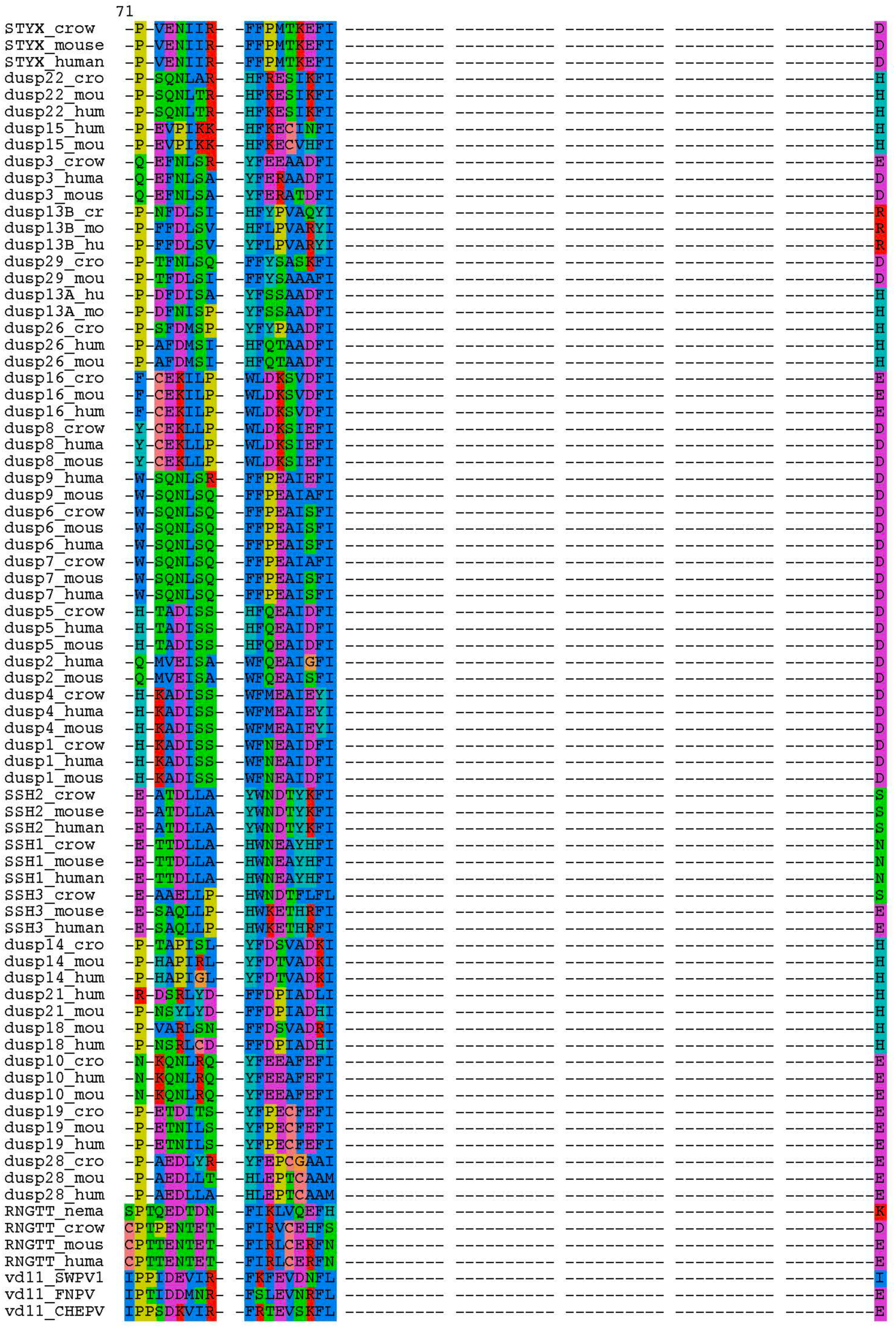
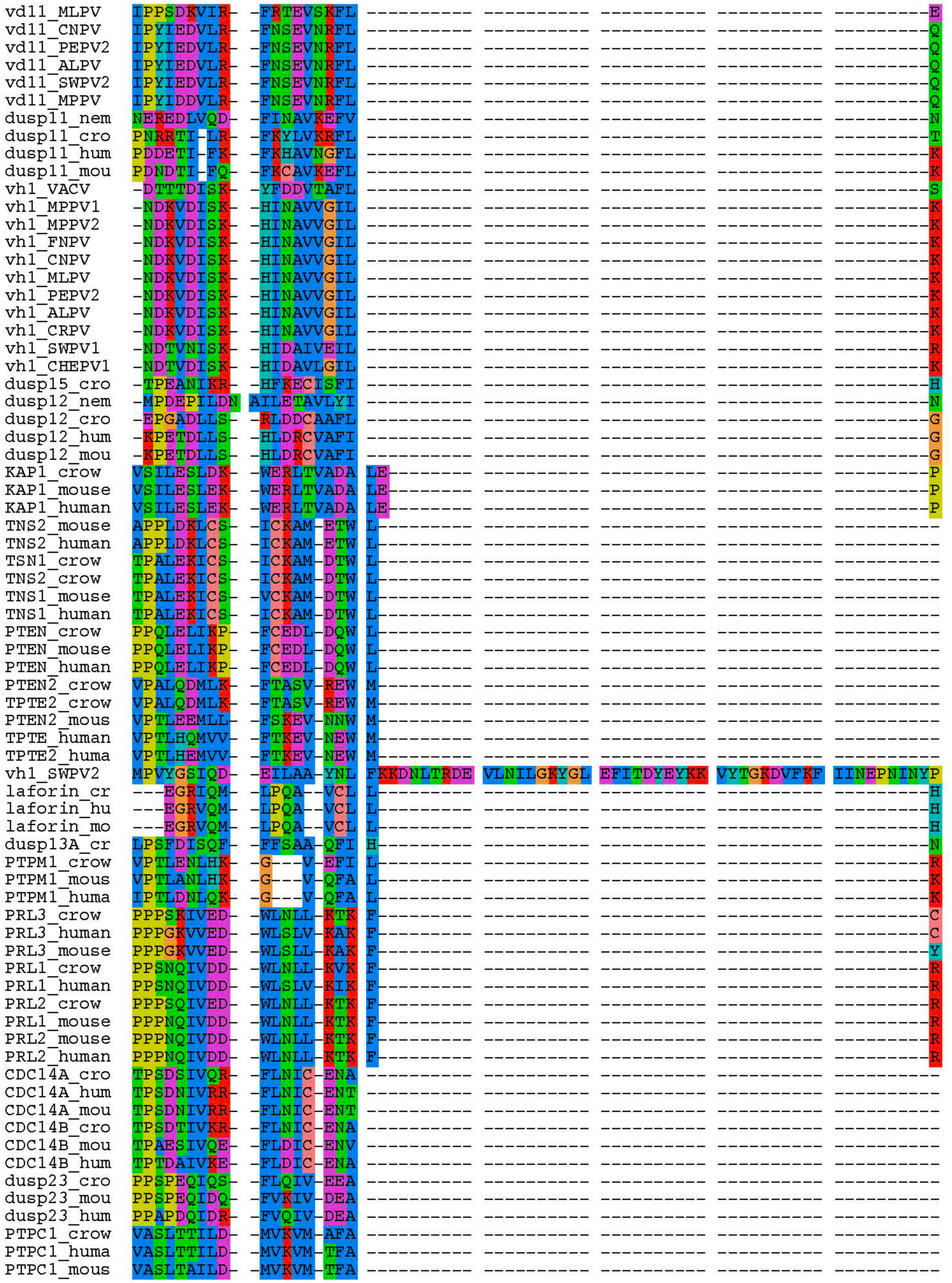
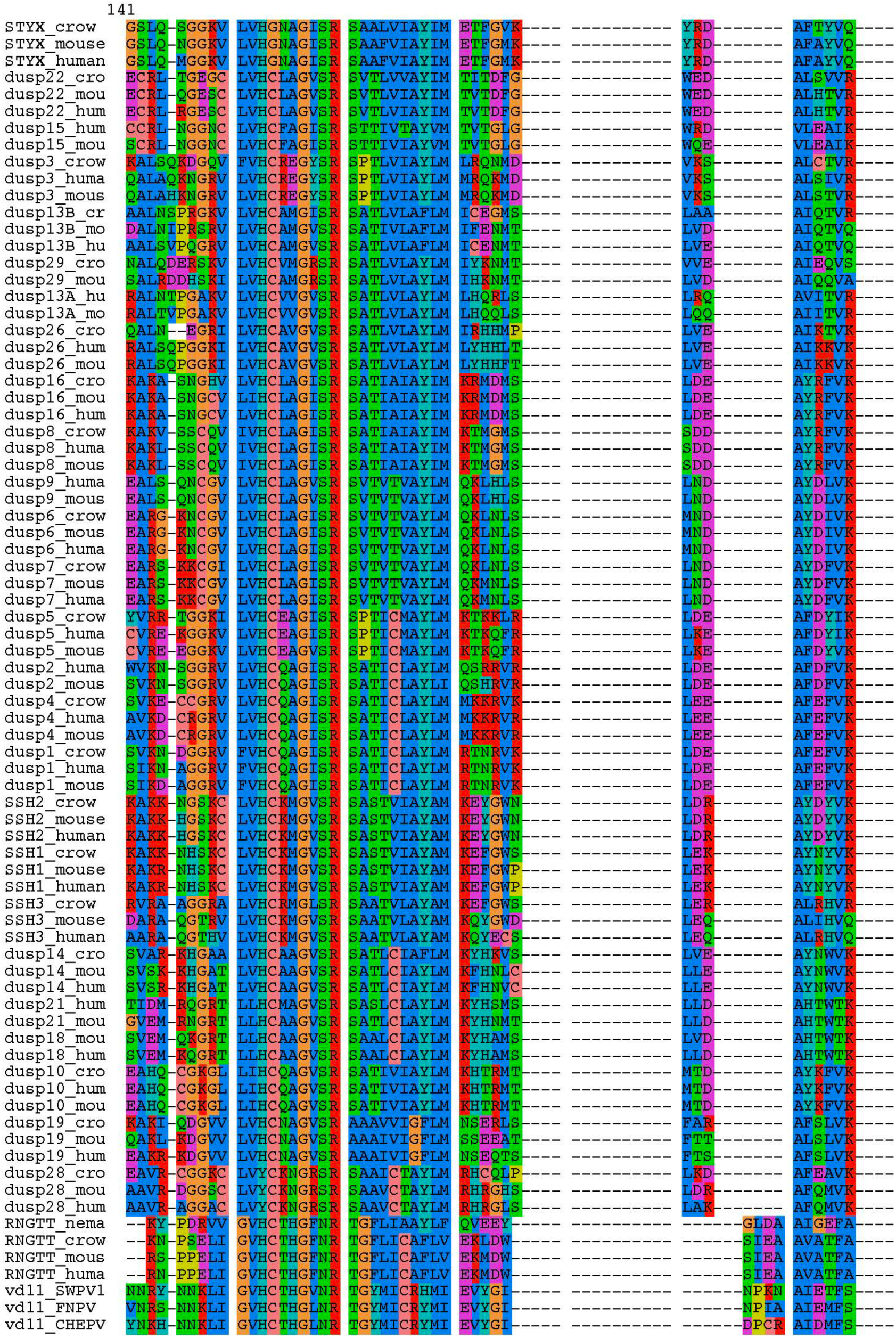
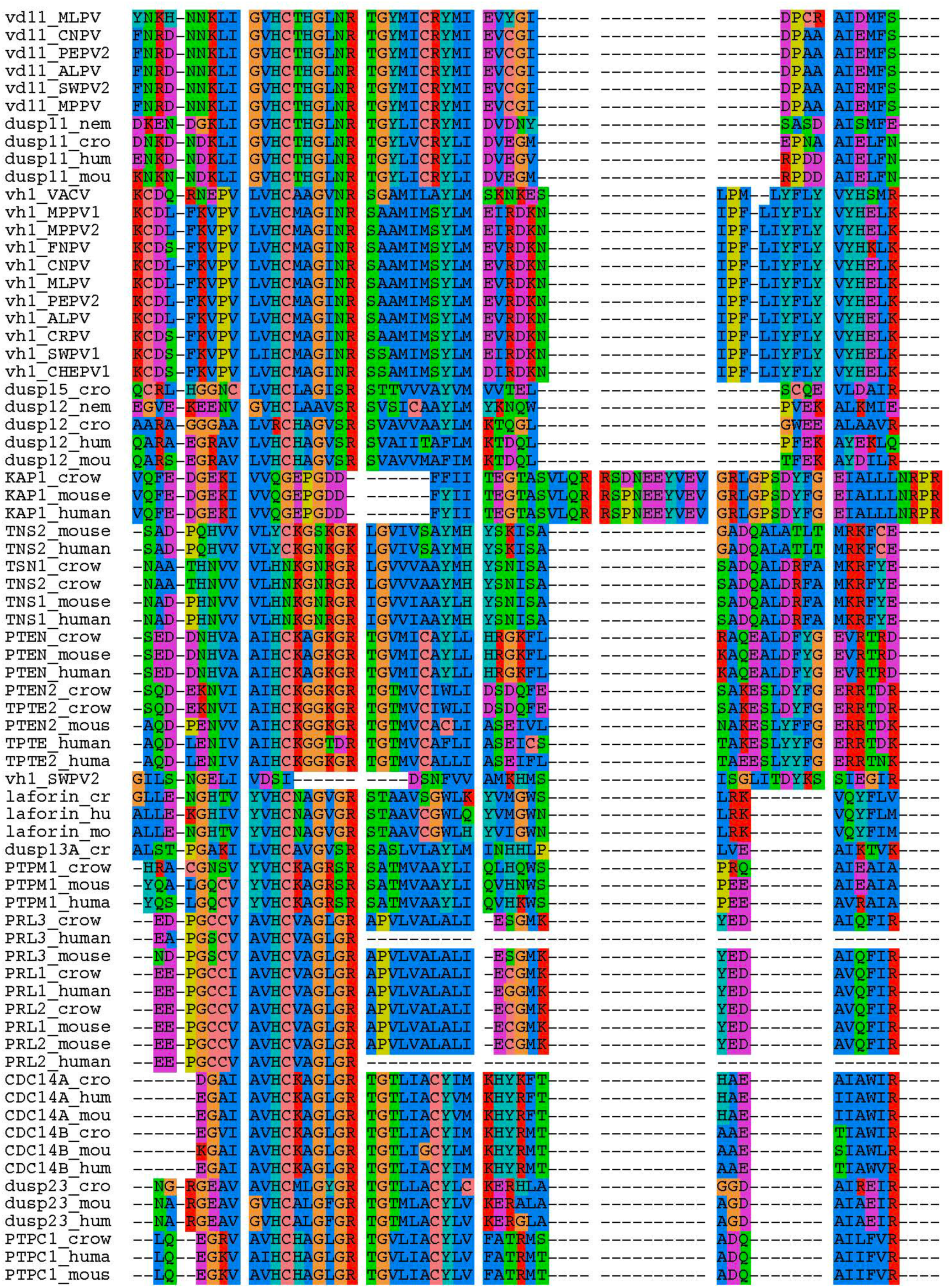
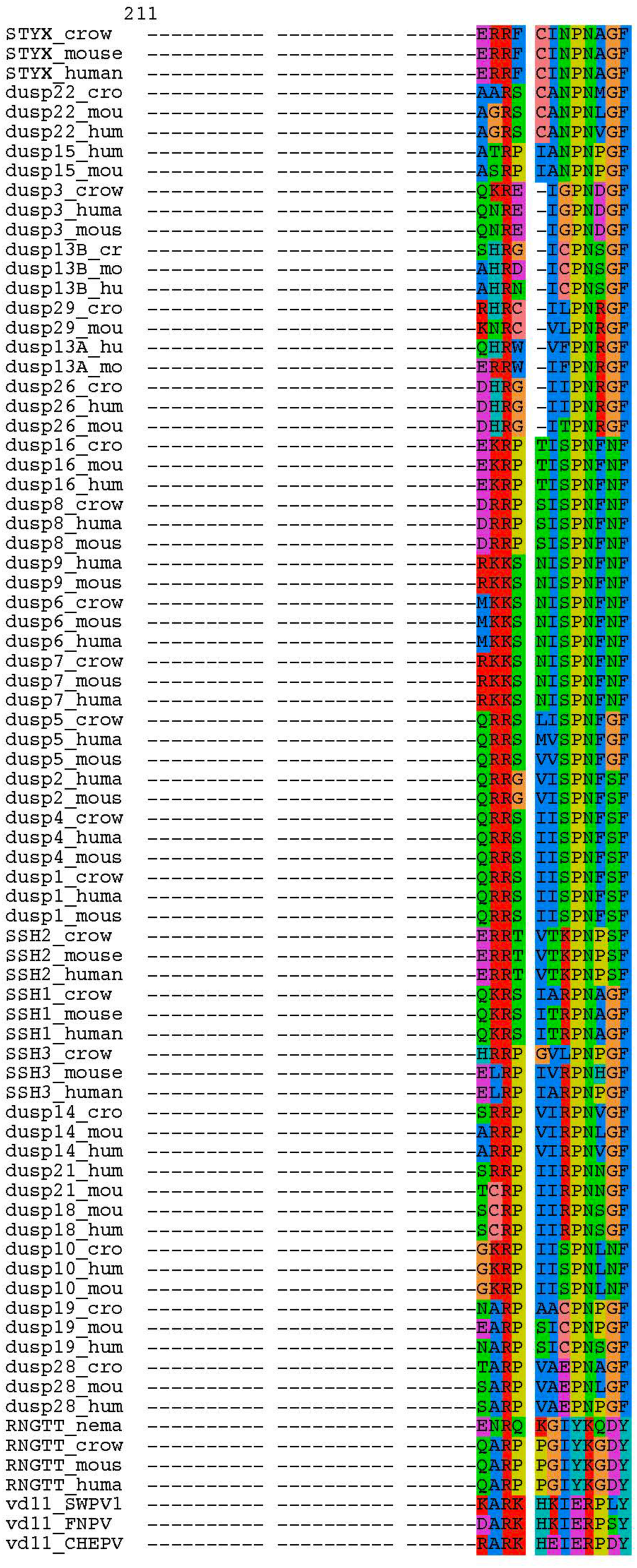
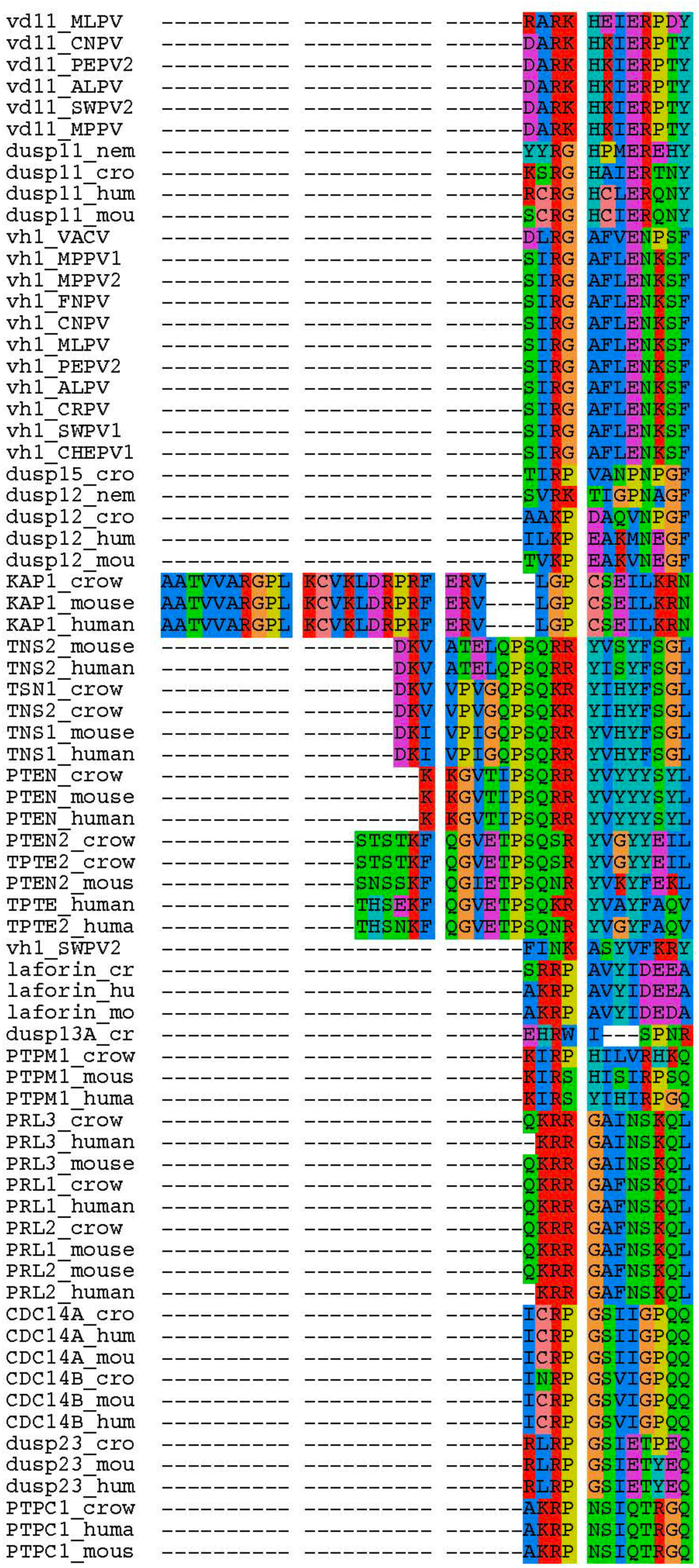
Clustal Omega alignment of host/poxviral DUSP sequences (trimmed) used for PhyML analysis.

**Figure S4.**
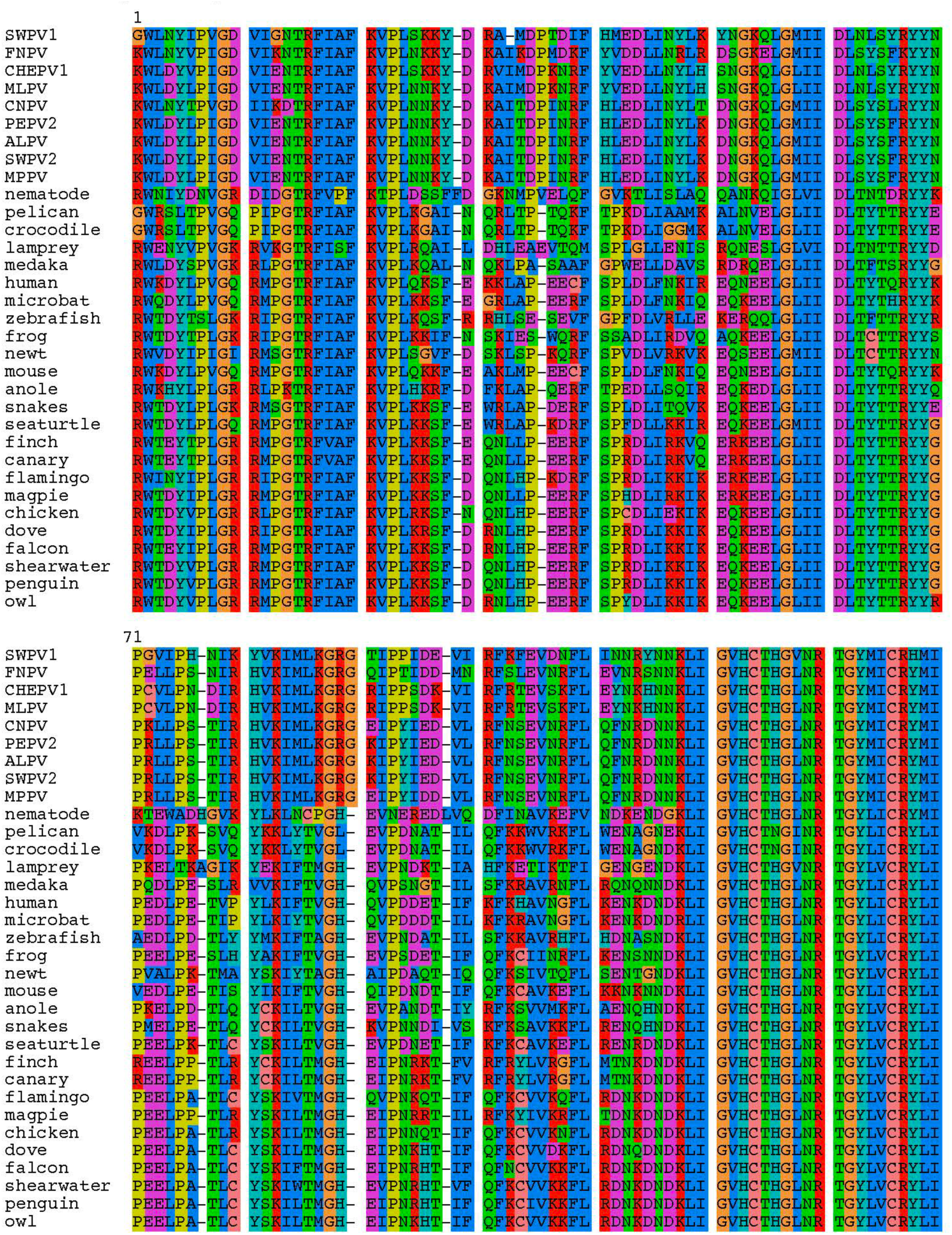
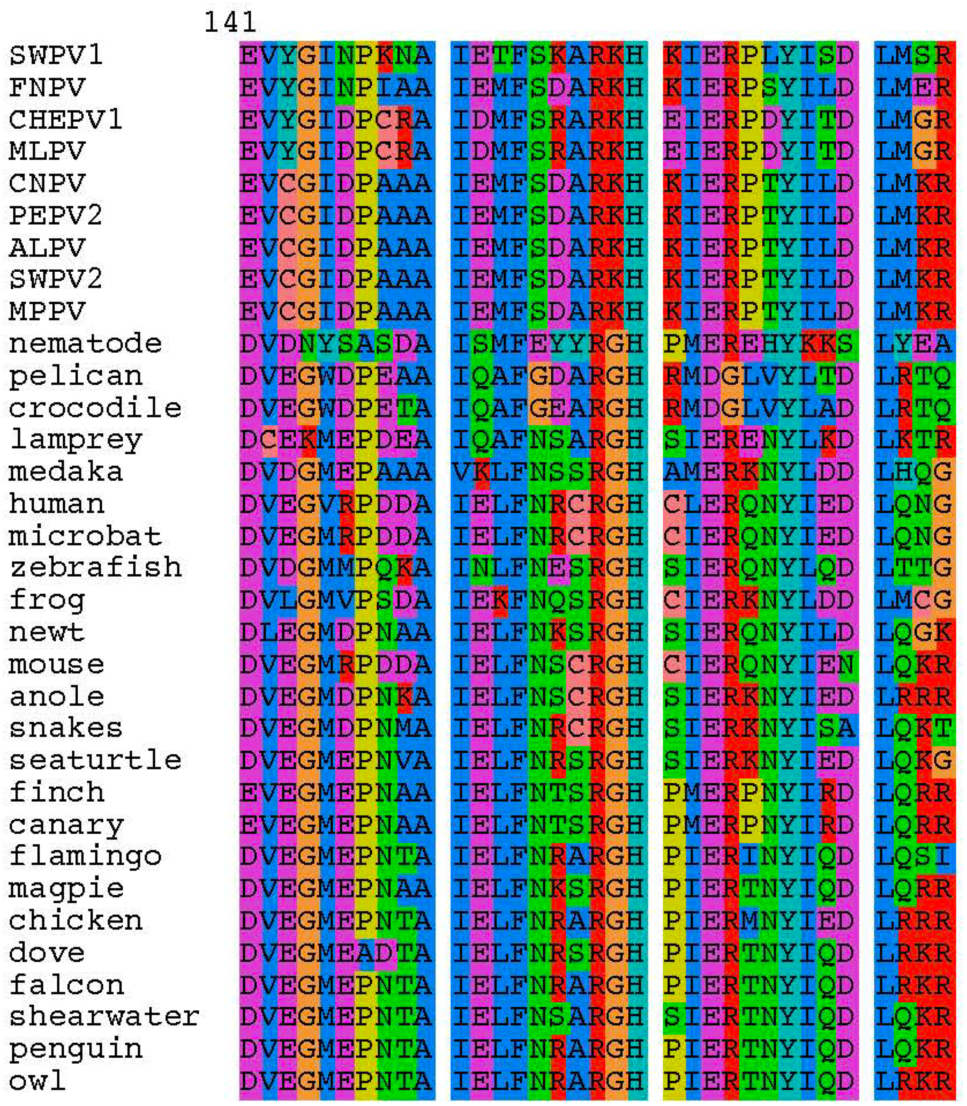
Clustal Omega alignment of host DUSP11 and avipox vDUSP11 sequences (trimmed) used for PhyML analysis.

**Figure S5:**
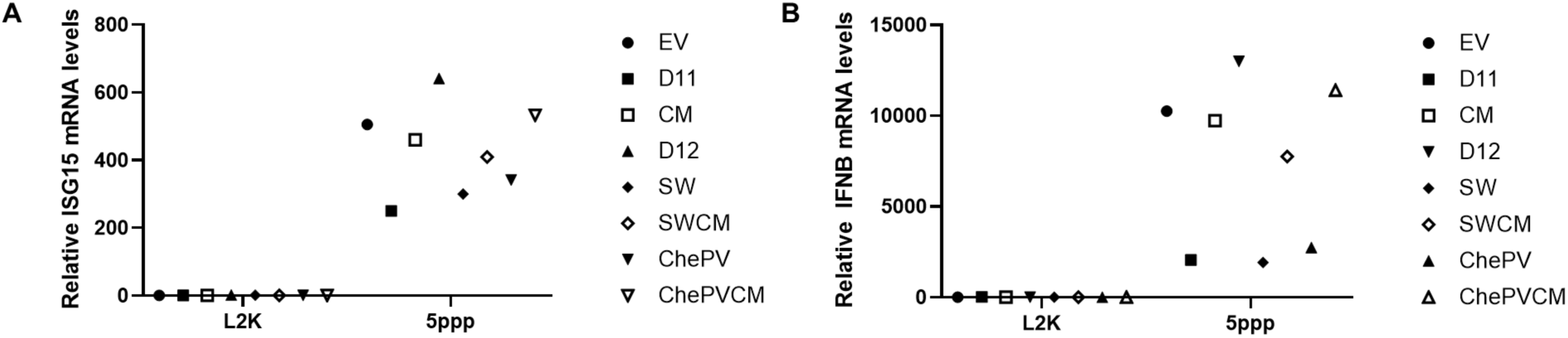
vDUSP11 modulates immune activation in response to liposomal 5’-ppp-RNAs. Confirmation of key result trends from Figure 4 via a different wet bench scientist. A549 DUSP11 knockout (KO) reconstituted cells (12-well) were transfected with 5-10 ng of *in vitro* transcribed 5’-ppp-RNA for 18 hours followed by RT-qPCR to assay induction of ISGs. (A) RT-qPCR analysis of *ISG15* and (B) *IFNB1* mRNA normalized to *GAPDH* mRNA in 5’-ppp-RNA transfected in A549 DUSP11 KO reconstituted cells. Results are represented relative to those of empty vector-expressing cells. Data are derived n = 1 replicates.

**Figure S6.**
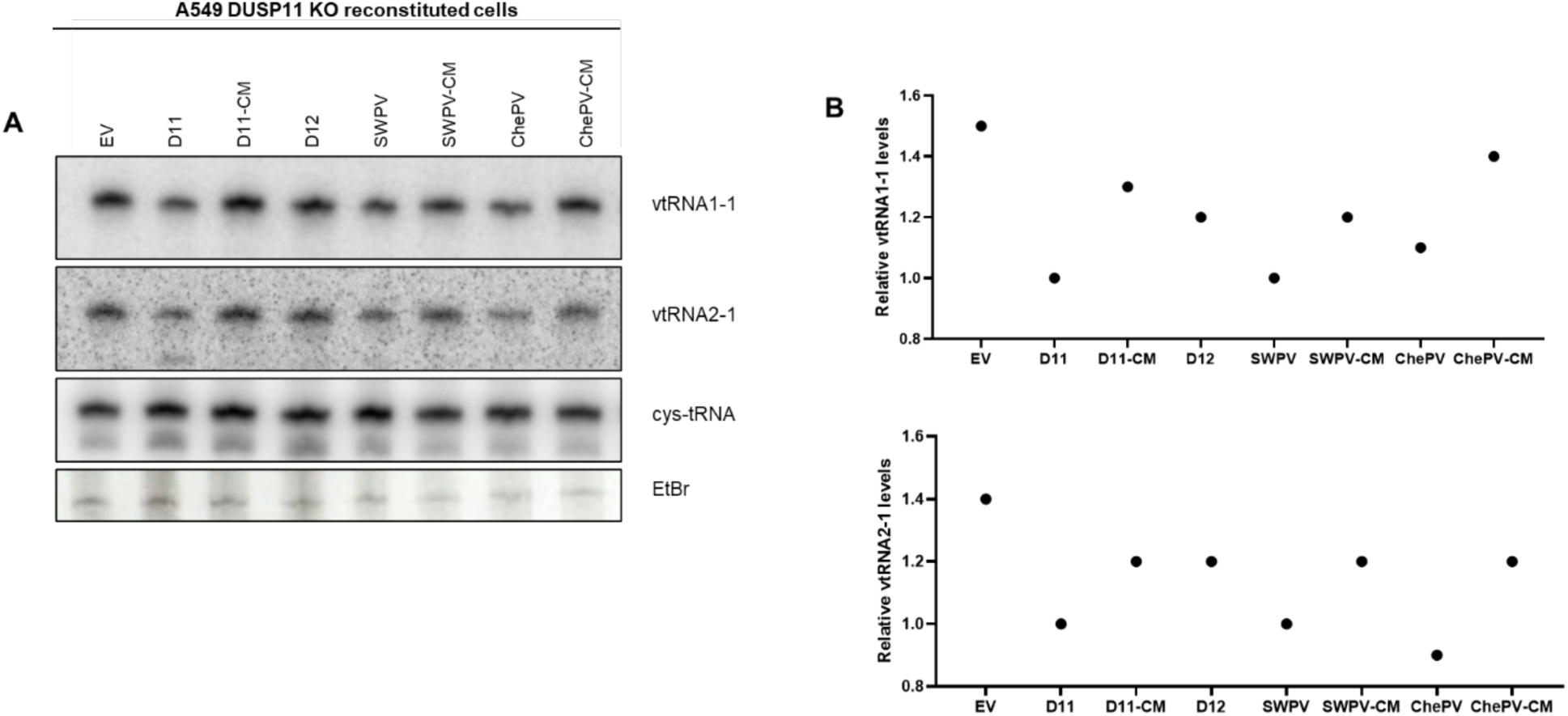
vDUSP11 modulates steady-state RNA levels of endogenous RNAP III transcripts. Confirmation of key result trends from Figure 6 via a different wet bench scientist. (A) Northern blot analysis of vtRNA1-1 and vtRNA2-1 using RNA from A549 DUSP11 KO reconstituted cells. (B) Graphical representation of relative band intensity of vtRNA1-1 and vtRNA2-1 normalized to the relative band intensity of the cysteine-tRNA. Data are derived from n = 1 replicates.

**Figure S7.**
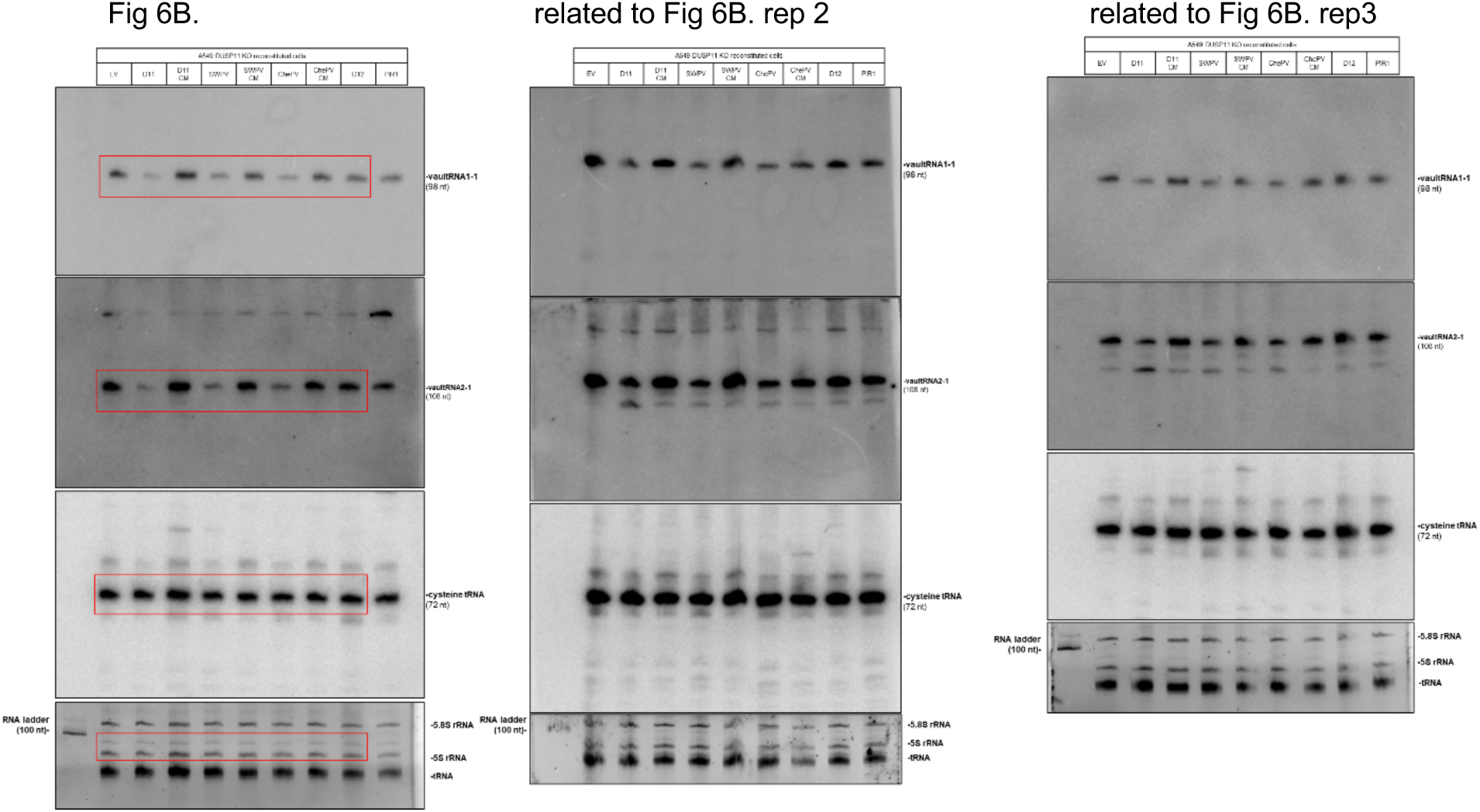
uncropped northern blots.

**Figure S8.**
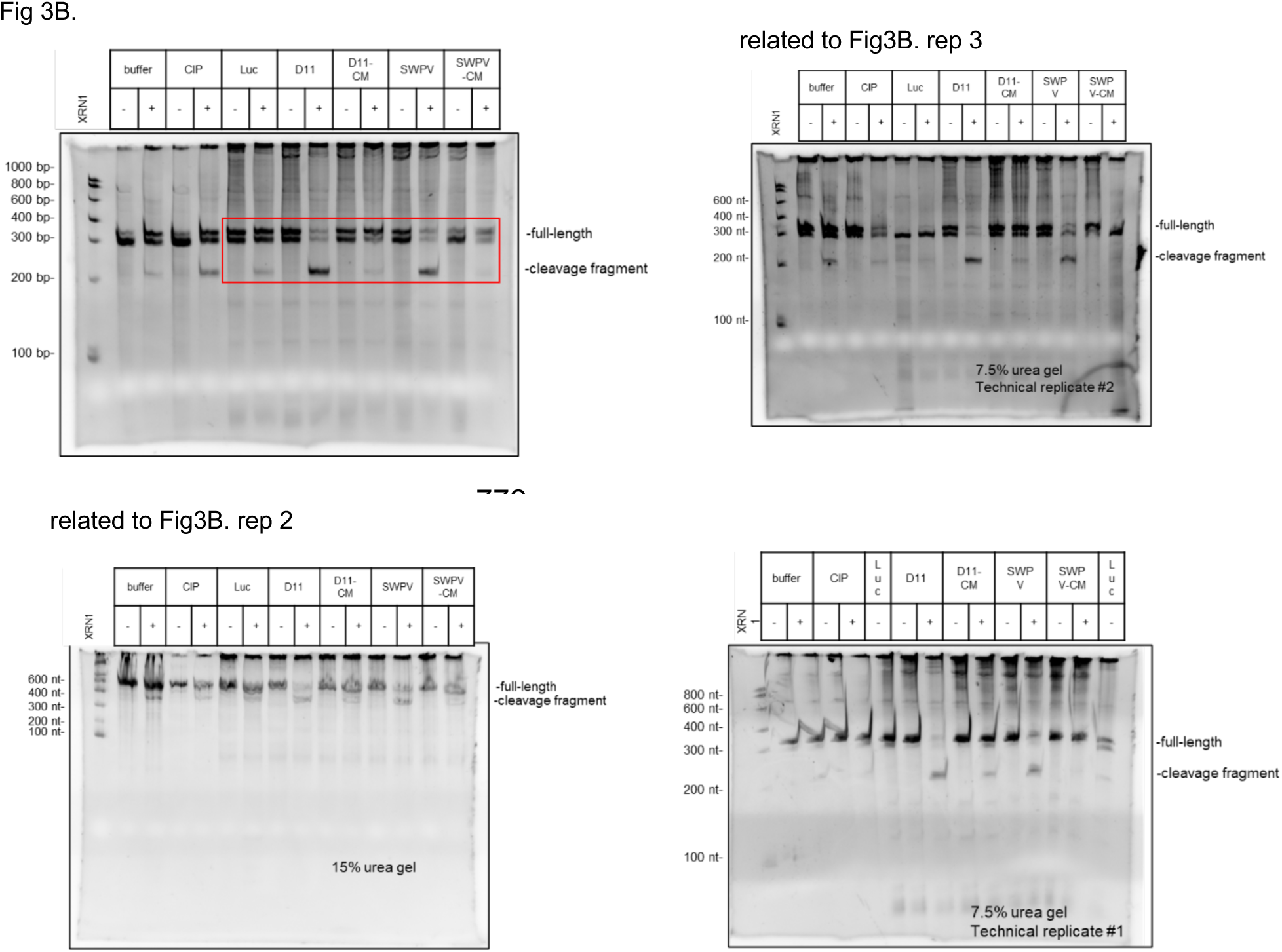
uncropped ethidium bromide-stained urea gels.

**Figure S9:**
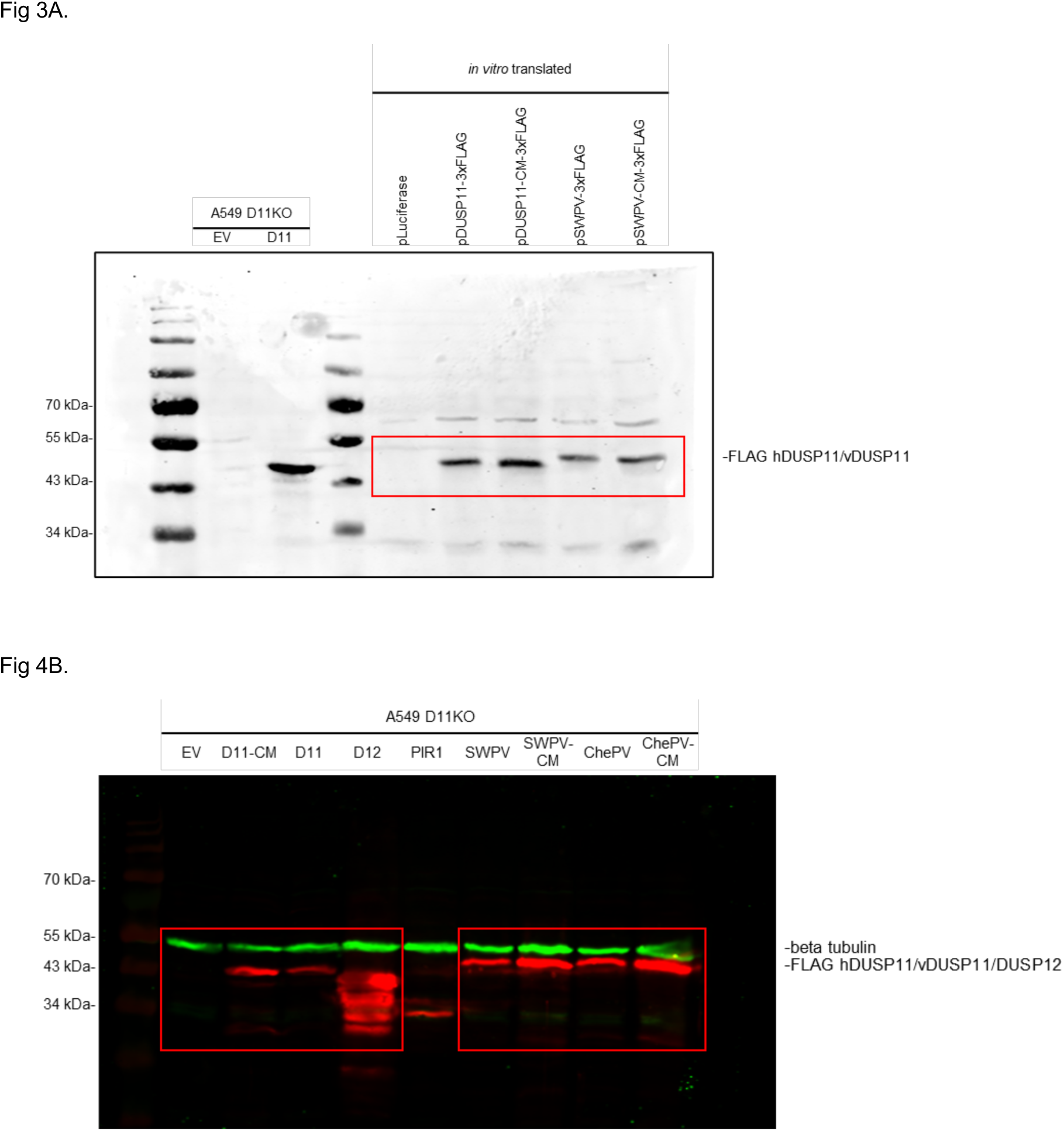
uncropped western blots.

**Figure S10:**
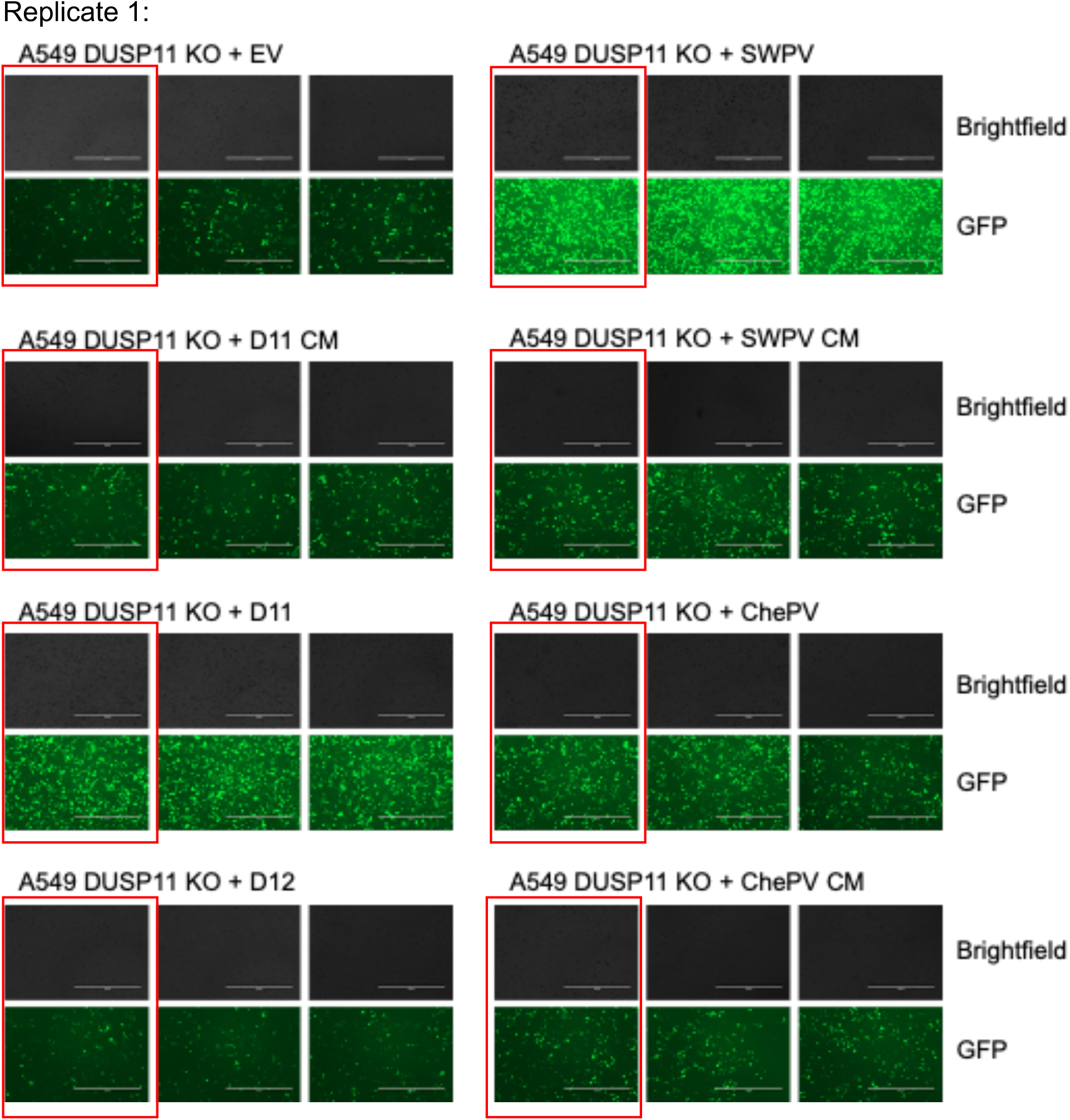
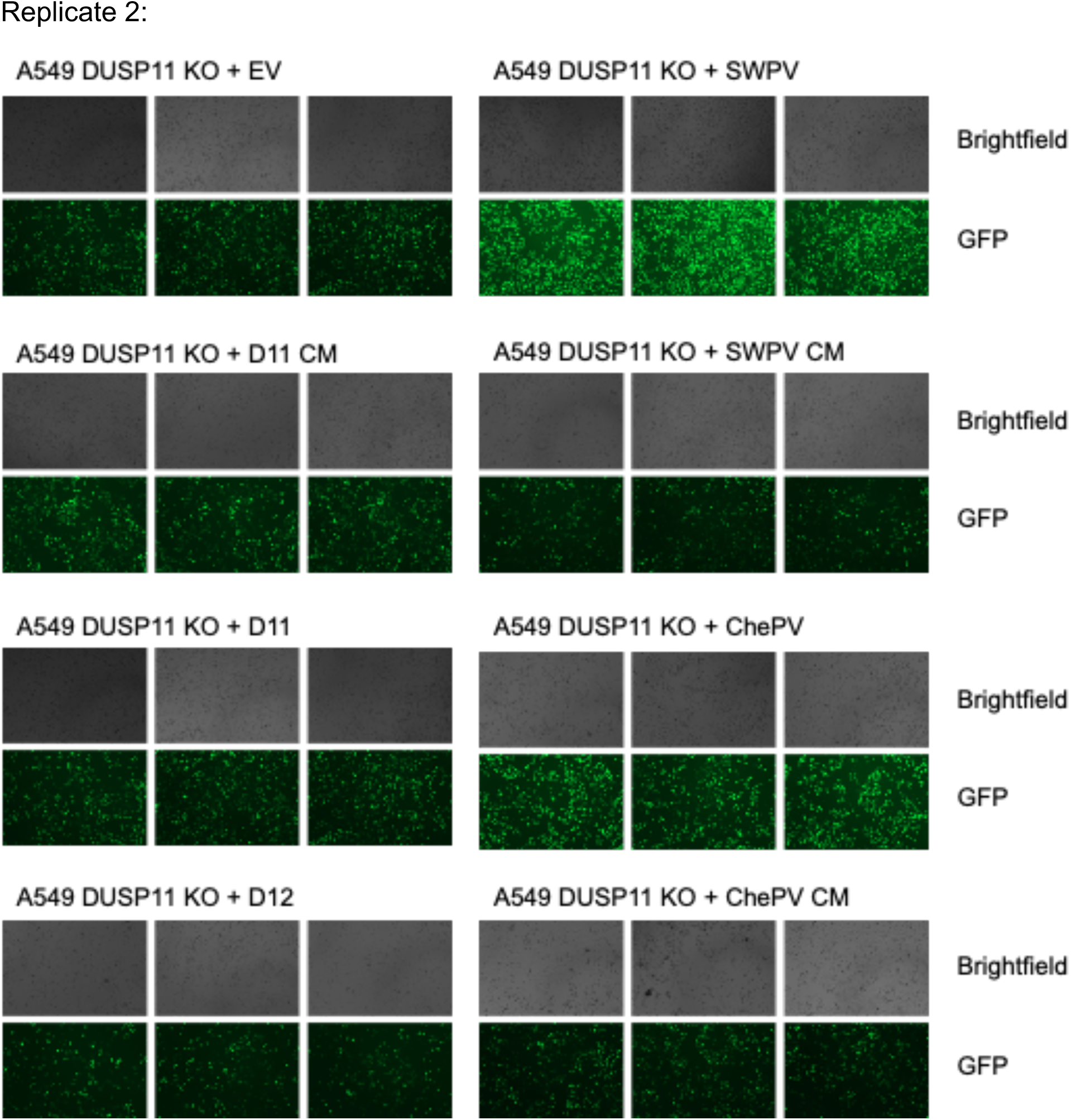
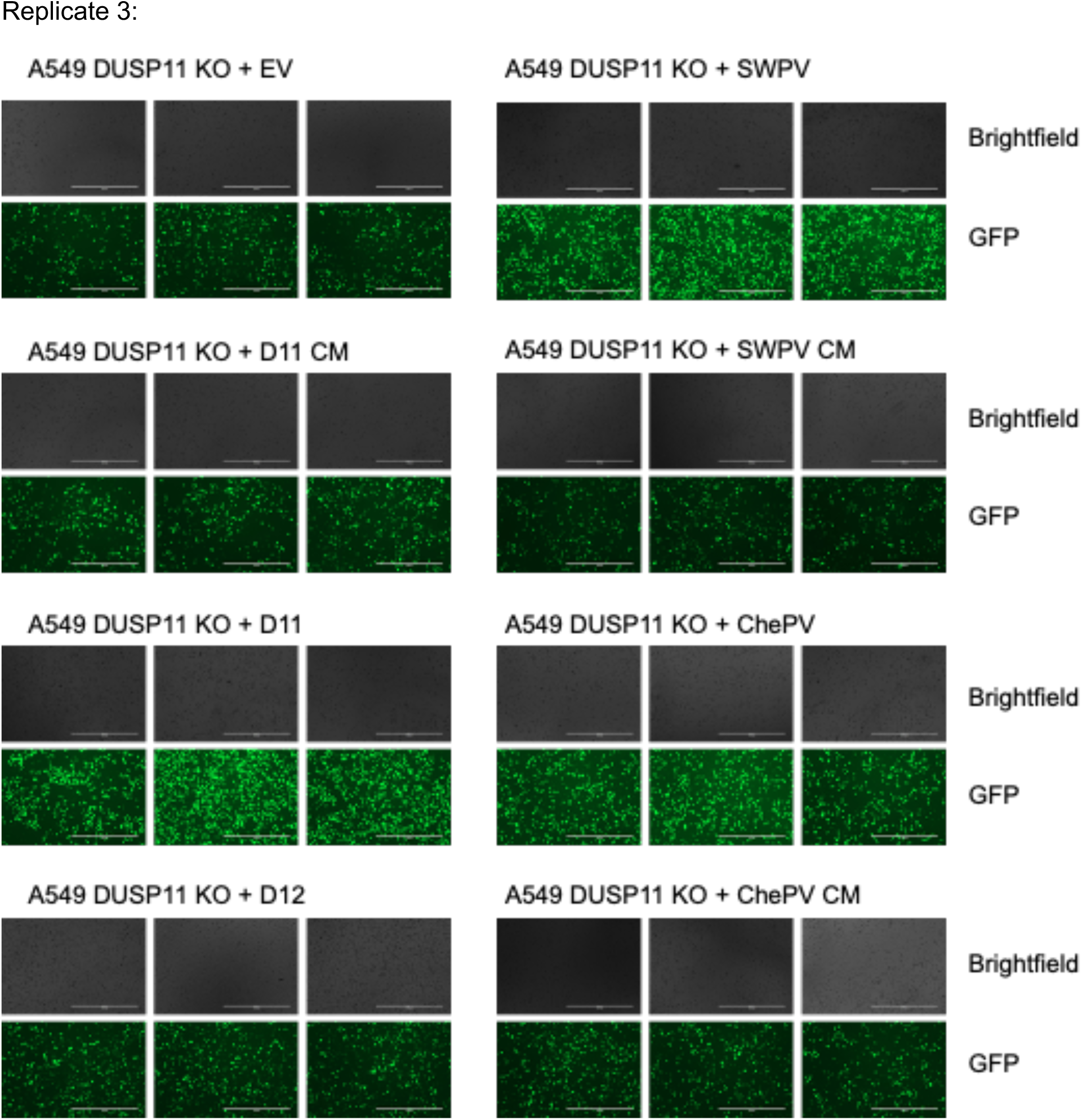
M51R VSV infection GFP images.

**Table S1:**
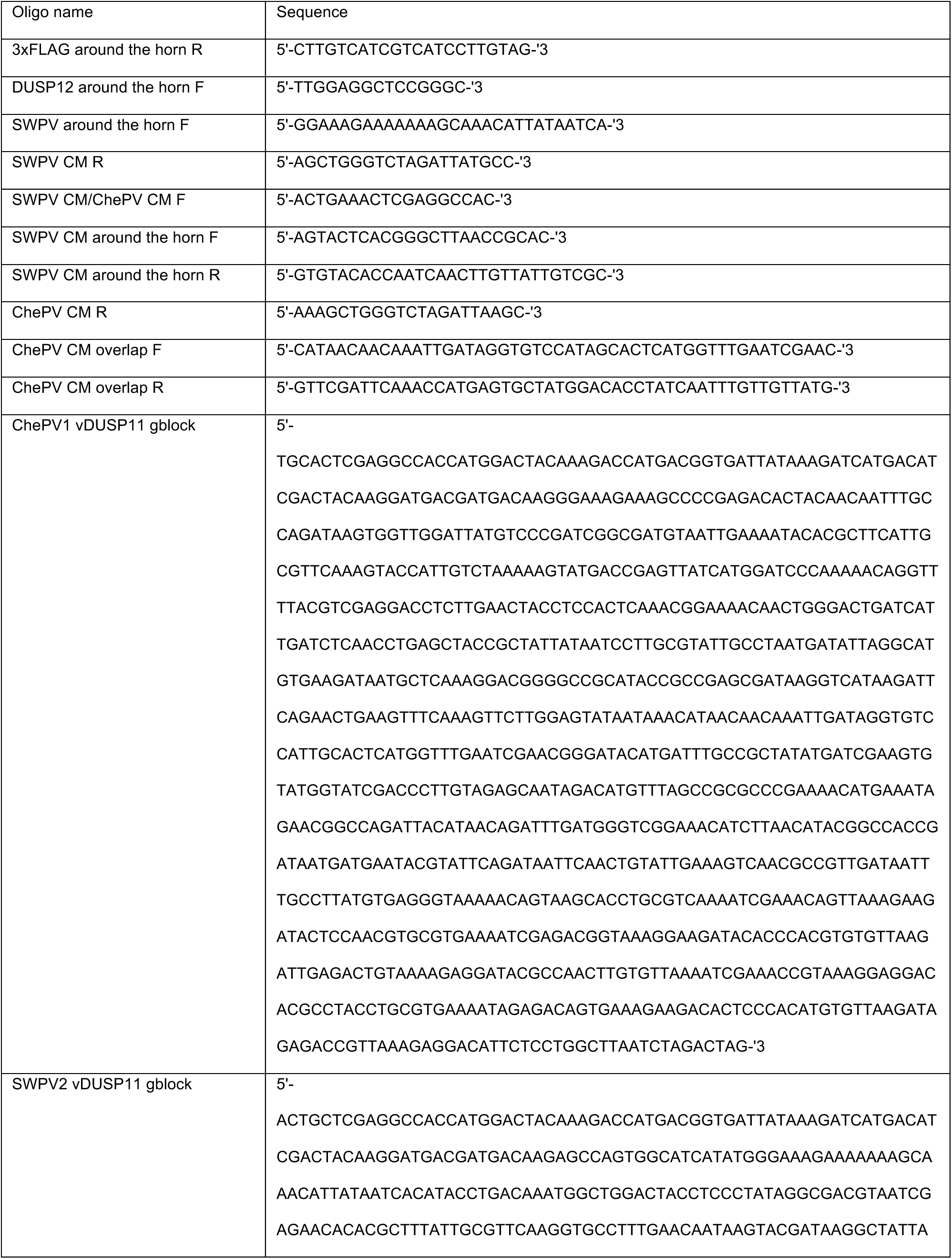
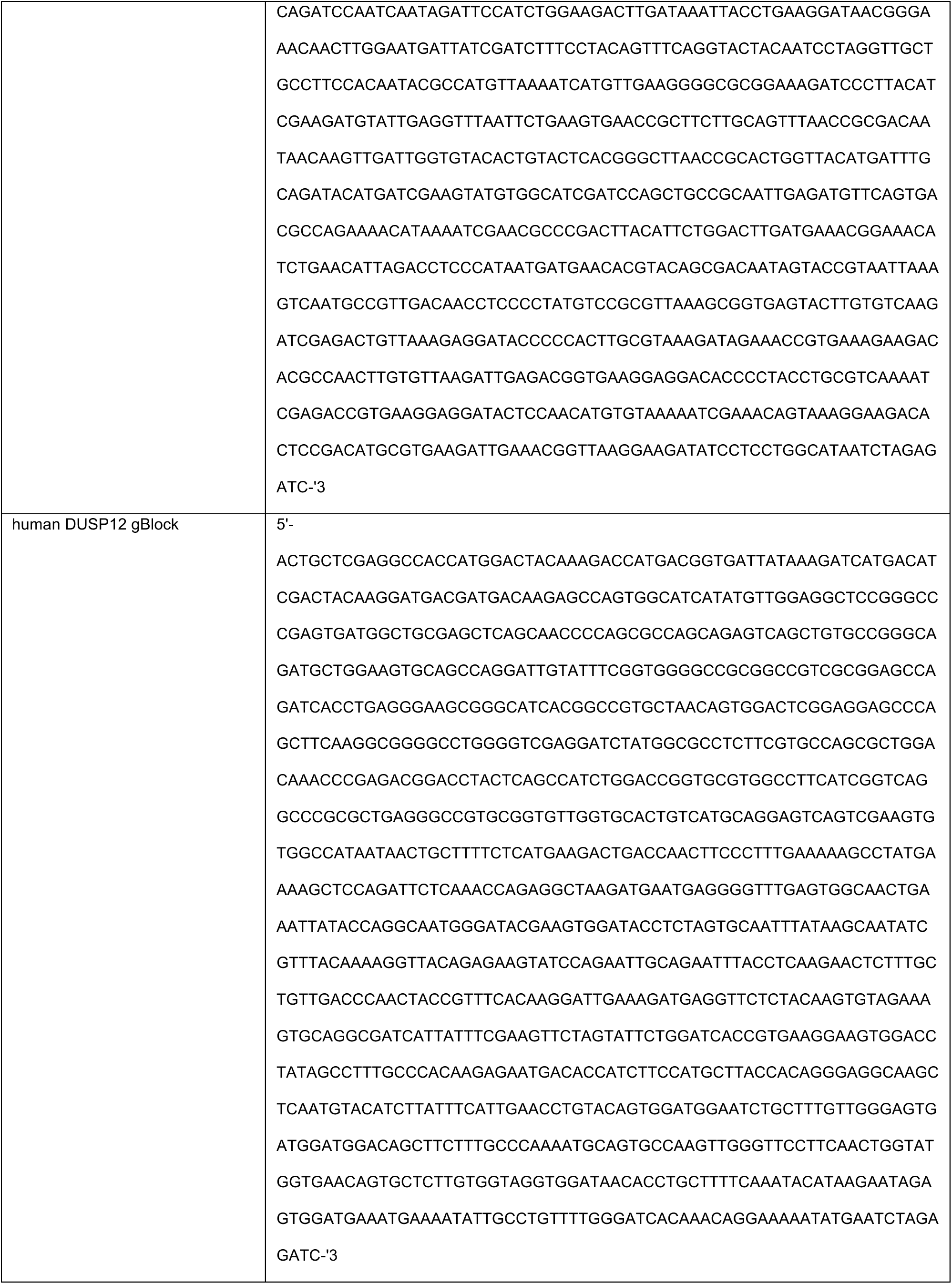
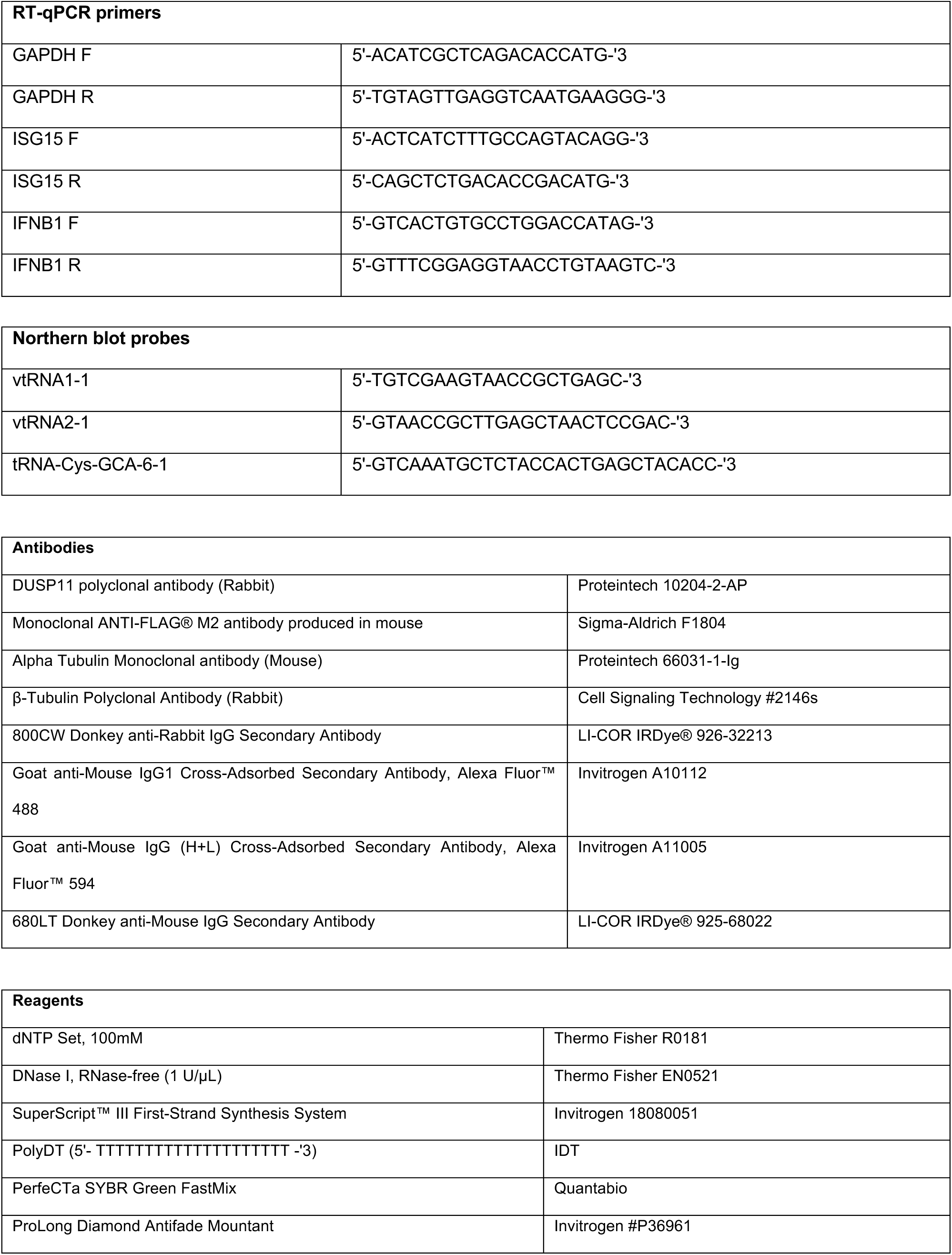
Sequence information, synthetic oligonucleotides and DNA, and reagents.

**Table S2.**
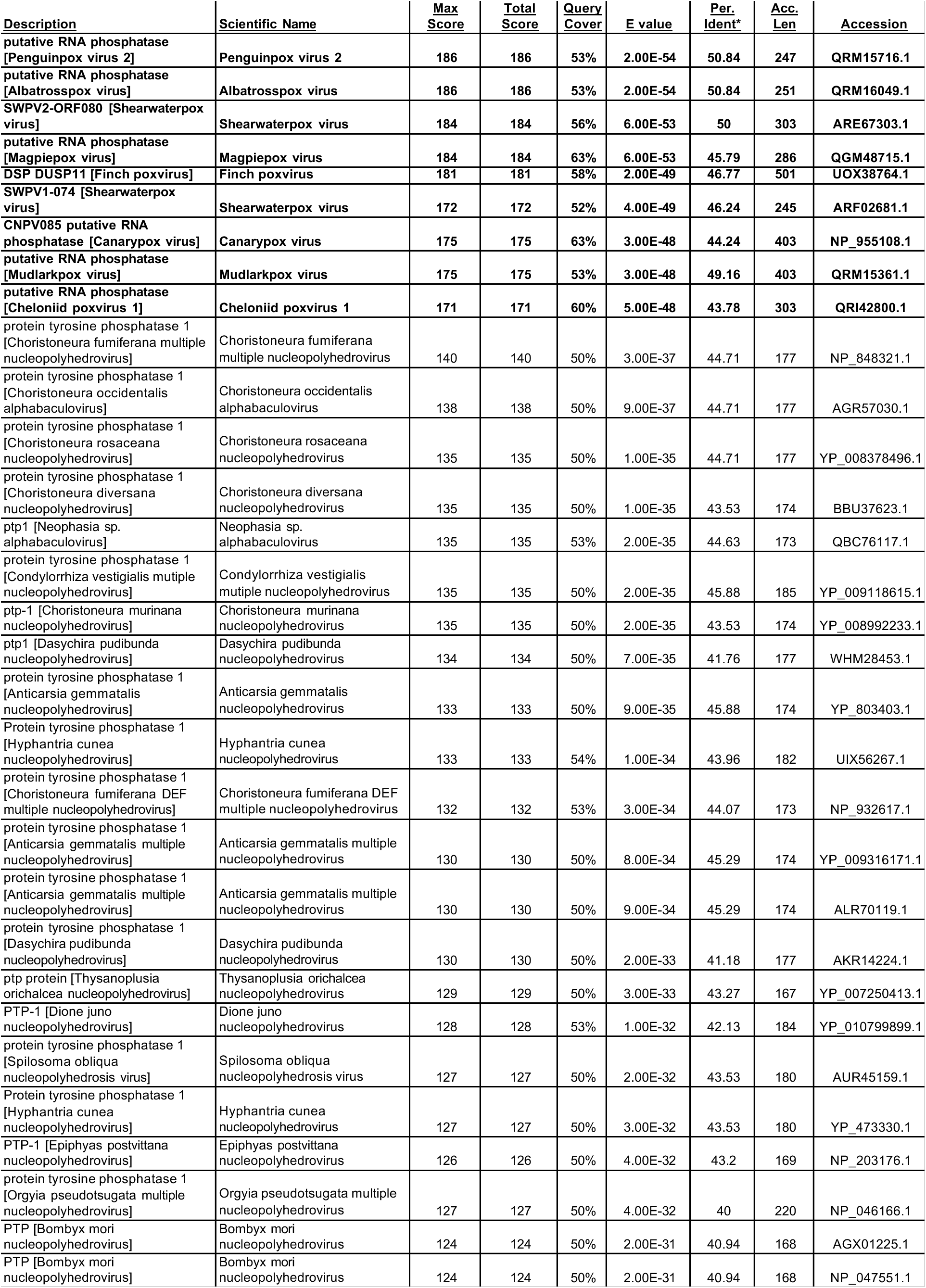
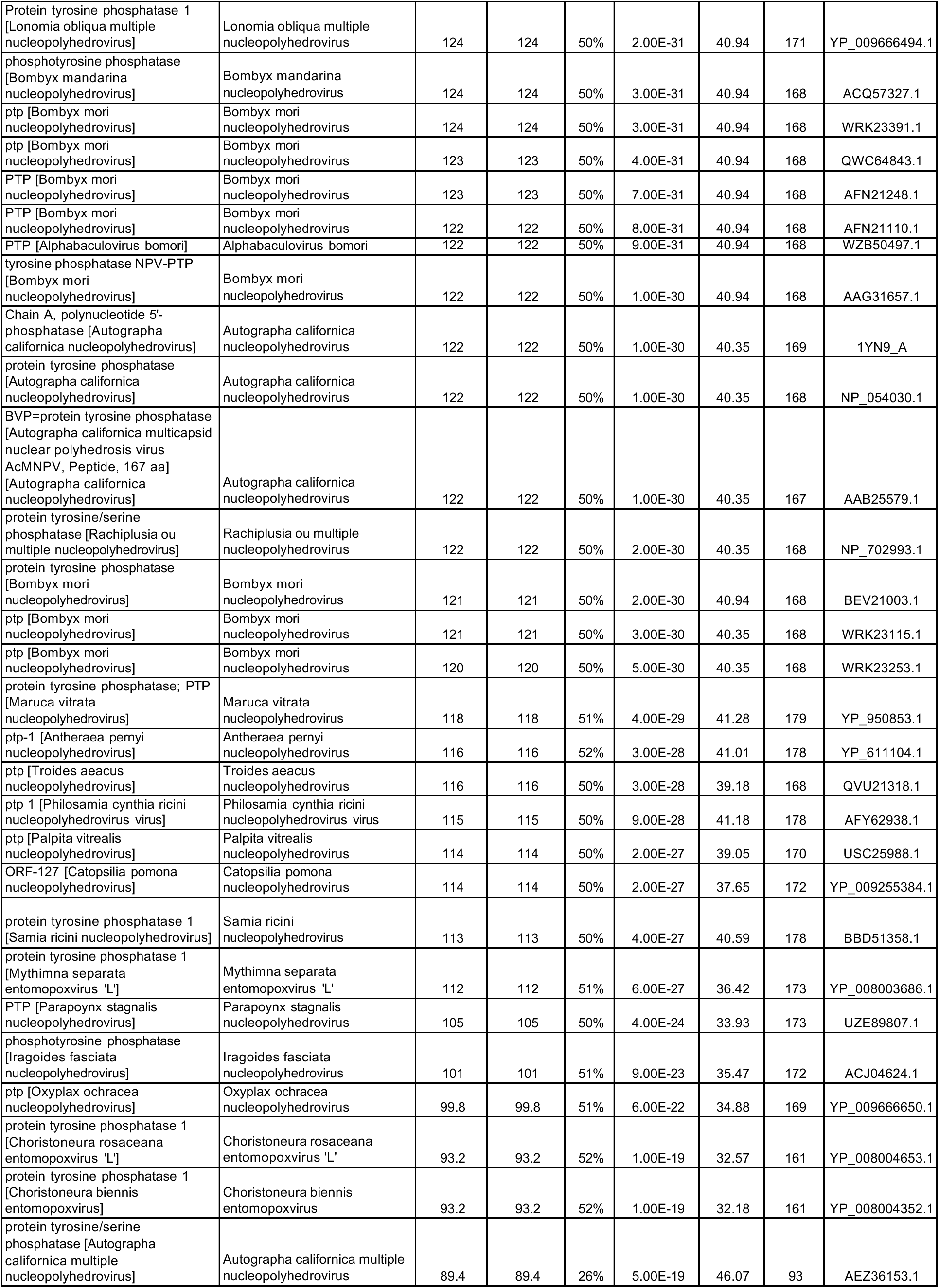
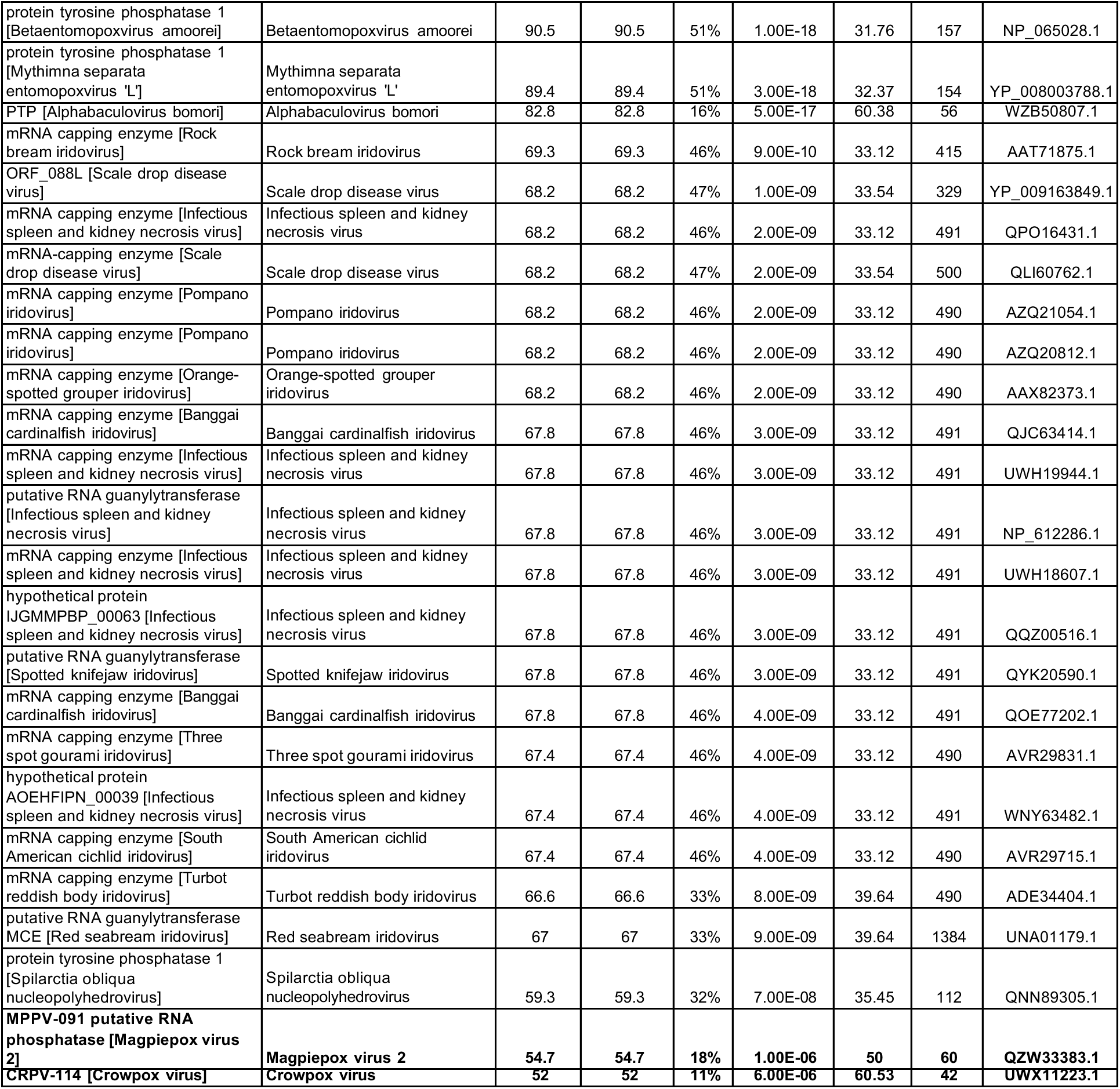
Blastp results using human DUSP11 (uniprot o75319) as query limiting results to viruses (taxid: 10239)

**Table S3.**
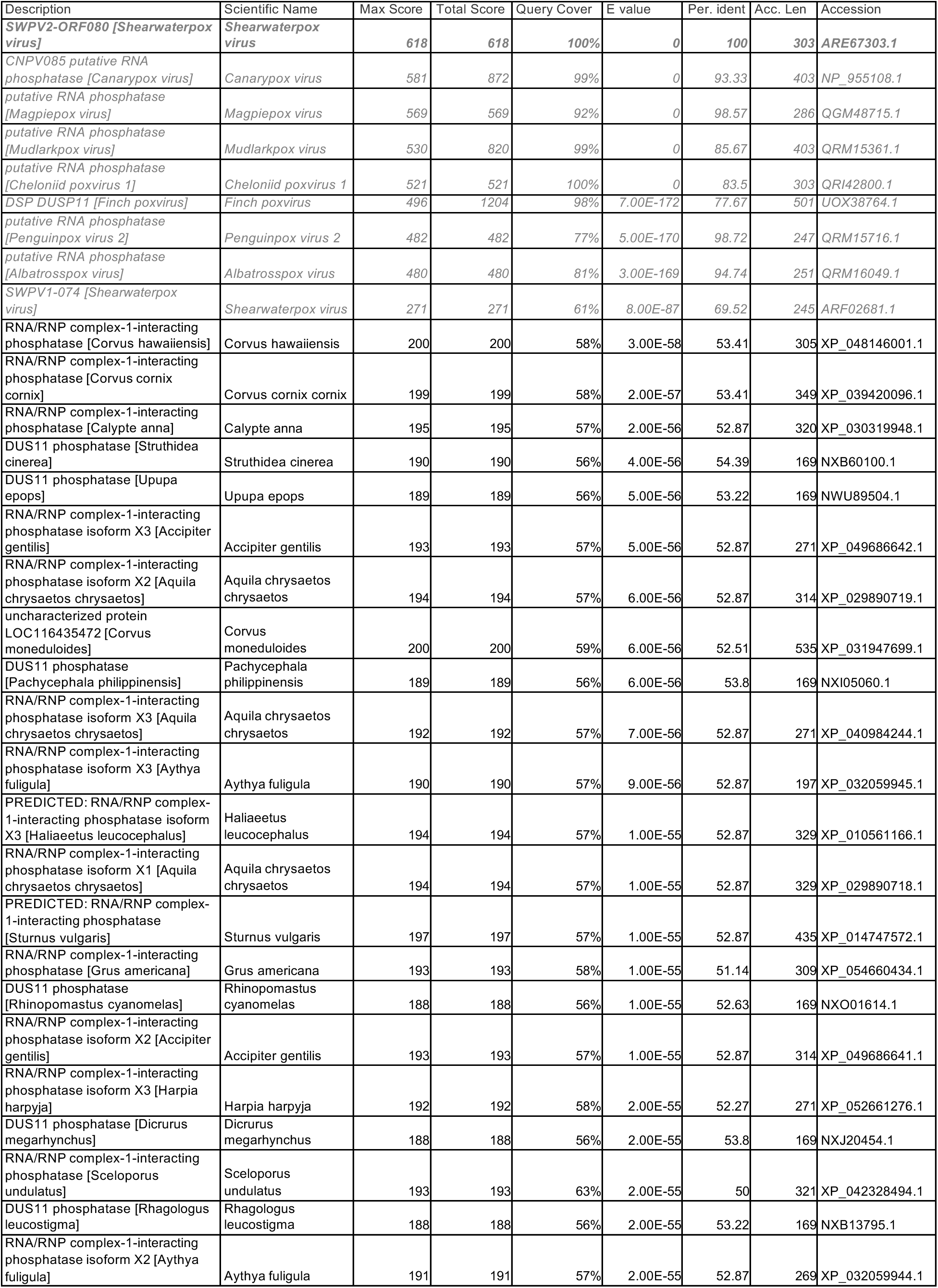
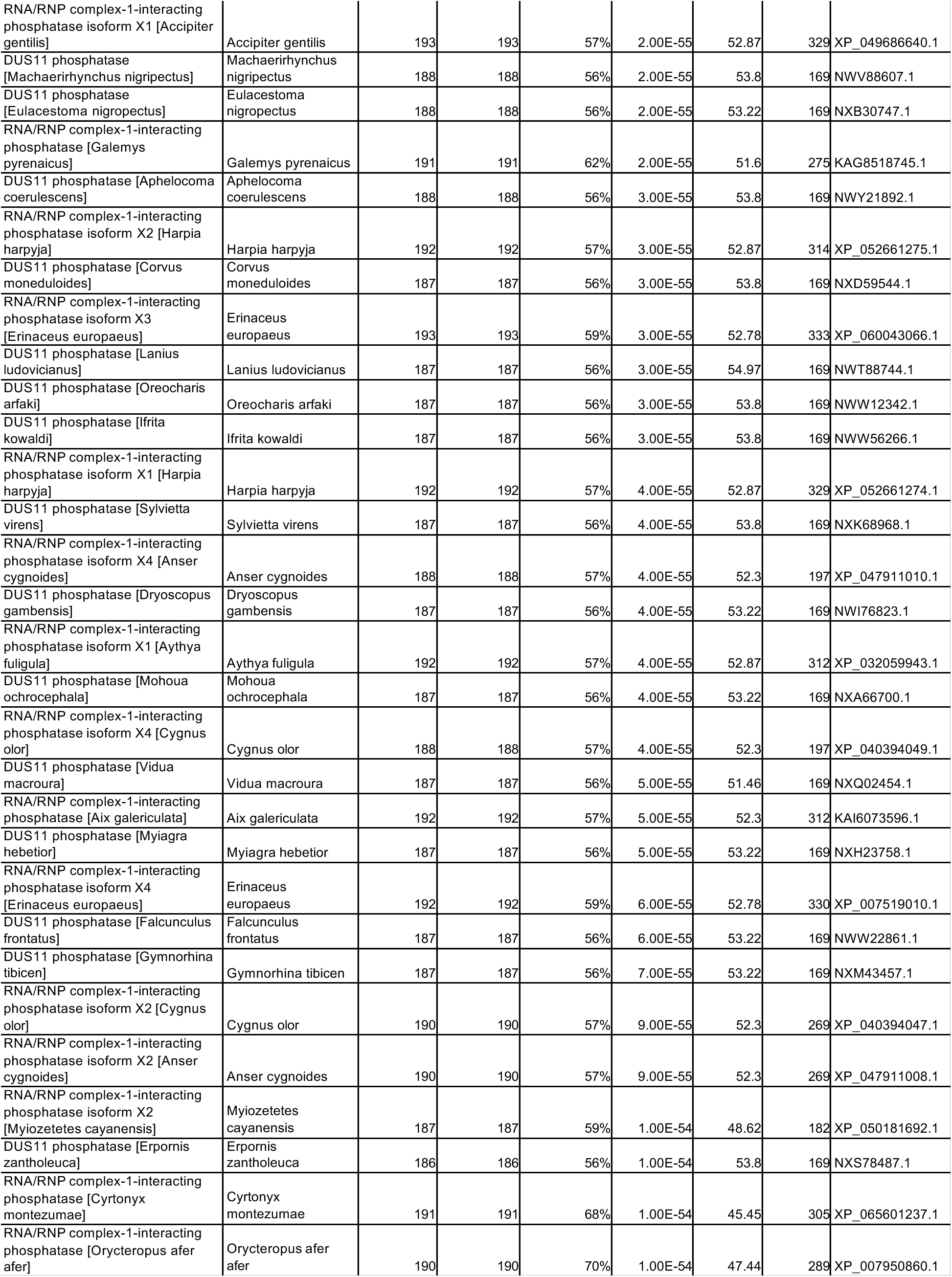
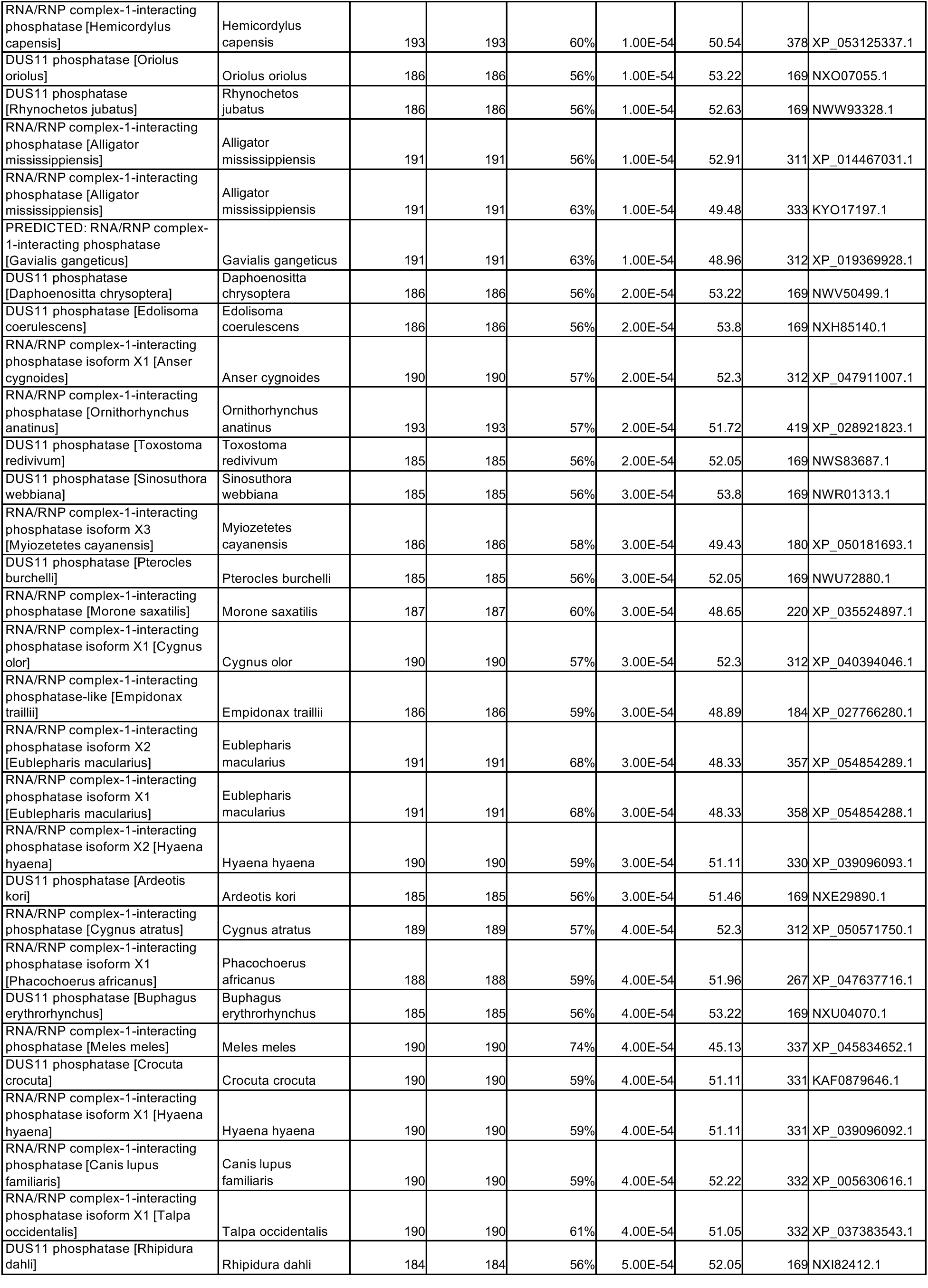
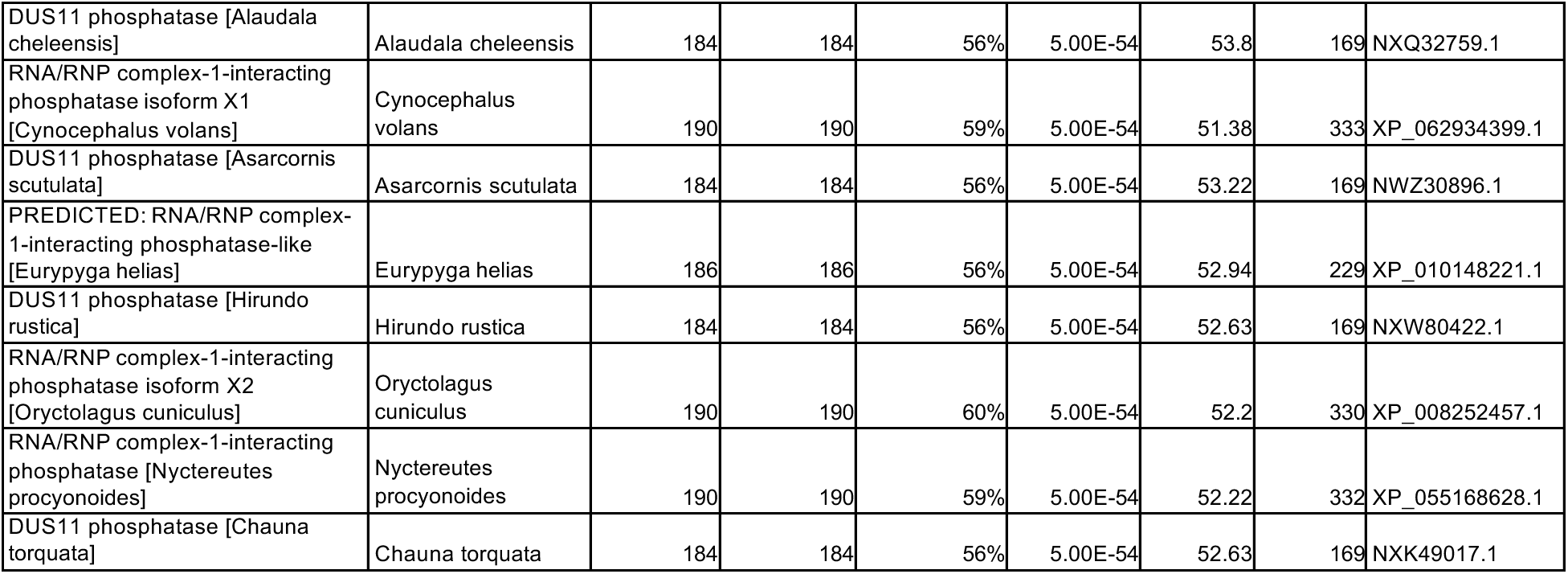
Blastp results using vDUSP11 from SWPV2 as query (ARE67303.1)

**Table S4.**
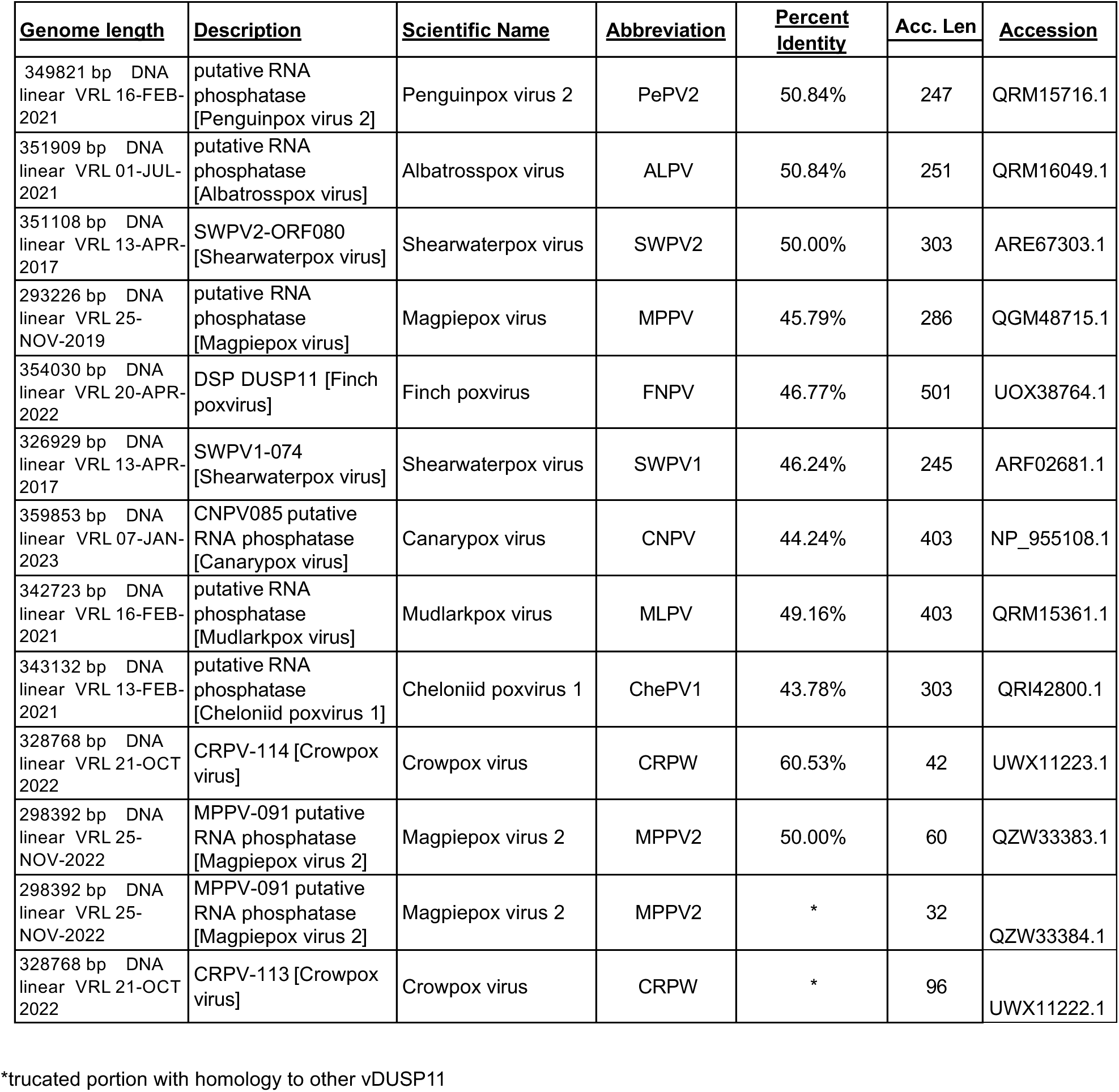
Avipox and avipox-adjacent viral DUSP11 accession IDs.

**Table S5.**
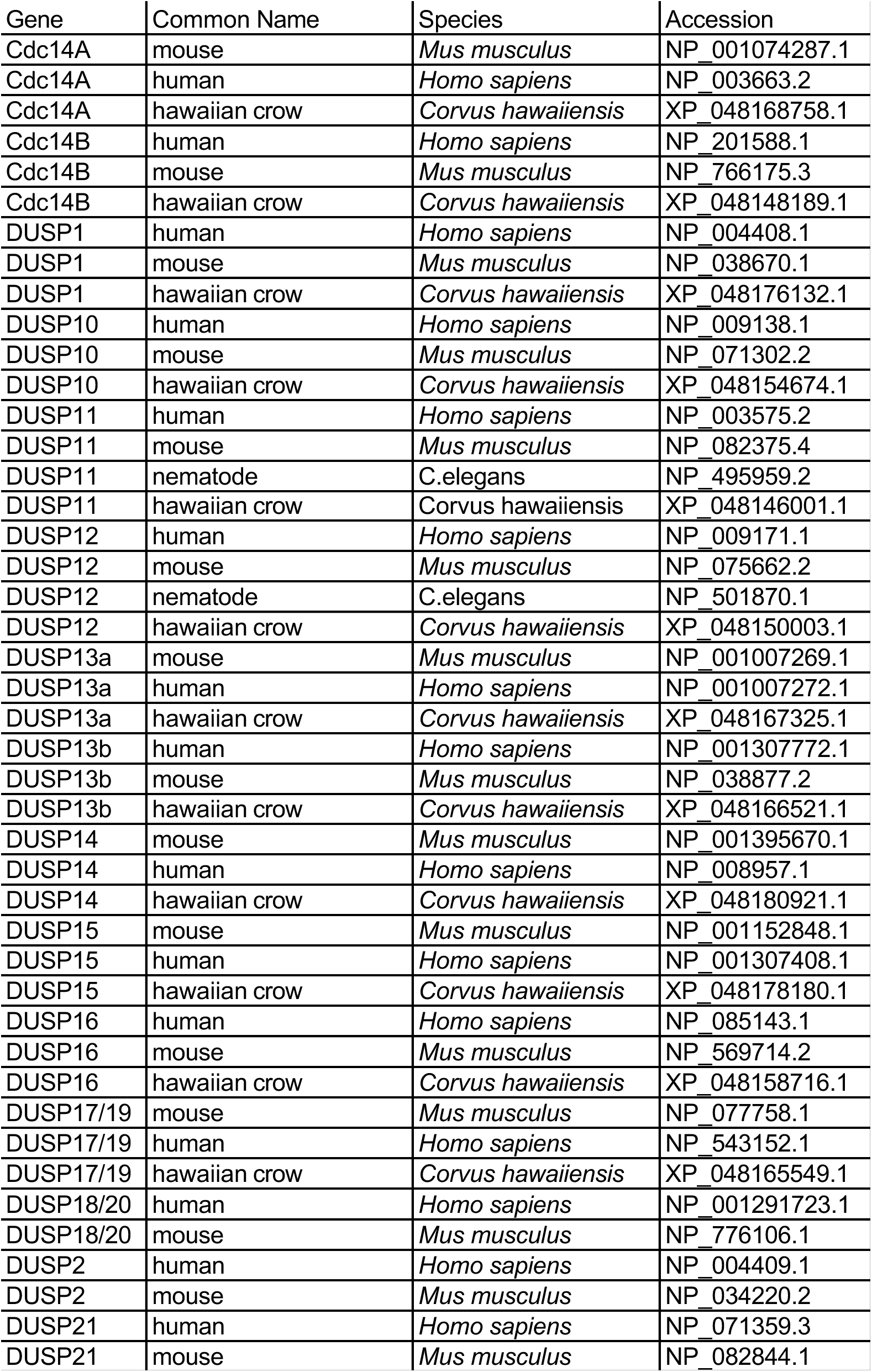
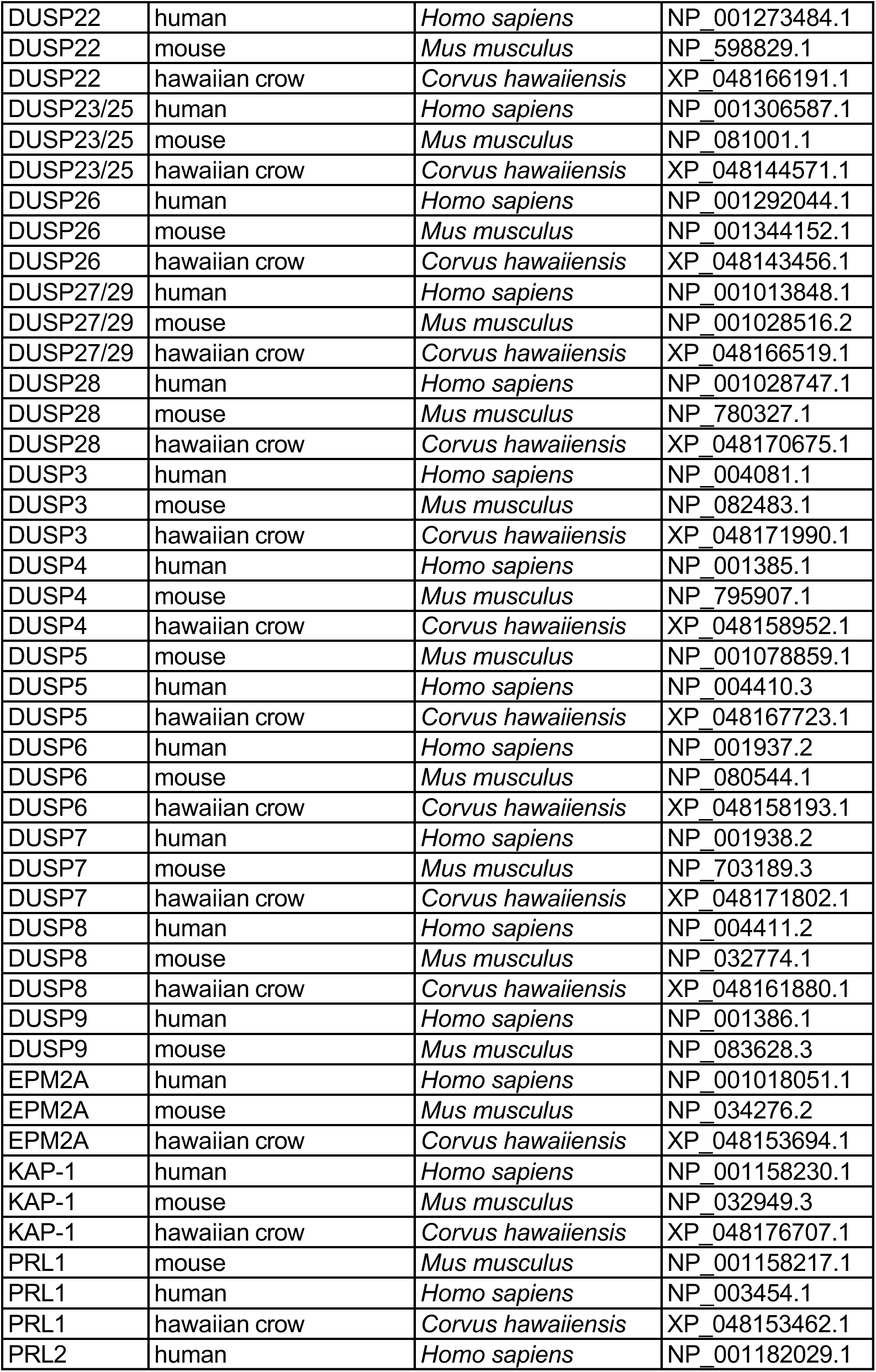
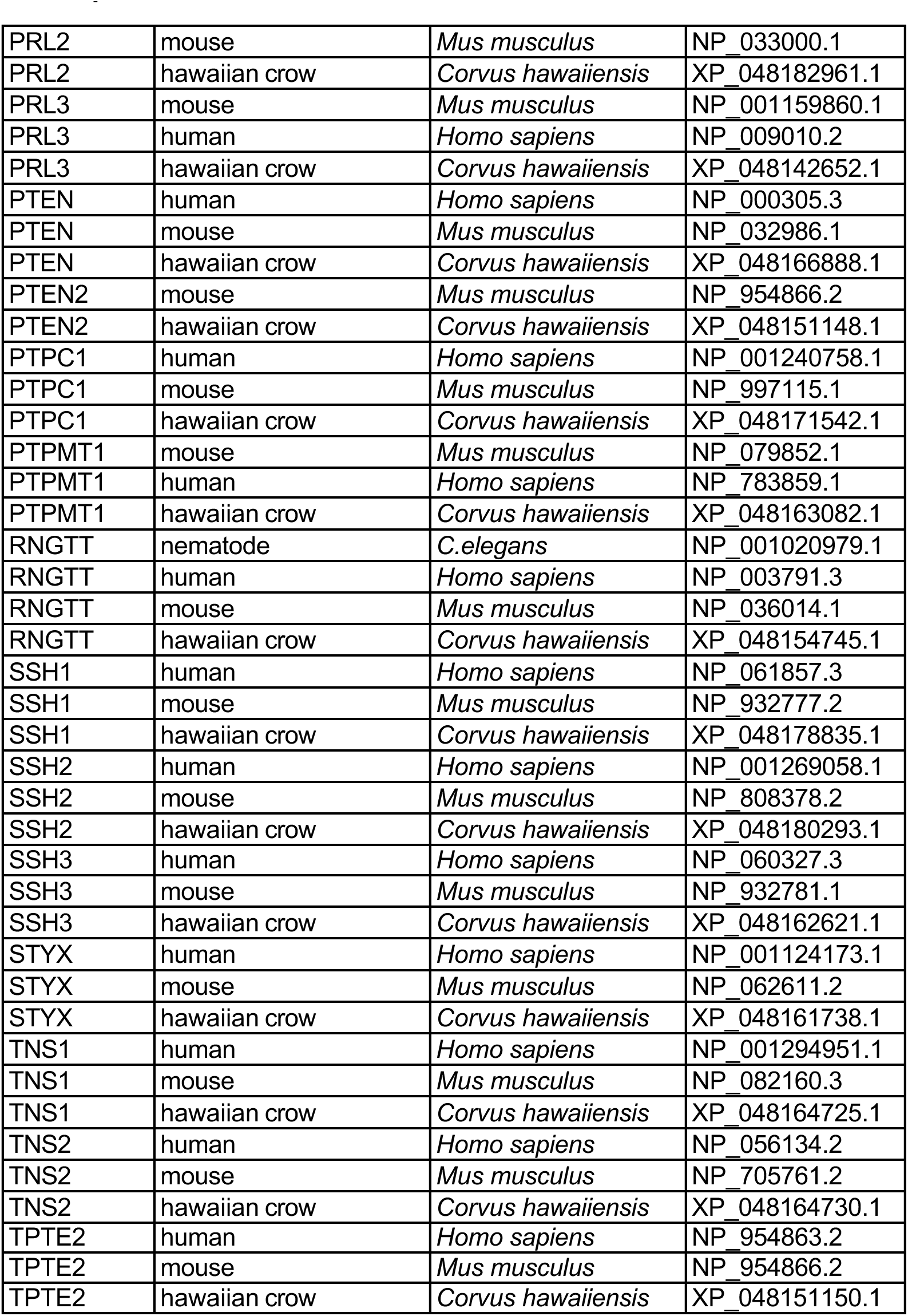
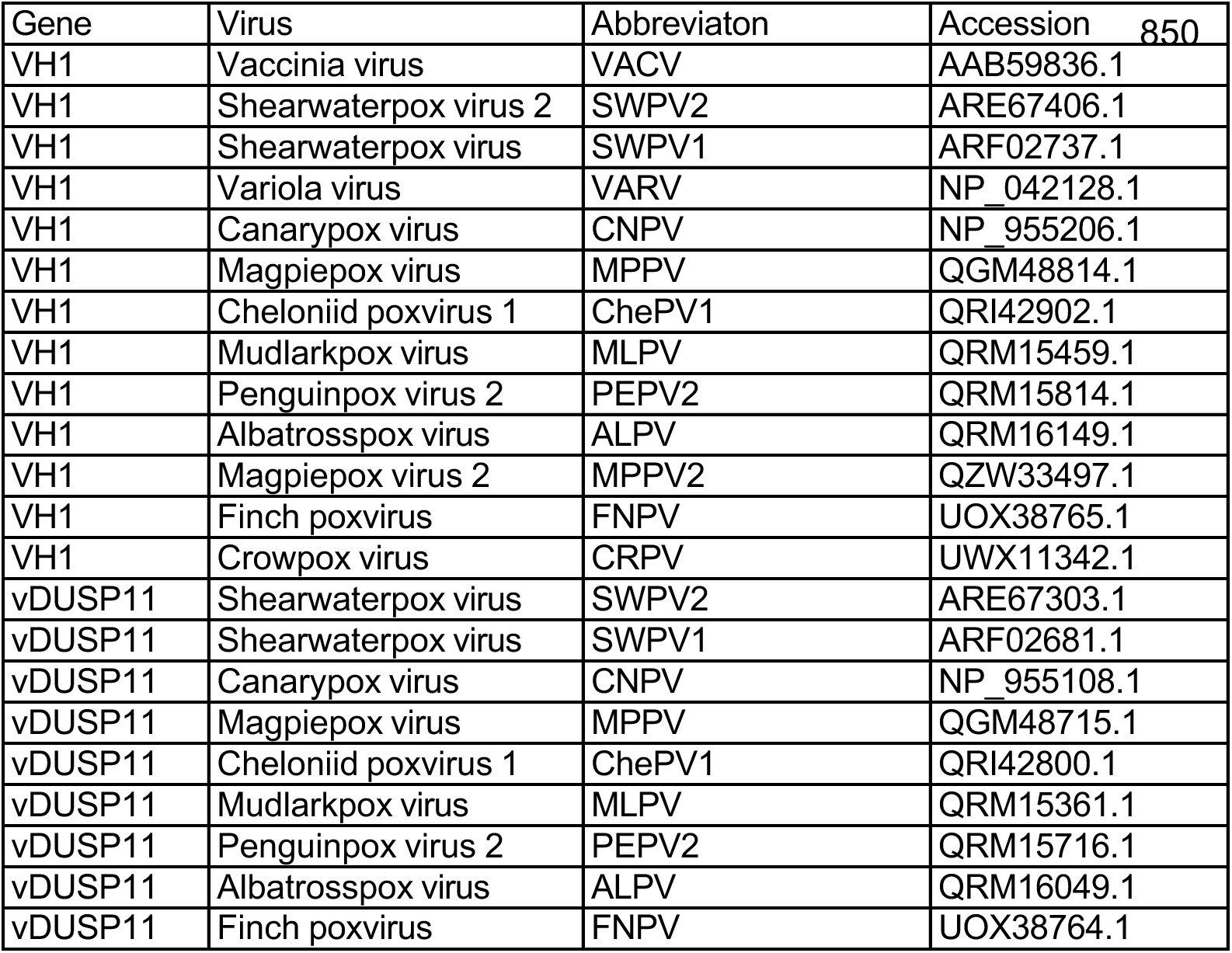
Accession IDs used for phylogenetic analysis of host DUSPs and related poxviral DUSPs.

**Table S6.**
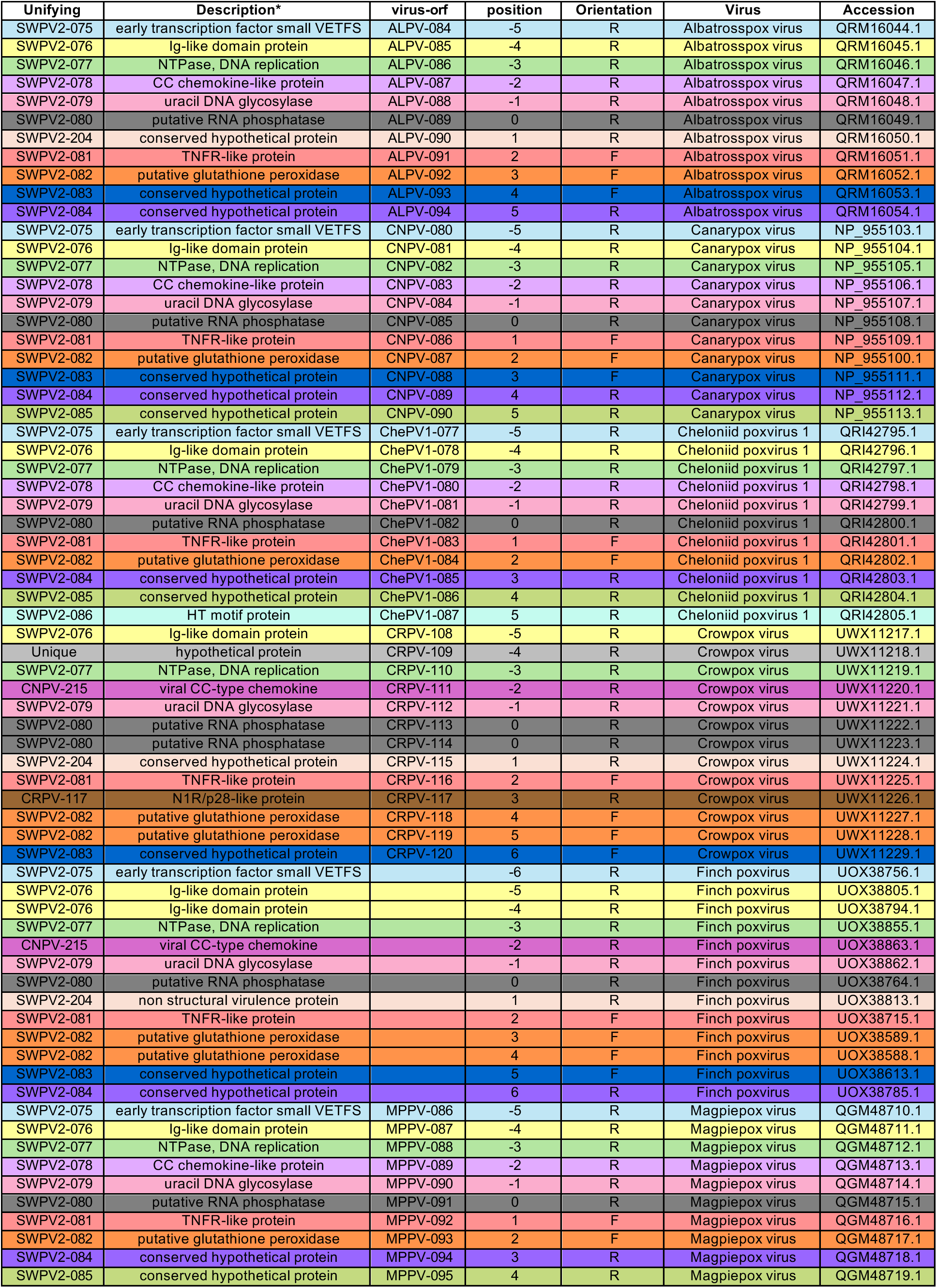
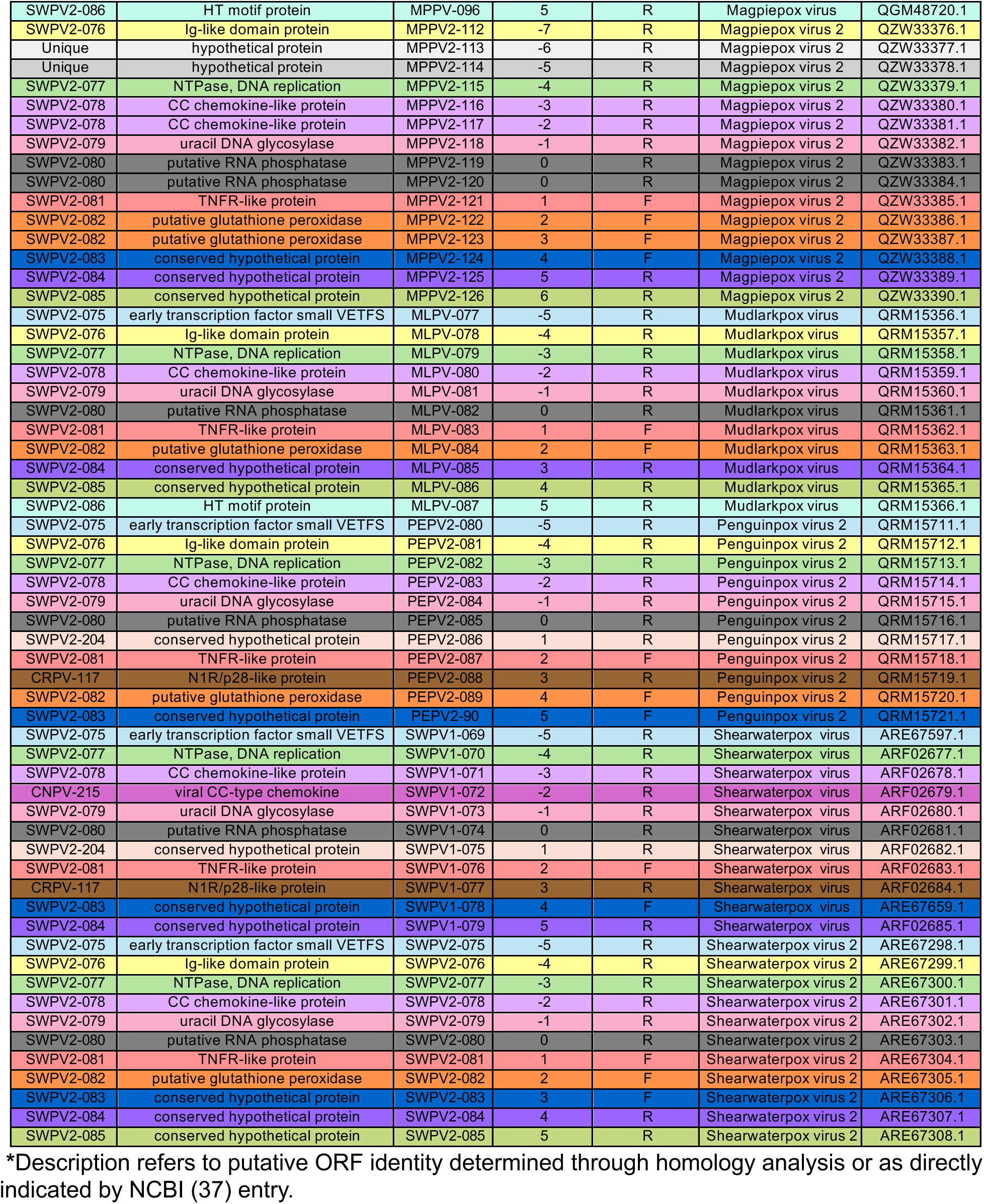
Genes surrounding the genomic locations of avipox vDUSP11 from synteny analysis.

**Table S7.**
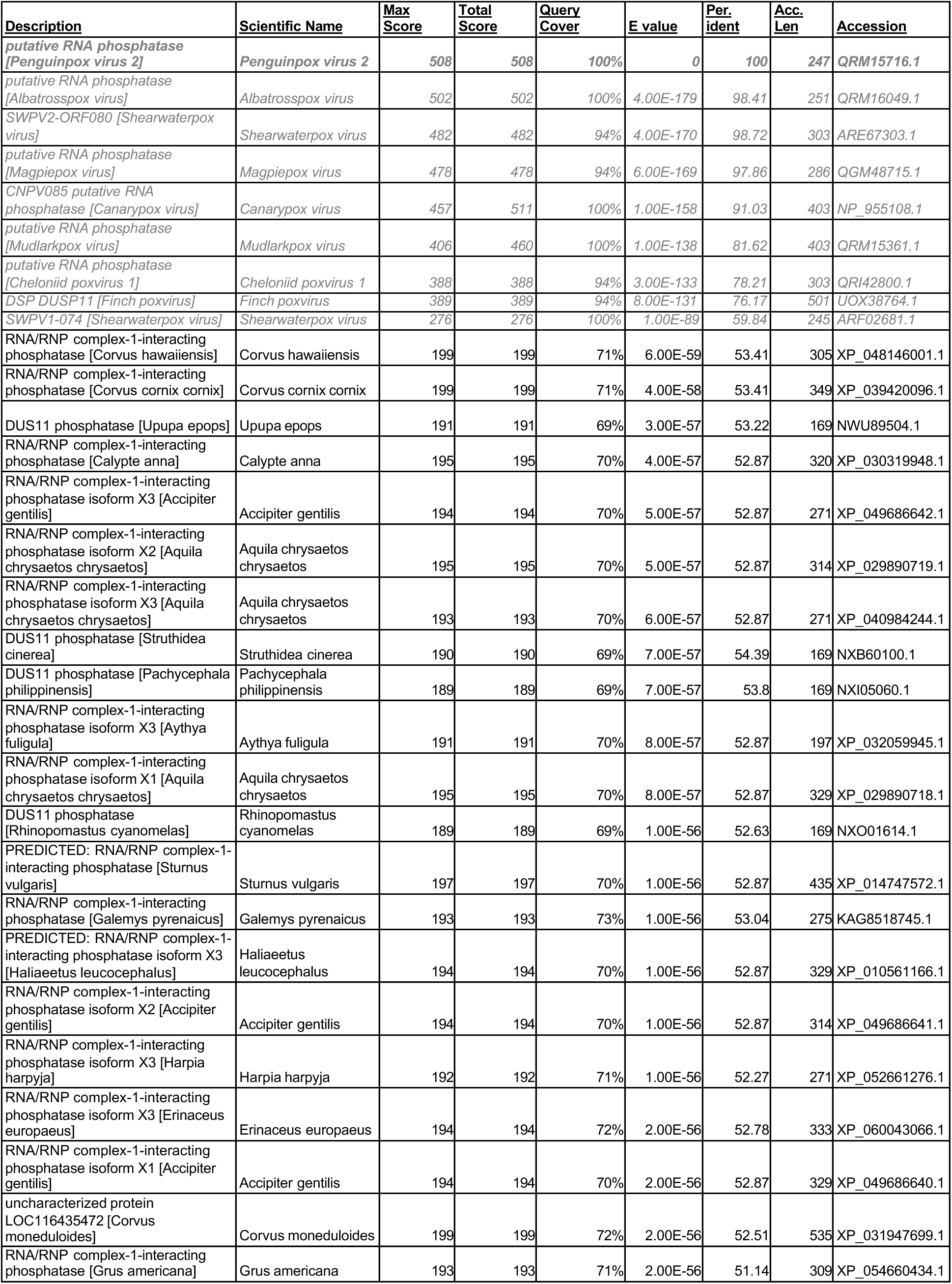
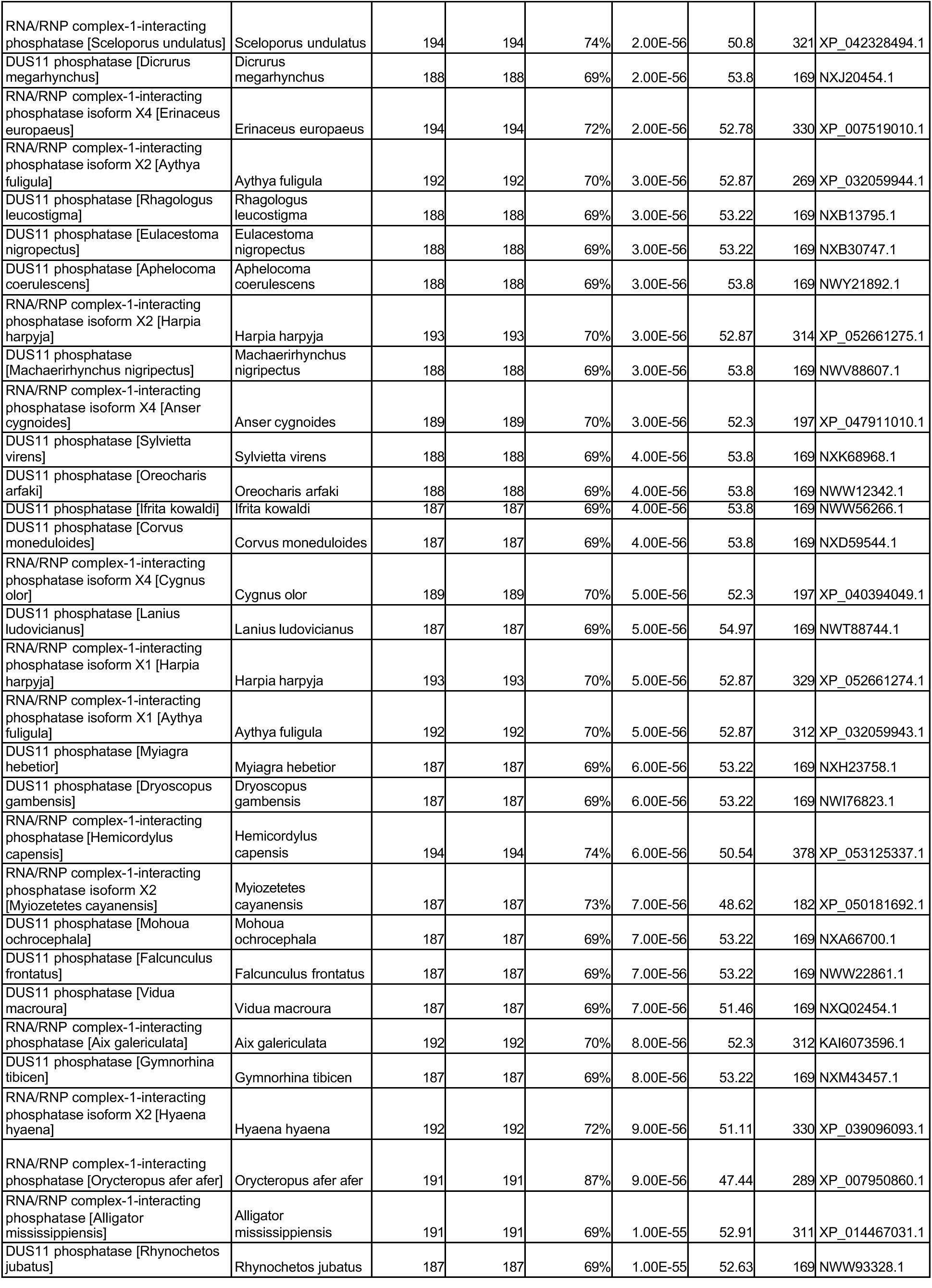
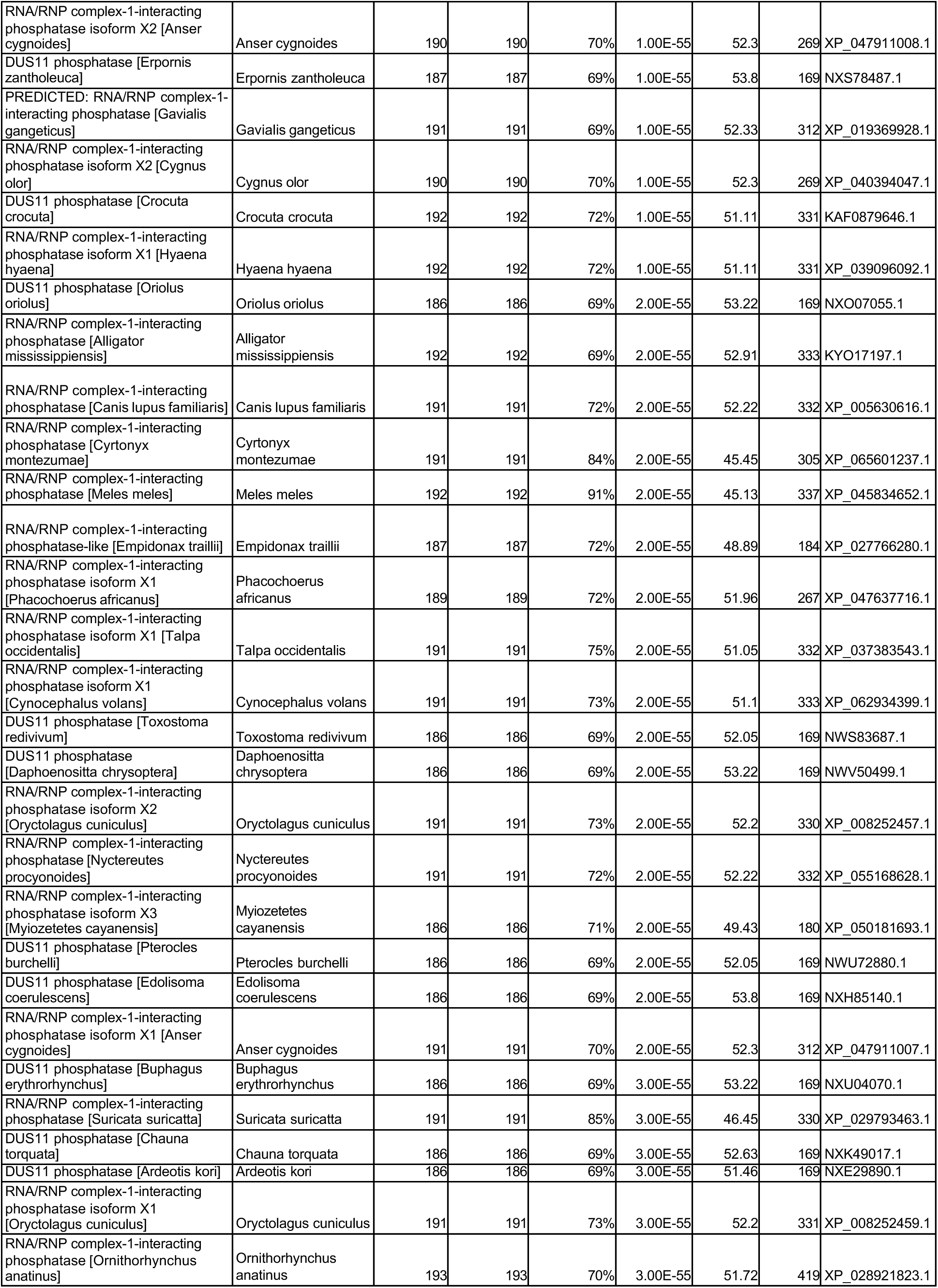
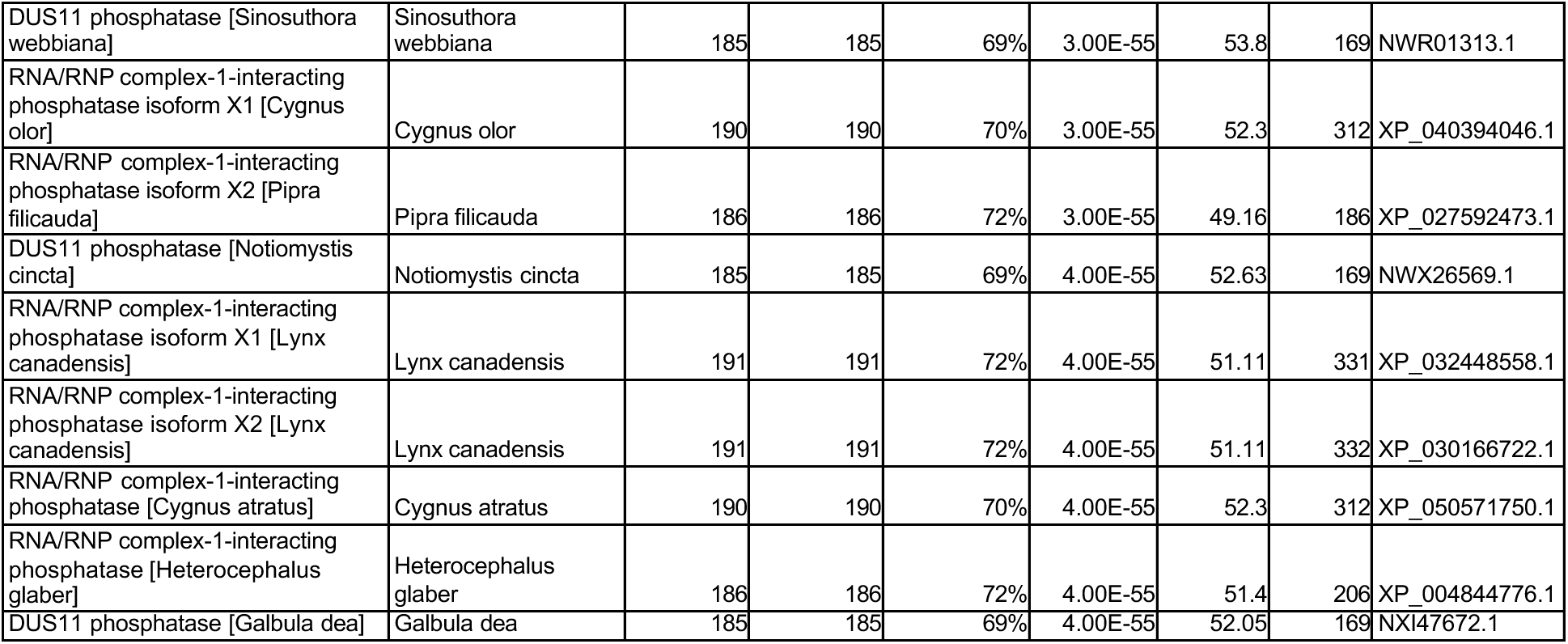
Blastp results using penguinpox virus 2 (PEPV2) viral DUSP11 as query (QRM15716.1)

**Table S8.**
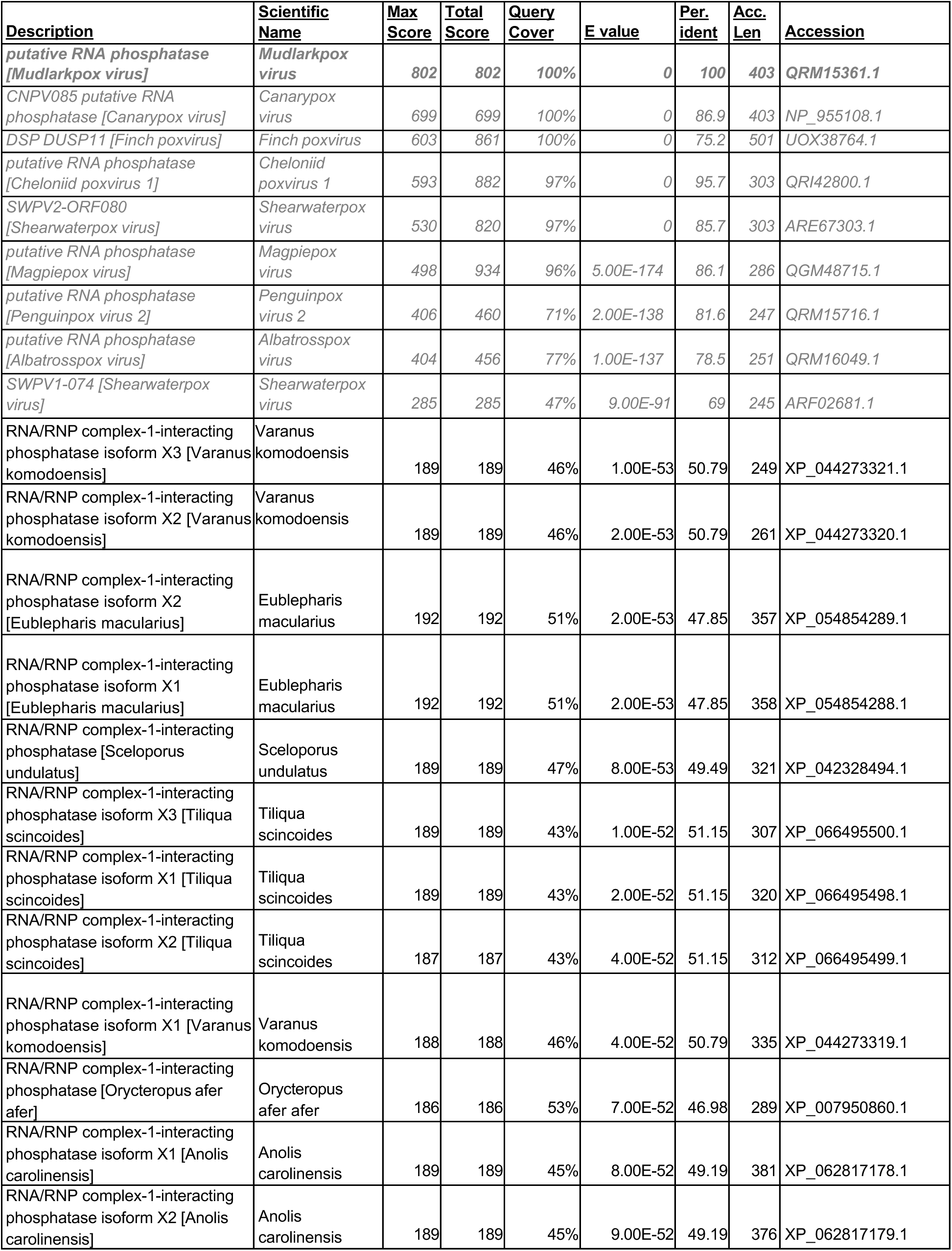
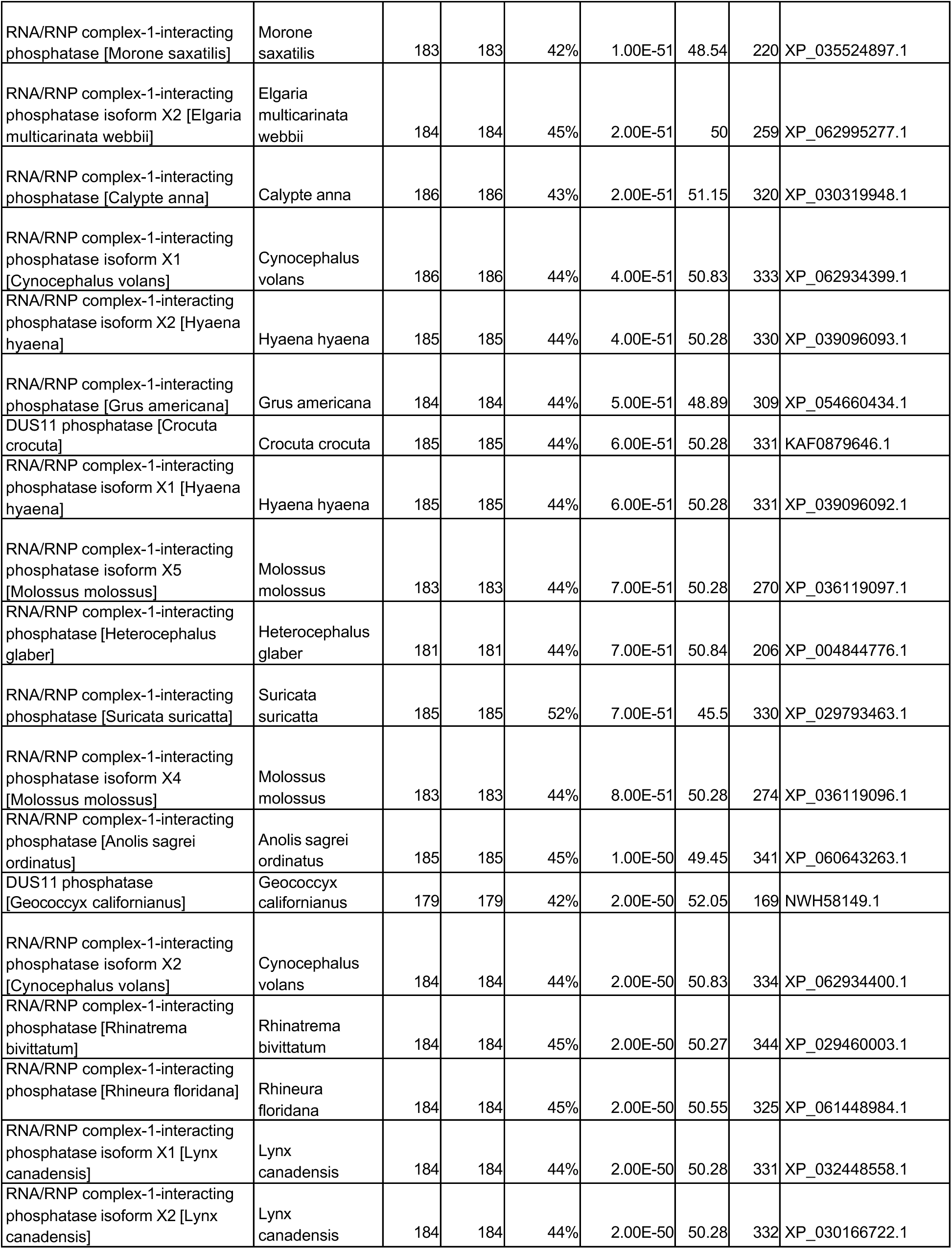
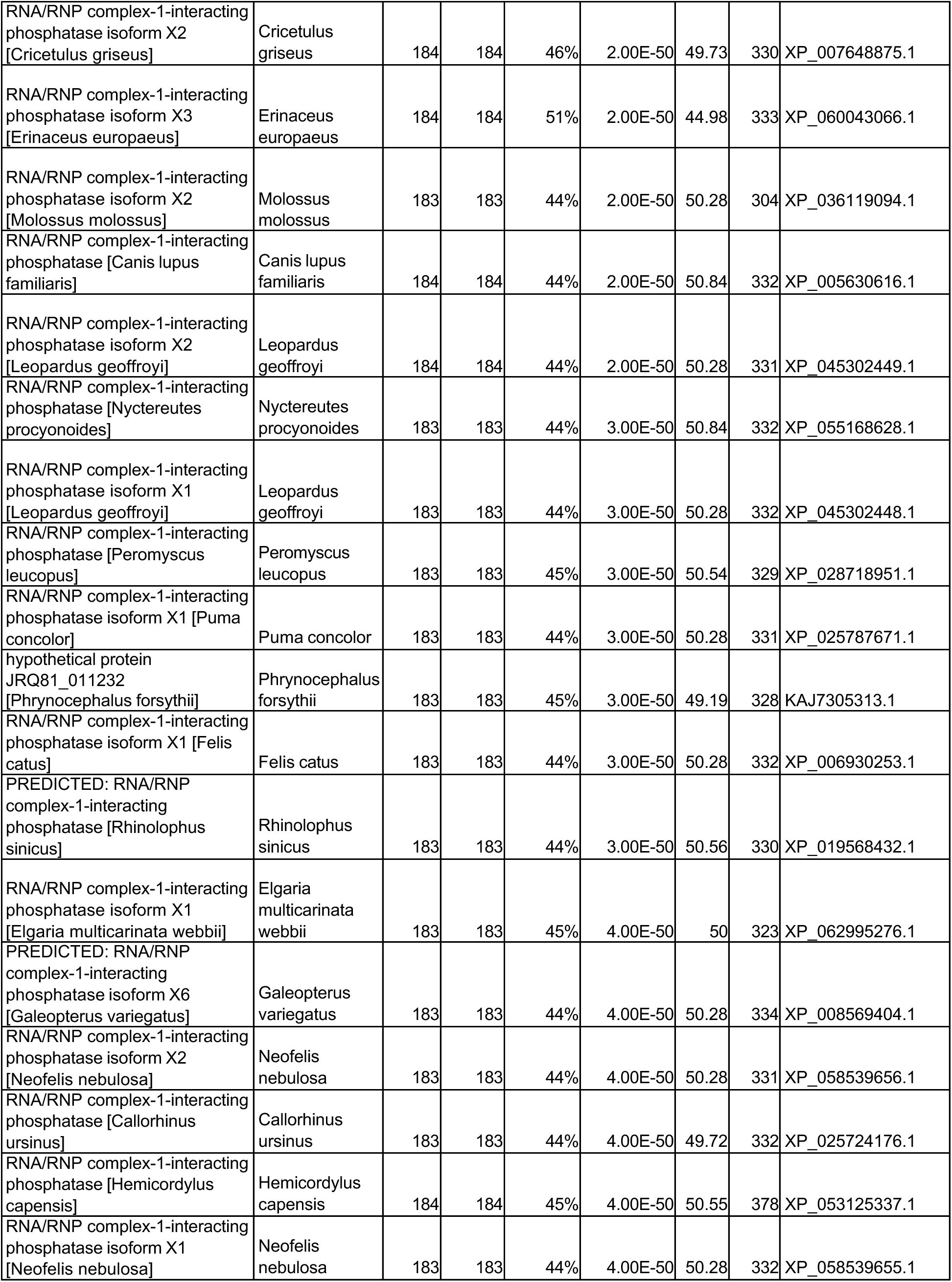
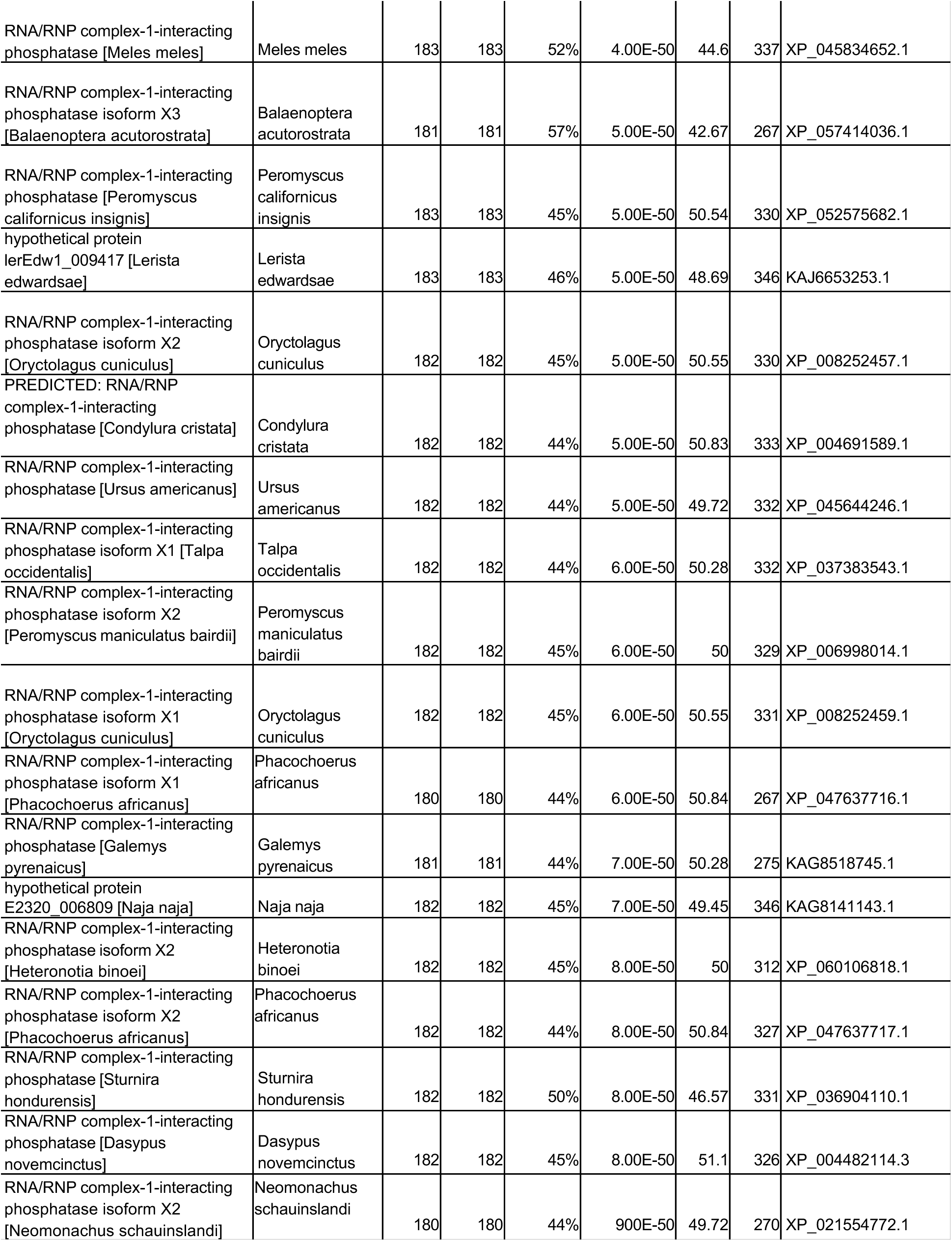
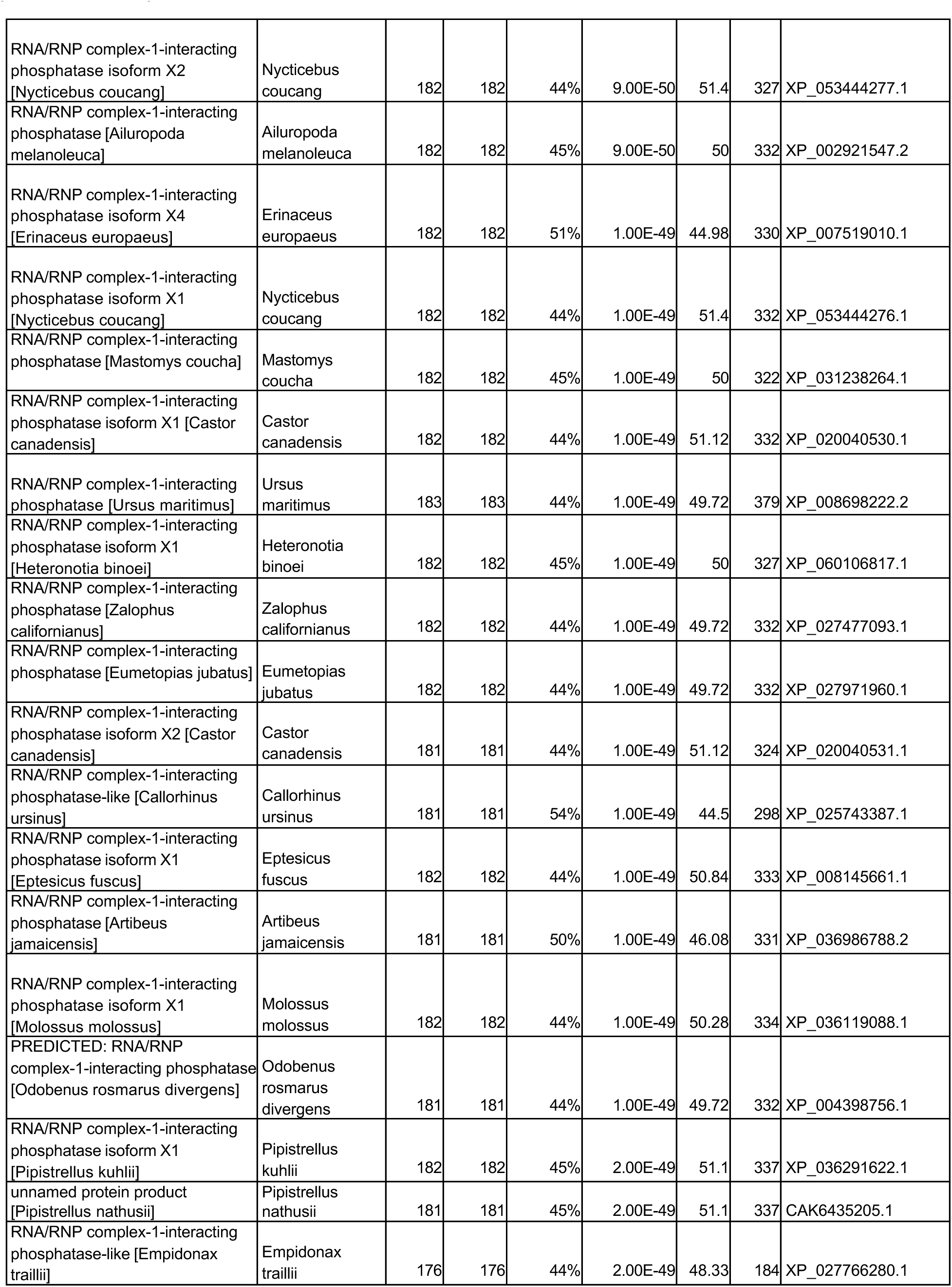
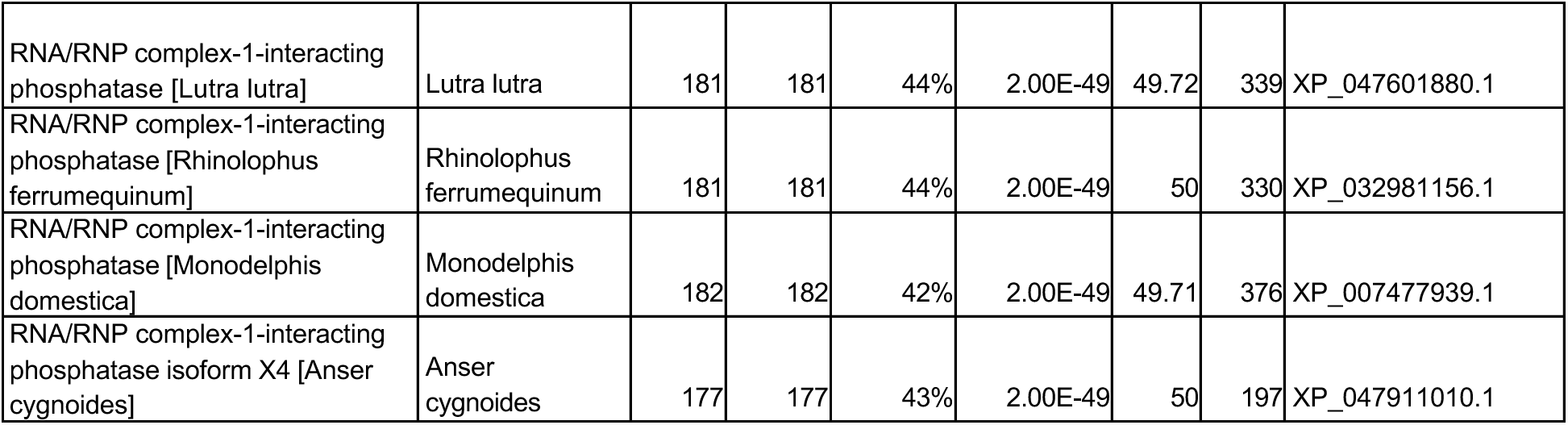
Blastp results using vDUSP11 for mudlarkpox virus as query (QRM15361.1)

**Table S9.**
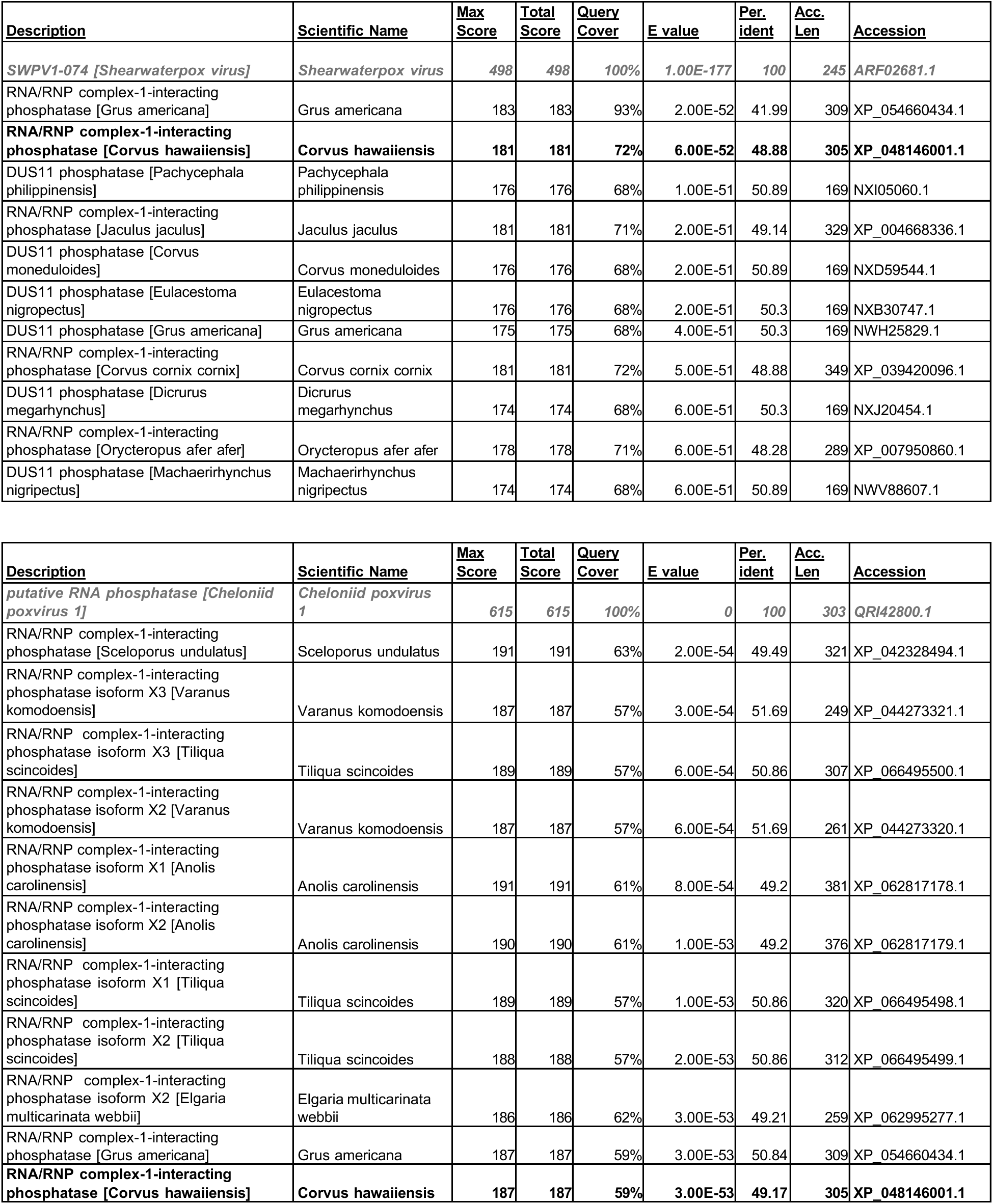
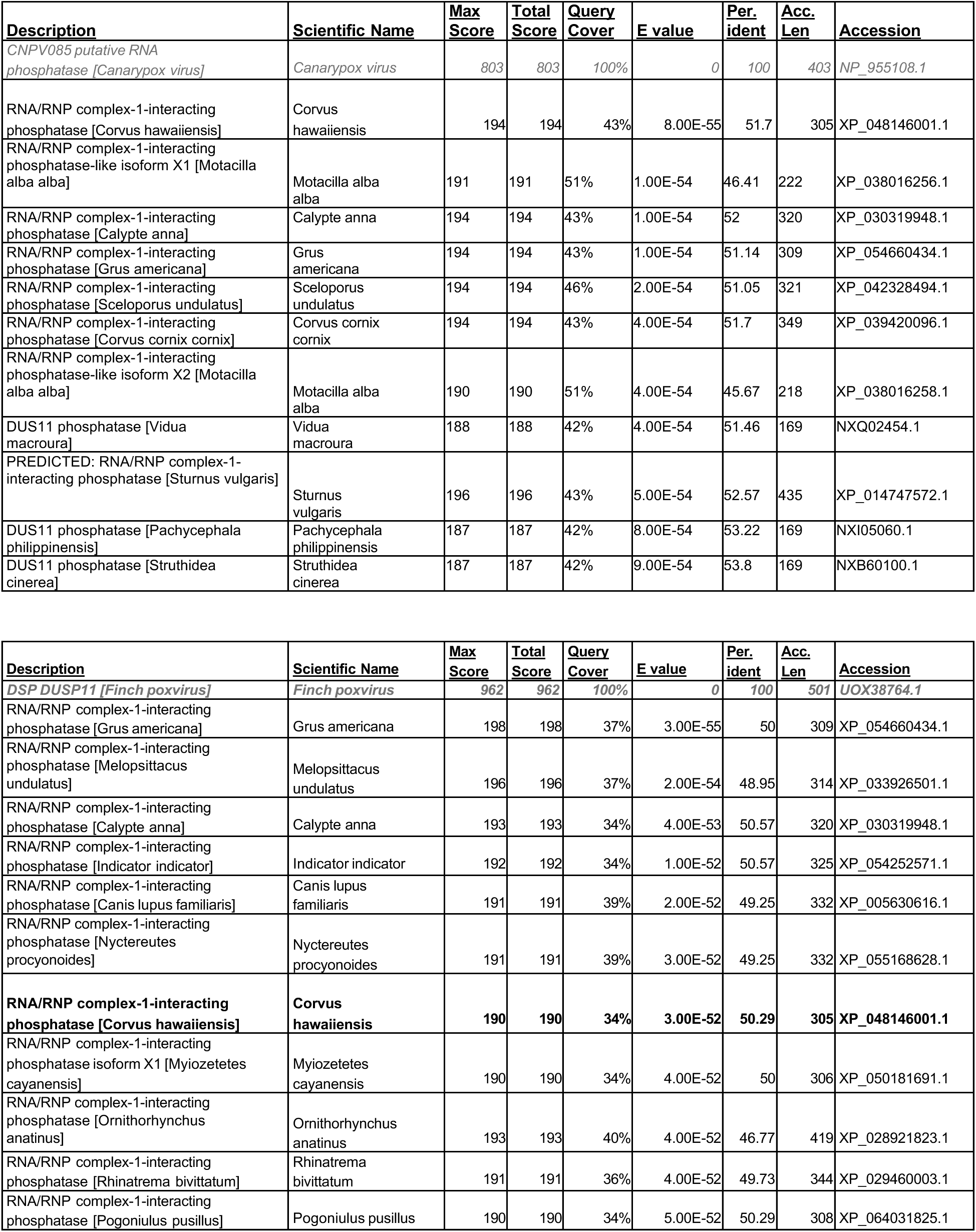
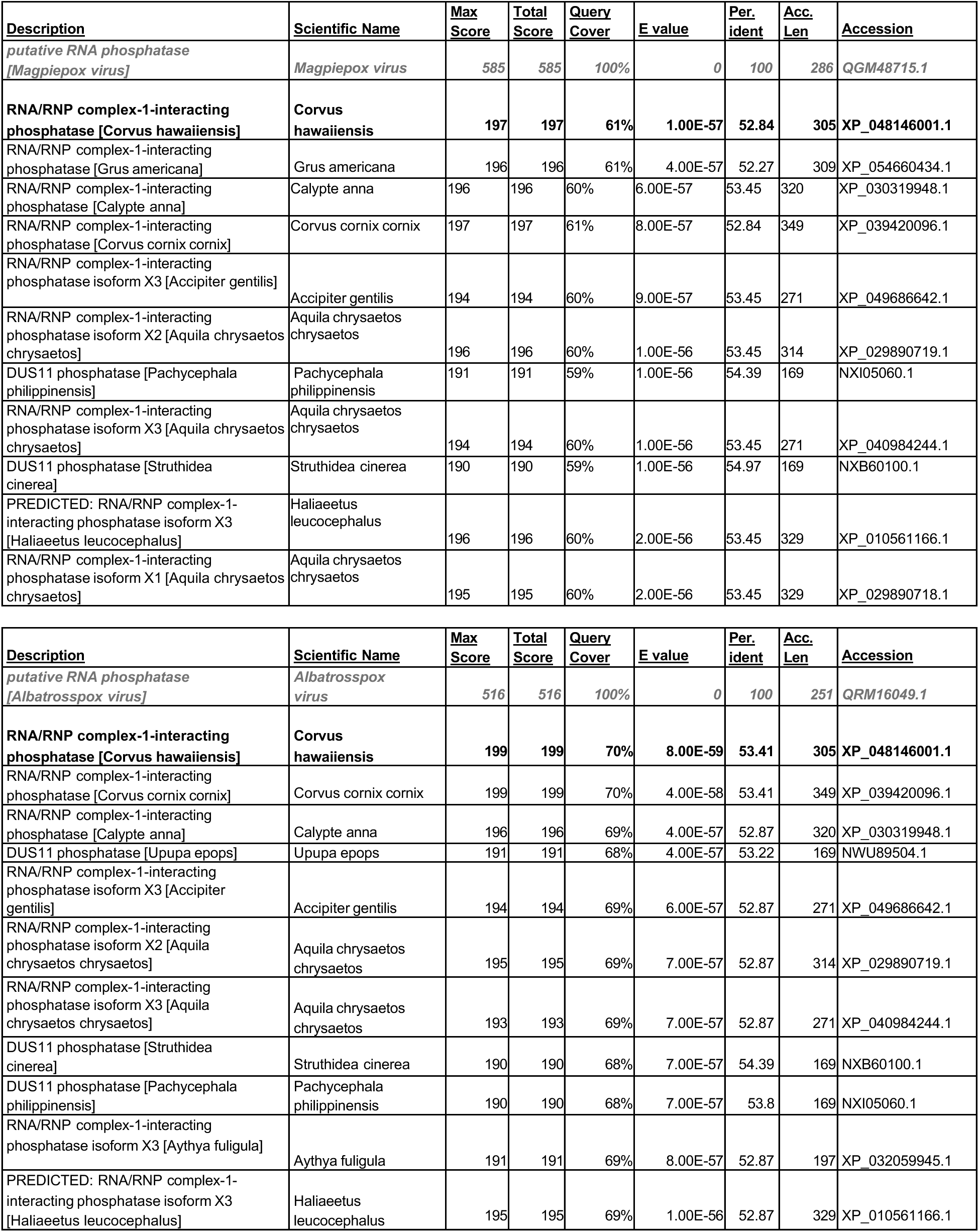
Reverse Blastp results using remaining vDUSP11 as query. vDUSP11 used as query is listed first in each table. First 10 results shown for each virus excluding other avipox vDUSP11s.

**Table S10.**
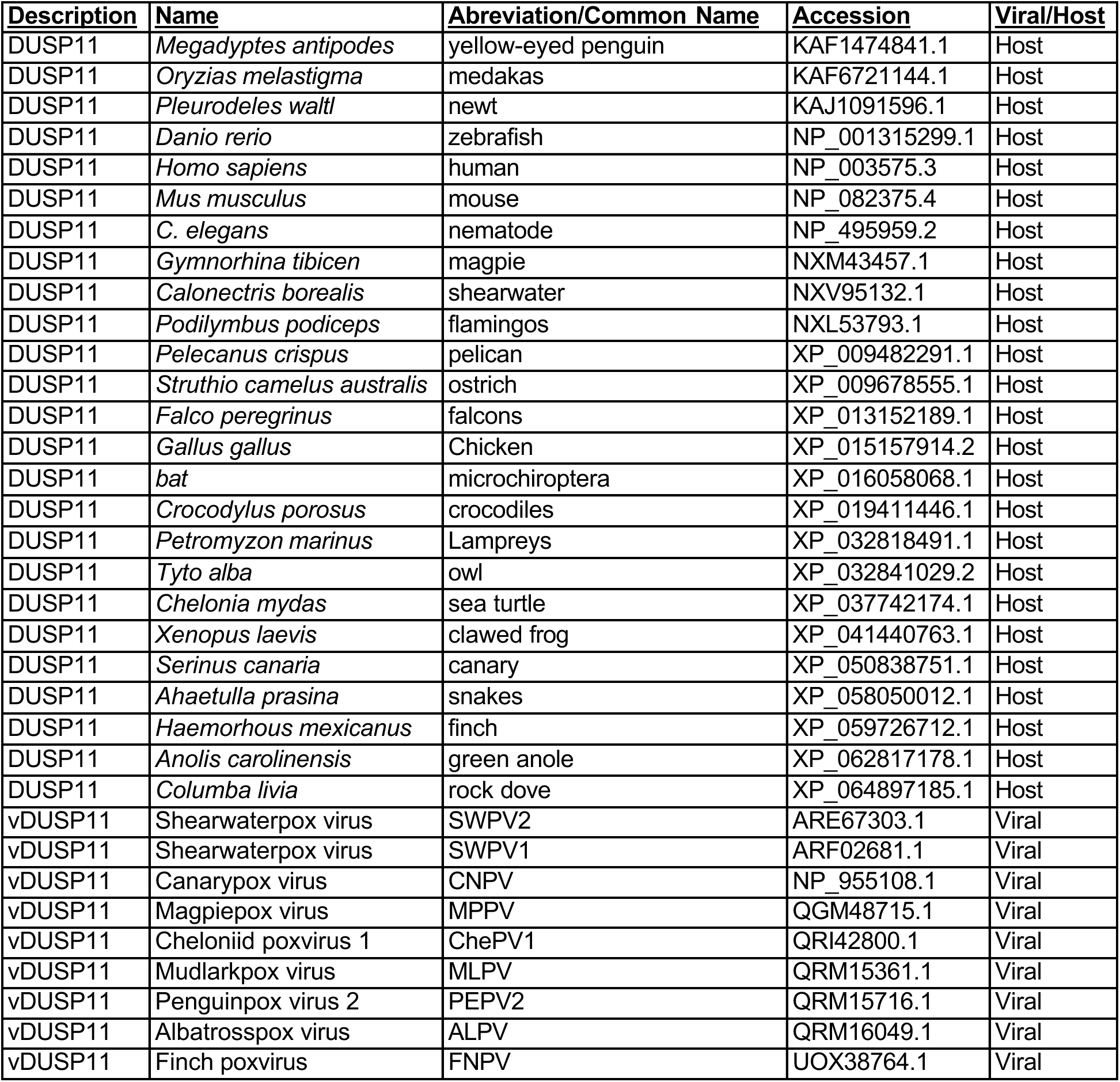
Accession IDs used to retrieve FASTA sequence for construction of phylogenetic tree.

