## Supplementary Materials for "Viral piracy of host RNA phosphatase DUSP11 by avipoxviruses"

**[1]Department of Molecular Biosciences**

**LaMontagne Center for Infectious Disease**

**The University of Texas at Austin, Austin, Texas, USA.**

**[2]Department of Immunology**

**UT. Southwestern Medical Center, Dallas, TX, USA.**

**Funding**

This work was supported by 1R35GM142689-04 to DCH and NIH 1R01AI134980 and a Burroughs Wellcome Investigators in Pathogenesis Award 1011070 to CSS.

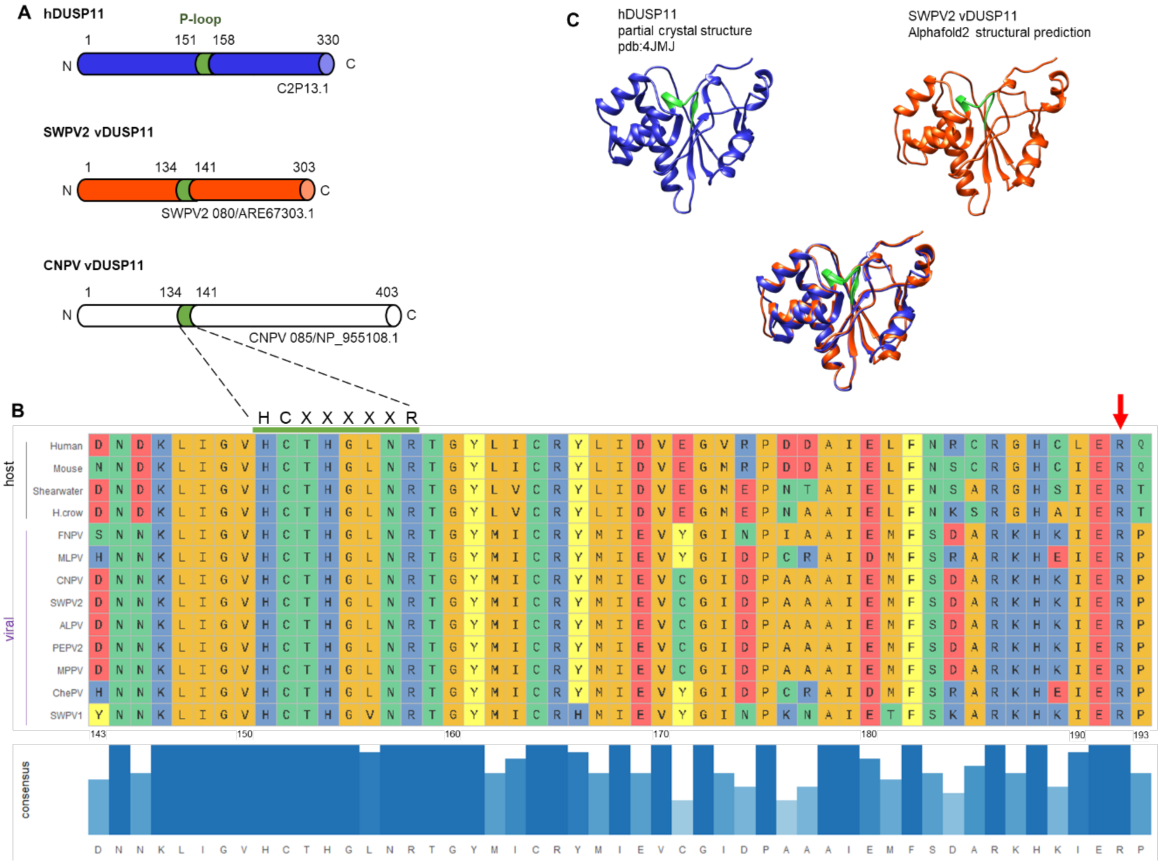


**Figure 1: Avipoxviruses and related unclassified avipox-adjacent poxviruses encode putative homolog of DUSP11.**

(A). Schematic of human *DUSP11* (hDUSP11) and select putative viral *DUSP11s* with predicted domains indicated. Colored domain indicates hDUSP11 P-loop with corresponding identical sequences present in putative viral DUSP11 homologs (vDUSP11). Putative vDUSP11 include shearwaterpox virus 2 vDUSP11 (SWPV2 vDUSP11) and canarypox virus vDUSP11 (CNPV vDUSP11). (B). Key catalytic residues are conserved between hDUSP11 and putative avipox vDUSP11. Multiple sequence comparison by log-expectation (Clustal) amino acid (AA) sequence alignment of human DUSP11, mouse DUSP11, *C. borealis* (Shearwater), and putative vDUSP11s from shearwaterpox virus 1 (SWPV1), finchpox virus (FNPV), mudlarkpox virus (MLPV), cheloniidpox virus (ChePV), canarypox virus (CNPV), penguinpox virus 2 (PEPV2), albatrosspox virus (ALPV), magpiepox virus (MPPV), and shearwaterpox virus 2 (SWPV2). A subset of AAs surrounding the hDUSP11 P-loop are shown. Amino acids are colored based on side-chain chemistry. Bar graph indicating relative sequence conservation with consensus amino acid sequence determined from listed sequences. Taller bar height and darker blue color correspond to relative correlation to consensus sequence. The numbering below the consensus sequence corresponds to hDUSP11 amino acid numbering. hDUSP11 residue R192 is indicated by a red arrow. (C). Partial crystal structure of hDUSP11 (pdb:4JMJ) (blue) and alphafold2 structural prediction of corresponding region from putative shearwaterpox virus 2 vDUSP11 (orange) with structural overlay. Key catalytic residues (P-loop) present in hDUSP11 are indicated in green, and corresponding residues are also indicated on the putative SWPV2 vDUSP11 structural prediction.

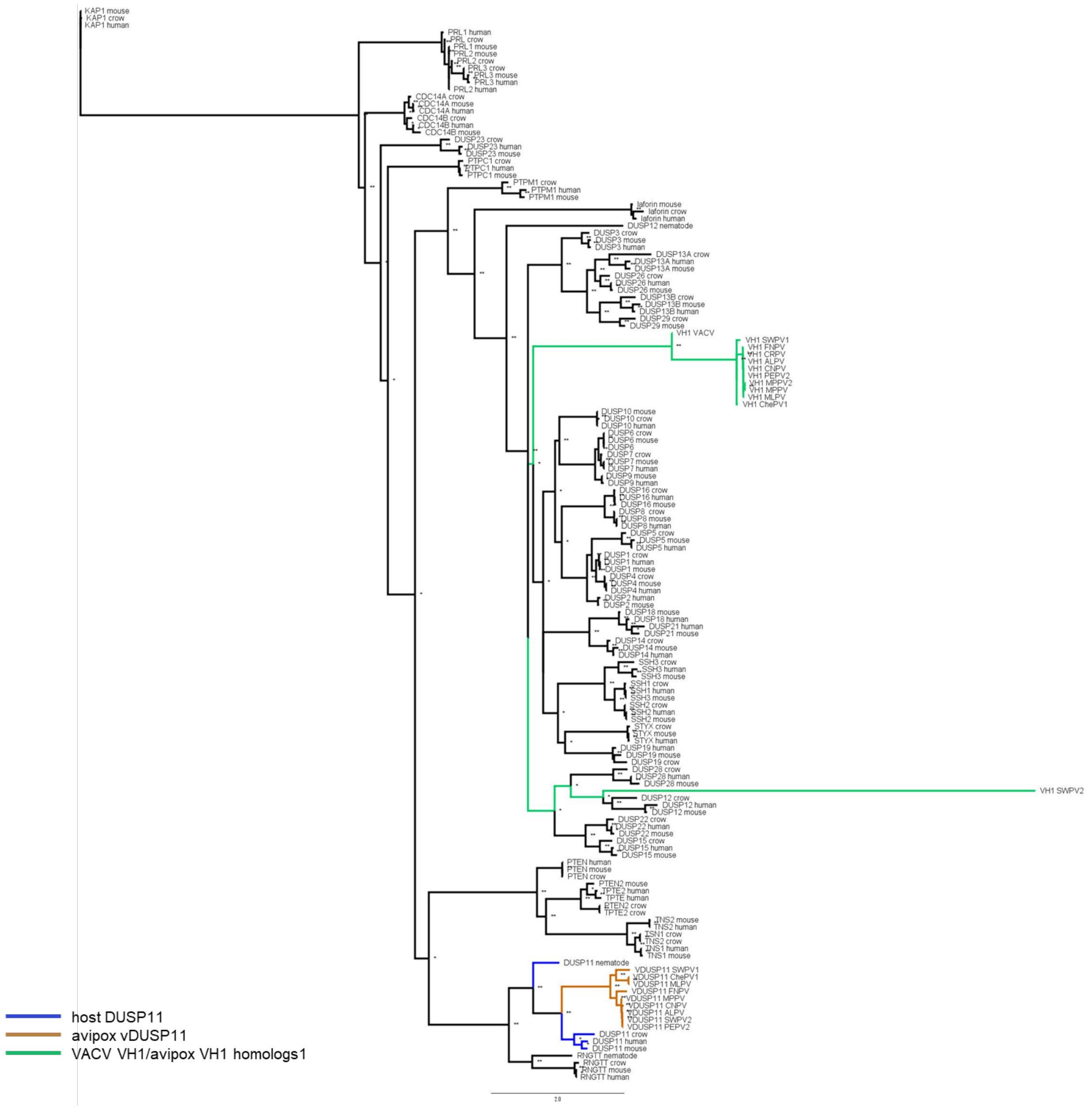


**Figure 2: Phylogenetic analysis supports putative vDUSP11 acquisition from host DUSP11 sequence.**

An inferred tree built using 150 DUSP amino acid (AA) sequences by maximum-likelihood analysis phylogenetic tree using PhyML (38). AA sequences for host DUSPs, poxviral putative vDUSP11s, and Vaccinia virus DUSP *H1L* protein phosphatase were aligned using Clustal Omega (Fig S3) (49). Clustal alignment was used to run PhyML analysis with LG +R model selected by SMS (50). Sequences were retrieved from the NCBI sequence database (37) and Uniprot (www.uniprot.org/) (Table S5). Putative APV/AdjPV vDUSP11s (orange) cluster with host DUSP11s (blue) and not with other related host protein DUSPs or Vaccinia VH1 or avipox homolog protein DUSP (green). 100 bootstrap replicates were performed; branch support >50% (*) or >70% (**) are indicated.

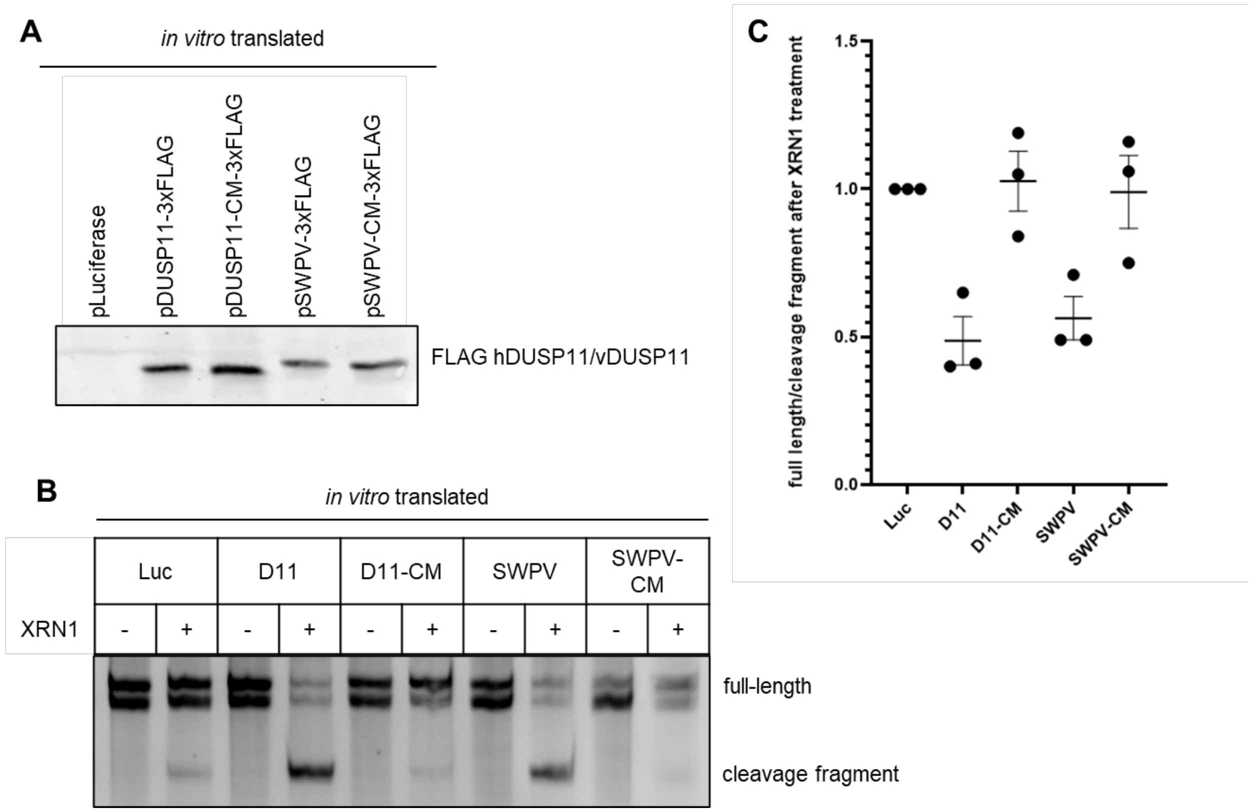


**Figure 3: vDUSP11 sensitizes HCV 5' UTR RNA to XRN-mediated degradation.**

(A). Confirmation of *in vitro* translated constructs via immunoblot analysis. Membrane was probed for FLAG-tagged proteins. Plasmid encoding luciferase was used for negative control reactions. (B). *In vitro* XRN susceptibility assay. *In vitro* transcribed HCV 5' UTR RNA was incubated with *in vitro* translated products from (A) (luciferase (Luc), hDUSP11 (D11), hDUSP11 catalytic mutant (D11-CM), shearwaterpox virus 2 vDUSP11 (SWPV) or shearwaterpox virus 2 vDUSP11 catalytic mutant (SWPV-CM)) and RNA was purified. Purified RNA was then subjected to treatment ± recombinant XRN1. Products were separated using urea PAGE and stained with EtBr. Migration of full-length HCV 5' UTR is indicated, which for unknown reasons migrates as a doublet, the position of the faster-migrating cleavage fragment is indicated. (C). Graphical representation of (B), displaying relative band intensity of HCV 5' UTR full-length to cleavage fragment following treatment with recombinant XRN1 (+XRN1/-XRN1). Values are normalized to the luciferase negative control. Data are derived from n = 3 independent replicates. In all panels, data are represented as mean ± SEM.

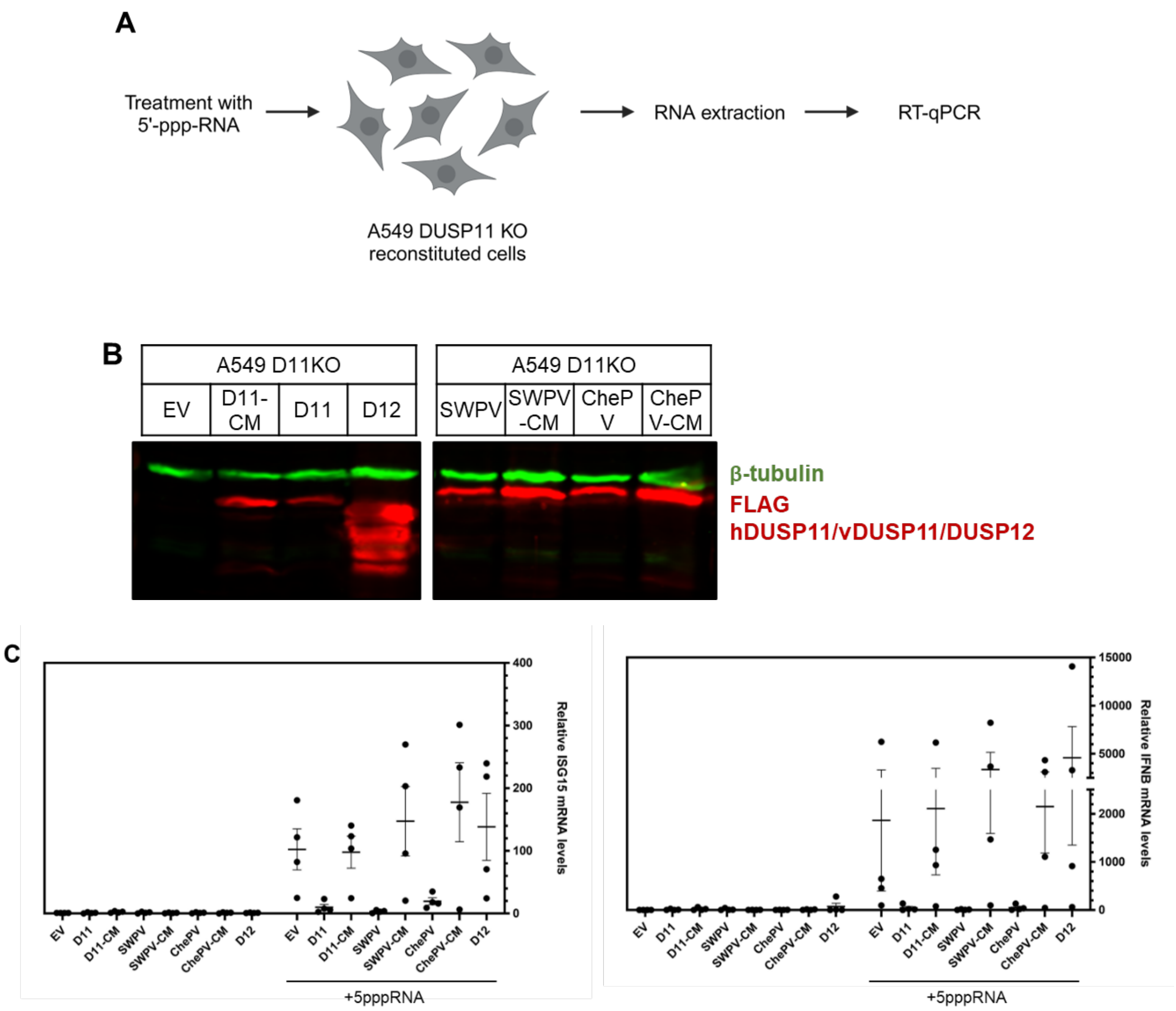


**Figure 4: vDUSP11 modulates immune activation in response to liposomal 5'-ppp-RNAs.**

(A). Schematic diagram of the 5'-triphosphate RNA (5'-ppp-RNA) transfection assay. A549 DUSP11 knockout (KO) reconstituted cells (12-well) were transfected with 5-10 ng of *in vitro* transcribed 5'-ppp-RNA for 18 hours followed by RT-qPCR to assay induction of ISGs. (B). Immunoblot analysis of A549 DUSP11 knockout (KO) cells transduced with pLenti empty vector (EV), pLenti hDUSP11-3xFLAG (D11), pLenti hDUSP11 catalytic mutant-3xFLAG (D11-CM), pLenti shearwaterpox virus 2 vDUSP11-3xFLAG (SWPV), pLenti shearwaterpox virus 2 vDUSP11 catalytic mutant-3xFLAG (SWPV-CM), pLenti cheloniidpox virus 1 vDUSP11-3xFLAG (ChePV), pLenti 3xFLAG-cheloniidpox virus 1 vDUSP11 catalytic mutant-3xFLAG (ChePV-CM), or negative control pLenti DUSP12-3xFLAG (D12). (C). RT-qPCR analysis of *ISG15* and *IFNB1* mRNA normalized to *GAPDH* mRNA in 5'-ppp-RNA transfected in A549 DUSP11 KO reconstituted cells. Results are represented relative to those of empty vector-expressing cells. Data are derived from n = 4 independent replicates in and are represented as mean ± SEM.

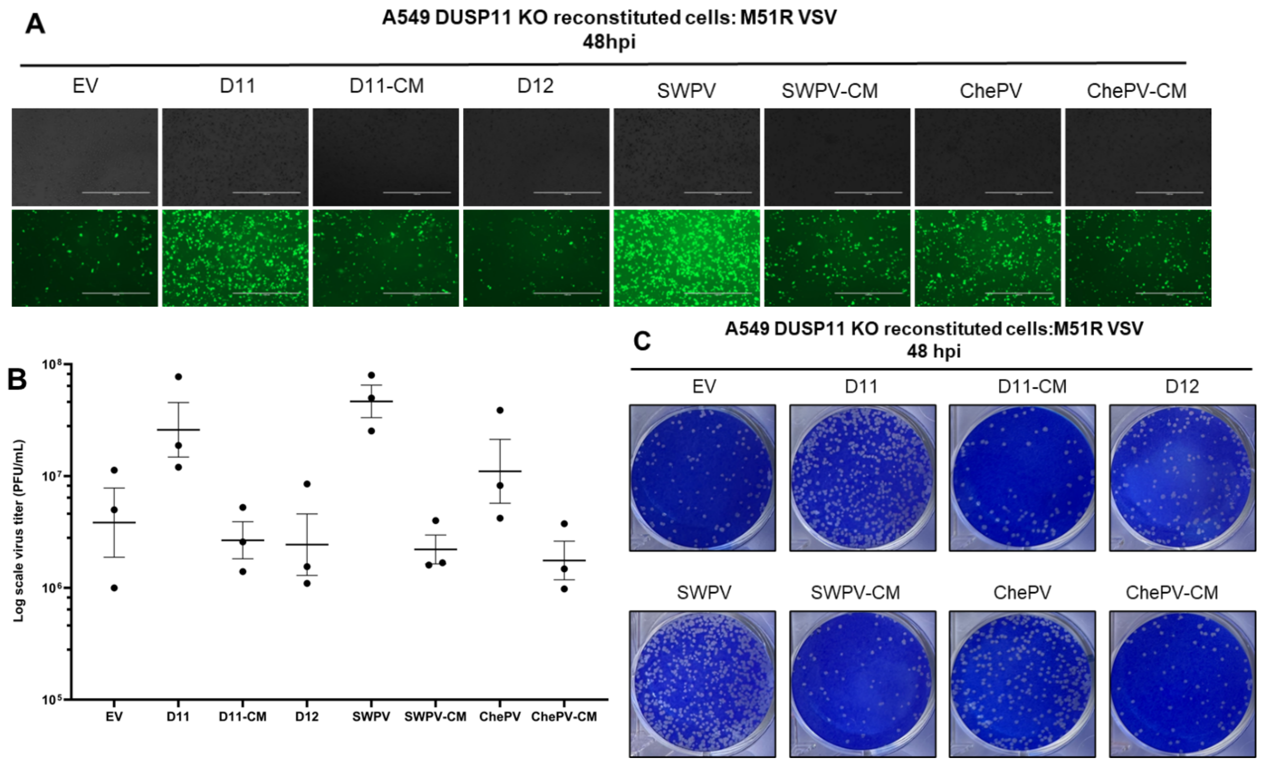


**Figure 5: vDUSP11 catalytic activity promotes VSV virus replication.**

(A). Representative GFP images of A549 DUSP11 knockout (KO) cells stably reconstituted with either empty vector (EV) plasmid, hDUSP11 (D11), hDUSP11 catalytic mutant (D11-CM), shearwaterpox virus 2 vDUSP11 (SWPV), shearwaterpox virus 2 vDUSP11 catalytic mutant (SWPV-CM), cheloniidpox virus vDUSP11 (ChePV), cheloniidpox virus vDUSP11 catalytic mutant (ChePV-CM), or protein phosphatase DUSP12 (D12) infected with GFP-M51R VSV at 48 hours post-infection (hpi). Cells were infected with GFP-M51R VSV at an MOI of 0.01 PFU/cell. (B). VSV viral titer of indicated A549 DUSP11 KO reconstituted cells at 48 hours post-infection (hpi). Cells were infected with GFP-M51R VSV at an MOI of 0.01 PFU/cell and virus supernatant was collected for plaque assay analysis. (C). Representative images of plaque assay analysis of virus supernatant at 48 hours post-infection. Data are derived from n = 3 independent replicates. Data are presented as mean ± SEM.

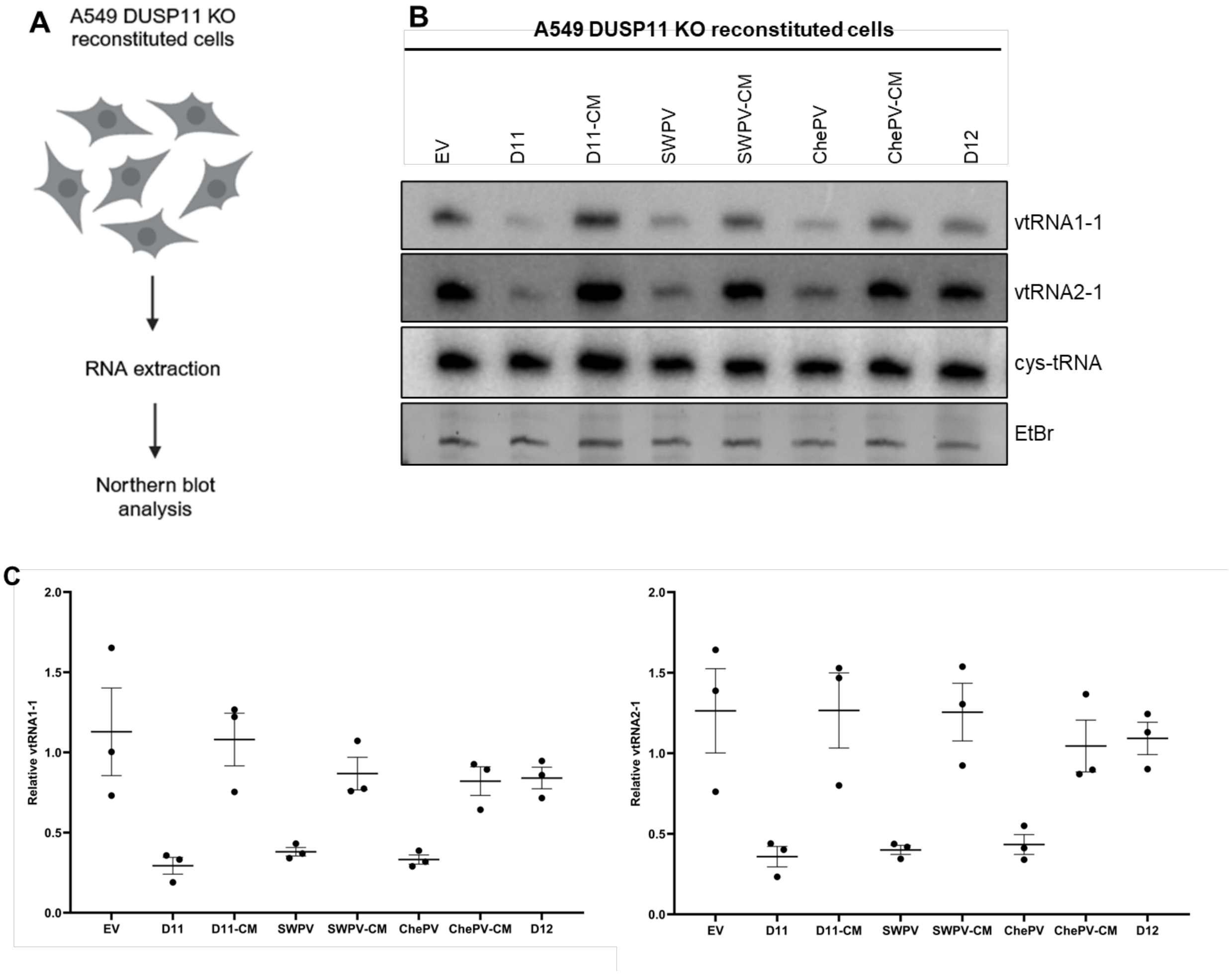


**Figure 6: vDUSP11 modulates steady-state RNA levels of endogenous RNAP III transcripts.**

(A) Schematic diagram of vtRNA Northern blot. RNA was collected from resting A549 DUSP11 KO cells stably reconstituted with empty vector (EV) plasmid, hDUSP11 (D11), hDUSP11 catalytic mutant (D11-CM), shearwaterpox virus 2 vDUSP11 (SWPV), shearwaterpox virus 2 vDUSP11 catalytic mutant (SWPV-CM), cheloniidpox virus vDUSP11 (ChePV), cheloniidpox virus vDUSP11 catalytic mutant (ChePV-CM), or negative control protein phosphatase DUSP12 (D12). Purified RNA was then subjected to Northern blot analysis. (B) Northern blot analysis of vtRNA1-1 and vtRNA2-1 using RNA from A549 DUSP11 KO reconstituted cells. (C) Graphical representation of relative band intensity of vtRNA1-1 and vtRNA2-1 normalized to the relative band intensity of the 5’ monophosphate control cysteine-tRNA. Data are derived from n =3 independent replicates. In all panels, data are represented as mean ± SEM.

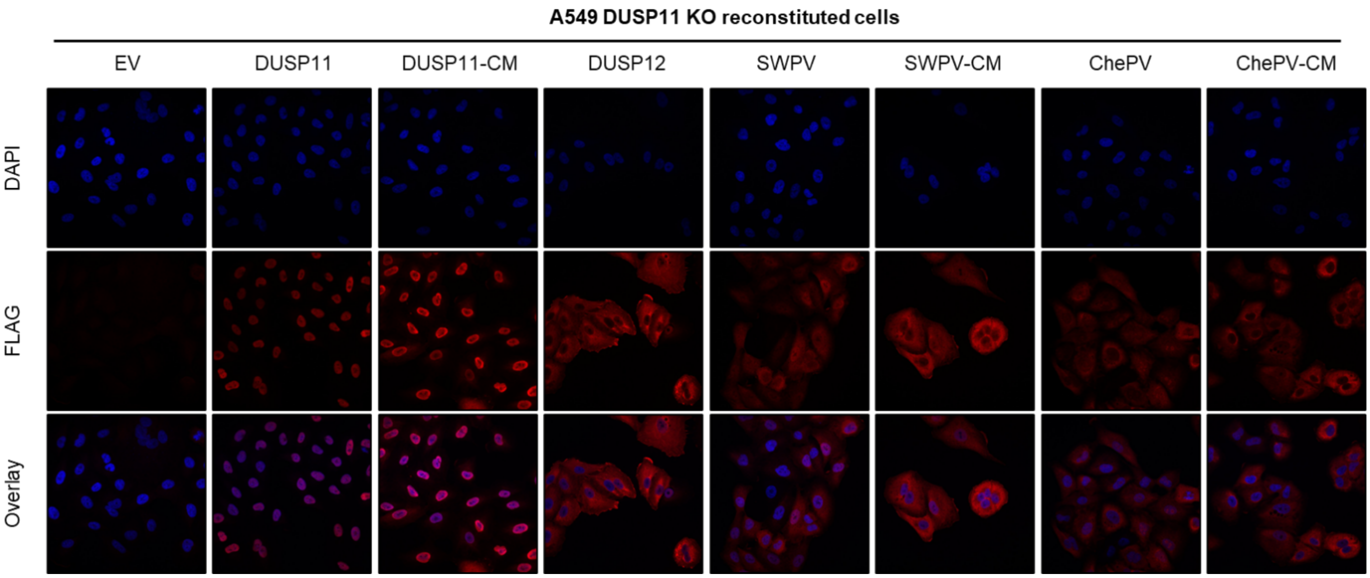


**Figure 7: vDUSP11s differ in subcellular localization compared to DUSP11.**

Representative confocal images of A549 DUSP11 KO cells stably expressing either an empty vector (EV) plasmid or individual 3xFLAG tagged proteins including hDUSP11 (D11), hDUSP11 catalytic mutant (D11-CM), SWPV2 vDUSP11 (SWPV), SWPV2-CM (SWPV-CM), ChePV1 vDUSP11 (ChePV), ChePV1 vDUSP11-CM (ChePV-CM), or negative control protein phosphatase DUSP12 (D12). Prolong™ Gold Antifade Mountant (Thermo Fisher Scientific) was used in slide preparation to visualize nuclei.

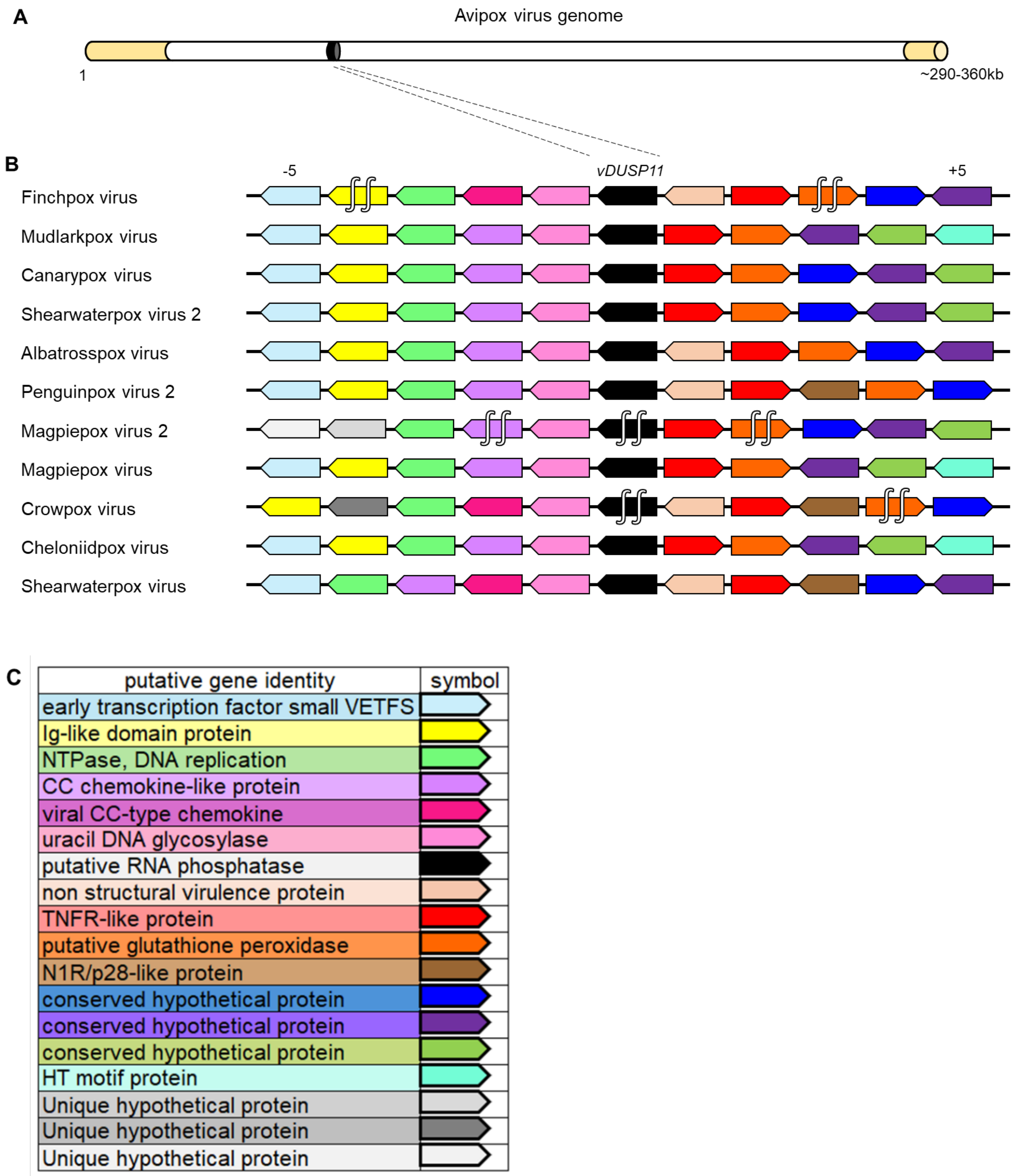


**Figure 8: A single acquisition event for vDUSP11 is indicated by synteny analysis.**

(A) Graphical representation of avipoxvirus genome using published information from the canarypox virus genome as a model (58). Inverted terminal repeats (ITRs) are indicated in yellow, relative genome position of *vDUSP11* indicated in black. (B) Analysis of upstream and downstream flanking genes indicated *vDUSP11*s (black) reside at similar genomic locations across poxvirus genomes. Protein homologs were determined using reciprocal BLAST hits for the 5 genes upstream (left of *vDUSP11*) and 5 genes downstream (right of *vDUSP11*) of *vDUSP11*. Homologous genes are indicated by shared color. Genes oriented to the right indicated ORFs on the top strand while genes oriented to the left indicated genes on the bottom strand. (C) Corresponding identity for genes indicated in (B). Putative gene identity was determined using NCBI database genome annotations (37). For genes lacking descriptive annotations, more detailed annotations based on identified homologous genes were used (Table S6). ∫∫ indicates *vDUSP11* ORFs disrupted by stop codons, resulting in multiple separate coding regions.

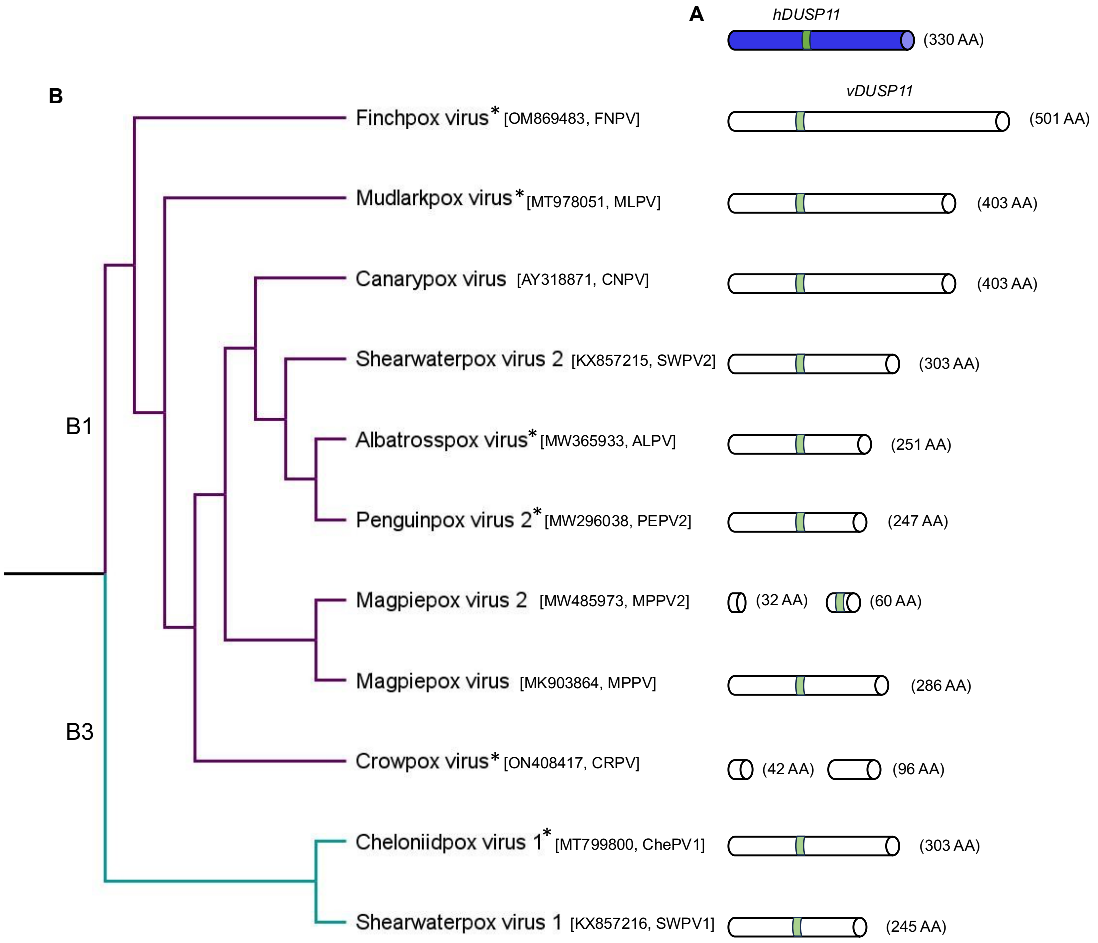


**Figure S1: A broad phylogenetic distribution of avipox and related avipox-adjacent poxviruses containing vDUSP11.**

(A) A graphical representation of *hDUSP11* with AA sequence length. (B) A cladogram built from published phylogenetic data (59), focusing on a subset of poxviruses encoding vDUSP11. Sub-clades (B1 and B3) are designated according to Gyuranecz et al. (2013) (60). Graphical representations of *vDUSP11* are to the right of each virus, demonstrating variations in vDUSP11 AA sequence length between viruses. Magpiepox virus 2 and crowpox virus encode truncated vDUSP11 as indicated. The presence of * indicates unclassified poxviruses. vDUSP11 p-loop indicated in green.

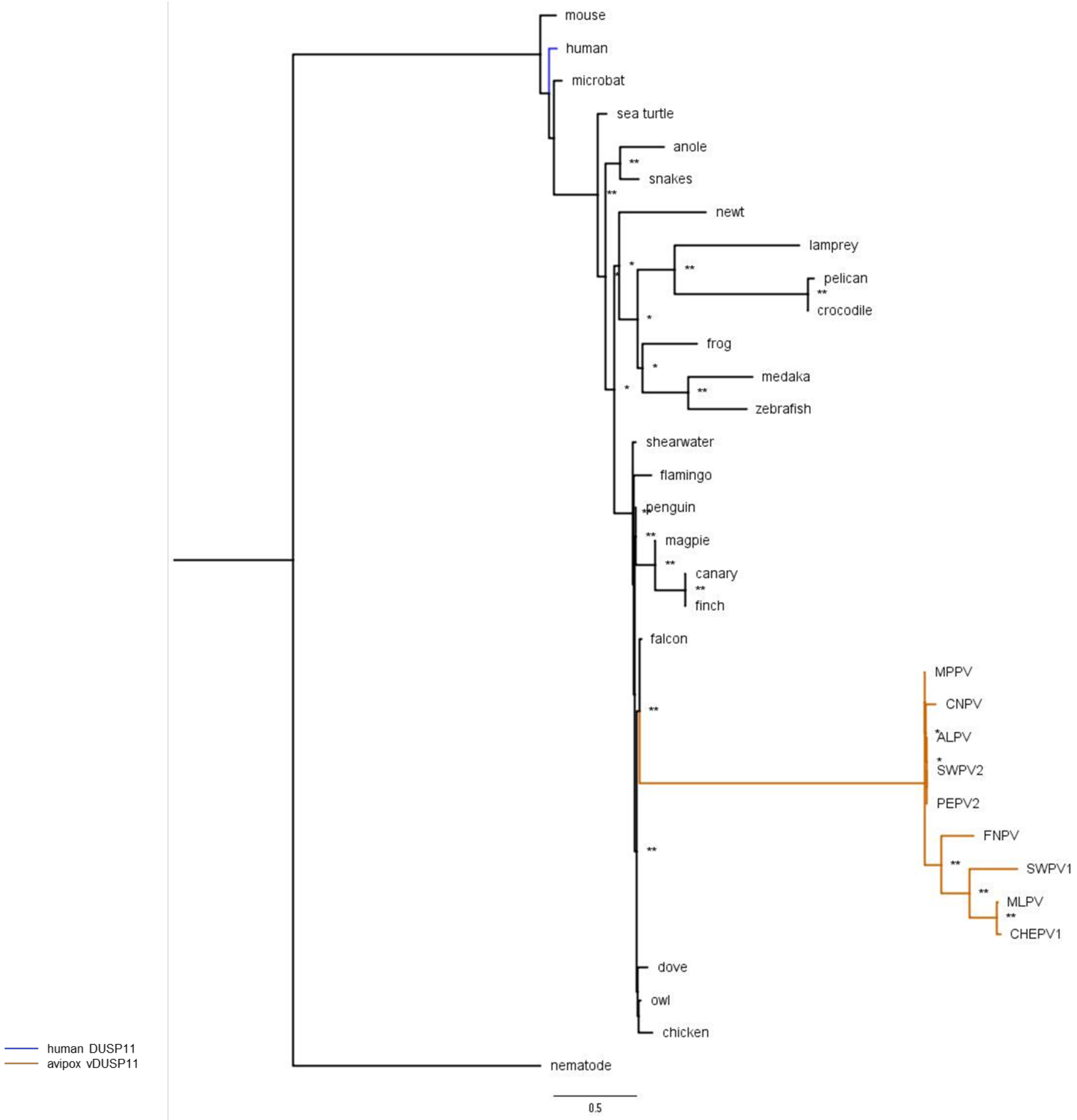


**Figure S2. An inferred phylogenetic tree highlighting likely avian origin of APV/AdjPV vDUSP11s.**

An inferred tree built using 33 host and viral DUSP11 amino acid (AA) sequences by maximum-likelihood analysis phylogenetic tree using PhyML (38). AA sequences for host DUSP11s and APV/AdjPV putative vDUSP11s were aligned using Clustal Omega (Fig S4) (49). Clustal alignment was used to run PhyML analysis with the Q.plant +G+I model selected by SMS (50). Sequences were retrieved from the NCBI sequence database (37) and Uniprot (www.uniprot.org/) (Table S10). Putative APV/AdjPV vDUSP11s (orange) cluster with avian host DUSP11s. 100 bootstrap replicates were performed; branch support >50% (*) or >70% (**) are indicated.

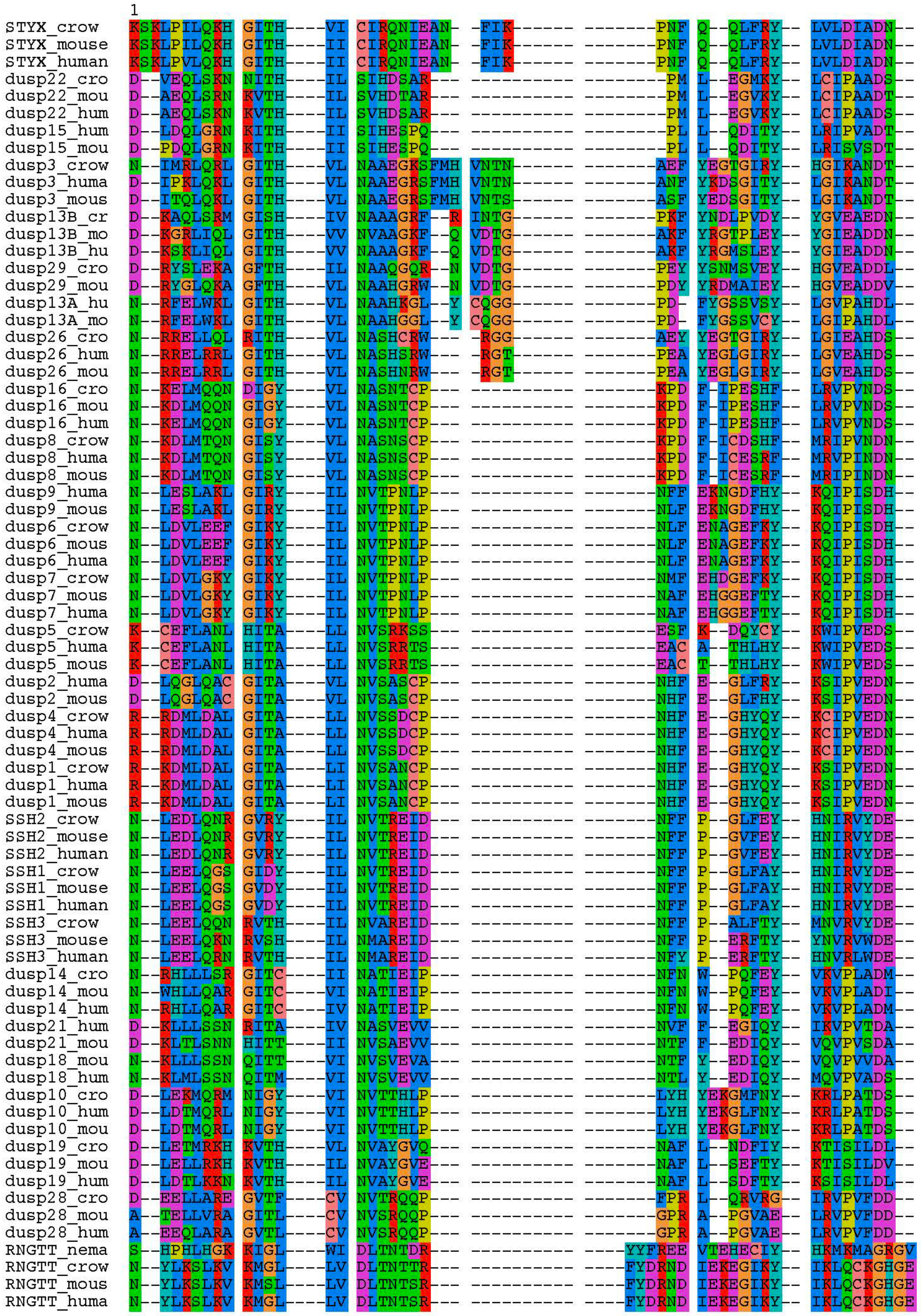


**Figure S3. Clustal Omega alignment of host/poxviral DUSP sequences (trimmed) used for PhyML analysis**

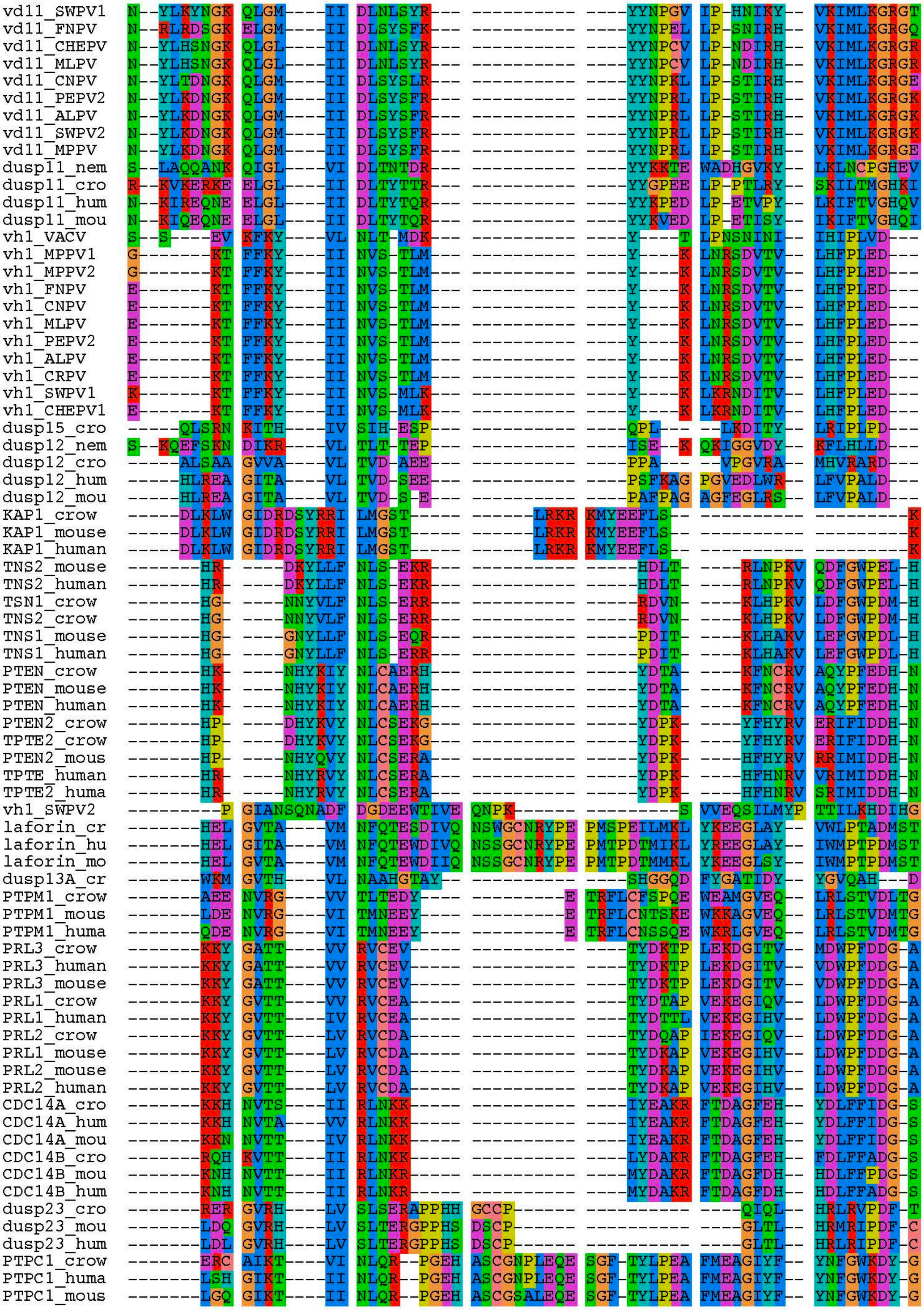


**Figure S3 cont. Clustal Omega alignment of host/poxviral DUSP sequences (trimmed) used for PhyML analysis**

```
               71
STYX_crow     -P-VENIIR- -FFPMTKEFI ---------- ---------- ---------- ---------- ---------D
STYX_mouse    -P-VENIIR- -FFPMTKEFI ---------- ---------- ---------- ---------- ---------D
STYX_human    -P-VENIIR- -FFPMTKEFI ---------- ---------- ---------- ---------- ---------D
dusp22_cro    -P-SQNLAR- -HFRESIKFI ---------- ---------- ---------- ---------- ---------H
dusp22_mou    -P-SQNLTR- -HFKESIKFI ---------- ---------- ---------- ---------- ---------H
dusp22_hum    -P-SQNLTR- -HFKESIKFI ---------- ---------- ---------- ---------- ---------H
dusp15_hum    -P-EVPIKK- -HFKECINFI ---------- ---------- ---------- ---------- ---------H
dusp15_mou    -P-EVPIKK- -HFKECVHFI ---------- ---------- ---------- ---------- ---------H
dusp3_crow    -Q-EFNLSR- -YFEEAADFI ---------- ---------- ---------- ---------- ---------E
dusp3_huma    -Q-EFNLSA- -YFERAADFI ---------- ---------- ---------- ---------- ---------D
dusp3_mous    -Q-EFNLSA- -YFERATDFI ---------- ---------- ---------- ---------- ---------D
dusp13B_cr    -P-NFDLSI- -HFYPVAQYI ---------- ---------- ---------- ---------- ---------R
dusp13B_mo    -P-FFDLSV- -HFLPVARYI ---------- ---------- ---------- ---------- ---------R
dusp13B_hu    -P-FFDLSV- -YFLPVARYI ---------- ---------- ---------- ---------- ---------R
dusp29_cro    -P-TFNLSQ- -FFYSASKFI ---------- ---------- ---------- ---------- ---------D
dusp29_mou    -P-TFDLSI- -FFYSAAAFI ---------- ---------- ---------- ---------- ---------D
dusp13A_hu    -P-DFDISA- -YFSSAADFI ---------- ---------- ---------- ---------- ---------H
dusp13A_mo    -P-DFNISP- -YFSSAADFI ---------- ---------- ---------- ---------- ---------H
dusp26_cro    -P-SFDMSP- -YFYPAADFI ---------- ---------- ---------- ---------- ---------H
dusp26_hum    -P-AFDMSI- -HFQTAADFI ---------- ---------- ---------- ---------- ---------H
dusp26_mou    -P-AFDMSI- -HFQTAADFI ---------- ---------- ---------- ---------- ---------H
dusp16_cro    -F-CEKILP- -WLDKSVDFI ---------- ---------- ---------- ---------- ---------E
dusp16_mou    -F-CEKILP- -WLDKSVDFI ---------- ---------- ---------- ---------- ---------E
dusp16_hum    -F-CEKILP- -WLDKSVDFI ---------- ---------- ---------- ---------- ---------E
dusp8_crow    -Y-CEKLLP- -WLDKSIEFI ---------- ---------- ---------- ---------- ---------D
dusp8_huma    -Y-CEKLLP- -WLDKSIEFI ---------- ---------- ---------- ---------- ---------D
dusp8_mous    -Y-CEKLLP- -WLDKSIEFI ---------- ---------- ---------- ---------- ---------D
dusp9_huma    -W-SQNLSR- -FFPEAIEFI ---------- ---------- ---------- ---------- ---------D
dusp9_mous    -W-SQNLSQ- -FFPEAIAFI ---------- ---------- ---------- ---------- ---------D
dusp6_crow    -W-SQNLSQ- -FFPEAISFI ---------- ---------- ---------- ---------- ---------D
dusp6_mous    -W-SQNLSQ- -FFPEAISFI ---------- ---------- ---------- ---------- ---------D
dusp6_huma    -W-SQNLSQ- -FFPEAISFI ---------- ---------- ---------- ---------- ---------D
dusp7_crow    -W-SQNLSQ- -FFPEAIAFI ---------- ---------- ---------- ---------- ---------D
dusp7_mous    -W-SQNLSQ- -FFPEAISFI ---------- ---------- ---------- ---------- ---------D
dusp7_huma    -W-SQNLSQ- -FFPEAISFI ---------- ---------- ---------- ---------- ---------D
dusp5_crow    -H-TADISS- -HFQEAIDFI ---------- ---------- ---------- ---------- ---------D
dusp5_huma    -H-TADISS- -HFQEAIDFI ---------- ---------- ---------- ---------- ---------D
dusp5_mous    -H-TADISS- -HFQEAIDFI ---------- ---------- ---------- ---------- ---------D
dusp2_huma    -Q-MVEISA- -WFQEAIGFI ---------- ---------- ---------- ---------- ---------D
dusp2_mous    -Q-MVEISA- -WFQEAISFI ---------- ---------- ---------- ---------- ---------D
dusp4_crow    -H-KADISS- -WFMEAIEYI ---------- ---------- ---------- ---------- ---------D
dusp4_huma    -H-KADISS- -WFMEAIEYI ---------- ---------- ---------- ---------- ---------D
dusp4_mous    -H-KADISS- -WFMEAIEYI ---------- ---------- ---------- ---------- ---------D
dusp1_crow    -H-KADISS- -WFNEAIDFI ---------- ---------- ---------- ---------- ---------D
dusp1_huma    -H-KADISS- -WFNEAIDFI ---------- ---------- ---------- ---------- ---------D
dusp1_mous    -H-KADISS- -WFNEAIDFI ---------- ---------- ---------- ---------- ---------D
SSH2_crow     -E-ATDLLA- -YWNDTYKFI ---------- ---------- ---------- ---------- ---------S
SSH2_mouse    -E-ATDLLA- -YWNDTYKFI ---------- ---------- ---------- ---------- ---------S
SSH2_human    -E-ATDLLA- -YWNDTYKFI ---------- ---------- ---------- ---------- ---------S
SSH1_crow     -E-TTDLLA- -HWNEAYHFI ---------- ---------- ---------- ---------- ---------N
SSH1_mouse    -E-TTDLLA- -HWNEAYHFI ---------- ---------- ---------- ---------- ---------N
SSH1_human    -E-TTDLLA- -HWNEAYHFI ---------- ---------- ---------- ---------- ---------N
SSH3_crow     -E-AAELLP- -HWNDTFLFL ---------- ---------- ---------- ---------- ---------S
SSH3_mouse    -E-SAQLLP- -HWKETHRFI ---------- ---------- ---------- ---------- ---------E
SSH3_human    -E-SAQLLP- -HWKETHRFI ---------- ---------- ---------- ---------- ---------E
dusp14_cro    -P-TAPISL- -YFDSVADKI ---------- ---------- ---------- ---------- ---------H
dusp14_mou    -P-HAPIRL- -YFDTVADKI ---------- ---------- ---------- ---------- ---------H
dusp14_hum    -P-HAPIGL- -YFDTVADKI ---------- ---------- ---------- ---------- ---------H
dusp21_hum    -R-DSRLYD- -FFDPIADLI ---------- ---------- ---------- ---------- ---------H
dusp21_mou    -P-NSYLYD- -FFDPIADHI ---------- ---------- ---------- ---------- ---------H
dusp18_mou    -P-VARLSN- -FFDSVADRI ---------- ---------- ---------- ---------- ---------H
dusp18_hum    -P-NSRLCD- -FFDPIADHI ---------- ---------- ---------- ---------- ---------H
dusp10_cro    -N-KQNLRQ- -YFEEAFEFI ---------- ---------- ---------- ---------- ---------E
dusp10_hum    -N-KQNLRQ- -YFEEAFEFI ---------- ---------- ---------- ---------- ---------E
dusp10_mou    -N-KQNLRQ- -YFEEAFEFI ---------- ---------- ---------- ---------- ---------E
dusp19_cro    -P-ETDITS- -YFPECFEFI ---------- ---------- ---------- ---------- ---------E
dusp19_mou    -P-ETNILS- -YFPECFEFI ---------- ---------- ---------- ---------- ---------E
dusp19_hum    -P-ETNILS- -YFPECFEFI ---------- ---------- ---------- ---------- ---------E
dusp28_cro    -P-AEDLYR- -YFEPCGAAI ---------- ---------- ---------- ---------- ---------E
dusp28_mou    -P-AEDLLT- -HLEPTCAAM ---------- ---------- ---------- ---------- ---------E
dusp28_hum    -P-AEDLLA- -HLEPTCAAM ---------- ---------- ---------- ---------- ---------E
RNGTT_nema    SPTQEDTDN- -FIKLVQEFH ---------- ---------- ---------- ---------- ---------K
RNGTT_crow    CPTPENTET- -FIRVCEHFS ---------- ---------- ---------- ---------- ---------D
RNGTT_mous    CPTTENTET- -FIRLCERFN ---------- ---------- ---------- ---------- ---------E
RNGTT_huma    CPTTENTET- -FIRLCERFN ---------- ---------- ---------- ---------- ---------E
vd11_SWPV1    IPPIDEVIR- -FKFEVDNFL ---------- ---------- ---------- ---------- ---------I
vd11_FNPV     IPTIDDMNR- -FSLEVNRFL ---------- ---------- ---------- ---------- ---------E
vd11_CHEPV    IPPSDKVIR- -FRTEVSKFL ---------- ---------- ---------- ---------- ---------E
```

**Figure S3 cont. Clustal Omega alignment of host/poxviral DUSP sequences (trimmed) used for PhyML analysis**

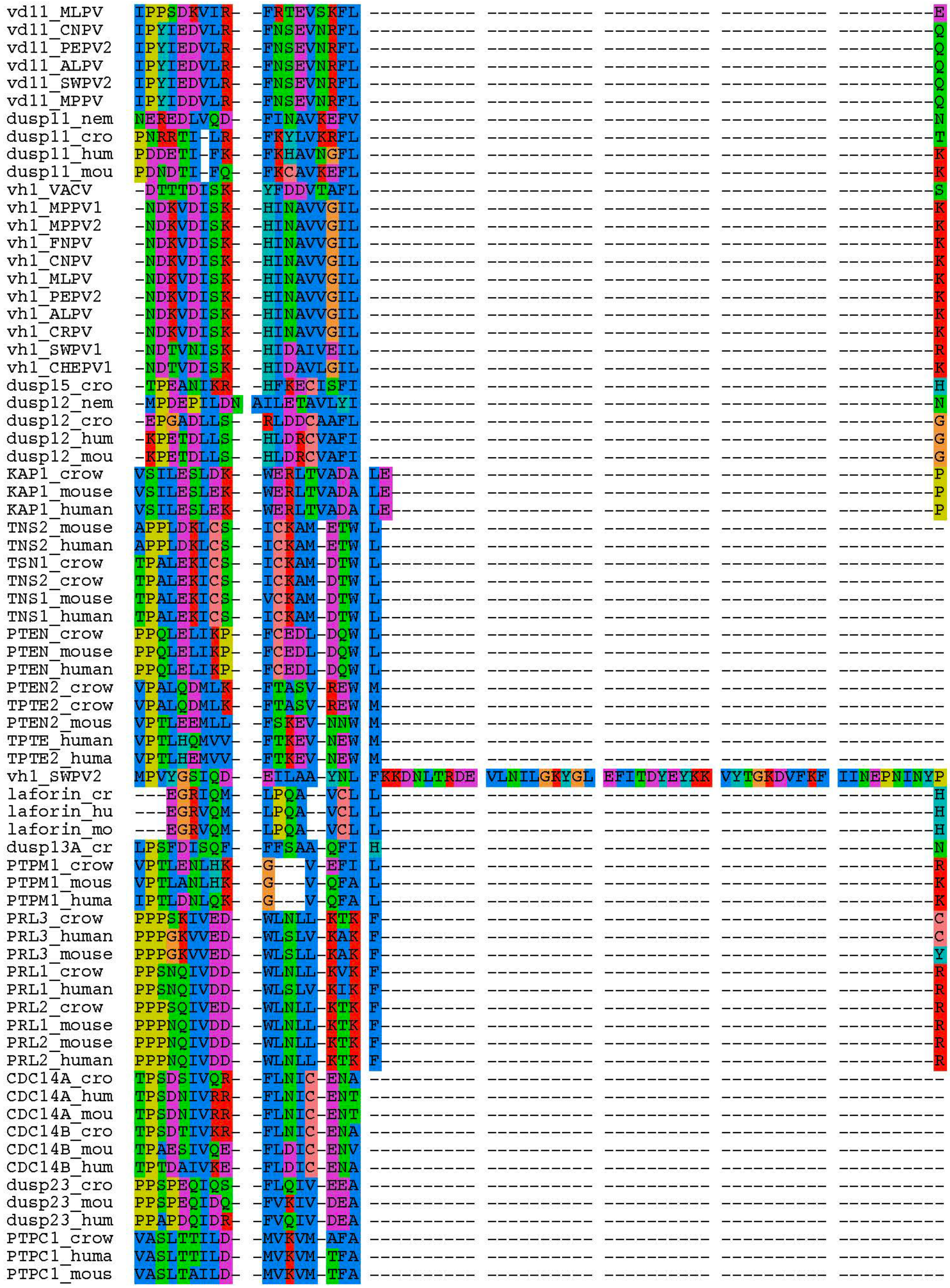


**Figure S3 cont. Clustal Omega alignment of host/poxviral DUSP sequences (trimmed) used for PhyML analysis**

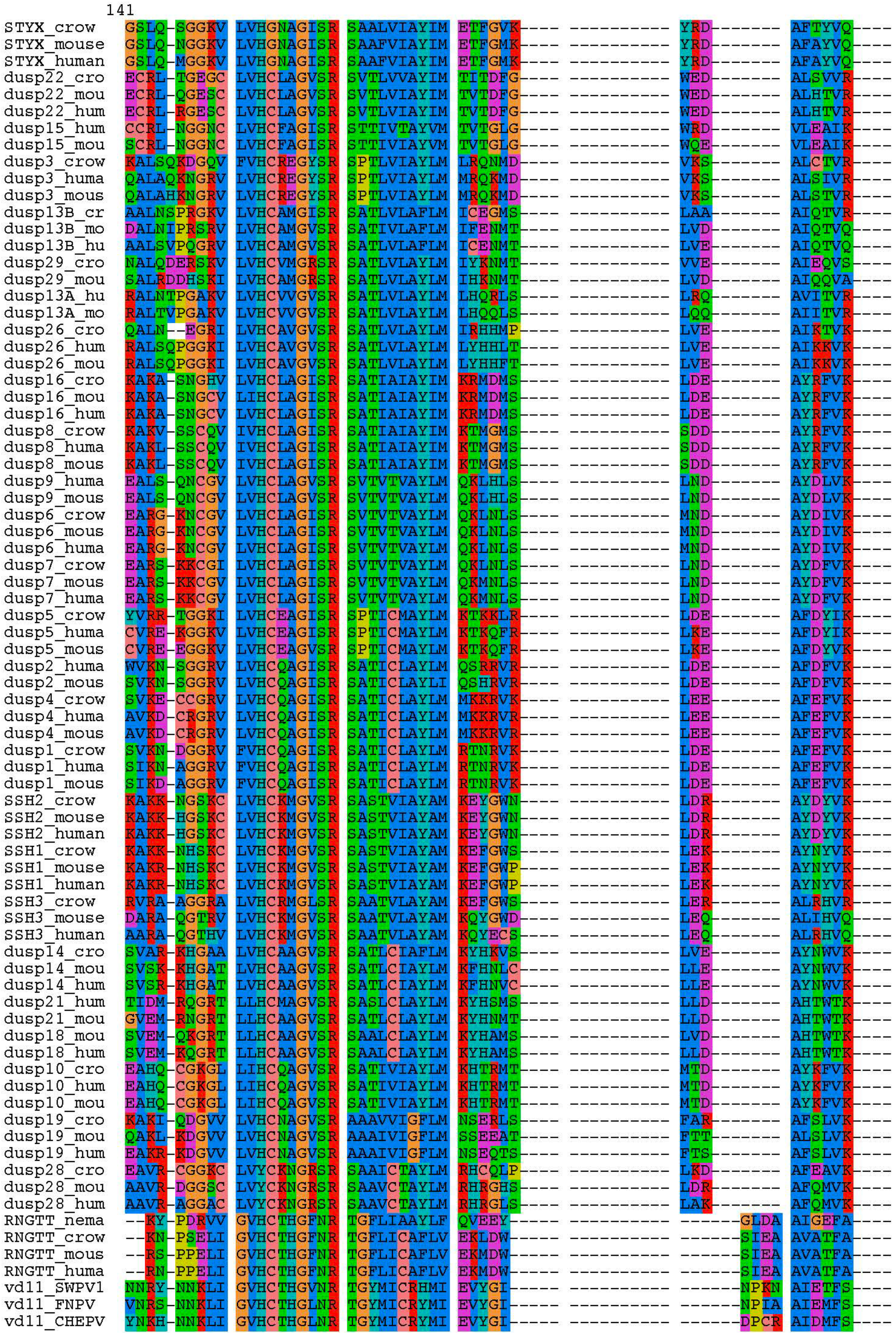


**Figure S3 cont. Clustal Omega alignment of host/poxviral DUSP sequences (trimmed) used for PhyML analysis**

```
vd11_MLPV   YNKH-NNKLI GVHCTHGLNR TGYMICRYMI EVYGI----- ---------- ------DPCR AIDMFS----
vd11_CNPV   FNRD-NNKLI GVHCTHGLNR TGYMICRYMI EVCGI----- ---------- ------DPAA AIEMFS----
vd11_PEPV2  FNRD-NNKLI GVHCTHGLNR TGYMICRYMI EVCGI----- ---------- ------DPAA AIEMFS----
vd11_ALPV   FNRD-NNKLI GVHCTHGLNR TGYMICRYMI EVCGI----- ---------- ------DPAA AIEMFS----
vd11_SWPV2  FNRD-NNKLI GVHCTHGLNR TGYMICRYMI EVCGI----- ---------- ------DPAA AIEMFS----
vd11_MPPV   FNRD-NNKLI GVHCTHGLNR TGYMICRYMI EVCGI----- ---------- ------DPAA AIEMFS----
dusp11_nem  DKEN-DGKLI GVHCTHGLNR TGYLICRYMI DVDNY----- ---------- ------SASD AISMFE----
dusp11_cro  DNKD-NDKLI GVHCTHGLNR TGYLVCRYLI DVEGM----- ---------- ------EPNA AIELFN----
dusp11_hum  ENKD-NDKLI GVHCTHGLNR TGYLICRYLI DVEGV----- ---------- ------RPDD AIELFN----
dusp11_mou  KNKN-NDKLI GVHCTHGLNR TGYLICRYLI DVEGM----- ---------- ------RPDD AIELFN----
vh1_VACV    KCDQ-RNEPV LVHCAAGVNR SGAMILAYLM SKNKES---- ---------- LPM--LYFLY VYHSMR----
vh1_MPPV1   KCDL-FKVPV LVHCMAGINR SAAMIMSYLM EIRDKN---- ---------- IPF-LIYFLY VYHELK----
vh1_MPPV2   KCDL-FKVPV LVHCMAGINR SAAMIMSYLM EIRDKN---- ---------- IPF-LIYFLY VYHELK----
vh1_FNPV    KCDS-FKVPV LVHCMAGINR SAAMIMSYLM EVRDKN---- ---------- IPF-LIYFLY VYHKLK----
vh1_CNPV    KCDL-FKVPV LVHCMAGINR SAAMIMSYLM EVRDKN---- ---------- IPF-LIYFLY VYHELK----
vh1_MLPV    KCDL-FKVPV LVHCMAGINR SAAMIMSYLM EVRDKN---- ---------- IPF-LIYFLY VYHELK----
vh1_PEPV2   KCDL-FKVPV LVHCMAGINR SAAMIMSYLM EVRDKN---- ---------- IPF-LIYFLY VYHELK----
vh1_ALPV    KCDL-FKVPV LVHCMAGINR SAAMIMSYLM EVRDKN---- ---------- IPF-LIYFLY VYHELK----
vh1_CRPV    KCDS-FKVPV LVHCMAGINR SAAMIMSYLM EVRDKN---- ---------- IPF-LIYFLY VYHELK----
vh1_SWPV1   KCDS-FKVPV LIHCMAGINR SSAMIMSYLM EIRDKN---- ---------- IPF-LIYFLY VYHELK----
vh1_CHEPV1  KCDS-FKVPV LVHCMAGINR SSAMIMSYLM DIRDKN---- ---------- IPF-LIYFLY VYHELK----
dusp15_cro  QCRL-HGGNC LVHCLAGISR STTVVVAYVM VVTEL----- ---------- ------SCQE VLDAIR----
dusp12_nem  EGVE-KEENV GVHCLAAVSR SVSICAAYLM YKNQW----- ---------- ------PVEK ALKMIE----
dusp12_cro  AARA-GGGAA LVRCHAGVSR SVAVVAAYLM KTQGL----- ---------- ------GWEE ALAAVR----
dusp12_hum  QARA-EGRAV LVHCHAGVSR SVAIITAFLM KTDQL----- ---------- ------PFEK AYEKLQ----
dusp12_mou  QARS-EGRAV LVHCHAGVSR SVAVVMAFIM KTDQL----- ---------- ------TFEK AYDILR----
KAP1_crow   VQFE-DGEKI VVQGEPGDD- ------FFII TEGTASVLQR RSDNEEYVEV GRLGPSDYFG EIALLLNRPR
KAP1_mouse  VQFE-DGEKI VVQGEPGDD- ------FYII TEGTASVLQR RSPNEEYVEV GRLGPSDYFG EIALLLNRPR
KAP1_human  VQFE-DGEKI VVQGEPGDD- ------FYII TEGTASVLQR RSPNEEYVEV GRLGPSDYFG EIALLLNRPR
TNS2_mouse  -SAD-PQHVV VLYCKGSKGK LGVIVSAYMH YSKISA---- ---------- GADQALATLT MRKFCE----
TNS2_human  -SAD-PQHVV VLYCKGNKGK LGVIVSAYMH YSKISA---- ---------- GADQALATLT MRKFCE----
TSN1_crow   -NAA-THNVV VLHNKGNRGR LGVVVAAYMH YSNISA---- ---------- SADQALDRFA MKRFYE----
TNS2_crow   -NAA-THNVV VLHNKGNRGR LGVVVAAYMH YSNISA---- ---------- SADQALDRFA MKRFYE----
TNS1_mouse  -NAD-PHNVV VLHNKGNRGR IGVVIAAYLH YSNISA---- ---------- SADQALDRFA MKRFYE----
TNS1_human  -NAD-PHNVV VLHNKGNRGR IGVVIAAYMH YSNISA---- ---------- SADQALDRFA MKRFYE----
PTEN_crow   -SED-DNHVA AIHCKAGKGR TGVMICAYLL HRGKFL---- ---------- RAQEALDFYG EVRTRD----
PTEN_mouse  -SED-DNHVA AIHCKAGKGR TGVMICAYLL HRGKFL---- ---------- KAQEALDFYG EVRTRD----
PTEN_human  -SED-DNHVA AIHCKAGKGR TGVMICAYLL HRGKFL---- ---------- KAQEALDFYG EVRTRD----
PTEN2_crow  -SQD-EKNVI AIHCKGGKGR TGTMVCIWLI DSDQFE---- ---------- SAKESLDYFG ERRTDR----
TPTE2_crow  -SQD-EKNVI AIHCKGGKGR TGTMVCIWLI DSDQFE---- ---------- SAKESLDYFG ERRTDR----
PTEN2_mous  -AQD-PENVV AIHCKGGKGR TGTMVCACLI ASEIVL---- ---------- NAKESLYFFG ERRTDK----
TPTE_human  -AQD-LENIV AIHCKGGTDR TGTMVCAFLI ASEICS---- ---------- TAKESLYYFG ERRTDK----
TPTE2_huma  -AQD-LENIV AIHCKGGKGR TGTMVCALLI ASEIFL---- ---------- TAEESLYYFG ERRTNK----
vh1_SWPV2   GILS-NGELI VDSI------ ----DSNFVV AMKHMS---- ---------- ISGLITDYKS SIEGIR----
laforin_cr  GLLE-NGHTV YVHCNAGVGR STAAVSGWLK YVMGWS---- ---------- LRK------- VQYFLV----
laforin_hu  ALLE-KGHIV YVHCNAGVGR STAAVCGWLQ YVMGWN---- ---------- LRK------- VQYFLM----
laforin_mo  ALLE-NGHTV YVHCNAGVGR STAAVCGWLH YVIGWN---- ---------- LRK------- VQYFIM----
dusp13A_cr  ALST-PGAKI LVHCAVGVSR SASLVLAYLM INHHLP---- ---------- LVE------- AIKTVK----
PTPM1_crow  -HRA-CGNSV YVHCKAGRSR SATMVAAYLI QLHQWS---- ---------- PRQ------- AIEAIA----
PTPM1_mous  -YQA-LGQCV YVHCKAGRSR SATMVAAYLI QVHNWS---- ---------- PEE------- AIEAIA----
PTPM1_huma  -YQS-LGQCV YVHCKAGRSR SATMVAAYLI QVHKWS---- ---------- PEE------- AVRAIA----
PRL3_crow   --ED-PGCCV AVHCVAGLGR APVLVALALI -ESGMK---- ---------- YED------- AIQFIR----
PRL3_human  --EA-PGSCV AVHCVAGLGR ---------- ---------- ---------- ---------- ----------
PRL3_mouse  --ND-PGSCV AVHCVAGLGR APVLVALALI -ESGMK---- ---------- YED------- AIQFIR----
PRL1_crow   --EE-PGCCI AVHCVAGLGR APVLVALALI -ECGMK---- ---------- YED------- AVQFIR----
PRL1_human  --EE-PGCCI AVHCVAGLGR APVLVALALI -EGGMK---- ---------- YED------- AVQFIR----
PRL2_crow   --EE-PGCCV AVHCVAGLGR APVLVALALI -ECGMK---- ---------- YED------- AVQFIR----
PRL1_mouse  --EE-PGCCV AVHCVAGLGR APVLVALALI -ECGMK---- ---------- YED------- AVQFIR----
PRL2_mouse  --EE-PGCCV AVHCVAGLGR APVLVALALI -ECGMK---- ---------- YED------- AVQFIR----
PRL2_human  --EE-PGCCV AVHCVAGLGR ---------- ---------- ---------- ---------- ----------
CDC14A_cro  ------DGAI AVHCKAGLGR TGTLIACYIM KHYKFT---- ---------- HAE------- AIAWIR----
CDC14A_hum  ------EGAI AVHCKAGLGR TGTLIACYVM KHYRFT---- ---------- HAE------- IIAWIR----
CDC14A_mou  ------EGAI AVHCKAGLGR TGTLIACYVM KHYRFT---- ---------- HAE------- IIAWIR----
CDC14B_cro  ------EGVI AVHCKAGLGR TGTLIACYIM KHYRMT---- ---------- AAE------- TIAWIR----
CDC14B_mou  ------KGAI AVHCKAGLGR TGTLIGCYLM KHYRMT---- ---------- AAE------- SIAWLR----
CDC14B_hum  ------EGAI AVHCKAGLGR TGTLIACYIM KHYRMT---- ---------- AAE------- TIAWVR----
dusp23_cro  --NG-RGEAV AVHCMLGYGR TGTLLACYLC KERHLA---- ---------- GGD------- AIREIR----
dusp23_mou  --NA-RGEAV GVHCALGFGR TGTMLACYLV KERALA---- ---------- AGD------- AIAEIR----
dusp23_hum  --NA-RGEAV GVHCALGFGR TGTMLACYLV KERGLA---- ---------- AGD------- AIAEIR----
PTPC1_crow  --LQ--EGRV AVHCHAGLGR TGVLIACYLV FATRMS---- ---------- ADQ------- AILFVR----
PTPC1_huma  --LQ--EGKV AIHCHAGLGR TGVLIACYLV FATRMT---- ---------- ADQ------- AIIFVR----
PTPC1_mous  --LQ--EGKV AVHCHAGLGR TGVLIACYLV FATRMT---- ---------- ADQ------- AIIFVR----
```

**Figure S3 cont. Clustal Omega alignment of host/poxviral DUSP sequences (trimmed) used for PhyML analysis**

```
              211
STYX_crow     ---------- ---------- ------ERRF CINPNAGF
STYX_mouse    ---------- ---------- ------ERRF CINPNAGF
STYX_human    ---------- ---------- ------ERRF CINPNAGF
dusp22_cro    ---------- ---------- ------AARS CANPNMGF
dusp22_mou    ---------- ---------- ------AGRS CANPNLGF
dusp22_hum    ---------- ---------- ------AGRS CANPNVGF
dusp15_hum    ---------- ---------- ------ATRP IANPNPGF
dusp15_mou    ---------- ---------- ------ASRP IANPNPGF
dusp3_crow    ---------- ---------- ------QKRE -IGPNDGF
dusp3_huma    ---------- ---------- ------QNRE -IGPNDGF
dusp3_mous    ---------- ---------- ------QNRE -IGPNDGF
dusp13B_cr    ---------- ---------- ------SHRG -ICPNSGF
dusp13B_mo    ---------- ---------- ------AHRD -ICPNSGF
dusp13B_hu    ---------- ---------- ------AHRN -ICPNSGF
dusp29_cro    ---------- ---------- ------RHRC -ILPNRGF
dusp29_mou    ---------- ---------- ------KNRC -VLPNRGF
dusp13A_hu    ---------- ---------- ------QHRW -VFPNRGF
dusp13A_mo    ---------- ---------- ------ERRW -IFPNRGF
dusp26_cro    ---------- ---------- ------DHRG -IIPNRGF
dusp26_hum    ---------- ---------- ------DHRG -IIPNRGF
dusp26_mou    ---------- ---------- ------DHRG -ITPNRGF
dusp16_cro    ---------- ---------- ------EKRP TISPNFNF
dusp16_mou    ---------- ---------- ------EKRP TISPNFNF
dusp16_hum    ---------- ---------- ------EKRP TISPNFNF
dusp8_crow    ---------- ---------- ------DRRP SISPNFNF
dusp8_huma    ---------- ---------- ------DRRP SISPNFNF
dusp8_mous    ---------- ---------- ------DRRP SISPNFNF
dusp9_huma    ---------- ---------- ------RKKS NISPNFNF
dusp9_mous    ---------- ---------- ------RKKS NISPNFNF
dusp6_crow    ---------- ---------- ------MKKS NISPNFNF
dusp6_mous    ---------- ---------- ------MKKS NISPNFNF
dusp6_huma    ---------- ---------- ------MKKS NISPNFNF
dusp7_crow    ---------- ---------- ------RKKS NISPNFNF
dusp7_mous    ---------- ---------- ------RKKS NISPNFNF
dusp7_huma    ---------- ---------- ------RKKS NISPNFNF
dusp5_crow    ---------- ---------- ------QRRS LISPNFGF
dusp5_huma    ---------- ---------- ------QRRS MVSPNFGF
dusp5_mous    ---------- ---------- ------QRRS VVSPNFGF
dusp2_huma    ---------- ---------- ------QRRG VISPNFSF
dusp2_mous    ---------- ---------- ------QRRG VISPNFSF
dusp4_crow    ---------- ---------- ------QRRS IISPNFSF
dusp4_huma    ---------- ---------- ------QRRS IISPNFSF
dusp4_mous    ---------- ---------- ------QRRS IISPNFSF
dusp1_crow    ---------- ---------- ------QRRS IISPNFSF
dusp1_huma    ---------- ---------- ------QRRS IISPNFSF
dusp1_mous    ---------- ---------- ------QRRS IISPNFSF
SSH2_crow     ---------- ---------- ------ERRT VTKPNPSF
SSH2_mouse    ---------- ---------- ------ERRT VTKPNPSF
SSH2_human    ---------- ---------- ------ERRT VTKPNPSF
SSH1_crow     ---------- ---------- ------QKRS IARPNAGF
SSH1_mouse    ---------- ---------- ------QKRS ITRPNAGF
SSH1_human    ---------- ---------- ------QKRS ITRPNAGF
SSH3_crow     ---------- ---------- ------HRRP GVLPNPGF
SSH3_mouse    ---------- ---------- ------ELRP IVRPNHGF
SSH3_human    ---------- ---------- ------ELRP IARPNPGF
dusp14_cro    ---------- ---------- ------SRRP VIRPNVGF
dusp14_mou    ---------- ---------- ------ARRP VIRPNLGF
dusp14_hum    ---------- ---------- ------ARRP VIRPNVGF
dusp21_hum    ---------- ---------- ------SRRP IIRPNNGF
dusp21_mou    ---------- ---------- ------TCRP IIRPNNGF
dusp18_mou    ---------- ---------- ------SCRP IIRPNSGF
dusp18_hum    ---------- ---------- ------SCRP IIRPNSGF
dusp10_cro    ---------- ---------- ------GKRP IISPNLNF
dusp10_hum    ---------- ---------- ------GKRP IISPNLNF
dusp10_mou    ---------- ---------- ------GKRP IISPNLNF
dusp19_cro    ---------- ---------- ------NARP AACPNPGF
dusp19_mou    ---------- ---------- ------EARP SICPNPGF
dusp19_hum    ---------- ---------- ------NARP SICPNSGF
dusp28_cro    ---------- ---------- ------TARP VAEPNAGF
dusp28_mou    ---------- ---------- ------SARP VAEPNLGF
dusp28_hum    ---------- ---------- ------SARP VAEPNPGF
RNGTT_nema    ---------- ---------- ------ENRQ KGIYKQDY
RNGTT_crow    ---------- ---------- ------QARP PGIYKGDY
RNGTT_mous    ---------- ---------- ------QARP PGIYKGDY
RNGTT_huma    ---------- ---------- ------QARP PGIYKGDY
vd11_SWPV1    ---------- ---------- ------KARK HKIERPLY
vd11_FNPV     ---------- ---------- ------DARK HKIERPSY
vd11_CHEPV    ---------- ---------- ------RARK HEIERPDY
```

**Figure S3 cont. Clustal Omega alignment of host/poxviral DUSP sequences (trimmed) used for PhyML analysis**

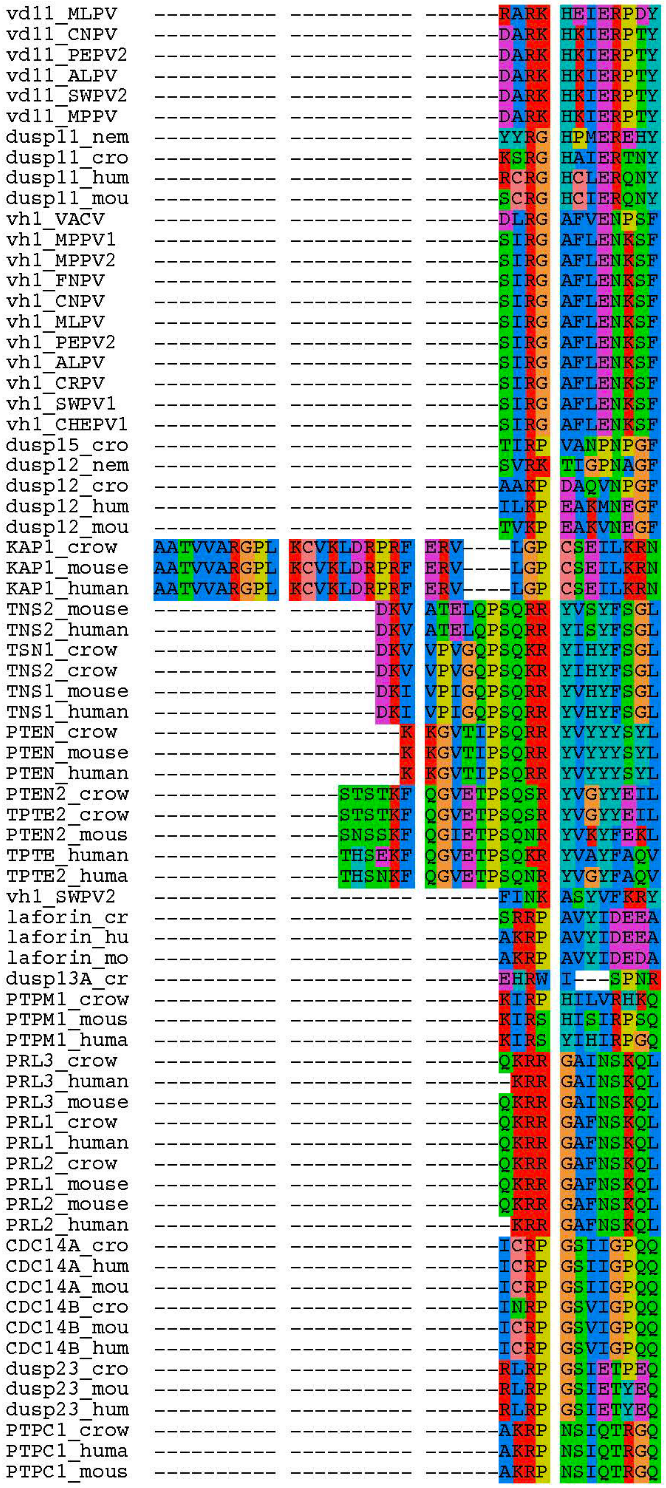


**Figure S3 cont. Clustal Omega alignment of host/poxviral DUSP sequences (trimmed) used for PhyML analysis**

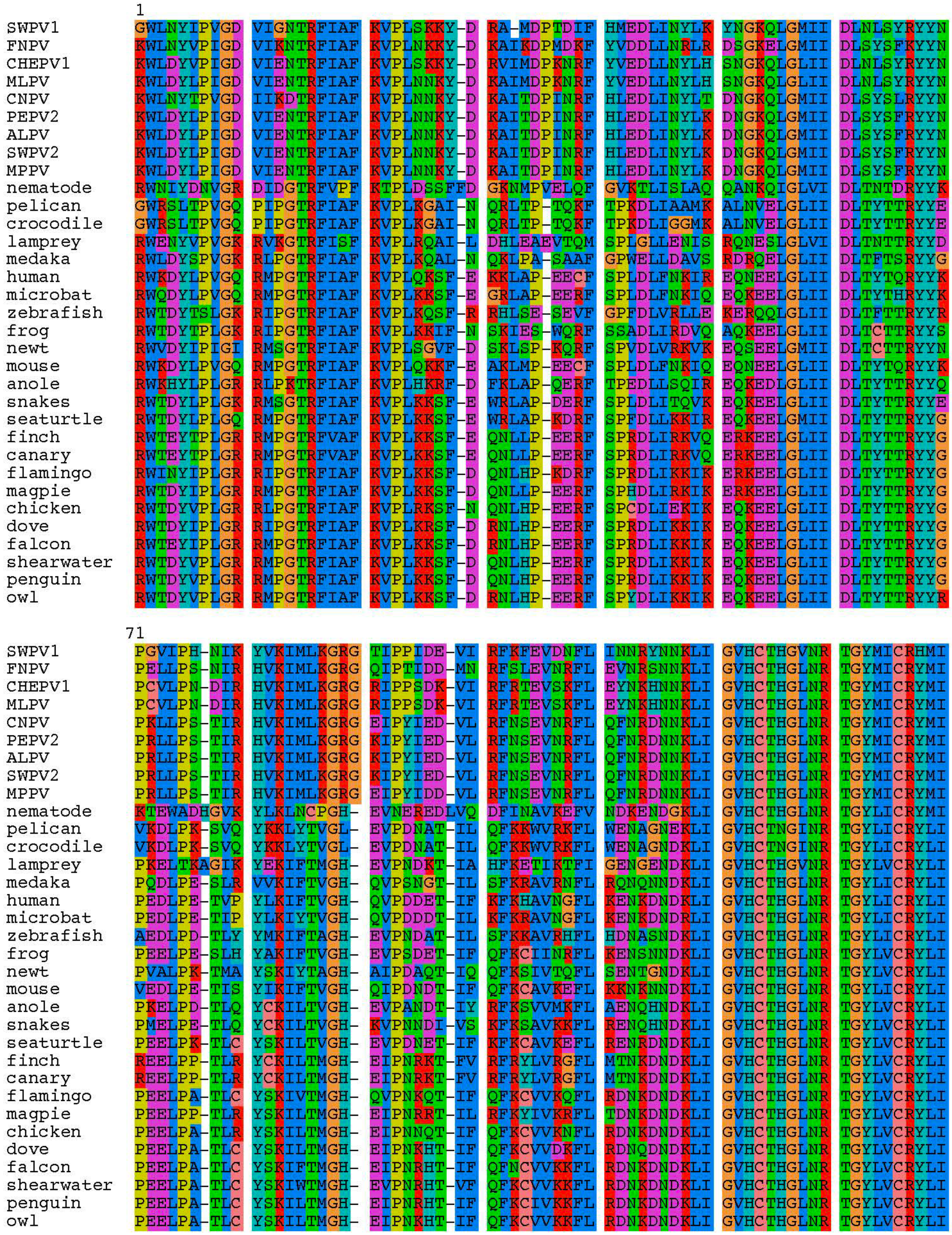


**Figure S4. Clustal Omega alignment of host DUSP11 and avipox vDUSP11 sequences (trimmed) used for PhyML analysis**

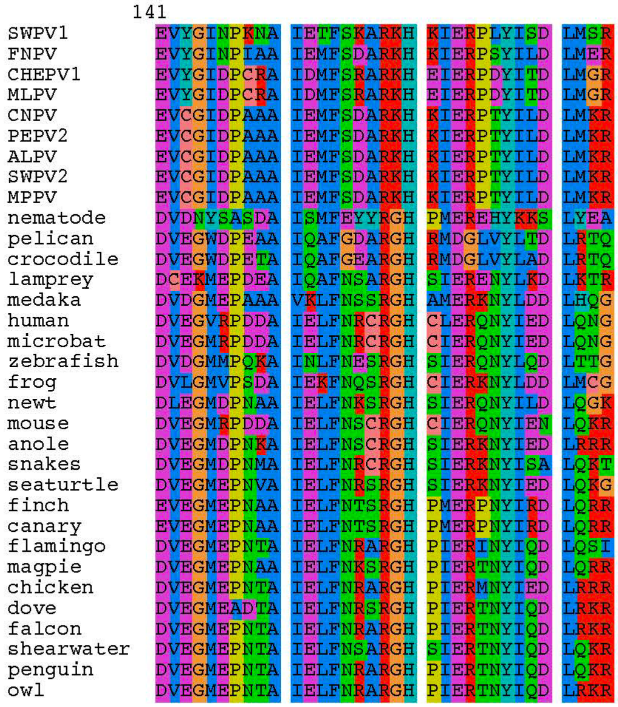


**Figure S4 cont. Clustal Omega alignment of host DUSP11 and avipox vDUSP11 sequences (trimmed) used for PhyML analysis**

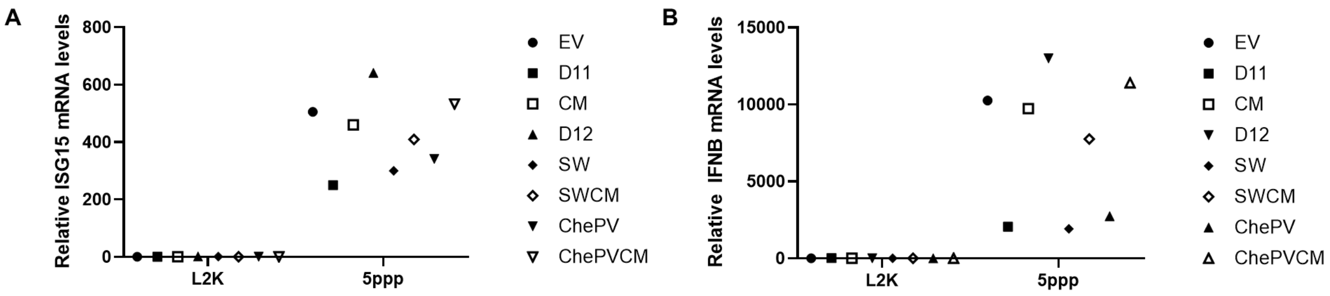


**Figure S5: vDUSP11 modulates immune activation in response to liposomal 5'-ppp-RNAs.**

Confirmation of key result trends from Figure 4 via a different wet bench scientist. A549 DUSP11 knockout (KO) reconstituted cells (12-well) were transfected with 5-10 ng of *in vitro* transcribed 5'-ppp-RNA for 18 hours followed by RT-qPCR to assay induction of ISGs. (A) RT-qPCR analysis of *ISG15* and (B) *IFNB1* mRNA normalized to *GAPDH* mRNA in 5'-ppp-RNA transfected in A549 DUSP11 KO reconstituted cells. Results are represented relative to those of empty vector-expressing cells. Data are derived n = 1 replicates.

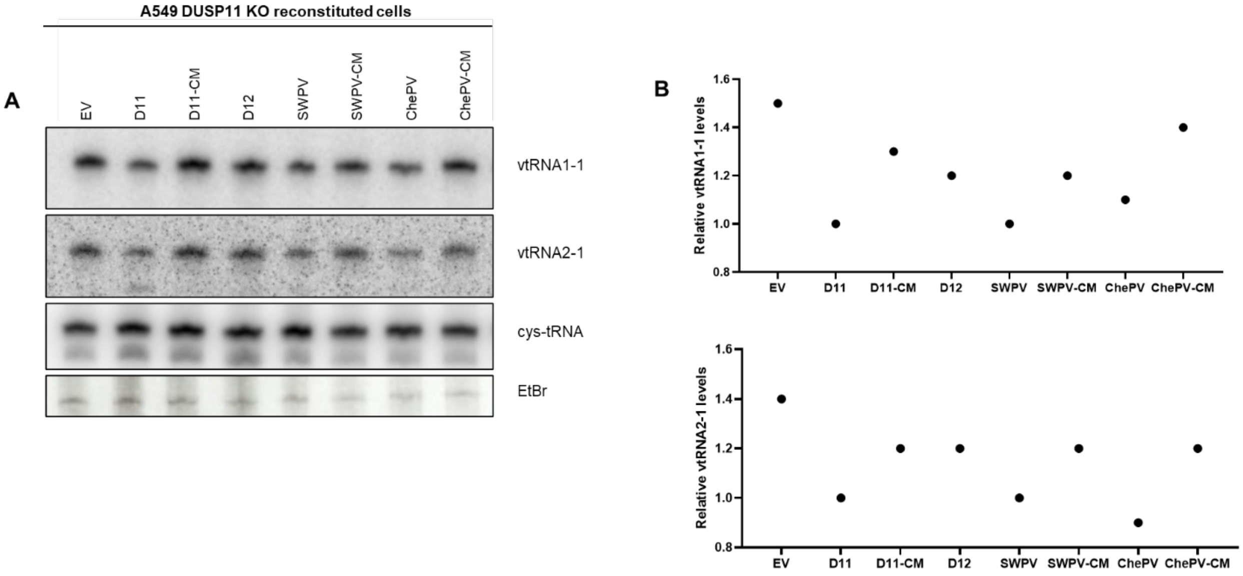


**Figure S6. vDUSP11 modulates steady-state RNA levels of endogenous RNAP III transcripts.**

Confirmation of key result trends from Figure 6 via a different wet bench scientist. (A) Northern blot analysis of vtRNA1-1 and vtRNA2-1 using RNA from A549 DUSP11 KO reconstituted cells. (B) Graphical representation of relative band intensity of vtRNA1-1 and vtRNA2-1 normalized to the relative band intensity of the cysteine-tRNA. Data are derived from n = 1 replicates.

Fig 6B. related to Fig 6B. rep 2 related to Fig 6B. rep3

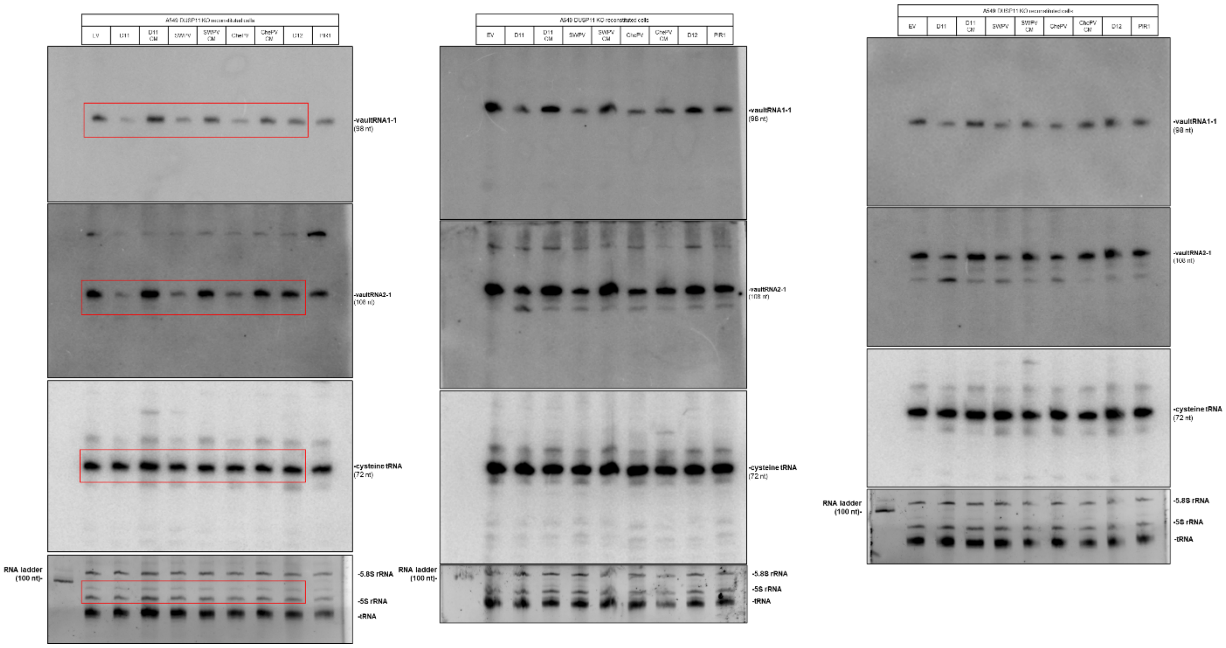


**Figure S7. uncropped northern blots**

Fig 3B.

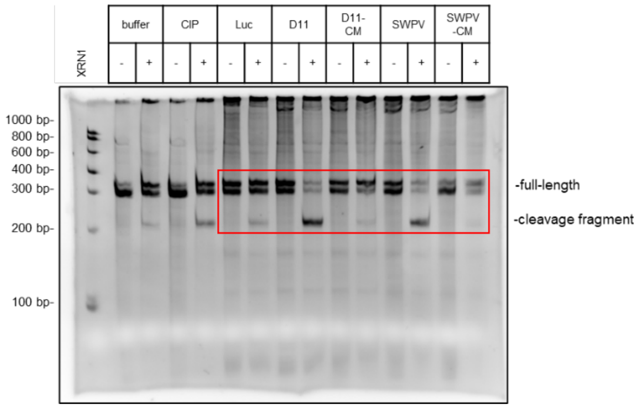


related to Fig3B. rep 3

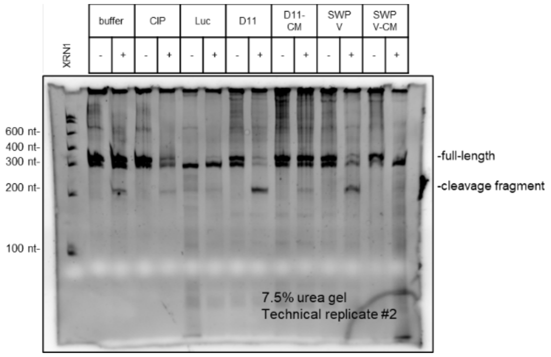


related to Fig3B. rep 2

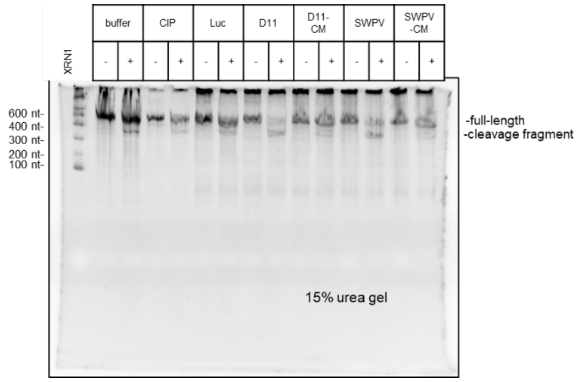


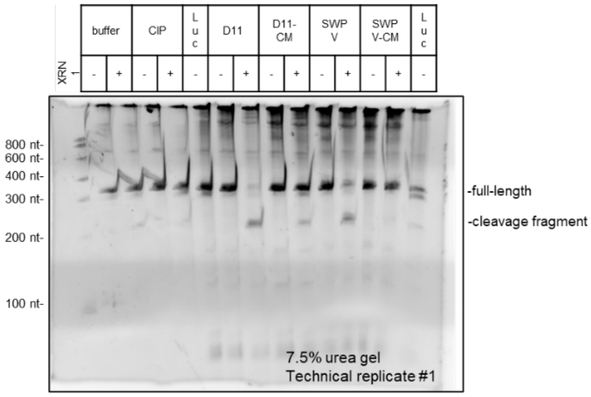


**Figure S8. uncropped ethidium bromide-stained urea gels**

Fig 3A.

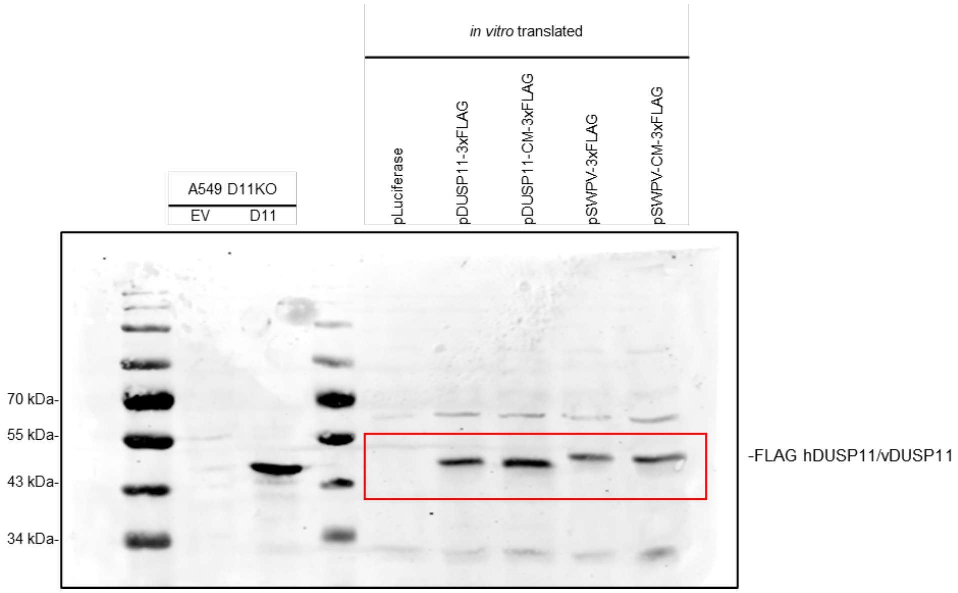


Fig 4B.

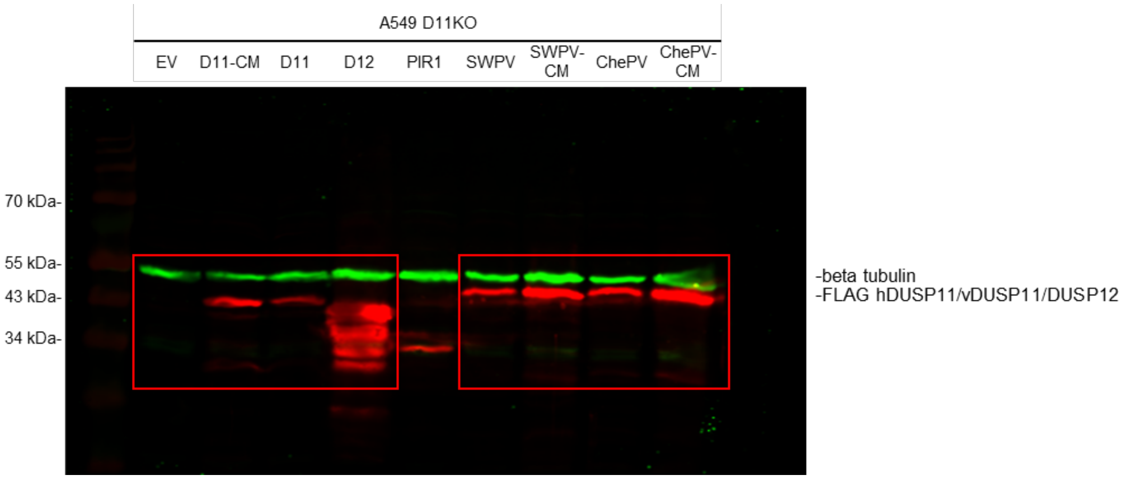


**Figure S9: uncropped western blots**

Replicate 1:

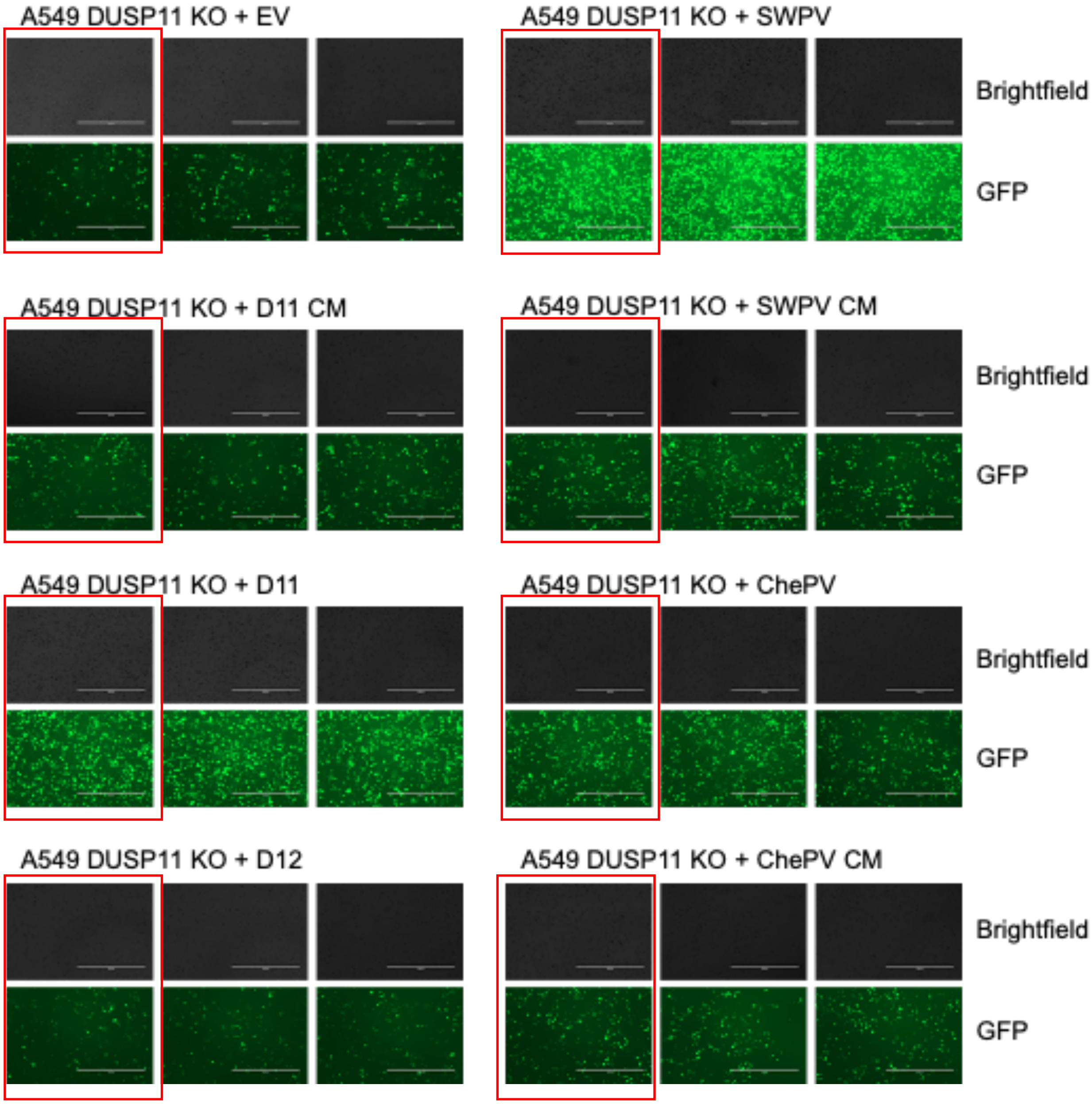


**Figure S10: M51R VSV infection GFP images**

Replicate 2:

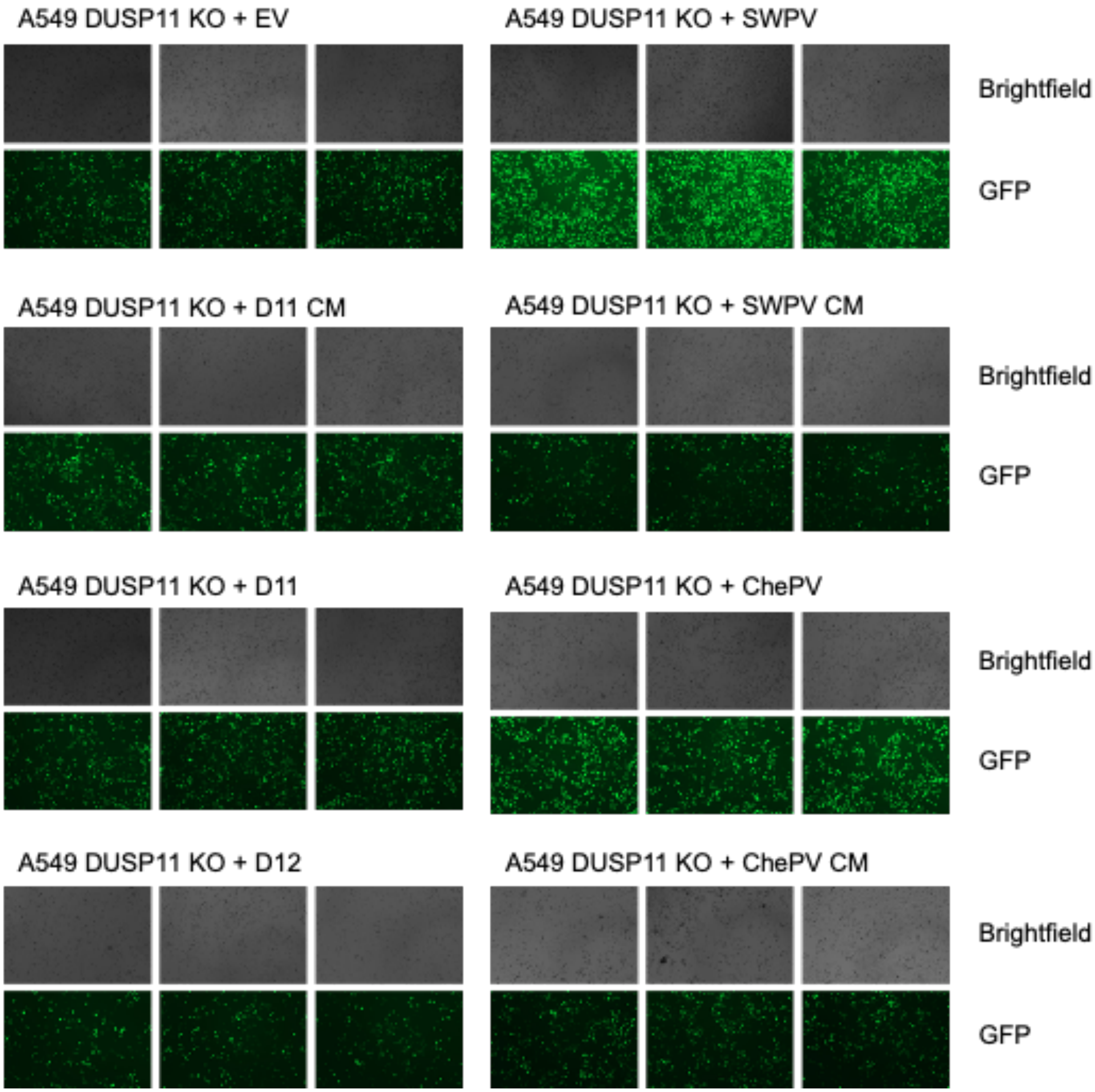


**Figure S10 cont.: M51R VSV infection GFP images**

Replicate 3:

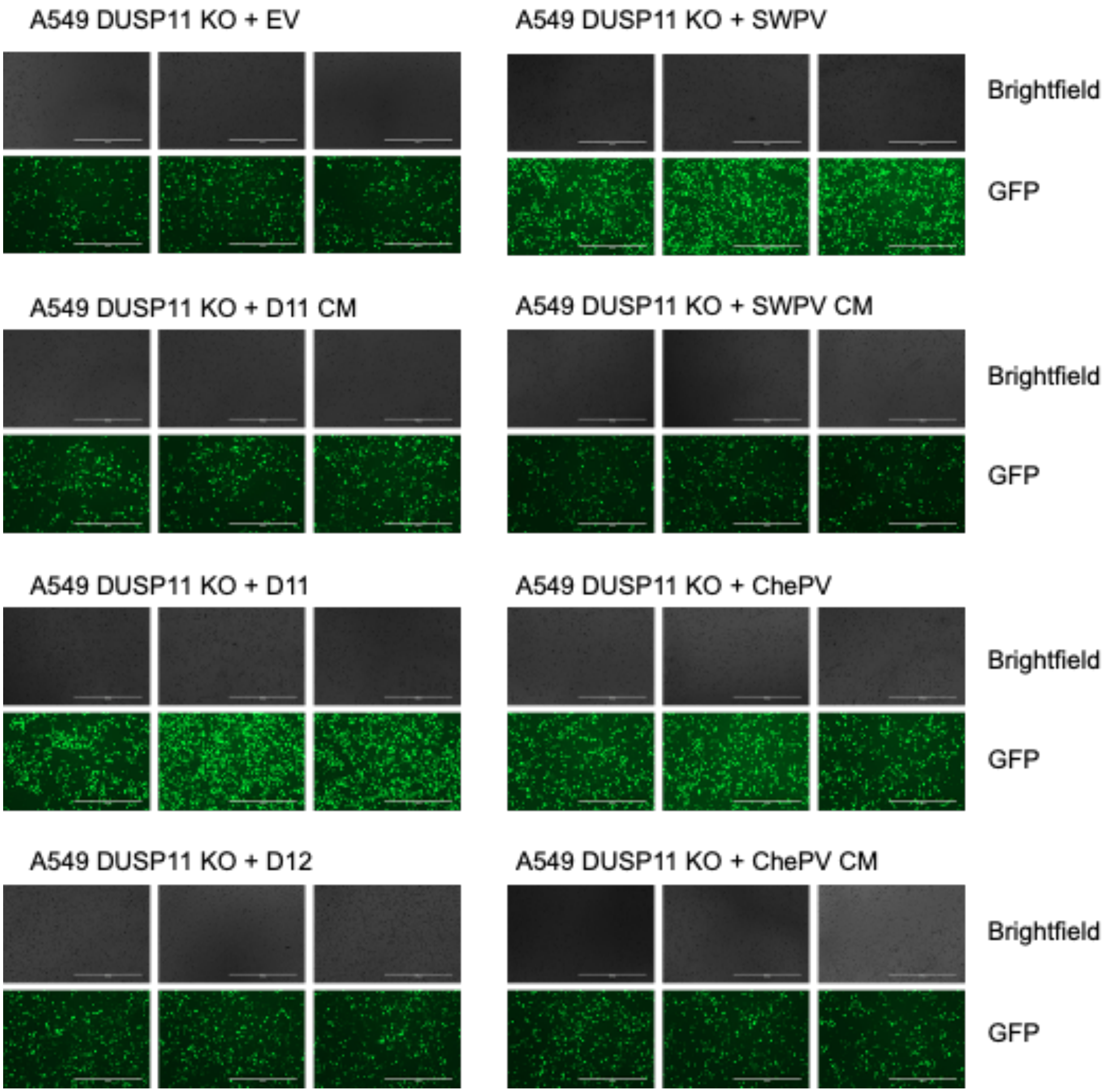


**Figure S10 cont.: M51R VSV infection GFP images**

#### Table S1: Sequence information, synthetic oligonucleotides and DNA, and reagents

| Oligo name | Sequence |
|---|---|
| 3xFLAG around the horn R | 5'-CTTGTCATCGTCATCCTTGTAG-'3 |
| DUSP12 around the horn F | 5'-TTGGAGGCTCCGGGC-'3 |
| SWPV around the horn F | 5'-GGAAAGAAAAAAAGCAAACATTATAATCA-'3 |
| SWPV CM R | 5'-AGCTGGGTCTAGATTATGCC-'3 |
| SWPV CM/ChePV CM F | 5'-ACTGAAACTCGAGGCCAC-'3 |
| SWPV CM around the horn F | 5'-AGTACTCACGGGCTTAACCGCAC-'3 |
| SWPV CM around the horn R | 5'-GTGTACACCAATCAACTTGTTATTGTCGC-'3 |
| ChePV CM R | 5'-AAAGCTGGGTCTAGATTAAGC-'3 |
| ChePV CM overlap F | 5'-CATAACAACAAATTGATAGGTGTCCATAGCACTCATGGTTTGAATCGAAC-'3 |
| ChePV CM overlap R | 5'-GTTCGATTCAAACCATGAGTGCTATGGACACCTATCAATTTGTTGTTATG-'3 |
| ChePV1 vDUSP11 gblock | 5'-<br>TGCACTCGAGGCCACCATGGACTACAAAGACCATGACGGTGATTATAAAGATCATGACAT<br>CGACTACAAGGATGACGATGACAAGGGAAAGAAAGCCCCGAGACACTACAACAATTTGC<br>CAGATAAGTGGTTGGATTATGTCCCGATCGGCGATGTAATTGAAAATACACGCTTCATTG<br>CGTTCAAAGTACCATTGTCTAAAAAGTATGACCGAGTTATCATGGATCCCAAAAACAGGTT<br>TTACGTCGAGGACCTCTTGAACTACCTCCACTCAAACGGAAAACAACTGGGACTGATCAT<br>TGATCTCAACCTGAGCTACCGCTATTATAATCCTTGCGTATTGCCTAATGATATTAGGCAT<br>GTGAAGATAATGCTCAAAGGACGGGGCCGCATACCGCCGAGCGATAAGGTCATAAGATT<br>CAGAACTGAAGTTTCAAAGTTCTTGGAGTATAATAAACATAACAACAAATTGATAGGTGTC<br>CATTGCACTCATGGTTTGAATCGAACGGGATACATGATTTGCCGCTATATGATCGAAGTG<br>TATGGTATCGACCCTTGTAGAGCAATAGACATGTTTAGCCGCGCCCGAAAACATGAAATA<br>GAACGGCCAGATTACATAACAGATTTGATGGGTCGGAAACATCTTAACATACGGCCACCG<br>ATAATGATGAATACGTATTCAGATAATTCAACTGTATTGAAAGTCAACGCCGTTGATAATT<br>TGCCTTATGTGAGGGTAAAAACAGTAAGCACCTGCGTCAAAATCGAAACAGTTAAAGAAG<br>ATACTCCAACGTGCGTGAAAATCGAGACGGTAAAGGAAGATACACCCACGTGTGTTAAG<br>ATTGAGACTGTAAAAGAGGATACGCCAACTTGTGTTAAAATCGAAACCGTAAAGGAGGAC<br>ACGCCTACCTGCGTGAAAATAGAGACAGTGAAAGAAGACACTCCCACATGTGTTAAGATA<br>GAGACCGTTAAAGAGGACATTCTCCTGGCTTAATCTAGACTAG-'3 |
| SWPV2 vDUSP11 gblock | 5'-<br>ACTGCTCGAGGCCACCATGGACTACAAAGACCATGACGGTGATTATAAAGATCATGACAT<br>CGACTACAAGGATGACGATGACAAGAGCCAGTGGCATCATATGGGAAAGAAAAAAAGCA<br>AACATTATAATCACATACCTGACAAATGGCTGGACTACCTCCCTATAGGCGACGTAATCG<br>AGAACACACGCTTTATTGCGTTCAAGGTGCCTTTGAACAATAAGTACGATAAGGCTATTA |

| | |
|---|---|
| | CAGATCCAATCAATAGATTCCATCTGGAAGACTTGATAAATTACCTGAAGGATAACGGGA<br>AACAACTTGGAATGATTATCGATCTTTCCTACAGTTTCAGGTACTACAATCCTAGGTTGCT<br>GCCTTCCACAATACGCCATGTTAAAATCATGTTGAAGGGGCGCGGAAAGATCCCTTACAT<br>CGAAGATGTATTGAGGTTTAATTCTGAAGTGAACCGCTTCTTGCAGTTTAACCGCGACAA<br>TAACAAGTTGATTGGTGTACACTGTACTCACGGGCTTAACCGCACTGGTTACATGATTTG<br>CAGATACATGATCGAAGTATGTGGCATCGATCCAGCTGCCGCAATTGAGATGTTCAGTGA<br>CGCCAGAAAACATAAAATCGAACGCCCGACTTACATTCTGGACTTGATGAAACGGAAACA<br>TCTGAACATTAGACCTCCCATAATGATGAACACGTACAGCGACAATAGTACCGTAATTAAA<br>GTCAATGCCGTTGACAACCTCCCCTATGTCCGCGTTAAAGCGGTGAGTACTTGTGTCAAG<br>ATCGAGACTGTTAAAGAGGATACCCCCACTTGCGTAAAGATAGAAACCGTGAAAGAAGAC<br>ACGCCAACTTGTGTTAAGATTGAGACGGTGAAGGAGGACACCCCTACCTGCGTCAAAAT<br>CGAGACCGTGAAGGAGGATACTCCAACATGTGTAAAAATCGAAACAGTAAAGGAAGACA<br>CTCCGACATGCGTGAAGATTGAAACGGTTAAGGAAGATATCCTCCTGGCATAATCTAGAG<br>ATC-'3 |
| human DUSP12 gBlock | 5'-<br>ACTGCTCGAGGCCACCATGGACTACAAAGACCATGACGGTGATTATAAAGATCATGACAT<br>CGACTACAAGGATGACGATGACAAGAGCCAGTGGCATCATATGTTGGAGGCTCCGGGCC<br>CGAGTGATGGCTGCGAGCTCAGCAACCCCAGCGCCAGCAGAGTCAGCTGTGCCGGGCA<br>GATGCTGGAAGTGCAGCCAGGATTGTATTTCGGTGGGGCCGCGGCCGTCGCGGAGCCA<br>GATCACCTGAGGGAAGCGGGCATCACGGCCGTGCTAACAGTGGACTCGGAGGAGCCCA<br>GCTTCAAGGCGGGGCCTGGGGTCGAGGATCTATGGCGCCTCTTCGTGCCAGCGCTGGA<br>CAAACCCGAGACGGACCTACTCAGCCATCTGGACCGGTGCGTGGCCTTCATCGGTCAG<br>GCCCGCGCTGAGGGCCGTGCGGTGTTGGTGCACTGTCATGCAGGAGTCAGTCGAAGTG<br>TGGCCATAATAACTGCTTTTCTCATGAAGACTGACCAACTTCCCTTTGAAAAAGCCTATGA<br>AAAGCTCCAGATTCTCAAACCAGAGGCTAAGATGAATGAGGGGTTTGAGTGGCAACTGA<br>AATTATACCAGGCAATGGGATACGAAGTGGATACCTCTAGTGCAATTTATAAGCAATATC<br>GTTTACAAAAGGTTACAGAGAAGTATCCAGAATTGCAGAATTTACCTCAAGAACTCTTTGC<br>TGTTGACCCAACTACCGTTTCACAAGGATTGAAAGATGAGGTTCTCTACAAGTGTAGAAA<br>GTGCAGGCGATCATTATTTCGAAGTTCTAGTATTCTGGATCACCGTGAAGGAAGTGGACC<br>TATAGCCTTTGCCCACAAGAGAATGACACCATCTTCCATGCTTACCACAGGGAGGCAAGC<br>TCAATGTACATCTTATTTCATTGAACCTGTACAGTGGATGGAATCTGCTTTGTTGGGAGTG<br>ATGGATGGACAGCTTCTTTGCCCAAAATGCAGTGCCAAGTTGGGTTCCTTCAACTGGTAT<br>GGTGAACAGTGCTCTTGTGGTAGGTGGATAACACCTGCTTTTCAAATACATAAGAATAGA<br>GTGGATGAAATGAAAATATTGCCTGTTTTGGGATCACAAACAGGAAAAATATGAATCTAGA<br>GATC-'3 |

Table S1 cont. Sequence information, synthetic oligonucleotides and DNA, and reagents

| **RT-qPCR primers** | |
|---|---|
| GAPDH F | 5'-ACATCGCTCAGACACCATG-'3 |
| GAPDH R | 5'-TGTAGTTGAGGTCAATGAAGGG-'3 |
| ISG15 F | 5'-ACTCATCTTTGCCAGTACAGG-'3 |
| ISG15 R | 5'-CAGCTCTGACACCGACATG-'3 |
| IFNB1 F | 5'-GTCACTGTGCCTGGACCATAG-'3 |
| IFNB1 R | 5'-GTTTCGGAGGTAACCTGTAAGTC-'3 |

| **Northern blot probes** | |
|---|---|
| vtRNA1-1 | 5'-TGTCGAAGTAACCGCTGAGC-'3 |
| vtRNA2-1 | 5'-GTAACCGCTTGAGCTAACTCCGAC-'3 |
| tRNA-Cys-GCA-6-1 | 5'-GTCAAATGCTCTACCACTGAGCTACACC-'3 |

| **Antibodies** | |
|---|---|
| DUSP11 polyclonal antibody (Rabbit) | Proteintech 10204-2-AP |
| Monoclonal ANTI-FLAG® M2 antibody produced in mouse | Sigma-Aldrich F1804 |
| Alpha Tubulin Monoclonal antibody (Mouse) | Proteintech 66031-1-Ig |
| β-Tubulin Polyclonal Antibody (Rabbit) | Cell Signaling Technology #2146s |
| 800CW Donkey anti-Rabbit IgG Secondary Antibody | LI-COR IRDye® 926-32213 |
| Goat anti-Mouse IgG1 Cross-Adsorbed Secondary Antibody, Alexa Fluor™ 488 | Invitrogen A10112 |
| Goat anti-Mouse IgG (H+L) Cross-Adsorbed Secondary Antibody, Alexa Fluor™ 594 | Invitrogen A11005 |
| 680LT Donkey anti-Mouse IgG Secondary Antibody | LI-COR IRDye® 925-68022 |

| **Reagents** | |
|---|---|
| dNTP Set, 100mM | Thermo Fisher R0181 |
| DNase I, RNase-free (1 U/µL) | Thermo Fisher EN0521 |
| SuperScript™ III First-Strand Synthesis System | Invitrogen 18080051 |
| PolyDT (5'- TTTTTTTTTTTTTTTTTTTT -'3) | IDT |
| PerfeCTa SYBR Green FastMix | Quantabio |
| ProLong Diamond Antifade Mountant | Invitrogen #P36961 |

#### Table S2. Blastp results using human DUSP11 (uniprot o75319) as query limiting results to viruses (taxid: 10239)

| Description | Scientific Name | Max Score | Total Score | Query Cover | E value | Per. Ident* | Acc. Len | Accession |
|---|---|---|---|---|---|---|---|---|
| putative RNA phosphatase [Penguinpox virus 2] | Penguinpox virus 2 | 186 | 186 | 53% | 2.00E-54 | 50.84 | 247 | QRM15716.1 |
| putative RNA phosphatase [Albatrosspox virus] | Albatrosspox virus | 186 | 186 | 53% | 2.00E-54 | 50.84 | 251 | QRM16049.1 |
| SWPV2-ORF080 [Shearwaterpox virus] | Shearwaterpox virus | 184 | 184 | 56% | 6.00E-53 | 50 | 303 | ARE67303.1 |
| putative RNA phosphatase [Magpiepox virus] | Magpiepox virus | 184 | 184 | 63% | 6.00E-53 | 45.79 | 286 | QGM48715.1 |
| DSP DUSP11 [Finch poxvirus] | Finch poxvirus | 181 | 181 | 58% | 2.00E-49 | 46.77 | 501 | UOX38764.1 |
| SWPV1-074 [Shearwaterpox virus] | Shearwaterpox virus | 172 | 172 | 52% | 4.00E-49 | 46.24 | 245 | ARF02681.1 |
| CNPV085 putative RNA phosphatase [Canarypox virus] | Canarypox virus | 175 | 175 | 63% | 3.00E-48 | 44.24 | 403 | NP_955108.1 |
| putative RNA phosphatase [Mudlarkpox virus] | Mudlarkpox virus | 175 | 175 | 53% | 3.00E-48 | 49.16 | 403 | QRM15361.1 |
| putative RNA phosphatase [Cheloniid poxvirus 1] | Cheloniid poxvirus 1 | 171 | 171 | 60% | 5.00E-48 | 43.78 | 303 | QRI42800.1 |
| protein tyrosine phosphatase 1 [Choristoneura fumiferana multiple nucleopolyhedrovirus] | Choristoneura fumiferana multiple nucleopolyhedrovirus | 140 | 140 | 50% | 3.00E-37 | 44.71 | 177 | NP_848321.1 |
| protein tyrosine phosphatase 1 [Choristoneura occidentalis alphabaculovirus] | Choristoneura occidentalis alphabaculovirus | 138 | 138 | 50% | 9.00E-37 | 44.71 | 177 | AGR57030.1 |
| protein tyrosine phosphatase 1 [Choristoneura rosaceana nucleopolyhedrovirus] | Choristoneura rosaceana nucleopolyhedrovirus | 135 | 135 | 50% | 1.00E-35 | 44.71 | 177 | YP_008378496.1 |
| protein tyrosine phosphatase 1 [Choristoneura diversana nucleopolyhedrovirus] | Choristoneura diversana nucleopolyhedrovirus | 135 | 135 | 50% | 1.00E-35 | 43.53 | 174 | BBU37623.1 |
| ptp1 [Neophasia sp. alphabaculovirus] | Neophasia sp. alphabaculovirus | 135 | 135 | 53% | 2.00E-35 | 44.63 | 173 | QBC76117.1 |
| protein tyrosine phosphatase 1 [Condylorrhiza vestigialis mutiple nucleopolyhedrovirus] | Condylorrhiza vestigialis mutiple nucleopolyhedrovirus | 135 | 135 | 50% | 2.00E-35 | 45.88 | 185 | YP_009118615.1 |
| ptp-1 [Choristoneura murinana nucleopolyhedrovirus] | Choristoneura murinana nucleopolyhedrovirus | 135 | 135 | 50% | 2.00E-35 | 43.53 | 174 | YP_008992233.1 |
| ptp1 [Dasychira pudibunda nucleopolyhedrovirus] | Dasychira pudibunda nucleopolyhedrovirus | 134 | 134 | 50% | 7.00E-35 | 41.76 | 177 | WHM28453.1 |
| protein tyrosine phosphatase 1 [Anticarsia gemmatalis nucleopolyhedrovirus] | Anticarsia gemmatalis nucleopolyhedrovirus | 133 | 133 | 50% | 9.00E-35 | 45.88 | 174 | YP_803403.1 |
| Protein tyrosine phosphatase 1 [Hyphantria cunea nucleopolyhedrovirus] | Hyphantria cunea nucleopolyhedrovirus | 133 | 133 | 54% | 1.00E-34 | 43.96 | 182 | UIX56267.1 |
| protein tyrosine phosphatase 1 [Choristoneura fumiferana DEF multiple nucleopolyhedrovirus] | Choristoneura fumiferana DEF multiple nucleopolyhedrovirus | 132 | 132 | 53% | 3.00E-34 | 44.07 | 173 | NP_932617.1 |
| protein tyrosine phosphatase 1 [Anticarsia gemmatalis multiple nucleopolyhedrovirus] | Anticarsia gemmatalis multiple nucleopolyhedrovirus | 130 | 130 | 50% | 8.00E-34 | 45.29 | 174 | YP_009316171.1 |
| protein tyrosine phosphatase 1 [Anticarsia gemmatalis multiple nucleopolyhedrovirus] | Anticarsia gemmatalis multiple nucleopolyhedrovirus | 130 | 130 | 50% | 9.00E-34 | 45.29 | 174 | ALR70119.1 |
| protein tyrosine phosphatase 1 [Dasychira pudibunda nucleopolyhedrovirus] | Dasychira pudibunda nucleopolyhedrovirus | 130 | 130 | 50% | 2.00E-33 | 41.18 | 177 | AKR14224.1 |
| ptp protein [Thysanoplusia orichalcea nucleopolyhedrovirus] | Thysanoplusia orichalcea nucleopolyhedrovirus | 129 | 129 | 50% | 3.00E-33 | 43.27 | 167 | YP_007250413.1 |
| PTP-1 [Dione juno nucleopolyhedrovirus] | Dione juno nucleopolyhedrovirus | 128 | 128 | 53% | 1.00E-32 | 42.13 | 184 | YP_010799899.1 |
| protein tyrosine phosphatase 1 [Spilosoma obliqua nucleopolyhedrosis virus] | Spilosoma obliqua nucleopolyhedrosis virus | 127 | 127 | 50% | 2.00E-32 | 43.53 | 180 | AUR45159.1 |
| Protein tyrosine phosphatase 1 [Hyphantria cunea nucleopolyhedrovirus] | Hyphantria cunea nucleopolyhedrovirus | 127 | 127 | 50% | 3.00E-32 | 43.53 | 180 | YP_473330.1 |
| PTP-1 [Epiphyas postvittana nucleopolyhedrovirus] | Epiphyas postvittana nucleopolyhedrovirus | 126 | 126 | 50% | 4.00E-32 | 43.2 | 169 | NP_203176.1 |
| protein tyrosine phosphatase 1 [Orgyia pseudotsugata multiple nucleopolyhedrovirus] | Orgyia pseudotsugata multiple nucleopolyhedrovirus | 127 | 127 | 50% | 4.00E-32 | 40 | 220 | NP_046166.1 |
| PTP [Bombyx mori nucleopolyhedrovirus] | Bombyx mori nucleopolyhedrovirus | 124 | 124 | 50% | 2.00E-31 | 40.94 | 168 | AGX01225.1 |
| PTP [Bombyx mori nucleopolyhedrovirus] | Bombyx mori nucleopolyhedrovirus | 124 | 124 | 50% | 2.00E-31 | 40.94 | 168 | NP_047551.1 |

#### Table S2 cont. Blastp results using human DUSP11 (uniprot o75319) as query limiting results to viruses (taxid: 10239)

| | | | | | | | | |
|---|---|---|---|---|---|---|---|---|
| Protein tyrosine phosphatase 1 [Lonomia obliqua multiple nucleopolyhedrovirus] | Lonomia obliqua multiple nucleopolyhedrovirus | 124 | 124 | 50% | 2.00E-31 | 40.94 | 171 | YP_009666494.1 |
| phosphotyrosine phosphatase [Bombyx mandarina nucleopolyhedrovirus] | Bombyx mandarina nucleopolyhedrovirus | 124 | 124 | 50% | 3.00E-31 | 40.94 | 168 | ACQ57327.1 |
| ptp [Bombyx mori nucleopolyhedrovirus] | Bombyx mori nucleopolyhedrovirus | 124 | 124 | 50% | 3.00E-31 | 40.94 | 168 | WRK23391.1 |
| ptp [Bombyx mori nucleopolyhedrovirus] | Bombyx mori nucleopolyhedrovirus | 123 | 123 | 50% | 4.00E-31 | 40.94 | 168 | QWC64843.1 |
| PTP [Bombyx mori nucleopolyhedrovirus] | Bombyx mori nucleopolyhedrovirus | 123 | 123 | 50% | 7.00E-31 | 40.94 | 168 | AFN21248.1 |
| PTP [Bombyx mori nucleopolyhedrovirus] | Bombyx mori nucleopolyhedrovirus | 122 | 122 | 50% | 8.00E-31 | 40.94 | 168 | AFN21110.1 |
| PTP [Alphabaculovirus bomori] | Alphabaculovirus bomori | 122 | 122 | 50% | 9.00E-31 | 40.94 | 168 | WZB50497.1 |
| tyrosine phosphatase NPV-PTP [Bombyx mori nucleopolyhedrovirus] | Bombyx mori nucleopolyhedrovirus | 122 | 122 | 50% | 1.00E-30 | 40.94 | 168 | AAG31657.1 |
| Chain A, polynucleotide 5'-phosphatase [Autographa californica nucleopolyhedrovirus] | Autographa californica nucleopolyhedrovirus | 122 | 122 | 50% | 1.00E-30 | 40.35 | 169 | 1YN9_A |
| protein tyrosine phosphatase [Autographa californica nucleopolyhedrovirus] | Autographa californica nucleopolyhedrovirus | 122 | 122 | 50% | 1.00E-30 | 40.35 | 168 | NP_054030.1 |
| BVP=protein tyrosine phosphatase [Autographa californica multicapsid nuclear polyhedrosis virus AcMNPV, Peptide, 167 aa] [Autographa californica nucleopolyhedrovirus] | Autographa californica nucleopolyhedrovirus | 122 | 122 | 50% | 1.00E-30 | 40.35 | 167 | AAB25579.1 |
| protein tyrosine/serine phosphatase [Rachiplusia ou multiple nucleopolyhedrovirus] | Rachiplusia ou multiple nucleopolyhedrovirus | 122 | 122 | 50% | 2.00E-30 | 40.35 | 168 | NP_702993.1 |
| protein tyrosine phosphatase [Bombyx mori nucleopolyhedrovirus] | Bombyx mori nucleopolyhedrovirus | 121 | 121 | 50% | 2.00E-30 | 40.94 | 168 | BEV21003.1 |
| ptp [Bombyx mori nucleopolyhedrovirus] | Bombyx mori nucleopolyhedrovirus | 121 | 121 | 50% | 3.00E-30 | 40.35 | 168 | WRK23115.1 |
| ptp [Bombyx mori nucleopolyhedrovirus] | Bombyx mori nucleopolyhedrovirus | 120 | 120 | 50% | 5.00E-30 | 40.35 | 168 | WRK23253.1 |
| protein tyrosine phosphatase; PTP [Maruca vitrata nucleopolyhedrovirus] | Maruca vitrata nucleopolyhedrovirus | 118 | 118 | 51% | 4.00E-29 | 41.28 | 179 | YP_950853.1 |
| ptp-1 [Antheraea pernyi nucleopolyhedrovirus] | Antheraea pernyi nucleopolyhedrovirus | 116 | 116 | 52% | 3.00E-28 | 41.01 | 178 | YP_611104.1 |
| ptp [Troides aeacus nucleopolyhedrovirus] | Troides aeacus nucleopolyhedrovirus | 116 | 116 | 50% | 3.00E-28 | 39.18 | 168 | QVU21318.1 |
| ptp 1 [Philosamia cynthia ricini nucleopolyhedrovirus virus] | Philosamia cynthia ricini nucleopolyhedrovirus virus | 115 | 115 | 50% | 9.00E-28 | 41.18 | 178 | AFY62938.1 |
| ptp [Palpita vitrealis nucleopolyhedrovirus] | Palpita vitrealis nucleopolyhedrovirus | 114 | 114 | 50% | 2.00E-27 | 39.05 | 170 | USC25988.1 |
| ORF-127 [Catopsilia pomona nucleopolyhedrovirus] | Catopsilia pomona nucleopolyhedrovirus | 114 | 114 | 50% | 2.00E-27 | 37.65 | 172 | YP_009255384.1 |
| protein tyrosine phosphatase 1 [Samia ricini nucleopolyhedrovirus] | Samia ricini nucleopolyhedrovirus | 113 | 113 | 50% | 4.00E-27 | 40.59 | 178 | BBD51358.1 |
| protein tyrosine phosphatase 1 [Mythimna separata entomopoxvirus 'L'] | Mythimna separata entomopoxvirus 'L' | 112 | 112 | 51% | 6.00E-27 | 36.42 | 173 | YP_008003686.1 |
| PTP [Parapoynx stagnalis nucleopolyhedrovirus] | Parapoynx stagnalis nucleopolyhedrovirus | 105 | 105 | 50% | 4.00E-24 | 33.93 | 173 | UZE89807.1 |
| phosphotyrosine phosphatase [Iragoides fasciata nucleopolyhedrovirus] | Iragoides fasciata nucleopolyhedrovirus | 101 | 101 | 51% | 9.00E-23 | 35.47 | 172 | ACJ04624.1 |
| ptp [Oxyplax ochracea nucleopolyhedrovirus] | Oxyplax ochracea nucleopolyhedrovirus | 99.8 | 99.8 | 51% | 6.00E-22 | 34.88 | 169 | YP_009666650.1 |
| protein tyrosine phosphatase 1 [Choristoneura rosaceana entomopoxvirus 'L'] | Choristoneura rosaceana entomopoxvirus 'L' | 93.2 | 93.2 | 52% | 1.00E-19 | 32.57 | 161 | YP_008004653.1 |
| protein tyrosine phosphatase 1 [Choristoneura biennis entomopoxvirus] | Choristoneura biennis entomopoxvirus | 93.2 | 93.2 | 52% | 1.00E-19 | 32.18 | 161 | YP_008004352.1 |
| protein tyrosine/serine phosphatase [Autographa californica multiple nucleopolyhedrovirus] | Autographa californica multiple nucleopolyhedrovirus | 89.4 | 89.4 | 26% | 5.00E-19 | 46.07 | 93 | AEZ36153.1 |

**Table S2 cont. Blastp results using human DUSP11 (uniprot o75319) as query limiting results to viruses (taxid: 10239)**

| | | | | | | | | |
|---|---|---|---|---|---|---|---|---|
| protein tyrosine phosphatase 1 [Betaentomopoxvirus amoorei] | Betaentomopoxvirus amoorei | 90.5 | 90.5 | 51% | 1.00E-18 | 31.76 | 157 | NP_065028.1 |
| protein tyrosine phosphatase 1 [Mythimna separata entomopoxvirus 'L'] | Mythimna separata entomopoxvirus 'L' | 89.4 | 89.4 | 51% | 3.00E-18 | 32.37 | 154 | YP_008003788.1 |
| PTP [Alphabaculovirus bomori] | Alphabaculovirus bomori | 82.8 | 82.8 | 16% | 5.00E-17 | 60.38 | 56 | WZB50807.1 |
| mRNA capping enzyme [Rock bream iridovirus] | Rock bream iridovirus | 69.3 | 69.3 | 46% | 9.00E-10 | 33.12 | 415 | AAT71875.1 |
| ORF_088L [Scale drop disease virus] | Scale drop disease virus | 68.2 | 68.2 | 47% | 1.00E-09 | 33.54 | 329 | YP_009163849.1 |
| mRNA capping enzyme [Infectious spleen and kidney necrosis virus] | Infectious spleen and kidney necrosis virus | 68.2 | 68.2 | 46% | 2.00E-09 | 33.12 | 491 | QPO16431.1 |
| mRNA-capping enzyme [Scale drop disease virus] | Scale drop disease virus | 68.2 | 68.2 | 47% | 2.00E-09 | 33.54 | 500 | QLI60762.1 |
| mRNA capping enzyme [Pompano iridovirus] | Pompano iridovirus | 68.2 | 68.2 | 46% | 2.00E-09 | 33.12 | 490 | AZQ21054.1 |
| mRNA capping enzyme [Pompano iridovirus] | Pompano iridovirus | 68.2 | 68.2 | 46% | 2.00E-09 | 33.12 | 490 | AZQ20812.1 |
| mRNA capping enzyme [Orange-spotted grouper iridovirus] | Orange-spotted grouper iridovirus | 68.2 | 68.2 | 46% | 2.00E-09 | 33.12 | 490 | AAX82373.1 |
| mRNA capping enzyme [Banggai cardinalfish iridovirus] | Banggai cardinalfish iridovirus | 67.8 | 67.8 | 46% | 3.00E-09 | 33.12 | 491 | QJC63414.1 |
| mRNA capping enzyme [Infectious spleen and kidney necrosis virus] | Infectious spleen and kidney necrosis virus | 67.8 | 67.8 | 46% | 3.00E-09 | 33.12 | 491 | UWH19944.1 |
| putative RNA guanylytransferase [Infectious spleen and kidney necrosis virus] | Infectious spleen and kidney necrosis virus | 67.8 | 67.8 | 46% | 3.00E-09 | 33.12 | 491 | NP_612286.1 |
| mRNA capping enzyme [Infectious spleen and kidney necrosis virus] | Infectious spleen and kidney necrosis virus | 67.8 | 67.8 | 46% | 3.00E-09 | 33.12 | 491 | UWH18607.1 |
| hypothetical protein IJGMMPBP_00063 [Infectious spleen and kidney necrosis virus] | Infectious spleen and kidney necrosis virus | 67.8 | 67.8 | 46% | 3.00E-09 | 33.12 | 491 | QQZ00516.1 |
| putative RNA guanylytransferase [Spotted knifejaw iridovirus] | Spotted knifejaw iridovirus | 67.8 | 67.8 | 46% | 3.00E-09 | 33.12 | 491 | QYK20590.1 |
| mRNA capping enzyme [Banggai cardinalfish iridovirus] | Banggai cardinalfish iridovirus | 67.8 | 67.8 | 46% | 4.00E-09 | 33.12 | 491 | QOE77202.1 |
| mRNA capping enzyme [Three spot gourami iridovirus] | Three spot gourami iridovirus | 67.4 | 67.4 | 46% | 4.00E-09 | 33.12 | 490 | AVR29831.1 |
| hypothetical protein AOEHFIPN_00039 [Infectious spleen and kidney necrosis virus] | Infectious spleen and kidney necrosis virus | 67.4 | 67.4 | 46% | 4.00E-09 | 33.12 | 491 | WNY63482.1 |
| mRNA capping enzyme [South American cichlid iridovirus] | South American cichlid iridovirus | 67.4 | 67.4 | 46% | 4.00E-09 | 33.12 | 490 | AVR29715.1 |
| mRNA capping enzyme [Turbot reddish body iridovirus] | Turbot reddish body iridovirus | 66.6 | 66.6 | 33% | 8.00E-09 | 39.64 | 490 | ADE34404.1 |
| putative RNA guanylytransferase MCE [Red seabream iridovirus] | Red seabream iridovirus | 67 | 67 | 33% | 9.00E-09 | 39.64 | 1384 | UNA01179.1 |
| protein tyrosine phosphatase 1 [Spilarctia obliqua nucleopolyhedrovirus] | Spilarctia obliqua nucleopolyhedrovirus | 59.3 | 59.3 | 32% | 7.00E-08 | 35.45 | 112 | QNN89305.1 |
| **MPPV-091 putative RNA phosphatase [Magpiepox virus 2]** | **Magpiepox virus 2** | **54.7** | **54.7** | **18%** | **1.00E-06** | **50** | **60** | **QZW33383.1** |
| **CRPV-114 [Crowpox virus]** | **Crowpox virus** | **52** | **52** | **11%** | **6.00E-06** | **60.53** | **42** | **UWX11223.1** |

#### Table S3. Blastp results using vDUSP11 from SWPV2 as query (ARE67303.1)

| Description | Scientific Name | Max Score | Total Score | Query Cover | E value | Per. ident | Acc. Len | Accession |
|---|---|---|---|---|---|---|---|---|
| SWPV2-ORF080 [Shearwaterpox virus] | Shearwaterpox virus | 618 | 618 | 100% | 0 | 100 | 303 | ARE67303.1 |
| CNPV085 putative RNA phosphatase [Canarypox virus] | Canarypox virus | 581 | 872 | 99% | 0 | 93.33 | 403 | NP_955108.1 |
| putative RNA phosphatase [Magpiepox virus] | Magpiepox virus | 569 | 569 | 92% | 0 | 98.57 | 286 | QGM48715.1 |
| putative RNA phosphatase [Mudlarkpox virus] | Mudlarkpox virus | 530 | 820 | 99% | 0 | 85.67 | 403 | QRM15361.1 |
| putative RNA phosphatase [Cheloniid poxvirus 1] | Cheloniid poxvirus 1 | 521 | 521 | 100% | 0 | 83.5 | 303 | QRI42800.1 |
| DSP DUSP11 [Finch poxvirus] | Finch poxvirus | 496 | 1204 | 98% | 7.00E-172 | 77.67 | 501 | UOX38764.1 |
| putative RNA phosphatase [Penguinpox virus 2] | Penguinpox virus 2 | 482 | 482 | 77% | 5.00E-170 | 98.72 | 247 | QRM15716.1 |
| putative RNA phosphatase [Albatrosspox virus] | Albatrosspox virus | 480 | 480 | 81% | 3.00E-169 | 94.74 | 251 | QRM16049.1 |
| SWPV1-074 [Shearwaterpox virus] | Shearwaterpox virus | 271 | 271 | 61% | 8.00E-87 | 69.52 | 245 | ARF02681.1 |
| RNA/RNP complex-1-interacting phosphatase [Corvus hawaiiensis] | Corvus hawaiiensis | 200 | 200 | 58% | 3.00E-58 | 53.41 | 305 | XP_048146001.1 |
| RNA/RNP complex-1-interacting phosphatase [Corvus cornix cornix] | Corvus cornix cornix | 199 | 199 | 58% | 2.00E-57 | 53.41 | 349 | XP_039420096.1 |
| RNA/RNP complex-1-interacting phosphatase [Calypte anna] | Calypte anna | 195 | 195 | 57% | 2.00E-56 | 52.87 | 320 | XP_030319948.1 |
| DUS11 phosphatase [Struthidea cinerea] | Struthidea cinerea | 190 | 190 | 56% | 4.00E-56 | 54.39 | 169 | NXB60100.1 |
| DUS11 phosphatase [Upupa epops] | Upupa epops | 189 | 189 | 56% | 5.00E-56 | 53.22 | 169 | NWU89504.1 |
| RNA/RNP complex-1-interacting phosphatase isoform X3 [Accipiter gentilis] | Accipiter gentilis | 193 | 193 | 57% | 5.00E-56 | 52.87 | 271 | XP_049686642.1 |
| RNA/RNP complex-1-interacting phosphatase isoform X2 [Aquila chrysaetos chrysaetos] | Aquila chrysaetos chrysaetos | 194 | 194 | 57% | 6.00E-56 | 52.87 | 314 | XP_029890719.1 |
| uncharacterized protein LOC116435472 [Corvus moneduloides] | Corvus moneduloides | 200 | 200 | 59% | 6.00E-56 | 52.51 | 535 | XP_031947699.1 |
| DUS11 phosphatase [Pachycephala philippinensis] | Pachycephala philippinensis | 189 | 189 | 56% | 6.00E-56 | 53.8 | 169 | NXI05060.1 |
| RNA/RNP complex-1-interacting phosphatase isoform X3 [Aquila chrysaetos chrysaetos] | Aquila chrysaetos chrysaetos | 192 | 192 | 57% | 7.00E-56 | 52.87 | 271 | XP_040984244.1 |
| RNA/RNP complex-1-interacting phosphatase isoform X3 [Aythya fuligula] | Aythya fuligula | 190 | 190 | 57% | 9.00E-56 | 52.87 | 197 | XP_032059945.1 |
| PREDICTED: RNA/RNP complex-1-interacting phosphatase isoform X3 [Haliaeetus leucocephalus] | Haliaeetus leucocephalus | 194 | 194 | 57% | 1.00E-55 | 52.87 | 329 | XP_010561166.1 |
| RNA/RNP complex-1-interacting phosphatase isoform X1 [Aquila chrysaetos chrysaetos] | Aquila chrysaetos chrysaetos | 194 | 194 | 57% | 1.00E-55 | 52.87 | 329 | XP_029890718.1 |
| PREDICTED: RNA/RNP complex-1-interacting phosphatase [Sturnus vulgaris] | Sturnus vulgaris | 197 | 197 | 57% | 1.00E-55 | 52.87 | 435 | XP_014747572.1 |
| RNA/RNP complex-1-interacting phosphatase [Grus americana] | Grus americana | 193 | 193 | 58% | 1.00E-55 | 51.14 | 309 | XP_054660434.1 |
| DUS11 phosphatase [Rhinopomastus cyanomelas] | Rhinopomastus cyanomelas | 188 | 188 | 56% | 1.00E-55 | 52.63 | 169 | NXO01614.1 |
| RNA/RNP complex-1-interacting phosphatase isoform X2 [Accipiter gentilis] | Accipiter gentilis | 193 | 193 | 57% | 1.00E-55 | 52.87 | 314 | XP_049686641.1 |
| RNA/RNP complex-1-interacting phosphatase isoform X3 [Harpia harpyja] | Harpia harpyja | 192 | 192 | 58% | 2.00E-55 | 52.27 | 271 | XP_052661276.1 |
| DUS11 phosphatase [Dicrurus megarhynchus] | Dicrurus megarhynchus | 188 | 188 | 56% | 2.00E-55 | 53.8 | 169 | NXJ20454.1 |
| RNA/RNP complex-1-interacting phosphatase [Sceloporus undulatus] | Sceloporus undulatus | 193 | 193 | 63% | 2.00E-55 | 50 | 321 | XP_042328494.1 |
| DUS11 phosphatase [Rhagologus leucostigma] | Rhagologus leucostigma | 188 | 188 | 56% | 2.00E-55 | 53.22 | 169 | NXB13795.1 |
| RNA/RNP complex-1-interacting phosphatase isoform X2 [Aythya fuligula] | Aythya fuligula | 191 | 191 | 57% | 2.00E-55 | 52.87 | 269 | XP_032059944.1 |

#### Table S3 cont. Blastp results using vDUSP11 from SWPV2 as query (ARE67303.1)

| | | | | | | | | |
|---|---|---|---|---|---|---|---|---|
| RNA/RNP complex-1-interacting phosphatase isoform X1 [Accipiter gentilis] | Accipiter gentilis | 193 | 193 | 57% | 2.00E-55 | 52.87 | 329 | XP_049686640.1 |
| DUS11 phosphatase [Machaerirhynchus nigripectus] | Machaerirhynchus nigripectus | 188 | 188 | 56% | 2.00E-55 | 53.8 | 169 | NWV88607.1 |
| DUS11 phosphatase [Eulacestoma nigropectus] | Eulacestoma nigropectus | 188 | 188 | 56% | 2.00E-55 | 53.22 | 169 | NXB30747.1 |
| RNA/RNP complex-1-interacting phosphatase [Galemys pyrenaicus] | Galemys pyrenaicus | 191 | 191 | 62% | 2.00E-55 | 51.6 | 275 | KAG8518745.1 |
| DUS11 phosphatase [Aphelocoma coerulescens] | Aphelocoma coerulescens | 188 | 188 | 56% | 3.00E-55 | 53.8 | 169 | NWY21892.1 |
| RNA/RNP complex-1-interacting phosphatase isoform X2 [Harpia harpyja] | Harpia harpyja | 192 | 192 | 57% | 3.00E-55 | 52.87 | 314 | XP_052661275.1 |
| DUS11 phosphatase [Corvus moneduloides] | Corvus moneduloides | 187 | 187 | 56% | 3.00E-55 | 53.8 | 169 | NXD59544.1 |
| RNA/RNP complex-1-interacting phosphatase isoform X3 [Erinaceus europaeus] | Erinaceus europaeus | 193 | 193 | 59% | 3.00E-55 | 52.78 | 333 | XP_060043066.1 |
| DUS11 phosphatase [Lanius ludovicianus] | Lanius ludovicianus | 187 | 187 | 56% | 3.00E-55 | 54.97 | 169 | NWT88744.1 |
| DUS11 phosphatase [Oreocharis arfaki] | Oreocharis arfaki | 187 | 187 | 56% | 3.00E-55 | 53.8 | 169 | NWW12342.1 |
| DUS11 phosphatase [Ifrita kowaldi] | Ifrita kowaldi | 187 | 187 | 56% | 3.00E-55 | 53.8 | 169 | NWW56266.1 |
| RNA/RNP complex-1-interacting phosphatase isoform X1 [Harpia harpyja] | Harpia harpyja | 192 | 192 | 57% | 4.00E-55 | 52.87 | 329 | XP_052661274.1 |
| DUS11 phosphatase [Sylvietta virens] | Sylvietta virens | 187 | 187 | 56% | 4.00E-55 | 53.8 | 169 | NXK68968.1 |
| RNA/RNP complex-1-interacting phosphatase isoform X4 [Anser cygnoides] | Anser cygnoides | 188 | 188 | 57% | 4.00E-55 | 52.3 | 197 | XP_047911010.1 |
| DUS11 phosphatase [Dryoscopus gambensis] | Dryoscopus gambensis | 187 | 187 | 56% | 4.00E-55 | 53.22 | 169 | NWI76823.1 |
| RNA/RNP complex-1-interacting phosphatase isoform X1 [Aythya fuligula] | Aythya fuligula | 192 | 192 | 57% | 4.00E-55 | 52.87 | 312 | XP_032059943.1 |
| DUS11 phosphatase [Mohoua ochrocephala] | Mohoua ochrocephala | 187 | 187 | 56% | 4.00E-55 | 53.22 | 169 | NXA66700.1 |
| RNA/RNP complex-1-interacting phosphatase isoform X4 [Cygnus olor] | Cygnus olor | 188 | 188 | 57% | 4.00E-55 | 52.3 | 197 | XP_040394049.1 |
| DUS11 phosphatase [Vidua macroura] | Vidua macroura | 187 | 187 | 56% | 5.00E-55 | 51.46 | 169 | NXQ02454.1 |
| RNA/RNP complex-1-interacting phosphatase [Aix galericulata] | Aix galericulata | 192 | 192 | 57% | 5.00E-55 | 52.3 | 312 | KAI6073596.1 |
| DUS11 phosphatase [Myiagra hebetior] | Myiagra hebetior | 187 | 187 | 56% | 5.00E-55 | 53.22 | 169 | NXH23758.1 |
| RNA/RNP complex-1-interacting phosphatase isoform X4 [Erinaceus europaeus] | Erinaceus europaeus | 192 | 192 | 59% | 6.00E-55 | 52.78 | 330 | XP_007519010.1 |
| DUS11 phosphatase [Falcunculus frontatus] | Falcunculus frontatus | 187 | 187 | 56% | 6.00E-55 | 53.22 | 169 | NWW22861.1 |
| DUS11 phosphatase [Gymnorhina tibicen] | Gymnorhina tibicen | 187 | 187 | 56% | 7.00E-55 | 53.22 | 169 | NXM43457.1 |
| RNA/RNP complex-1-interacting phosphatase isoform X2 [Cygnus olor] | Cygnus olor | 190 | 190 | 57% | 9.00E-55 | 52.3 | 269 | XP_040394047.1 |
| RNA/RNP complex-1-interacting phosphatase isoform X2 [Anser cygnoides] | Anser cygnoides | 190 | 190 | 57% | 9.00E-55 | 52.3 | 269 | XP_047911008.1 |
| RNA/RNP complex-1-interacting phosphatase isoform X2 [Myiozetetes cayanensis] | Myiozetetes cayanensis | 187 | 187 | 59% | 1.00E-54 | 48.62 | 182 | XP_050181692.1 |
| DUS11 phosphatase [Erpornis zantholeuca] | Erpornis zantholeuca | 186 | 186 | 56% | 1.00E-54 | 53.8 | 169 | NXS78487.1 |
| RNA/RNP complex-1-interacting phosphatase [Cyrtonyx montezumae] | Cyrtonyx montezumae | 191 | 191 | 68% | 1.00E-54 | 45.45 | 305 | XP_065601237.1 |
| RNA/RNP complex-1-interacting phosphatase [Orycteropus afer afer] | Orycteropus afer afer | 190 | 190 | 70% | 1.00E-54 | 47.44 | 289 | XP_007950860.1 |

### Table S3 cont. Blastp results using vDUSP11 from SWPV2 as query (ARE67303.1)

| | | | | | | | | |
|---|---|---|---|---|---|---|---|---|
| RNA/RNP complex-1-interacting phosphatase [Hemicordylus capensis] | Hemicordylus capensis | 193 | 193 | 60% | 1.00E-54 | 50.54 | 378 | XP_053125337.1 |
| DUS11 phosphatase [Oriolus oriolus] | Oriolus oriolus | 186 | 186 | 56% | 1.00E-54 | 53.22 | 169 | NXO07055.1 |
| DUS11 phosphatase [Rhynochetos jubatus] | Rhynochetos jubatus | 186 | 186 | 56% | 1.00E-54 | 52.63 | 169 | NWW93328.1 |
| RNA/RNP complex-1-interacting phosphatase [Alligator mississippiensis] | Alligator mississippiensis | 191 | 191 | 56% | 1.00E-54 | 52.91 | 311 | XP_014467031.1 |
| RNA/RNP complex-1-interacting phosphatase [Alligator mississippiensis] | Alligator mississippiensis | 191 | 191 | 63% | 1.00E-54 | 49.48 | 333 | KYO17197.1 |
| PREDICTED: RNA/RNP complex-1-interacting phosphatase [Gavialis gangeticus] | Gavialis gangeticus | 191 | 191 | 63% | 1.00E-54 | 48.96 | 312 | XP_019369928.1 |
| DUS11 phosphatase [Daphoenositta chrysoptera] | Daphoenositta chrysoptera | 186 | 186 | 56% | 2.00E-54 | 53.22 | 169 | NWV50499.1 |
| DUS11 phosphatase [Edolisoma coerulescens] | Edolisoma coerulescens | 186 | 186 | 56% | 2.00E-54 | 53.8 | 169 | NXH85140.1 |
| RNA/RNP complex-1-interacting phosphatase isoform X1 [Anser cygnoides] | Anser cygnoides | 190 | 190 | 57% | 2.00E-54 | 52.3 | 312 | XP_047911007.1 |
| RNA/RNP complex-1-interacting phosphatase [Ornithorhynchus anatinus] | Ornithorhynchus anatinus | 193 | 193 | 57% | 2.00E-54 | 51.72 | 419 | XP_028921823.1 |
| DUS11 phosphatase [Toxostoma redivivum] | Toxostoma redivivum | 185 | 185 | 56% | 2.00E-54 | 52.05 | 169 | NWS83687.1 |
| DUS11 phosphatase [Sinosuthora webbiana] | Sinosuthora webbiana | 185 | 185 | 56% | 3.00E-54 | 53.8 | 169 | NWR01313.1 |
| RNA/RNP complex-1-interacting phosphatase isoform X3 [Myiozetetes cayanensis] | Myiozetetes cayanensis | 186 | 186 | 58% | 3.00E-54 | 49.43 | 180 | XP_050181693.1 |
| DUS11 phosphatase [Pterocles burchelli] | Pterocles burchelli | 185 | 185 | 56% | 3.00E-54 | 52.05 | 169 | NWU72880.1 |
| RNA/RNP complex-1-interacting phosphatase [Morone saxatilis] | Morone saxatilis | 187 | 187 | 60% | 3.00E-54 | 48.65 | 220 | XP_035524897.1 |
| RNA/RNP complex-1-interacting phosphatase isoform X1 [Cygnus olor] | Cygnus olor | 190 | 190 | 57% | 3.00E-54 | 52.3 | 312 | XP_040394046.1 |
| RNA/RNP complex-1-interacting phosphatase-like [Empidonax traillii] | Empidonax traillii | 186 | 186 | 59% | 3.00E-54 | 48.89 | 184 | XP_027766280.1 |
| RNA/RNP complex-1-interacting phosphatase isoform X2 [Eublepharis macularius] | Eublepharis macularius | 191 | 191 | 68% | 3.00E-54 | 48.33 | 357 | XP_054854289.1 |
| RNA/RNP complex-1-interacting phosphatase isoform X1 [Eublepharis macularius] | Eublepharis macularius | 191 | 191 | 68% | 3.00E-54 | 48.33 | 358 | XP_054854288.1 |
| RNA/RNP complex-1-interacting phosphatase isoform X2 [Hyaena hyaena] | Hyaena hyaena | 190 | 190 | 59% | 3.00E-54 | 51.11 | 330 | XP_039096093.1 |
| DUS11 phosphatase [Ardeotis kori] | Ardeotis kori | 185 | 185 | 56% | 3.00E-54 | 51.46 | 169 | NXE29890.1 |
| RNA/RNP complex-1-interacting phosphatase [Cygnus atratus] | Cygnus atratus | 189 | 189 | 57% | 4.00E-54 | 52.3 | 312 | XP_050571750.1 |
| RNA/RNP complex-1-interacting phosphatase isoform X1 [Phacochoerus africanus] | Phacochoerus africanus | 188 | 188 | 59% | 4.00E-54 | 51.96 | 267 | XP_047637716.1 |
| DUS11 phosphatase [Buphagus erythrorhynchus] | Buphagus erythrorhynchus | 185 | 185 | 56% | 4.00E-54 | 53.22 | 169 | NXU04070.1 |
| RNA/RNP complex-1-interacting phosphatase [Meles meles] | Meles meles | 190 | 190 | 74% | 4.00E-54 | 45.13 | 337 | XP_045834652.1 |
| DUS11 phosphatase [Crocuta crocuta] | Crocuta crocuta | 190 | 190 | 59% | 4.00E-54 | 51.11 | 331 | KAF0879646.1 |
| RNA/RNP complex-1-interacting phosphatase isoform X1 [Hyaena hyaena] | Hyaena hyaena | 190 | 190 | 59% | 4.00E-54 | 51.11 | 331 | XP_039096092.1 |
| RNA/RNP complex-1-interacting phosphatase [Canis lupus familiaris] | Canis lupus familiaris | 190 | 190 | 59% | 4.00E-54 | 52.22 | 332 | XP_005630616.1 |
| RNA/RNP complex-1-interacting phosphatase isoform X1 [Talpa occidentalis] | Talpa occidentalis | 190 | 190 | 61% | 4.00E-54 | 51.05 | 332 | XP_037383543.1 |
| DUS11 phosphatase [Rhipidura dahli] | Rhipidura dahli | 184 | 184 | 56% | 5.00E-54 | 52.05 | 169 | NXI82412.1 |

#### Table S3. Blastp results using vDUSP11 from SWPV2 as query (ARE67303.1)

| | | | | | | | | |
|---|---|---|---|---|---|---|---|---|
| DUS11 phosphatase [Alaudala cheleensis] | Alaudala cheleensis | 184 | 184 | 56% | 5.00E-54 | 53.8 | 169 | NXQ32759.1 |
| RNA/RNP complex-1-interacting phosphatase isoform X1 [Cynocephalus volans] | Cynocephalus volans | 190 | 190 | 59% | 5.00E-54 | 51.38 | 333 | XP_062934399.1 |
| DUS11 phosphatase [Asarcornis scutulata] | Asarcornis scutulata | 184 | 184 | 56% | 5.00E-54 | 53.22 | 169 | NWZ30896.1 |
| PREDICTED: RNA/RNP complex-1-interacting phosphatase-like [Eurypyga helias] | Eurypyga helias | 186 | 186 | 56% | 5.00E-54 | 52.94 | 229 | XP_010148221.1 |
| DUS11 phosphatase [Hirundo rustica] | Hirundo rustica | 184 | 184 | 56% | 5.00E-54 | 52.63 | 169 | NXW80422.1 |
| RNA/RNP complex-1-interacting phosphatase isoform X2 [Oryctolagus cuniculus] | Oryctolagus cuniculus | 190 | 190 | 60% | 5.00E-54 | 52.2 | 330 | XP_008252457.1 |
| RNA/RNP complex-1-interacting phosphatase [Nyctereutes procyonoides] | Nyctereutes procyonoides | 190 | 190 | 59% | 5.00E-54 | 52.22 | 332 | XP_055168628.1 |
| DUS11 phosphatase [Chauna torquata] | Chauna torquata | 184 | 184 | 56% | 5.00E-54 | 52.63 | 169 | NXK49017.1 |

#### Table S4. Avipox and avipox-adjacent viral DUSP11 accession IDs

| Genome length | Description | Scientific Name | Abbreviation | Percent Identity | Acc. Len | Accession |
|---|---|---|---|---|---|---|
| 349821 bp DNA linear VRL 16-FEB-2021 | putative RNA phosphatase [Penguinpox virus 2] | Penguinpox virus 2 | PePV2 | 50.84% | 247 | QRM15716.1 |
| 351909 bp DNA linear VRL 01-JUL-2021 | putative RNA phosphatase [Albatrosspox virus] | Albatrosspox virus | ALPV | 50.84% | 251 | QRM16049.1 |
| 351108 bp DNA linear VRL 13-APR-2017 | SWPV2-ORF080 [Shearwaterpox virus] | Shearwaterpox virus | SWPV2 | 50.00% | 303 | ARE67303.1 |
| 293226 bp DNA linear VRL 25-NOV-2019 | putative RNA phosphatase [Magpiepox virus] | Magpiepox virus | MPPV | 45.79% | 286 | QGM48715.1 |
| 354030 bp DNA linear VRL 20-APR-2022 | DSP DUSP11 [Finch poxvirus] | Finch poxvirus | FNPV | 46.77% | 501 | UOX38764.1 |
| 326929 bp DNA linear VRL 13-APR-2017 | SWPV1-074 [Shearwaterpox virus] | Shearwaterpox virus | SWPV1 | 46.24% | 245 | ARF02681.1 |
| 359853 bp DNA linear VRL 07-JAN-2023 | CNPV085 putative RNA phosphatase [Canarypox virus] | Canarypox virus | CNPV | 44.24% | 403 | NP_955108.1 |
| 342723 bp DNA linear VRL 16-FEB-2021 | putative RNA phosphatase [Mudlarkpox virus] | Mudlarkpox virus | MLPV | 49.16% | 403 | QRM15361.1 |
| 343132 bp DNA linear VRL 13-FEB-2021 | putative RNA phosphatase [Cheloniid poxvirus 1] | Cheloniid poxvirus 1 | ChePV1 | 43.78% | 303 | QRI42800.1 |
| 328768 bp DNA linear VRL 21-OCT 2022 | CRPV-114 [Crowpox virus] | Crowpox virus | CRPW | 60.53% | 42 | UWX11223.1 |
| 298392 bp DNA linear VRL 25-NOV-2022 | MPPV-091 putative RNA phosphatase [Magpiepox virus 2] | Magpiepox virus 2 | MPPV2 | 50.00% | 60 | QZW33383.1 |
| 298392 bp DNA linear VRL 25-NOV-2022 | MPPV-091 putative RNA phosphatase [Magpiepox virus 2] | Magpiepox virus 2 | MPPV2 | * | 32 | QZW33384.1 |
| 328768 bp DNA linear VRL 21-OCT 2022 | CRPV-113 [Crowpox virus] | Crowpox virus | CRPW | * | 96 | UWX11222.1 |

*trucated portion with homology to other vDUSP11

**Table S5. Accession IDs used for phylogenetic analysis of host DUSPs and related poxviral DUSPs**

| Gene | Common Name | Species | Accession |
|---|---|---|---|
| Cdc14A | mouse | *Mus musculus* | NP_001074287.1 |
| Cdc14A | human | *Homo sapiens* | NP_003663.2 |
| Cdc14A | hawaiian crow | *Corvus hawaiiensis* | XP_048168758.1 |
| Cdc14B | human | *Homo sapiens* | NP_201588.1 |
| Cdc14B | mouse | *Mus musculus* | NP_766175.3 |
| Cdc14B | hawaiian crow | *Corvus hawaiiensis* | XP_048148189.1 |
| DUSP1 | human | *Homo sapiens* | NP_004408.1 |
| DUSP1 | mouse | *Mus musculus* | NP_038670.1 |
| DUSP1 | hawaiian crow | *Corvus hawaiiensis* | XP_048176132.1 |
| DUSP10 | human | *Homo sapiens* | NP_009138.1 |
| DUSP10 | mouse | *Mus musculus* | NP_071302.2 |
| DUSP10 | hawaiian crow | *Corvus hawaiiensis* | XP_048154674.1 |
| DUSP11 | human | *Homo sapiens* | NP_003575.2 |
| DUSP11 | mouse | *Mus musculus* | NP_082375.4 |
| DUSP11 | nematode | C.elegans | NP_495959.2 |
| DUSP11 | hawaiian crow | Corvus hawaiiensis | XP_048146001.1 |
| DUSP12 | human | *Homo sapiens* | NP_009171.1 |
| DUSP12 | mouse | *Mus musculus* | NP_075662.2 |
| DUSP12 | nematode | C.elegans | NP_501870.1 |
| DUSP12 | hawaiian crow | *Corvus hawaiiensis* | XP_048150003.1 |
| DUSP13a | mouse | *Mus musculus* | NP_001007269.1 |
| DUSP13a | human | *Homo sapiens* | NP_001007272.1 |
| DUSP13a | hawaiian crow | *Corvus hawaiiensis* | XP_048167325.1 |
| DUSP13b | human | *Homo sapiens* | NP_001307772.1 |
| DUSP13b | mouse | *Mus musculus* | NP_038877.2 |
| DUSP13b | hawaiian crow | *Corvus hawaiiensis* | XP_048166521.1 |
| DUSP14 | mouse | *Mus musculus* | NP_001395670.1 |
| DUSP14 | human | *Homo sapiens* | NP_008957.1 |
| DUSP14 | hawaiian crow | *Corvus hawaiiensis* | XP_048180921.1 |
| DUSP15 | mouse | *Mus musculus* | NP_001152848.1 |
| DUSP15 | human | *Homo sapiens* | NP_001307408.1 |
| DUSP15 | hawaiian crow | *Corvus hawaiiensis* | XP_048178180.1 |
| DUSP16 | human | *Homo sapiens* | NP_085143.1 |
| DUSP16 | mouse | *Mus musculus* | NP_569714.2 |
| DUSP16 | hawaiian crow | *Corvus hawaiiensis* | XP_048158716.1 |
| DUSP17/19 | mouse | *Mus musculus* | NP_077758.1 |
| DUSP17/19 | human | *Homo sapiens* | NP_543152.1 |
| DUSP17/19 | hawaiian crow | *Corvus hawaiiensis* | XP_048165549.1 |
| DUSP18/20 | human | *Homo sapiens* | NP_001291723.1 |
| DUSP18/20 | mouse | *Mus musculus* | NP_776106.1 |
| DUSP2 | human | *Homo sapiens* | NP_004409.1 |
| DUSP2 | mouse | *Mus musculus* | NP_034220.2 |
| DUSP21 | human | *Homo sapiens* | NP_071359.3 |
| DUSP21 | mouse | *Mus musculus* | NP_082844.1 |

**Table S5 cont. Accession IDs used for phylogenetic analysis of host DUSPs and related poxviral DUSPs**

| | | | |
|---|---|---|---|
| DUSP22 | human | *Homo sapiens* | NP_001273484.1 |
| DUSP22 | mouse | *Mus musculus* | NP_598829.1 |
| DUSP22 | hawaiian crow | *Corvus hawaiiensis* | XP_048166191.1 |
| DUSP23/25 | human | *Homo sapiens* | NP_001306587.1 |
| DUSP23/25 | mouse | *Mus musculus* | NP_081001.1 |
| DUSP23/25 | hawaiian crow | *Corvus hawaiiensis* | XP_048144571.1 |
| DUSP26 | human | *Homo sapiens* | NP_001292044.1 |
| DUSP26 | mouse | *Mus musculus* | NP_001344152.1 |
| DUSP26 | hawaiian crow | *Corvus hawaiiensis* | XP_048143456.1 |
| DUSP27/29 | human | *Homo sapiens* | NP_001013848.1 |
| DUSP27/29 | mouse | *Mus musculus* | NP_001028516.2 |
| DUSP27/29 | hawaiian crow | *Corvus hawaiiensis* | XP_048166519.1 |
| DUSP28 | human | *Homo sapiens* | NP_001028747.1 |
| DUSP28 | mouse | *Mus musculus* | NP_780327.1 |
| DUSP28 | hawaiian crow | *Corvus hawaiiensis* | XP_048170675.1 |
| DUSP3 | human | *Homo sapiens* | NP_004081.1 |
| DUSP3 | mouse | *Mus musculus* | NP_082483.1 |
| DUSP3 | hawaiian crow | *Corvus hawaiiensis* | XP_048171990.1 |
| DUSP4 | human | *Homo sapiens* | NP_001385.1 |
| DUSP4 | mouse | *Mus musculus* | NP_795907.1 |
| DUSP4 | hawaiian crow | *Corvus hawaiiensis* | XP_048158952.1 |
| DUSP5 | mouse | *Mus musculus* | NP_001078859.1 |
| DUSP5 | human | *Homo sapiens* | NP_004410.3 |
| DUSP5 | hawaiian crow | *Corvus hawaiiensis* | XP_048167723.1 |
| DUSP6 | human | *Homo sapiens* | NP_001937.2 |
| DUSP6 | mouse | *Mus musculus* | NP_080544.1 |
| DUSP6 | hawaiian crow | *Corvus hawaiiensis* | XP_048158193.1 |
| DUSP7 | human | *Homo sapiens* | NP_001938.2 |
| DUSP7 | mouse | *Mus musculus* | NP_703189.3 |
| DUSP7 | hawaiian crow | *Corvus hawaiiensis* | XP_048171802.1 |
| DUSP8 | human | *Homo sapiens* | NP_004411.2 |
| DUSP8 | mouse | *Mus musculus* | NP_032774.1 |
| DUSP8 | hawaiian crow | *Corvus hawaiiensis* | XP_048161880.1 |
| DUSP9 | human | *Homo sapiens* | NP_001386.1 |
| DUSP9 | mouse | *Mus musculus* | NP_083628.3 |
| EPM2A | human | *Homo sapiens* | NP_001018051.1 |
| EPM2A | mouse | *Mus musculus* | NP_034276.2 |
| EPM2A | hawaiian crow | *Corvus hawaiiensis* | XP_048153694.1 |
| KAP-1 | human | *Homo sapiens* | NP_001158230.1 |
| KAP-1 | mouse | *Mus musculus* | NP_032949.3 |
| KAP-1 | hawaiian crow | *Corvus hawaiiensis* | XP_048176707.1 |
| PRL1 | mouse | *Mus musculus* | NP_001158217.1 |
| PRL1 | human | *Homo sapiens* | NP_003454.1 |
| PRL1 | hawaiian crow | *Corvus hawaiiensis* | XP_048153462.1 |
| PRL2 | human | *Homo sapiens* | NP_001182029.1 |

**Table S5 cont. Accession IDs used for phylogenetic analysis of host DUSPs and related poxviral DUSPs**

| | | | |
|---|---|---|---|
| PRL2 | mouse | *Mus musculus* | NP_033000.1 |
| PRL2 | hawaiian crow | *Corvus hawaiiensis* | XP_048182961.1 |
| PRL3 | mouse | *Mus musculus* | NP_001159860.1 |
| PRL3 | human | *Homo sapiens* | NP_009010.2 |
| PRL3 | hawaiian crow | *Corvus hawaiiensis* | XP_048142652.1 |
| PTEN | human | *Homo sapiens* | NP_000305.3 |
| PTEN | mouse | *Mus musculus* | NP_032986.1 |
| PTEN | hawaiian crow | *Corvus hawaiiensis* | XP_048166888.1 |
| PTEN2 | mouse | *Mus musculus* | NP_954866.2 |
| PTEN2 | hawaiian crow | *Corvus hawaiiensis* | XP_048151148.1 |
| PTPC1 | human | *Homo sapiens* | NP_001240758.1 |
| PTPC1 | mouse | *Mus musculus* | NP_997115.1 |
| PTPC1 | hawaiian crow | *Corvus hawaiiensis* | XP_048171542.1 |
| PTPMT1 | mouse | *Mus musculus* | NP_079852.1 |
| PTPMT1 | human | *Homo sapiens* | NP_783859.1 |
| PTPMT1 | hawaiian crow | *Corvus hawaiiensis* | XP_048163082.1 |
| RNGTT | nematode | *C.elegans* | NP_001020979.1 |
| RNGTT | human | *Homo sapiens* | NP_003791.3 |
| RNGTT | mouse | *Mus musculus* | NP_036014.1 |
| RNGTT | hawaiian crow | *Corvus hawaiiensis* | XP_048154745.1 |
| SSH1 | human | *Homo sapiens* | NP_061857.3 |
| SSH1 | mouse | *Mus musculus* | NP_932777.2 |
| SSH1 | hawaiian crow | *Corvus hawaiiensis* | XP_048178835.1 |
| SSH2 | human | *Homo sapiens* | NP_001269058.1 |
| SSH2 | mouse | *Mus musculus* | NP_808378.2 |
| SSH2 | hawaiian crow | *Corvus hawaiiensis* | XP_048180293.1 |
| SSH3 | human | *Homo sapiens* | NP_060327.3 |
| SSH3 | mouse | *Mus musculus* | NP_932781.1 |
| SSH3 | hawaiian crow | *Corvus hawaiiensis* | XP_048162621.1 |
| STYX | human | *Homo sapiens* | NP_001124173.1 |
| STYX | mouse | *Mus musculus* | NP_062611.2 |
| STYX | hawaiian crow | *Corvus hawaiiensis* | XP_048161738.1 |
| TNS1 | human | *Homo sapiens* | NP_001294951.1 |
| TNS1 | mouse | *Mus musculus* | NP_082160.3 |
| TNS1 | hawaiian crow | *Corvus hawaiiensis* | XP_048164725.1 |
| TNS2 | human | *Homo sapiens* | NP_056134.2 |
| TNS2 | mouse | *Mus musculus* | NP_705761.2 |
| TNS2 | hawaiian crow | *Corvus hawaiiensis* | XP_048164730.1 |
| TPTE2 | human | *Homo sapiens* | NP_954863.2 |
| TPTE2 | mouse | *Mus musculus* | NP_954866.2 |
| TPTE2 | hawaiian crow | *Corvus hawaiiensis* | XP_048151150.1 |

**Table S5 cont. Accession IDs used for phylogenetic analysis of host DUSPs and related poxviral DUSPs**

| Gene | Virus | Abbreviaton | Accession |
|---|---|---|---|
| VH1 | Vaccinia virus | VACV | AAB59836.1 |
| VH1 | Shearwaterpox virus 2 | SWPV2 | ARE67406.1 |
| VH1 | Shearwaterpox virus | SWPV1 | ARF02737.1 |
| VH1 | Variola virus | VARV | NP_042128.1 |
| VH1 | Canarypox virus | CNPV | NP_955206.1 |
| VH1 | Magpiepox virus | MPPV | QGM48814.1 |
| VH1 | Cheloniid poxvirus 1 | ChePV1 | QRI42902.1 |
| VH1 | Mudlarkpox virus | MLPV | QRM15459.1 |
| VH1 | Penguinpox virus 2 | PEPV2 | QRM15814.1 |
| VH1 | Albatrosspox virus | ALPV | QRM16149.1 |
| VH1 | Magpiepox virus 2 | MPPV2 | QZW33497.1 |
| VH1 | Finch poxvirus | FNPV | UOX38765.1 |
| VH1 | Crowpox virus | CRPV | UWX11342.1 |
| vDUSP11 | Shearwaterpox virus | SWPV2 | ARE67303.1 |
| vDUSP11 | Shearwaterpox virus | SWPV1 | ARF02681.1 |
| vDUSP11 | Canarypox virus | CNPV | NP_955108.1 |
| vDUSP11 | Magpiepox virus | MPPV | QGM48715.1 |
| vDUSP11 | Cheloniid poxvirus 1 | ChePV1 | QRI42800.1 |
| vDUSP11 | Mudlarkpox virus | MLPV | QRM15361.1 |
| vDUSP11 | Penguinpox virus 2 | PEPV2 | QRM15716.1 |
| vDUSP11 | Albatrosspox virus | ALPV | QRM16049.1 |
| vDUSP11 | Finch poxvirus | FNPV | UOX38764.1 |

Table S6. Genes surrounding the genomic locations of avipox vDUSP11 from synteny analysis

| Unifying | Description* | virus-orf | position | Orientation | Virus | Accession |
|---|---|---|---|---|---|---|
| SWPV2-075 | early transcription factor small VETFS | ALPV-084 | -5 | R | Albatrosspox virus | QRM16044.1 |
| SWPV2-076 | Ig-like domain protein | ALPV-085 | -4 | R | Albatrosspox virus | QRM16045.1 |
| SWPV2-077 | NTPase, DNA replication | ALPV-086 | -3 | R | Albatrosspox virus | QRM16046.1 |
| SWPV2-078 | CC chemokine-like protein | ALPV-087 | -2 | R | Albatrosspox virus | QRM16047.1 |
| SWPV2-079 | uracil DNA glycosylase | ALPV-088 | -1 | R | Albatrosspox virus | QRM16048.1 |
| SWPV2-080 | putative RNA phosphatase | ALPV-089 | 0 | R | Albatrosspox virus | QRM16049.1 |
| SWPV2-204 | conserved hypothetical protein | ALPV-090 | 1 | R | Albatrosspox virus | QRM16050.1 |
| SWPV2-081 | TNFR-like protein | ALPV-091 | 2 | F | Albatrosspox virus | QRM16051.1 |
| SWPV2-082 | putative glutathione peroxidase | ALPV-092 | 3 | F | Albatrosspox virus | QRM16052.1 |
| SWPV2-083 | conserved hypothetical protein | ALPV-093 | 4 | F | Albatrosspox virus | QRM16053.1 |
| SWPV2-084 | conserved hypothetical protein | ALPV-094 | 5 | R | Albatrosspox virus | QRM16054.1 |
| SWPV2-075 | early transcription factor small VETFS | CNPV-080 | -5 | R | Canarypox virus | NP_955103.1 |
| SWPV2-076 | Ig-like domain protein | CNPV-081 | -4 | R | Canarypox virus | NP_955104.1 |
| SWPV2-077 | NTPase, DNA replication | CNPV-082 | -3 | R | Canarypox virus | NP_955105.1 |
| SWPV2-078 | CC chemokine-like protein | CNPV-083 | -2 | R | Canarypox virus | NP_955106.1 |
| SWPV2-079 | uracil DNA glycosylase | CNPV-084 | -1 | R | Canarypox virus | NP_955107.1 |
| SWPV2-080 | putative RNA phosphatase | CNPV-085 | 0 | R | Canarypox virus | NP_955108.1 |
| SWPV2-081 | TNFR-like protein | CNPV-086 | 1 | F | Canarypox virus | NP_955109.1 |
| SWPV2-082 | putative glutathione peroxidase | CNPV-087 | 2 | F | Canarypox virus | NP_955100.1 |
| SWPV2-083 | conserved hypothetical protein | CNPV-088 | 3 | F | Canarypox virus | NP_955111.1 |
| SWPV2-084 | conserved hypothetical protein | CNPV-089 | 4 | R | Canarypox virus | NP_955112.1 |
| SWPV2-085 | conserved hypothetical protein | CNPV-090 | 5 | R | Canarypox virus | NP_955113.1 |
| SWPV2-075 | early transcription factor small VETFS | ChePV1-077 | -5 | R | Cheloniid poxvirus 1 | QRI42795.1 |
| SWPV2-076 | Ig-like domain protein | ChePV1-078 | -4 | R | Cheloniid poxvirus 1 | QRI42796.1 |
| SWPV2-077 | NTPase, DNA replication | ChePV1-079 | -3 | R | Cheloniid poxvirus 1 | QRI42797.1 |
| SWPV2-078 | CC chemokine-like protein | ChePV1-080 | -2 | R | Cheloniid poxvirus 1 | QRI42798.1 |
| SWPV2-079 | uracil DNA glycosylase | ChePV1-081 | -1 | R | Cheloniid poxvirus 1 | QRI42799.1 |
| SWPV2-080 | putative RNA phosphatase | ChePV1-082 | 0 | R | Cheloniid poxvirus 1 | QRI42800.1 |
| SWPV2-081 | TNFR-like protein | ChePV1-083 | 1 | F | Cheloniid poxvirus 1 | QRI42801.1 |
| SWPV2-082 | putative glutathione peroxidase | ChePV1-084 | 2 | F | Cheloniid poxvirus 1 | QRI42802.1 |
| SWPV2-084 | conserved hypothetical protein | ChePV1-085 | 3 | R | Cheloniid poxvirus 1 | QRI42803.1 |
| SWPV2-085 | conserved hypothetical protein | ChePV1-086 | 4 | R | Cheloniid poxvirus 1 | QRI42804.1 |
| SWPV2-086 | HT motif protein | ChePV1-087 | 5 | R | Cheloniid poxvirus 1 | QRI42805.1 |
| SWPV2-076 | Ig-like domain protein | CRPV-108 | -5 | R | Crowpox virus | UWX11217.1 |
| Unique | hypothetical protein | CRPV-109 | -4 | R | Crowpox virus | UWX11218.1 |
| SWPV2-077 | NTPase, DNA replication | CRPV-110 | -3 | R | Crowpox virus | UWX11219.1 |
| CNPV-215 | viral CC-type chemokine | CRPV-111 | -2 | R | Crowpox virus | UWX11220.1 |
| SWPV2-079 | uracil DNA glycosylase | CRPV-112 | -1 | R | Crowpox virus | UWX11221.1 |
| SWPV2-080 | putative RNA phosphatase | CRPV-113 | 0 | R | Crowpox virus | UWX11222.1 |
| SWPV2-080 | putative RNA phosphatase | CRPV-114 | 0 | R | Crowpox virus | UWX11223.1 |
| SWPV2-204 | conserved hypothetical protein | CRPV-115 | 1 | R | Crowpox virus | UWX11224.1 |
| SWPV2-081 | TNFR-like protein | CRPV-116 | 2 | F | Crowpox virus | UWX11225.1 |
| CRPV-117 | N1R/p28-like protein | CRPV-117 | 3 | R | Crowpox virus | UWX11226.1 |
| SWPV2-082 | putative glutathione peroxidase | CRPV-118 | 4 | F | Crowpox virus | UWX11227.1 |
| SWPV2-082 | putative glutathione peroxidase | CRPV-119 | 5 | F | Crowpox virus | UWX11228.1 |
| SWPV2-083 | conserved hypothetical protein | CRPV-120 | 6 | F | Crowpox virus | UWX11229.1 |
| SWPV2-075 | early transcription factor small VETFS |  | -6 | R | Finch poxvirus | UOX38756.1 |
| SWPV2-076 | Ig-like domain protein |  | -5 | R | Finch poxvirus | UOX38805.1 |
| SWPV2-076 | Ig-like domain protein |  | -4 | R | Finch poxvirus | UOX38794.1 |
| SWPV2-077 | NTPase, DNA replication |  | -3 | R | Finch poxvirus | UOX38855.1 |
| CNPV-215 | viral CC-type chemokine |  | -2 | R | Finch poxvirus | UOX38863.1 |
| SWPV2-079 | uracil DNA glycosylase |  | -1 | R | Finch poxvirus | UOX38862.1 |
| SWPV2-080 | putative RNA phosphatase |  | 0 | R | Finch poxvirus | UOX38764.1 |
| SWPV2-204 | non structural virulence protein |  | 1 | R | Finch poxvirus | UOX38813.1 |
| SWPV2-081 | TNFR-like protein |  | 2 | F | Finch poxvirus | UOX38715.1 |
| SWPV2-082 | putative glutathione peroxidase |  | 3 | F | Finch poxvirus | UOX38589.1 |
| SWPV2-082 | putative glutathione peroxidase |  | 4 | F | Finch poxvirus | UOX38588.1 |
| SWPV2-083 | conserved hypothetical protein |  | 5 | F | Finch poxvirus | UOX38613.1 |
| SWPV2-084 | conserved hypothetical protein |  | 6 | R | Finch poxvirus | UOX38785.1 |
| SWPV2-075 | early transcription factor small VETFS | MPPV-086 | -5 | R | Magpiepox virus | QGM48710.1 |
| SWPV2-076 | Ig-like domain protein | MPPV-087 | -4 | R | Magpiepox virus | QGM48711.1 |
| SWPV2-077 | NTPase, DNA replication | MPPV-088 | -3 | R | Magpiepox virus | QGM48712.1 |
| SWPV2-078 | CC chemokine-like protein | MPPV-089 | -2 | R | Magpiepox virus | QGM48713.1 |
| SWPV2-079 | uracil DNA glycosylase | MPPV-090 | -1 | R | Magpiepox virus | QGM48714.1 |
| SWPV2-080 | putative RNA phosphatase | MPPV-091 | 0 | R | Magpiepox virus | QGM48715.1 |
| SWPV2-081 | TNFR-like protein | MPPV-092 | 1 | F | Magpiepox virus | QGM48716.1 |
| SWPV2-082 | putative glutathione peroxidase | MPPV-093 | 2 | F | Magpiepox virus | QGM48717.1 |
| SWPV2-084 | conserved hypothetical protein | MPPV-094 | 3 | R | Magpiepox virus | QGM48718.1 |
| SWPV2-085 | conserved hypothetical protein | MPPV-095 | 4 | R | Magpiepox virus | QGM48719.1 |

Table S6 cont. Genes surrounding the genomic locations of avipox vDUSP11 from synteny analysis

| | | | | | | |
|---|---|---|---|---|---|---|
| SWPV2-086 | HT motif protein | MPPV-096 | 5 | R | Magpiepox virus | QGM48720.1 |
| SWPV2-076 | Ig-like domain protein | MPPV2-112 | -7 | R | Magpiepox virus 2 | QZW33376.1 |
| Unique | hypothetical protein | MPPV2-113 | -6 | R | Magpiepox virus 2 | QZW33377.1 |
| Unique | hypothetical protein | MPPV2-114 | -5 | R | Magpiepox virus 2 | QZW33378.1 |
| SWPV2-077 | NTPase, DNA replication | MPPV2-115 | -4 | R | Magpiepox virus 2 | QZW33379.1 |
| SWPV2-078 | CC chemokine-like protein | MPPV2-116 | -3 | R | Magpiepox virus 2 | QZW33380.1 |
| SWPV2-078 | CC chemokine-like protein | MPPV2-117 | -2 | R | Magpiepox virus 2 | QZW33381.1 |
| SWPV2-079 | uracil DNA glycosylase | MPPV2-118 | -1 | R | Magpiepox virus 2 | QZW33382.1 |
| SWPV2-080 | putative RNA phosphatase | MPPV2-119 | 0 | R | Magpiepox virus 2 | QZW33383.1 |
| SWPV2-080 | putative RNA phosphatase | MPPV2-120 | 0 | R | Magpiepox virus 2 | QZW33384.1 |
| SWPV2-081 | TNFR-like protein | MPPV2-121 | 1 | F | Magpiepox virus 2 | QZW33385.1 |
| SWPV2-082 | putative glutathione peroxidase | MPPV2-122 | 2 | F | Magpiepox virus 2 | QZW33386.1 |
| SWPV2-082 | putative glutathione peroxidase | MPPV2-123 | 3 | F | Magpiepox virus 2 | QZW33387.1 |
| SWPV2-083 | conserved hypothetical protein | MPPV2-124 | 4 | F | Magpiepox virus 2 | QZW33388.1 |
| SWPV2-084 | conserved hypothetical protein | MPPV2-125 | 5 | R | Magpiepox virus 2 | QZW33389.1 |
| SWPV2-085 | conserved hypothetical protein | MPPV2-126 | 6 | R | Magpiepox virus 2 | QZW33390.1 |
| SWPV2-075 | early transcription factor small VETFS | MLPV-077 | -5 | R | Mudlarkpox virus | QRM15356.1 |
| SWPV2-076 | Ig-like domain protein | MLPV-078 | -4 | R | Mudlarkpox virus | QRM15357.1 |
| SWPV2-077 | NTPase, DNA replication | MLPV-079 | -3 | R | Mudlarkpox virus | QRM15358.1 |
| SWPV2-078 | CC chemokine-like protein | MLPV-080 | -2 | R | Mudlarkpox virus | QRM15359.1 |
| SWPV2-079 | uracil DNA glycosylase | MLPV-081 | -1 | R | Mudlarkpox virus | QRM15360.1 |
| SWPV2-080 | putative RNA phosphatase | MLPV-082 | 0 | R | Mudlarkpox virus | QRM15361.1 |
| SWPV2-081 | TNFR-like protein | MLPV-083 | 1 | F | Mudlarkpox virus | QRM15362.1 |
| SWPV2-082 | putative glutathione peroxidase | MLPV-084 | 2 | F | Mudlarkpox virus | QRM15363.1 |
| SWPV2-084 | conserved hypothetical protein | MLPV-085 | 3 | R | Mudlarkpox virus | QRM15364.1 |
| SWPV2-085 | conserved hypothetical protein | MLPV-086 | 4 | R | Mudlarkpox virus | QRM15365.1 |
| SWPV2-086 | HT motif protein | MLPV-087 | 5 | R | Mudlarkpox virus | QRM15366.1 |
| SWPV2-075 | early transcription factor small VETFS | PEPV2-080 | -5 | R | Penguinpox virus 2 | QRM15711.1 |
| SWPV2-076 | Ig-like domain protein | PEPV2-081 | -4 | R | Penguinpox virus 2 | QRM15712.1 |
| SWPV2-077 | NTPase, DNA replication | PEPV2-082 | -3 | R | Penguinpox virus 2 | QRM15713.1 |
| SWPV2-078 | CC chemokine-like protein | PEPV2-083 | -2 | R | Penguinpox virus 2 | QRM15714.1 |
| SWPV2-079 | uracil DNA glycosylase | PEPV2-084 | -1 | R | Penguinpox virus 2 | QRM15715.1 |
| SWPV2-080 | putative RNA phosphatase | PEPV2-085 | 0 | R | Penguinpox virus 2 | QRM15716.1 |
| SWPV2-204 | conserved hypothetical protein | PEPV2-086 | 1 | R | Penguinpox virus 2 | QRM15717.1 |
| SWPV2-081 | TNFR-like protein | PEPV2-087 | 2 | F | Penguinpox virus 2 | QRM15718.1 |
| CRPV-117 | N1R/p28-like protein | PEPV2-088 | 3 | R | Penguinpox virus 2 | QRM15719.1 |
| SWPV2-082 | putative glutathione peroxidase | PEPV2-089 | 4 | F | Penguinpox virus 2 | QRM15720.1 |
| SWPV2-083 | conserved hypothetical protein | PEPV2-90 | 5 | F | Penguinpox virus 2 | QRM15721.1 |
| SWPV2-075 | early transcription factor small VETFS | SWPV1-069 | -5 | R | Shearwaterpox virus | ARE67597.1 |
| SWPV2-077 | NTPase, DNA replication | SWPV1-070 | -4 | R | Shearwaterpox virus | ARF02677.1 |
| SWPV2-078 | CC chemokine-like protein | SWPV1-071 | -3 | R | Shearwaterpox virus | ARF02678.1 |
| CNPV-215 | viral CC-type chemokine | SWPV1-072 | -2 | R | Shearwaterpox virus | ARF02679.1 |
| SWPV2-079 | uracil DNA glycosylase | SWPV1-073 | -1 | R | Shearwaterpox virus | ARF02680.1 |
| SWPV2-080 | putative RNA phosphatase | SWPV1-074 | 0 | R | Shearwaterpox virus | ARF02681.1 |
| SWPV2-204 | conserved hypothetical protein | SWPV1-075 | 1 | R | Shearwaterpox virus | ARF02682.1 |
| SWPV2-081 | TNFR-like protein | SWPV1-076 | 2 | F | Shearwaterpox virus | ARF02683.1 |
| CRPV-117 | N1R/p28-like protein | SWPV1-077 | 3 | R | Shearwaterpox virus | ARF02684.1 |
| SWPV2-083 | conserved hypothetical protein | SWPV1-078 | 4 | F | Shearwaterpox virus | ARE67659.1 |
| SWPV2-084 | conserved hypothetical protein | SWPV1-079 | 5 | R | Shearwaterpox virus | ARF02685.1 |
| SWPV2-075 | early transcription factor small VETFS | SWPV2-075 | -5 | R | Shearwaterpox virus 2 | ARE67298.1 |
| SWPV2-076 | Ig-like domain protein | SWPV2-076 | -4 | R | Shearwaterpox virus 2 | ARE67299.1 |
| SWPV2-077 | NTPase, DNA replication | SWPV2-077 | -3 | R | Shearwaterpox virus 2 | ARE67300.1 |
| SWPV2-078 | CC chemokine-like protein | SWPV2-078 | -2 | R | Shearwaterpox virus 2 | ARE67301.1 |
| SWPV2-079 | uracil DNA glycosylase | SWPV2-079 | -1 | R | Shearwaterpox virus 2 | ARE67302.1 |
| SWPV2-080 | putative RNA phosphatase | SWPV2-080 | 0 | R | Shearwaterpox virus 2 | ARE67303.1 |
| SWPV2-081 | TNFR-like protein | SWPV2-081 | 1 | F | Shearwaterpox virus 2 | ARE67304.1 |
| SWPV2-082 | putative glutathione peroxidase | SWPV2-082 | 2 | F | Shearwaterpox virus 2 | ARE67305.1 |
| SWPV2-083 | conserved hypothetical protein | SWPV2-083 | 3 | F | Shearwaterpox virus 2 | ARE67306.1 |
| SWPV2-084 | conserved hypothetical protein | SWPV2-084 | 4 | R | Shearwaterpox virus 2 | ARE67307.1 |
| SWPV2-085 | conserved hypothetical protein | SWPV2-085 | 5 | R | Shearwaterpox virus 2 | ARE67308.1 |

*Description refers to putative ORF identity determined through homology analysis or as directly indicated by NCBI (37) entry.

#### Table S7. Blastp results using penguinpox virus 2 (PEPV2) viral DUSP11 as query (QRM15716.1)

| Description | Scientific Name | Max Score | Total Score | Query Cover | E value | Per. ident | Acc. Len | Accession |
|---|---|---|---|---|---|---|---|---|
| ***putative RNA phosphatase [Penguinpox virus 2]*** | ***Penguinpox virus 2*** | ***508*** | ***508*** | ***100%*** | ***0*** | ***100*** | ***247*** | ***QRM15716.1*** |
| *putative RNA phosphatase [Albatrosspox virus]* | *Albatrosspox virus* | *502* | *502* | *100%* | *4.00E-179* | *98.41* | *251* | *QRM16049.1* |
| *SWPV2-ORF080 [Shearwaterpox virus]* | *Shearwaterpox virus* | *482* | *482* | *94%* | *4.00E-170* | *98.72* | *303* | *ARE67303.1* |
| *putative RNA phosphatase [Magpiepox virus]* | *Magpiepox virus* | *478* | *478* | *94%* | *6.00E-169* | *97.86* | *286* | *QGM48715.1* |
| *CNPV085 putative RNA phosphatase [Canarypox virus]* | *Canarypox virus* | *457* | *511* | *100%* | *1.00E-158* | *91.03* | *403* | *NP_955108.1* |
| *putative RNA phosphatase [Mudlarkpox virus]* | *Mudlarkpox virus* | *406* | *460* | *100%* | *1.00E-138* | *81.62* | *403* | *QRM15361.1* |
| *putative RNA phosphatase [Cheloniid poxvirus 1]* | *Cheloniid poxvirus 1* | *388* | *388* | *94%* | *3.00E-133* | *78.21* | *303* | *QRI42800.1* |
| *DSP DUSP11 [Finch poxvirus]* | *Finch poxvirus* | *389* | *389* | *94%* | *8.00E-131* | *76.17* | *501* | *UOX38764.1* |
| *SWPV1-074 [Shearwaterpox virus]* | *Shearwaterpox virus* | *276* | *276* | *100%* | *1.00E-89* | *59.84* | *245* | *ARF02681.1* |
| RNA/RNP complex-1-interacting phosphatase [Corvus hawaiiensis] | Corvus hawaiiensis | 199 | 199 | 71% | 6.00E-59 | 53.41 | 305 | XP_048146001.1 |
| RNA/RNP complex-1-interacting phosphatase [Corvus cornix cornix] | Corvus cornix cornix | 199 | 199 | 71% | 4.00E-58 | 53.41 | 349 | XP_039420096.1 |
| DUS11 phosphatase [Upupa epops] | Upupa epops | 191 | 191 | 69% | 3.00E-57 | 53.22 | 169 | NWU89504.1 |
| RNA/RNP complex-1-interacting phosphatase [Calypte anna] | Calypte anna | 195 | 195 | 70% | 4.00E-57 | 52.87 | 320 | XP_030319948.1 |
| RNA/RNP complex-1-interacting phosphatase isoform X3 [Accipiter gentilis] | Accipiter gentilis | 194 | 194 | 70% | 5.00E-57 | 52.87 | 271 | XP_049686642.1 |
| RNA/RNP complex-1-interacting phosphatase isoform X2 [Aquila chrysaetos chrysaetos] | Aquila chrysaetos chrysaetos | 195 | 195 | 70% | 5.00E-57 | 52.87 | 314 | XP_029890719.1 |
| RNA/RNP complex-1-interacting phosphatase isoform X3 [Aquila chrysaetos chrysaetos] | Aquila chrysaetos chrysaetos | 193 | 193 | 70% | 6.00E-57 | 52.87 | 271 | XP_040984244.1 |
| DUS11 phosphatase [Struthidea cinerea] | Struthidea cinerea | 190 | 190 | 69% | 7.00E-57 | 54.39 | 169 | NXB60100.1 |
| DUS11 phosphatase [Pachycephala philippinensis] | Pachycephala philippinensis | 189 | 189 | 69% | 7.00E-57 | 53.8 | 169 | NXI05060.1 |
| RNA/RNP complex-1-interacting phosphatase isoform X3 [Aythya fuligula] | Aythya fuligula | 191 | 191 | 70% | 8.00E-57 | 52.87 | 197 | XP_032059945.1 |
| RNA/RNP complex-1-interacting phosphatase isoform X1 [Aquila chrysaetos chrysaetos] | Aquila chrysaetos chrysaetos | 195 | 195 | 70% | 8.00E-57 | 52.87 | 329 | XP_029890718.1 |
| DUS11 phosphatase [Rhinopomastus cyanomelas] | Rhinopomastus cyanomelas | 189 | 189 | 69% | 1.00E-56 | 52.63 | 169 | NXO01614.1 |
| PREDICTED: RNA/RNP complex-1-interacting phosphatase [Sturnus vulgaris] | Sturnus vulgaris | 197 | 197 | 70% | 1.00E-56 | 52.87 | 435 | XP_014747572.1 |
| RNA/RNP complex-1-interacting phosphatase [Galemys pyrenaicus] | Galemys pyrenaicus | 193 | 193 | 73% | 1.00E-56 | 53.04 | 275 | KAG8518745.1 |
| PREDICTED: RNA/RNP complex-1-interacting phosphatase isoform X3 [Haliaeetus leucocephalus] | Haliaeetus leucocephalus | 194 | 194 | 70% | 1.00E-56 | 52.87 | 329 | XP_010561166.1 |
| RNA/RNP complex-1-interacting phosphatase isoform X2 [Accipiter gentilis] | Accipiter gentilis | 194 | 194 | 70% | 1.00E-56 | 52.87 | 314 | XP_049686641.1 |
| RNA/RNP complex-1-interacting phosphatase isoform X3 [Harpia harpyja] | Harpia harpyja | 192 | 192 | 71% | 1.00E-56 | 52.27 | 271 | XP_052661276.1 |
| RNA/RNP complex-1-interacting phosphatase isoform X3 [Erinaceus europaeus] | Erinaceus europaeus | 194 | 194 | 72% | 2.00E-56 | 52.78 | 333 | XP_060043066.1 |
| RNA/RNP complex-1-interacting phosphatase isoform X1 [Accipiter gentilis] | Accipiter gentilis | 194 | 194 | 70% | 2.00E-56 | 52.87 | 329 | XP_049686640.1 |
| uncharacterized protein LOC116435472 [Corvus moneduloides] | Corvus moneduloides | 199 | 199 | 72% | 2.00E-56 | 52.51 | 535 | XP_031947699.1 |
| RNA/RNP complex-1-interacting phosphatase [Grus americana] | Grus americana | 193 | 193 | 71% | 2.00E-56 | 51.14 | 309 | XP_054660434.1 |

**Table S7 cont. Blastp results using penguinpox virus 2 (PEPV2) viral DUSP11 as query (QRM15716.1)**

| | | | | | | | | |
|---|---|---|---|---|---|---|---|---|
| RNA/RNP complex-1-interacting phosphatase [Sceloporus undulatus] | Sceloporus undulatus | 194 | 194 | 74% | 2.00E-56 | 50.8 | 321 | XP_042328494.1 |
| DUS11 phosphatase [Dicrurus megarhynchus] | Dicrurus megarhynchus | 188 | 188 | 69% | 2.00E-56 | 53.8 | 169 | NXJ20454.1 |
| RNA/RNP complex-1-interacting phosphatase isoform X4 [Erinaceus europaeus] | Erinaceus europaeus | 194 | 194 | 72% | 2.00E-56 | 52.78 | 330 | XP_007519010.1 |
| RNA/RNP complex-1-interacting phosphatase isoform X2 [Aythya fuligula] | Aythya fuligula | 192 | 192 | 70% | 3.00E-56 | 52.87 | 269 | XP_032059944.1 |
| DUS11 phosphatase [Rhagologus leucostigma] | Rhagologus leucostigma | 188 | 188 | 69% | 3.00E-56 | 53.22 | 169 | NXB13795.1 |
| DUS11 phosphatase [Eulacestoma nigropectus] | Eulacestoma nigropectus | 188 | 188 | 69% | 3.00E-56 | 53.22 | 169 | NXB30747.1 |
| DUS11 phosphatase [Aphelocoma coerulescens] | Aphelocoma coerulescens | 188 | 188 | 69% | 3.00E-56 | 53.8 | 169 | NWY21892.1 |
| RNA/RNP complex-1-interacting phosphatase isoform X2 [Harpia harpyja] | Harpia harpyja | 193 | 193 | 70% | 3.00E-56 | 52.87 | 314 | XP_052661275.1 |
| DUS11 phosphatase [Machaerirhynchus nigripectus] | Machaerirhynchus nigripectus | 188 | 188 | 69% | 3.00E-56 | 53.8 | 169 | NWV88607.1 |
| RNA/RNP complex-1-interacting phosphatase isoform X4 [Anser cygnoides] | Anser cygnoides | 189 | 189 | 70% | 3.00E-56 | 52.3 | 197 | XP_047911010.1 |
| DUS11 phosphatase [Sylvietta virens] | Sylvietta virens | 188 | 188 | 69% | 4.00E-56 | 53.8 | 169 | NXK68968.1 |
| DUS11 phosphatase [Oreocharis arfaki] | Oreocharis arfaki | 188 | 188 | 69% | 4.00E-56 | 53.8 | 169 | NWW12342.1 |
| DUS11 phosphatase [Ifrita kowaldi] | Ifrita kowaldi | 187 | 187 | 69% | 4.00E-56 | 53.8 | 169 | NWW56266.1 |
| DUS11 phosphatase [Corvus moneduloides] | Corvus moneduloides | 187 | 187 | 69% | 4.00E-56 | 53.8 | 169 | NXD59544.1 |
| RNA/RNP complex-1-interacting phosphatase isoform X4 [Cygnus olor] | Cygnus olor | 189 | 189 | 70% | 5.00E-56 | 52.3 | 197 | XP_040394049.1 |
| DUS11 phosphatase [Lanius ludovicianus] | Lanius ludovicianus | 187 | 187 | 69% | 5.00E-56 | 54.97 | 169 | NWT88744.1 |
| RNA/RNP complex-1-interacting phosphatase isoform X1 [Harpia harpyja] | Harpia harpyja | 193 | 193 | 70% | 5.00E-56 | 52.87 | 329 | XP_052661274.1 |
| RNA/RNP complex-1-interacting phosphatase isoform X1 [Aythya fuligula] | Aythya fuligula | 192 | 192 | 70% | 5.00E-56 | 52.87 | 312 | XP_032059943.1 |
| DUS11 phosphatase [Myiagra hebetior] | Myiagra hebetior | 187 | 187 | 69% | 6.00E-56 | 53.22 | 169 | NXH23758.1 |
| DUS11 phosphatase [Dryoscopus gambensis] | Dryoscopus gambensis | 187 | 187 | 69% | 6.00E-56 | 53.22 | 169 | NWI76823.1 |
| RNA/RNP complex-1-interacting phosphatase [Hemicordylus capensis] | Hemicordylus capensis | 194 | 194 | 74% | 6.00E-56 | 50.54 | 378 | XP_053125337.1 |
| RNA/RNP complex-1-interacting phosphatase isoform X2 [Myiozetetes cayanensis] | Myiozetetes cayanensis | 187 | 187 | 73% | 7.00E-56 | 48.62 | 182 | XP_050181692.1 |
| DUS11 phosphatase [Mohoua ochrocephala] | Mohoua ochrocephala | 187 | 187 | 69% | 7.00E-56 | 53.22 | 169 | NXA66700.1 |
| DUS11 phosphatase [Falcunculus frontatus] | Falcunculus frontatus | 187 | 187 | 69% | 7.00E-56 | 53.22 | 169 | NWW22861.1 |
| DUS11 phosphatase [Vidua macroura] | Vidua macroura | 187 | 187 | 69% | 7.00E-56 | 51.46 | 169 | NXQ02454.1 |
| RNA/RNP complex-1-interacting phosphatase [Aix galericulata] | Aix galericulata | 192 | 192 | 70% | 8.00E-56 | 52.3 | 312 | KAI6073596.1 |
| DUS11 phosphatase [Gymnorhina tibicen] | Gymnorhina tibicen | 187 | 187 | 69% | 8.00E-56 | 53.22 | 169 | NXM43457.1 |
| RNA/RNP complex-1-interacting phosphatase isoform X2 [Hyaena hyaena] | Hyaena hyaena | 192 | 192 | 72% | 9.00E-56 | 51.11 | 330 | XP_039096093.1 |
| RNA/RNP complex-1-interacting phosphatase [Orycteropus afer afer] | Orycteropus afer afer | 191 | 191 | 87% | 9.00E-56 | 47.44 | 289 | XP_007950860.1 |
| RNA/RNP complex-1-interacting phosphatase [Alligator mississippiensis] | Alligator mississippiensis | 191 | 191 | 69% | 1.00E-55 | 52.91 | 311 | XP_014467031.1 |
| DUS11 phosphatase [Rhynochetos jubatus] | Rhynochetos jubatus | 187 | 187 | 69% | 1.00E-55 | 52.63 | 169 | NWW93328.1 |

#### Table S7 cont. Blastp results using penguinpox virus 2 (PEPV2) viral DUSP11 as query (QRM15716.1)

| | | | | | | | | |
|---|---|---|---|---|---|---|---|---|
| RNA/RNP complex-1-interacting phosphatase isoform X2 [Anser cygnoides] | Anser cygnoides | 190 | 190 | 70% | 1.00E-55 | 52.3 | 269 | XP_047911008.1 |
| DUS11 phosphatase [Erpornis zantholeuca] | Erpornis zantholeuca | 187 | 187 | 69% | 1.00E-55 | 53.8 | 169 | NXS78487.1 |
| PREDICTED: RNA/RNP complex-1-interacting phosphatase [Gavialis gangeticus] | Gavialis gangeticus | 191 | 191 | 69% | 1.00E-55 | 52.33 | 312 | XP_019369928.1 |
| RNA/RNP complex-1-interacting phosphatase isoform X2 [Cygnus olor] | Cygnus olor | 190 | 190 | 70% | 1.00E-55 | 52.3 | 269 | XP_040394047.1 |
| DUS11 phosphatase [Crocuta crocuta] | Crocuta crocuta | 192 | 192 | 72% | 1.00E-55 | 51.11 | 331 | KAF0879646.1 |
| RNA/RNP complex-1-interacting phosphatase isoform X1 [Hyaena hyaena] | Hyaena hyaena | 192 | 192 | 72% | 1.00E-55 | 51.11 | 331 | XP_039096092.1 |
| DUS11 phosphatase [Oriolus oriolus] | Oriolus oriolus | 186 | 186 | 69% | 2.00E-55 | 53.22 | 169 | NXO07055.1 |
| RNA/RNP complex-1-interacting phosphatase [Alligator mississippiensis] | Alligator mississippiensis | 192 | 192 | 69% | 2.00E-55 | 52.91 | 333 | KYO17197.1 |
| RNA/RNP complex-1-interacting phosphatase [Canis lupus familiaris] | Canis lupus familiaris | 191 | 191 | 72% | 2.00E-55 | 52.22 | 332 | XP_005630616.1 |
| RNA/RNP complex-1-interacting phosphatase [Cyrtonyx montezumae] | Cyrtonyx montezumae | 191 | 191 | 84% | 2.00E-55 | 45.45 | 305 | XP_065601237.1 |
| RNA/RNP complex-1-interacting phosphatase [Meles meles] | Meles meles | 192 | 192 | 91% | 2.00E-55 | 45.13 | 337 | XP_045834652.1 |
| RNA/RNP complex-1-interacting phosphatase-like [Empidonax traillii] | Empidonax traillii | 187 | 187 | 72% | 2.00E-55 | 48.89 | 184 | XP_027766280.1 |
| RNA/RNP complex-1-interacting phosphatase isoform X1 [Phacochoerus africanus] | Phacochoerus africanus | 189 | 189 | 72% | 2.00E-55 | 51.96 | 267 | XP_047637716.1 |
| RNA/RNP complex-1-interacting phosphatase isoform X1 [Talpa occidentalis] | Talpa occidentalis | 191 | 191 | 75% | 2.00E-55 | 51.05 | 332 | XP_037383543.1 |
| RNA/RNP complex-1-interacting phosphatase isoform X1 [Cynocephalus volans] | Cynocephalus volans | 191 | 191 | 73% | 2.00E-55 | 51.1 | 333 | XP_062934399.1 |
| DUS11 phosphatase [Toxostoma redivivum] | Toxostoma redivivum | 186 | 186 | 69% | 2.00E-55 | 52.05 | 169 | NWS83687.1 |
| DUS11 phosphatase [Daphoenositta chrysoptera] | Daphoenositta chrysoptera | 186 | 186 | 69% | 2.00E-55 | 53.22 | 169 | NWV50499.1 |
| RNA/RNP complex-1-interacting phosphatase isoform X2 [Oryctolagus cuniculus] | Oryctolagus cuniculus | 191 | 191 | 73% | 2.00E-55 | 52.2 | 330 | XP_008252457.1 |
| RNA/RNP complex-1-interacting phosphatase [Nyctereutes procyonoides] | Nyctereutes procyonoides | 191 | 191 | 72% | 2.00E-55 | 52.22 | 332 | XP_055168628.1 |
| RNA/RNP complex-1-interacting phosphatase isoform X3 [Myiozetetes cayanensis] | Myiozetetes cayanensis | 186 | 186 | 71% | 2.00E-55 | 49.43 | 180 | XP_050181693.1 |
| DUS11 phosphatase [Pterocles burchelli] | Pterocles burchelli | 186 | 186 | 69% | 2.00E-55 | 52.05 | 169 | NWU72880.1 |
| DUS11 phosphatase [Edolisoma coerulescens] | Edolisoma coerulescens | 186 | 186 | 69% | 2.00E-55 | 53.8 | 169 | NXH85140.1 |
| RNA/RNP complex-1-interacting phosphatase isoform X1 [Anser cygnoides] | Anser cygnoides | 191 | 191 | 70% | 2.00E-55 | 52.3 | 312 | XP_047911007.1 |
| DUS11 phosphatase [Buphagus erythrorhynchus] | Buphagus erythrorhynchus | 186 | 186 | 69% | 3.00E-55 | 53.22 | 169 | NXU04070.1 |
| RNA/RNP complex-1-interacting phosphatase [Suricata suricatta] | Suricata suricatta | 191 | 191 | 85% | 3.00E-55 | 46.45 | 330 | XP_029793463.1 |
| DUS11 phosphatase [Chauna torquata] | Chauna torquata | 186 | 186 | 69% | 3.00E-55 | 52.63 | 169 | NXK49017.1 |
| DUS11 phosphatase [Ardeotis kori] | Ardeotis kori | 186 | 186 | 69% | 3.00E-55 | 51.46 | 169 | NXE29890.1 |
| RNA/RNP complex-1-interacting phosphatase isoform X1 [Oryctolagus cuniculus] | Oryctolagus cuniculus | 191 | 191 | 73% | 3.00E-55 | 52.2 | 331 | XP_008252459.1 |
| RNA/RNP complex-1-interacting phosphatase [Ornithorhynchus anatinus] | Ornithorhynchus anatinus | 193 | 193 | 70% | 3.00E-55 | 51.72 | 419 | XP_028921823.1 |

**Table S7 cont. Blastp results using penguinpox virus 2 (PEPV2) viral DUSP11 as query (QRM15716.1)**

| | | | | | | | | |
|---|---|---|---|---|---|---|---|---|
| DUS11 phosphatase [Sinosuthora webbiana] | Sinosuthora webbiana | 185 | 185 | 69% | 3.00E-55 | 53.8 | 169 | NWR01313.1 |
| RNA/RNP complex-1-interacting phosphatase isoform X1 [Cygnus olor] | Cygnus olor | 190 | 190 | 70% | 3.00E-55 | 52.3 | 312 | XP_040394046.1 |
| RNA/RNP complex-1-interacting phosphatase isoform X2 [Pipra filicauda] | Pipra filicauda | 186 | 186 | 72% | 3.00E-55 | 49.16 | 186 | XP_027592473.1 |
| DUS11 phosphatase [Notiomystis cincta] | Notiomystis cincta | 185 | 185 | 69% | 4.00E-55 | 52.63 | 169 | NWX26569.1 |
| RNA/RNP complex-1-interacting phosphatase isoform X1 [Lynx canadensis] | Lynx canadensis | 191 | 191 | 72% | 4.00E-55 | 51.11 | 331 | XP_032448558.1 |
| RNA/RNP complex-1-interacting phosphatase isoform X2 [Lynx canadensis] | Lynx canadensis | 191 | 191 | 72% | 4.00E-55 | 51.11 | 332 | XP_030166722.1 |
| RNA/RNP complex-1-interacting phosphatase [Cygnus atratus] | Cygnus atratus | 190 | 190 | 70% | 4.00E-55 | 52.3 | 312 | XP_050571750.1 |
| RNA/RNP complex-1-interacting phosphatase [Heterocephalus glaber] | Heterocephalus glaber | 186 | 186 | 72% | 4.00E-55 | 51.4 | 206 | XP_004844776.1 |
| DUS11 phosphatase [Galbula dea] | Galbula dea | 185 | 185 | 69% | 4.00E-55 | 52.05 | 169 | NXI47672.1 |

#### Table S8. Blastp results using vDUSP11 for mudlarkpox virus as query (QRM15361.1)

| Description | Scientific Name | Max Score | Total Score | Query Cover | E value | Per. ident | Acc. Len | Accession |
|---|---|---|---|---|---|---|---|---|
| putative RNA phosphatase [Mudlarkpox virus] | Mudlarkpox virus | 802 | 802 | 100% | 0 | 100 | 403 | QRM15361.1 |
| CNPV085 putative RNA phosphatase [Canarypox virus] | Canarypox virus | 699 | 699 | 100% | 0 | 86.9 | 403 | NP_955108.1 |
| DSP DUSP11 [Finch poxvirus] | Finch poxvirus | 603 | 861 | 100% | 0 | 75.2 | 501 | UOX38764.1 |
| putative RNA phosphatase [Cheloniid poxvirus 1] | Cheloniid poxvirus 1 | 593 | 882 | 97% | 0 | 95.7 | 303 | QRI42800.1 |
| SWPV2-ORF080 [Shearwaterpox virus] | Shearwaterpox virus | 530 | 820 | 97% | 0 | 85.7 | 303 | ARE67303.1 |
| putative RNA phosphatase [Magpiepox virus] | Magpiepox virus | 498 | 934 | 96% | 5.00E-174 | 86.1 | 286 | QGM48715.1 |
| putative RNA phosphatase [Penguinpox virus 2] | Penguinpox virus 2 | 406 | 460 | 71% | 2.00E-138 | 81.6 | 247 | QRM15716.1 |
| putative RNA phosphatase [Albatrosspox virus] | Albatrosspox virus | 404 | 456 | 77% | 1.00E-137 | 78.5 | 251 | QRM16049.1 |
| SWPV1-074 [Shearwaterpox virus] | Shearwaterpox virus | 285 | 285 | 47% | 9.00E-91 | 69 | 245 | ARF02681.1 |
| RNA/RNP complex-1-interacting phosphatase isoform X3 [Varanus komodoensis] | Varanus komodoensis | 189 | 189 | 46% | 1.00E-53 | 50.79 | 249 | XP_044273321.1 |
| RNA/RNP complex-1-interacting phosphatase isoform X2 [Varanus komodoensis] | Varanus komodoensis | 189 | 189 | 46% | 2.00E-53 | 50.79 | 261 | XP_044273320.1 |
| RNA/RNP complex-1-interacting phosphatase isoform X2 [Eublepharis macularius] | Eublepharis macularius | 192 | 192 | 51% | 2.00E-53 | 47.85 | 357 | XP_054854289.1 |
| RNA/RNP complex-1-interacting phosphatase isoform X1 [Eublepharis macularius] | Eublepharis macularius | 192 | 192 | 51% | 2.00E-53 | 47.85 | 358 | XP_054854288.1 |
| RNA/RNP complex-1-interacting phosphatase [Sceloporus undulatus] | Sceloporus undulatus | 189 | 189 | 47% | 8.00E-53 | 49.49 | 321 | XP_042328494.1 |
| RNA/RNP complex-1-interacting phosphatase isoform X3 [Tiliqua scincoides] | Tiliqua scincoides | 189 | 189 | 43% | 1.00E-52 | 51.15 | 307 | XP_066495500.1 |
| RNA/RNP complex-1-interacting phosphatase isoform X1 [Tiliqua scincoides] | Tiliqua scincoides | 189 | 189 | 43% | 2.00E-52 | 51.15 | 320 | XP_066495498.1 |
| RNA/RNP complex-1-interacting phosphatase isoform X2 [Tiliqua scincoides] | Tiliqua scincoides | 187 | 187 | 43% | 4.00E-52 | 51.15 | 312 | XP_066495499.1 |
| RNA/RNP complex-1-interacting phosphatase isoform X1 [Varanus komodoensis] | Varanus komodoensis | 188 | 188 | 46% | 4.00E-52 | 50.79 | 335 | XP_044273319.1 |
| RNA/RNP complex-1-interacting phosphatase [Orycteropus afer afer] | Orycteropus afer afer | 186 | 186 | 53% | 7.00E-52 | 46.98 | 289 | XP_007950860.1 |
| RNA/RNP complex-1-interacting phosphatase isoform X1 [Anolis carolinensis] | Anolis carolinensis | 189 | 189 | 45% | 8.00E-52 | 49.19 | 381 | XP_062817178.1 |
| RNA/RNP complex-1-interacting phosphatase isoform X2 [Anolis carolinensis] | Anolis carolinensis | 189 | 189 | 45% | 9.00E-52 | 49.19 | 376 | XP_062817179.1 |

**Table S8 cont. Blastp results using vDUSP11 for mudlarkpox virus as query (QRM15361.1)**

| | | | | | | | | |
|---|---|---|---|---|---|---|---|---|
| RNA/RNP complex-1-interacting phosphatase [Morone saxatilis] | Morone saxatilis | 183 | 183 | 42% | 1.00E-51 | 48.54 | 220 | XP_035524897.1 |
| RNA/RNP complex-1-interacting phosphatase isoform X2 [Elgaria multicarinata webbii] | Elgaria multicarinata webbii | 184 | 184 | 45% | 2.00E-51 | 50 | 259 | XP_062995277.1 |
| RNA/RNP complex-1-interacting phosphatase [Calypte anna] | Calypte anna | 186 | 186 | 43% | 2.00E-51 | 51.15 | 320 | XP_030319948.1 |
| RNA/RNP complex-1-interacting phosphatase isoform X1 [Cynocephalus volans] | Cynocephalus volans | 186 | 186 | 44% | 4.00E-51 | 50.83 | 333 | XP_062934399.1 |
| RNA/RNP complex-1-interacting phosphatase isoform X2 [Hyaena hyaena] | Hyaena hyaena | 185 | 185 | 44% | 4.00E-51 | 50.28 | 330 | XP_039096093.1 |
| RNA/RNP complex-1-interacting phosphatase [Grus americana] | Grus americana | 184 | 184 | 44% | 5.00E-51 | 48.89 | 309 | XP_054660434.1 |
| DUS11 phosphatase [Crocuta crocuta] | Crocuta crocuta | 185 | 185 | 44% | 6.00E-51 | 50.28 | 331 | KAF0879646.1 |
| RNA/RNP complex-1-interacting phosphatase isoform X1 [Hyaena hyaena] | Hyaena hyaena | 185 | 185 | 44% | 6.00E-51 | 50.28 | 331 | XP_039096092.1 |
| RNA/RNP complex-1-interacting phosphatase isoform X5 [Molossus molossus] | Molossus molossus | 183 | 183 | 44% | 7.00E-51 | 50.28 | 270 | XP_036119097.1 |
| RNA/RNP complex-1-interacting phosphatase [Heterocephalus glaber] | Heterocephalus glaber | 181 | 181 | 44% | 7.00E-51 | 50.84 | 206 | XP_004844776.1 |
| RNA/RNP complex-1-interacting phosphatase [Suricata suricatta] | Suricata suricatta | 185 | 185 | 52% | 7.00E-51 | 45.5 | 330 | XP_029793463.1 |
| RNA/RNP complex-1-interacting phosphatase isoform X4 [Molossus molossus] | Molossus molossus | 183 | 183 | 44% | 8.00E-51 | 50.28 | 274 | XP_036119096.1 |
| RNA/RNP complex-1-interacting phosphatase [Anolis sagrei ordinatus] | Anolis sagrei ordinatus | 185 | 185 | 45% | 1.00E-50 | 49.45 | 341 | XP_060643263.1 |
| DUS11 phosphatase [Geococcyx californianus] | Geococcyx californianus | 179 | 179 | 42% | 2.00E-50 | 52.05 | 169 | NWH58149.1 |
| RNA/RNP complex-1-interacting phosphatase isoform X2 [Cynocephalus volans] | Cynocephalus volans | 184 | 184 | 44% | 2.00E-50 | 50.83 | 334 | XP_062934400.1 |
| RNA/RNP complex-1-interacting phosphatase [Rhinatrema bivittatum] | Rhinatrema bivittatum | 184 | 184 | 45% | 2.00E-50 | 50.27 | 344 | XP_029460003.1 |
| RNA/RNP complex-1-interacting phosphatase [Rhineura floridana] | Rhineura floridana | 184 | 184 | 45% | 2.00E-50 | 50.55 | 325 | XP_061448984.1 |
| RNA/RNP complex-1-interacting phosphatase isoform X1 [Lynx canadensis] | Lynx canadensis | 184 | 184 | 44% | 2.00E-50 | 50.28 | 331 | XP_032448558.1 |
| RNA/RNP complex-1-interacting phosphatase isoform X2 [Lynx canadensis] | Lynx canadensis | 184 | 184 | 44% | 2.00E-50 | 50.28 | 332 | XP_030166722.1 |

#### Table S8 cont. Blastp results using vDUSP11 for mudlarkpox virus as query (QRM15361.1)

| | | | | | | | | |
|---|---|---|---|---|---|---|---|---|
| RNA/RNP complex-1-interacting phosphatase isoform X2 [Cricetulus griseus] | Cricetulus griseus | 184 | 184 | 46% | 2.00E-50 | 49.73 | 330 | XP_007648875.1 |
| RNA/RNP complex-1-interacting phosphatase isoform X3 [Erinaceus europaeus] | Erinaceus europaeus | 184 | 184 | 51% | 2.00E-50 | 44.98 | 333 | XP_060043066.1 |
| RNA/RNP complex-1-interacting phosphatase isoform X2 [Molossus molossus] | Molossus molossus | 183 | 183 | 44% | 2.00E-50 | 50.28 | 304 | XP_036119094.1 |
| RNA/RNP complex-1-interacting phosphatase [Canis lupus familiaris] | Canis lupus familiaris | 184 | 184 | 44% | 2.00E-50 | 50.84 | 332 | XP_005630616.1 |
| RNA/RNP complex-1-interacting phosphatase isoform X2 [Leopardus geoffroyi] | Leopardus geoffroyi | 184 | 184 | 44% | 2.00E-50 | 50.28 | 331 | XP_045302449.1 |
| RNA/RNP complex-1-interacting phosphatase [Nyctereutes procyonoides] | Nyctereutes procyonoides | 183 | 183 | 44% | 3.00E-50 | 50.84 | 332 | XP_055168628.1 |
| RNA/RNP complex-1-interacting phosphatase isoform X1 [Leopardus geoffroyi] | Leopardus geoffroyi | 183 | 183 | 44% | 3.00E-50 | 50.28 | 332 | XP_045302448.1 |
| RNA/RNP complex-1-interacting phosphatase [Peromyscus leucopus] | Peromyscus leucopus | 183 | 183 | 45% | 3.00E-50 | 50.54 | 329 | XP_028718951.1 |
| RNA/RNP complex-1-interacting phosphatase isoform X1 [Puma concolor] | Puma concolor | 183 | 183 | 44% | 3.00E-50 | 50.28 | 331 | XP_025787671.1 |
| hypothetical protein JRQ81_011232 [Phrynocephalus forsythii] | Phrynocephalus forsythii | 183 | 183 | 45% | 3.00E-50 | 49.19 | 328 | KAJ7305313.1 |
| RNA/RNP complex-1-interacting phosphatase isoform X1 [Felis catus] | Felis catus | 183 | 183 | 44% | 3.00E-50 | 50.28 | 332 | XP_006930253.1 |
| PREDICTED: RNA/RNP complex-1-interacting phosphatase [Rhinolophus sinicus] | Rhinolophus sinicus | 183 | 183 | 44% | 3.00E-50 | 50.56 | 330 | XP_019568432.1 |
| RNA/RNP complex-1-interacting phosphatase isoform X1 [Elgaria multicarinata webbii] | Elgaria multicarinata webbii | 183 | 183 | 45% | 4.00E-50 | 50 | 323 | XP_062995276.1 |
| PREDICTED: RNA/RNP complex-1-interacting phosphatase isoform X6 [Galeopterus variegatus] | Galeopterus variegatus | 183 | 183 | 44% | 4.00E-50 | 50.28 | 334 | XP_008569404.1 |
| RNA/RNP complex-1-interacting phosphatase isoform X2 [Neofelis nebulosa] | Neofelis nebulosa | 183 | 183 | 44% | 4.00E-50 | 50.28 | 331 | XP_058539656.1 |
| RNA/RNP complex-1-interacting phosphatase [Callorhinus ursinus] | Callorhinus ursinus | 183 | 183 | 44% | 4.00E-50 | 49.72 | 332 | XP_025724176.1 |
| RNA/RNP complex-1-interacting phosphatase [Hemicordylus capensis] | Hemicordylus capensis | 184 | 184 | 45% | 4.00E-50 | 50.55 | 378 | XP_053125337.1 |
| RNA/RNP complex-1-interacting phosphatase isoform X1 [Neofelis nebulosa] | Neofelis nebulosa | 183 | 183 | 44% | 4.00E-50 | 50.28 | 332 | XP_058539655.1 |

**Table S8 cont. Blastp results using vDUSP11 for mudlarkpox virus as query (QRM15361.1)**

| | | | | | | | | |
|---|---|---|---|---|---|---|---|---|
| RNA/RNP complex-1-interacting phosphatase [Meles meles] | Meles meles | 183 | 183 | 52% | 4.00E-50 | 44.6 | 337 | XP_045834652.1 |
| RNA/RNP complex-1-interacting phosphatase isoform X3 [Balaenoptera acutorostrata] | Balaenoptera acutorostrata | 181 | 181 | 57% | 5.00E-50 | 42.67 | 267 | XP_057414036.1 |
| RNA/RNP complex-1-interacting phosphatase [Peromyscus californicus insignis] | Peromyscus californicus insignis | 183 | 183 | 45% | 5.00E-50 | 50.54 | 330 | XP_052575682.1 |
| hypothetical protein lerEdw1_009417 [Lerista edwardsae] | Lerista edwardsae | 183 | 183 | 46% | 5.00E-50 | 48.69 | 346 | KAJ6653253.1 |
| RNA/RNP complex-1-interacting phosphatase isoform X2 [Oryctolagus cuniculus] | Oryctolagus cuniculus | 182 | 182 | 45% | 5.00E-50 | 50.55 | 330 | XP_008252457.1 |
| PREDICTED: RNA/RNP complex-1-interacting phosphatase [Condylura cristata] | Condylura cristata | 182 | 182 | 44% | 5.00E-50 | 50.83 | 333 | XP_004691589.1 |
| RNA/RNP complex-1-interacting phosphatase [Ursus americanus] | Ursus americanus | 182 | 182 | 44% | 5.00E-50 | 49.72 | 332 | XP_045644246.1 |
| RNA/RNP complex-1-interacting phosphatase isoform X1 [Talpa occidentalis] | Talpa occidentalis | 182 | 182 | 44% | 6.00E-50 | 50.28 | 332 | XP_037383543.1 |
| RNA/RNP complex-1-interacting phosphatase isoform X2 [Peromyscus maniculatus bairdii] | Peromyscus maniculatus bairdii | 182 | 182 | 45% | 6.00E-50 | 50 | 329 | XP_006998014.1 |
| RNA/RNP complex-1-interacting phosphatase isoform X1 [Oryctolagus cuniculus] | Oryctolagus cuniculus | 182 | 182 | 45% | 6.00E-50 | 50.55 | 331 | XP_008252459.1 |
| RNA/RNP complex-1-interacting phosphatase isoform X1 [Phacochoerus africanus] | Phacochoerus africanus | 180 | 180 | 44% | 6.00E-50 | 50.84 | 267 | XP_047637716.1 |
| RNA/RNP complex-1-interacting phosphatase [Galemys pyrenaicus] | Galemys pyrenaicus | 181 | 181 | 44% | 7.00E-50 | 50.28 | 275 | KAG8518745.1 |
| hypothetical protein E2320_006809 [Naja naja] | Naja naja | 182 | 182 | 45% | 7.00E-50 | 49.45 | 346 | KAG8141143.1 |
| RNA/RNP complex-1-interacting phosphatase isoform X2 [Heteronotia binoei] | Heteronotia binoei | 182 | 182 | 45% | 8.00E-50 | 50 | 312 | XP_060106818.1 |
| RNA/RNP complex-1-interacting phosphatase isoform X2 [Phacochoerus africanus] | Phacochoerus africanus | 182 | 182 | 44% | 8.00E-50 | 50.84 | 327 | XP_047637717.1 |
| RNA/RNP complex-1-interacting phosphatase [Sturnira hondurensis] | Sturnira hondurensis | 182 | 182 | 50% | 8.00E-50 | 46.57 | 331 | XP_036904110.1 |
| RNA/RNP complex-1-interacting phosphatase [Dasypus novemcinctus] | Dasypus novemcinctus | 182 | 182 | 45% | 8.00E-50 | 51.1 | 326 | XP_004482114.3 |
| RNA/RNP complex-1-interacting phosphatase isoform X2 [Neomonachus schauinslandi] | Neomonachus schauinslandi | 180 | 180 | 44% | 900E-50 | 49.72 | 270 | XP_021554772.1 |

**Table S8 cont. Blastp results using vDUSP11 for mudlarkpox virus as query (QRM15361.1)**

| | | | | | | | | |
|---|---|---|---|---|---|---|---|---|
| RNA/RNP complex-1-interacting phosphatase isoform X2 [Nycticebus coucang] | Nycticebus coucang | 182 | 182 | 44% | 9.00E-50 | 51.4 | 327 | XP_053444277.1 |
| RNA/RNP complex-1-interacting phosphatase [Ailuropoda melanoleuca] | Ailuropoda melanoleuca | 182 | 182 | 45% | 9.00E-50 | 50 | 332 | XP_002921547.2 |
| RNA/RNP complex-1-interacting phosphatase isoform X4 [Erinaceus europaeus] | Erinaceus europaeus | 182 | 182 | 51% | 1.00E-49 | 44.98 | 330 | XP_007519010.1 |
| RNA/RNP complex-1-interacting phosphatase isoform X1 [Nycticebus coucang] | Nycticebus coucang | 182 | 182 | 44% | 1.00E-49 | 51.4 | 332 | XP_053444276.1 |
| RNA/RNP complex-1-interacting phosphatase [Mastomys coucha] | Mastomys coucha | 182 | 182 | 45% | 1.00E-49 | 50 | 322 | XP_031238264.1 |
| RNA/RNP complex-1-interacting phosphatase isoform X1 [Castor canadensis] | Castor canadensis | 182 | 182 | 44% | 1.00E-49 | 51.12 | 332 | XP_020040530.1 |
| RNA/RNP complex-1-interacting phosphatase [Ursus maritimus] | Ursus maritimus | 183 | 183 | 44% | 1.00E-49 | 49.72 | 379 | XP_008698222.2 |
| RNA/RNP complex-1-interacting phosphatase isoform X1 [Heteronotia binoei] | Heteronotia binoei | 182 | 182 | 45% | 1.00E-49 | 50 | 327 | XP_060106817.1 |
| RNA/RNP complex-1-interacting phosphatase [Zalophus californianus] | Zalophus californianus | 182 | 182 | 44% | 1.00E-49 | 49.72 | 332 | XP_027477093.1 |
| RNA/RNP complex-1-interacting phosphatase [Eumetopias jubatus] | Eumetopias jubatus | 182 | 182 | 44% | 1.00E-49 | 49.72 | 332 | XP_027971960.1 |
| RNA/RNP complex-1-interacting phosphatase isoform X2 [Castor canadensis] | Castor canadensis | 181 | 181 | 44% | 1.00E-49 | 51.12 | 324 | XP_020040531.1 |
| RNA/RNP complex-1-interacting phosphatase-like [Callorhinus ursinus] | Callorhinus ursinus | 181 | 181 | 54% | 1.00E-49 | 44.5 | 298 | XP_025743387.1 |
| RNA/RNP complex-1-interacting phosphatase isoform X1 [Eptesicus fuscus] | Eptesicus fuscus | 182 | 182 | 44% | 1.00E-49 | 50.84 | 333 | XP_008145661.1 |
| RNA/RNP complex-1-interacting phosphatase [Artibeus jamaicensis] | Artibeus jamaicensis | 181 | 181 | 50% | 1.00E-49 | 46.08 | 331 | XP_036986788.2 |
| RNA/RNP complex-1-interacting phosphatase isoform X1 [Molossus molossus] | Molossus molossus | 182 | 182 | 44% | 1.00E-49 | 50.28 | 334 | XP_036119088.1 |
| PREDICTED: RNA/RNP complex-1-interacting phosphatase [Odobenus rosmarus divergens] | Odobenus rosmarus divergens | 181 | 181 | 44% | 1.00E-49 | 49.72 | 332 | XP_004398756.1 |
| RNA/RNP complex-1-interacting phosphatase isoform X1 [Pipistrellus kuhlii] | Pipistrellus kuhlii | 182 | 182 | 45% | 2.00E-49 | 51.1 | 337 | XP_036291622.1 |
| unnamed protein product [Pipistrellus nathusii] | Pipistrellus nathusii | 181 | 181 | 45% | 2.00E-49 | 51.1 | 337 | CAK6435205.1 |
| RNA/RNP complex-1-interacting phosphatase-like [Empidonax traillii] | Empidonax traillii | 176 | 176 | 44% | 2.00E-49 | 48.33 | 184 | XP_027766280.1 |

**Table S8 cont. Blastp results using vDUSP11 for mudlarkpox virus as query (QRM15361.1)**

| | | | | | | | | |
|---|---|---|---|---|---|---|---|---|
| RNA/RNP complex-1-interacting phosphatase [Lutra lutra] | Lutra lutra | 181 | 181 | 44% | 2.00E-49 | 49.72 | 339 | XP_047601880.1 |
| RNA/RNP complex-1-interacting phosphatase [Rhinolophus ferrumequinum] | Rhinolophus ferrumequinum | 181 | 181 | 44% | 2.00E-49 | 50 | 330 | XP_032981156.1 |
| RNA/RNP complex-1-interacting phosphatase [Monodelphis domestica] | Monodelphis domestica | 182 | 182 | 42% | 2.00E-49 | 49.71 | 376 | XP_007477939.1 |
| RNA/RNP complex-1-interacting phosphatase isoform X4 [Anser cygnoides] | Anser cygnoides | 177 | 177 | 43% | 2.00E-49 | 50 | 197 | XP_047911010.1 |

**Table S9. Reverse Blastp results using remaining vDUSP11 as query. vDUSP11 used as query is listed first in each table. First 10 results shown for each virus excluding other avipox vDUSP11s.**

| Description | Scientific Name | Max Score | Total Score | Query Cover | E value | Per. ident | Acc. Len | Accession |
|---|---|---|---|---|---|---|---|---|
| *SWPV1-074 [Shearwaterpox virus]* | *Shearwaterpox virus* | *498* | *498* | *100%* | *1.00E-177* | *100* | *245* | *ARF02681.1* |
| RNA/RNP complex-1-interacting phosphatase [Grus americana] | Grus americana | 183 | 183 | 93% | 2.00E-52 | 41.99 | 309 | XP_054660434.1 |
| **RNA/RNP complex-1-interacting phosphatase [Corvus hawaiiensis]** | **Corvus hawaiiensis** | **181** | **181** | **72%** | **6.00E-52** | **48.88** | **305** | **XP_048146001.1** |
| DUS11 phosphatase [Pachycephala philippinensis] | Pachycephala philippinensis | 176 | 176 | 68% | 1.00E-51 | 50.89 | 169 | NXI05060.1 |
| RNA/RNP complex-1-interacting phosphatase [Jaculus jaculus] | Jaculus jaculus | 181 | 181 | 71% | 2.00E-51 | 49.14 | 329 | XP_004668336.1 |
| DUS11 phosphatase [Corvus moneduloides] | Corvus moneduloides | 176 | 176 | 68% | 2.00E-51 | 50.89 | 169 | NXD59544.1 |
| DUS11 phosphatase [Eulacestoma nigropectus] | Eulacestoma nigropectus | 176 | 176 | 68% | 2.00E-51 | 50.3 | 169 | NXB30747.1 |
| DUS11 phosphatase [Grus americana] | Grus americana | 175 | 175 | 68% | 4.00E-51 | 50.3 | 169 | NWH25829.1 |
| RNA/RNP complex-1-interacting phosphatase [Corvus cornix cornix] | Corvus cornix cornix | 181 | 181 | 72% | 5.00E-51 | 48.88 | 349 | XP_039420096.1 |
| DUS11 phosphatase [Dicrurus megarhynchus] | Dicrurus megarhynchus | 174 | 174 | 68% | 6.00E-51 | 50.3 | 169 | NXJ20454.1 |
| RNA/RNP complex-1-interacting phosphatase [Orycteropus afer afer] | Orycteropus afer afer | 178 | 178 | 71% | 6.00E-51 | 48.28 | 289 | XP_007950860.1 |
| DUS11 phosphatase [Machaerirhynchus nigripectus] | Machaerirhynchus nigripectus | 174 | 174 | 68% | 6.00E-51 | 50.89 | 169 | NWV88607.1 |

| Description | Scientific Name | Max Score | Total Score | Query Cover | E value | Per. ident | Acc. Len | Accession |
|---|---|---|---|---|---|---|---|---|
| *putative RNA phosphatase [Cheloniid poxvirus 1]* | *Cheloniid poxvirus 1* | *615* | *615* | *100%* | *0* | *100* | *303* | *QRI42800.1* |
| RNA/RNP complex-1-interacting phosphatase [Sceloporus undulatus] | Sceloporus undulatus | 191 | 191 | 63% | 2.00E-54 | 49.49 | 321 | XP_042328494.1 |
| RNA/RNP complex-1-interacting phosphatase isoform X3 [Varanus komodoensis] | Varanus komodoensis | 187 | 187 | 57% | 3.00E-54 | 51.69 | 249 | XP_044273321.1 |
| RNA/RNP complex-1-interacting phosphatase isoform X3 [Tiliqua scincoides] | Tiliqua scincoides | 189 | 189 | 57% | 6.00E-54 | 50.86 | 307 | XP_066495500.1 |
| RNA/RNP complex-1-interacting phosphatase isoform X2 [Varanus komodoensis] | Varanus komodoensis | 187 | 187 | 57% | 6.00E-54 | 51.69 | 261 | XP_044273320.1 |
| RNA/RNP complex-1-interacting phosphatase isoform X1 [Anolis carolinensis] | Anolis carolinensis | 191 | 191 | 61% | 8.00E-54 | 49.2 | 381 | XP_062817178.1 |
| RNA/RNP complex-1-interacting phosphatase isoform X2 [Anolis carolinensis] | Anolis carolinensis | 190 | 190 | 61% | 1.00E-53 | 49.2 | 376 | XP_062817179.1 |
| RNA/RNP complex-1-interacting phosphatase isoform X1 [Tiliqua scincoides] | Tiliqua scincoides | 189 | 189 | 57% | 1.00E-53 | 50.86 | 320 | XP_066495498.1 |
| RNA/RNP complex-1-interacting phosphatase isoform X2 [Tiliqua scincoides] | Tiliqua scincoides | 188 | 188 | 57% | 2.00E-53 | 50.86 | 312 | XP_066495499.1 |
| RNA/RNP complex-1-interacting phosphatase isoform X2 [Elgaria multicarinata webbii] | Elgaria multicarinata webbii | 186 | 186 | 62% | 3.00E-53 | 49.21 | 259 | XP_062995277.1 |
| RNA/RNP complex-1-interacting phosphatase [Grus americana] | Grus americana | 187 | 187 | 59% | 3.00E-53 | 50.84 | 309 | XP_054660434.1 |
| **RNA/RNP complex-1-interacting phosphatase [Corvus hawaiiensis]** | **Corvus hawaiiensis** | **187** | **187** | **59%** | **3.00E-53** | **49.17** | **305** | **XP_048146001.1** |

**Table S9 cont. Reverse Blastp results using remaining vDUSP11 as query. vDUSP11 used as query is listed first in each table. First 10 results shown for each virus excluding other avipox vDUSP11s.**

| Description | Scientific Name | Max Score | Total Score | Query Cover | E value | Per. ident | Acc. Len | Accession |
|---|---|---|---|---|---|---|---|---|
| *CNPV085 putative RNA phosphatase [Canarypox virus]* | *Canarypox virus* | *803* | *803* | *100%* | *0* | *100* | *403* | *NP_955108.1* |
| RNA/RNP complex-1-interacting phosphatase [Corvus hawaiiensis] | Corvus hawaiiensis | 194 | 194 | 43% | 8.00E-55 | 51.7 | 305 | XP_048146001.1 |
| RNA/RNP complex-1-interacting phosphatase-like isoform X1 [Motacilla alba alba] | Motacilla alba alba | 191 | 191 | 51% | 1.00E-54 | 46.41 | 222 | XP_038016256.1 |
| RNA/RNP complex-1-interacting phosphatase [Calypte anna] | Calypte anna | 194 | 194 | 43% | 1.00E-54 | 52 | 320 | XP_030319948.1 |
| RNA/RNP complex-1-interacting phosphatase [Grus americana] | Grus americana | 194 | 194 | 43% | 1.00E-54 | 51.14 | 309 | XP_054660434.1 |
| RNA/RNP complex-1-interacting phosphatase [Sceloporus undulatus] | Sceloporus undulatus | 194 | 194 | 46% | 2.00E-54 | 51.05 | 321 | XP_042328494.1 |
| RNA/RNP complex-1-interacting phosphatase [Corvus cornix cornix] | Corvus cornix cornix | 194 | 194 | 43% | 4.00E-54 | 51.7 | 349 | XP_039420096.1 |
| RNA/RNP complex-1-interacting phosphatase-like isoform X2 [Motacilla alba alba] | Motacilla alba alba | 190 | 190 | 51% | 4.00E-54 | 45.67 | 218 | XP_038016258.1 |
| DUS11 phosphatase [Vidua macroura] | Vidua macroura | 188 | 188 | 42% | 4.00E-54 | 51.46 | 169 | NXQ02454.1 |
| PREDICTED: RNA/RNP complex-1-interacting phosphatase [Sturnus vulgaris] | Sturnus vulgaris | 196 | 196 | 43% | 5.00E-54 | 52.57 | 435 | XP_014747572.1 |
| DUS11 phosphatase [Pachycephala philippinensis] | Pachycephala philippinensis | 187 | 187 | 42% | 8.00E-54 | 53.22 | 169 | NXI05060.1 |
| DUS11 phosphatase [Struthidea cinerea] | Struthidea cinerea | 187 | 187 | 42% | 9.00E-54 | 53.8 | 169 | NXB60100.1 |

| Description | Scientific Name | Max Score | Total Score | Query Cover | E value | Per. ident | Acc. Len | Accession |
|---|---|---|---|---|---|---|---|---|
| ***DSP DUSP11 [Finch poxvirus]*** | ***Finch poxvirus*** | ***962*** | ***962*** | ***100%*** | ***0*** | ***100*** | ***501*** | ***UOX38764.1*** |
| RNA/RNP complex-1-interacting phosphatase [Grus americana] | Grus americana | 198 | 198 | 37% | 3.00E-55 | 50 | 309 | XP_054660434.1 |
| RNA/RNP complex-1-interacting phosphatase [Melopsittacus undulatus] | Melopsittacus undulatus | 196 | 196 | 37% | 2.00E-54 | 48.95 | 314 | XP_033926501.1 |
| RNA/RNP complex-1-interacting phosphatase [Calypte anna] | Calypte anna | 193 | 193 | 34% | 4.00E-53 | 50.57 | 320 | XP_030319948.1 |
| RNA/RNP complex-1-interacting phosphatase [Indicator indicator] | Indicator indicator | 192 | 192 | 34% | 1.00E-52 | 50.57 | 325 | XP_054252571.1 |
| RNA/RNP complex-1-interacting phosphatase [Canis lupus familiaris] | Canis lupus familiaris | 191 | 191 | 39% | 2.00E-52 | 49.25 | 332 | XP_005630616.1 |
| RNA/RNP complex-1-interacting phosphatase [Nyctereutes procyonoides] | Nyctereutes procyonoides | 191 | 191 | 39% | 3.00E-52 | 49.25 | 332 | XP_055168628.1 |
| **RNA/RNP complex-1-interacting phosphatase [Corvus hawaiiensis]** | **Corvus hawaiiensis** | **190** | **190** | **34%** | **3.00E-52** | **50.29** | **305** | **XP_048146001.1** |
| RNA/RNP complex-1-interacting phosphatase isoform X1 [Myiozetetes cayanensis] | Myiozetetes cayanensis | 190 | 190 | 34% | 4.00E-52 | 50 | 306 | XP_050181691.1 |
| RNA/RNP complex-1-interacting phosphatase [Ornithorhynchus anatinus] | Ornithorhynchus anatinus | 193 | 193 | 40% | 4.00E-52 | 46.77 | 419 | XP_028921823.1 |
| RNA/RNP complex-1-interacting phosphatase [Rhinatrema bivittatum] | Rhinatrema bivittatum | 191 | 191 | 36% | 4.00E-52 | 49.73 | 344 | XP_029460003.1 |
| RNA/RNP complex-1-interacting phosphatase [Pogoniulus pusillus] | Pogoniulus pusillus | 190 | 190 | 34% | 5.00E-52 | 50.29 | 308 | XP_064031825.1 |

**Table S9 cont. Reverse Blastp results using remaining vDUSP11 as query. vDUSP11 used as query is listed first in each table. First 10 results shown for each virus excluding other avipox vDUSP11s.**

| Description | Scientific Name | Max Score | Total Score | Query Cover | E value | Per. ident | Acc. Len | Accession |
|---|---|---|---|---|---|---|---|---|
| *putative RNA phosphatase [Magpiepox virus]* | *Magpiepox virus* | *585* | *585* | *100%* | *0* | *100* | *286* | *QGM48715.1* |
| **RNA/RNP complex-1-interacting phosphatase [Corvus hawaiiensis]** | **Corvus hawaiiensis** | **197** | **197** | **61%** | **1.00E-57** | **52.84** | **305** | **XP_048146001.1** |
| RNA/RNP complex-1-interacting phosphatase [Grus americana] | Grus americana | 196 | 196 | 61% | 4.00E-57 | 52.27 | 309 | XP_054660434.1 |
| RNA/RNP complex-1-interacting phosphatase [Calypte anna] | Calypte anna | 196 | 196 | 60% | 6.00E-57 | 53.45 | 320 | XP_030319948.1 |
| RNA/RNP complex-1-interacting phosphatase [Corvus cornix cornix] | Corvus cornix cornix | 197 | 197 | 61% | 8.00E-57 | 52.84 | 349 | XP_039420096.1 |
| RNA/RNP complex-1-interacting phosphatase isoform X3 [Accipiter gentilis] | Accipiter gentilis | 194 | 194 | 60% | 9.00E-57 | 53.45 | 271 | XP_049686642.1 |
| RNA/RNP complex-1-interacting phosphatase isoform X2 [Aquila chrysaetos chrysaetos] | Aquila chrysaetos chrysaetos | 196 | 196 | 60% | 1.00E-56 | 53.45 | 314 | XP_029890719.1 |
| DUS11 phosphatase [Pachycephala philippinensis] | Pachycephala philippinensis | 191 | 191 | 59% | 1.00E-56 | 54.39 | 169 | NXI05060.1 |
| RNA/RNP complex-1-interacting phosphatase isoform X3 [Aquila chrysaetos chrysaetos] | Aquila chrysaetos chrysaetos | 194 | 194 | 60% | 1.00E-56 | 53.45 | 271 | XP_040984244.1 |
| DUS11 phosphatase [Struthidea cinerea] | Struthidea cinerea | 190 | 190 | 59% | 1.00E-56 | 54.97 | 169 | NXB60100.1 |
| PREDICTED: RNA/RNP complex-1-interacting phosphatase isoform X3 [Haliaeetus leucocephalus] | Haliaeetus leucocephalus | 196 | 196 | 60% | 2.00E-56 | 53.45 | 329 | XP_010561166.1 |
| RNA/RNP complex-1-interacting phosphatase isoform X1 [Aquila chrysaetos chrysaetos] | Aquila chrysaetos chrysaetos | 195 | 195 | 60% | 2.00E-56 | 53.45 | 329 | XP_029890718.1 |

| Description | Scientific Name | Max Score | Total Score | Query Cover | E value | Per. ident | Acc. Len | Accession |
|---|---|---|---|---|---|---|---|---|
| *putative RNA phosphatase [Albatrosspox virus]* | *Albatrosspox virus* | *516* | *516* | *100%* | *0* | *100* | *251* | *QRM16049.1* |
| **RNA/RNP complex-1-interacting phosphatase [Corvus hawaiiensis]** | **Corvus hawaiiensis** | **199** | **199** | **70%** | **8.00E-59** | **53.41** | **305** | **XP_048146001.1** |
| RNA/RNP complex-1-interacting phosphatase [Corvus cornix cornix] | Corvus cornix cornix | 199 | 199 | 70% | 4.00E-58 | 53.41 | 349 | XP_039420096.1 |
| RNA/RNP complex-1-interacting phosphatase [Calypte anna] | Calypte anna | 196 | 196 | 69% | 4.00E-57 | 52.87 | 320 | XP_030319948.1 |
| DUS11 phosphatase [Upupa epops] | Upupa epops | 191 | 191 | 68% | 4.00E-57 | 53.22 | 169 | NWU89504.1 |
| RNA/RNP complex-1-interacting phosphatase isoform X3 [Accipiter gentilis] | Accipiter gentilis | 194 | 194 | 69% | 6.00E-57 | 52.87 | 271 | XP_049686642.1 |
| RNA/RNP complex-1-interacting phosphatase isoform X2 [Aquila chrysaetos chrysaetos] | Aquila chrysaetos chrysaetos | 195 | 195 | 69% | 7.00E-57 | 52.87 | 314 | XP_029890719.1 |
| RNA/RNP complex-1-interacting phosphatase isoform X3 [Aquila chrysaetos chrysaetos] | Aquila chrysaetos chrysaetos | 193 | 193 | 69% | 7.00E-57 | 52.87 | 271 | XP_040984244.1 |
| DUS11 phosphatase [Struthidea cinerea] | Struthidea cinerea | 190 | 190 | 68% | 7.00E-57 | 54.39 | 169 | NXB60100.1 |
| DUS11 phosphatase [Pachycephala philippinensis] | Pachycephala philippinensis | 190 | 190 | 68% | 7.00E-57 | 53.8 | 169 | NXI05060.1 |
| RNA/RNP complex-1-interacting phosphatase isoform X3 [Aythya fuligula] | Aythya fuligula | 191 | 191 | 69% | 8.00E-57 | 52.87 | 197 | XP_032059945.1 |
| PREDICTED: RNA/RNP complex-1-interacting phosphatase isoform X3 [Haliaeetus leucocephalus] | Haliaeetus leucocephalus | 195 | 195 | 69% | 1.00E-56 | 52.87 | 329 | XP_010561166.1 |

**Table S10. Accession IDs used to retrieve FASTA sequence for construction of phylogenetic tree.**

| Description | Name | Abreviation/Common Name | Accession | Viral/Host |
|---|---|---|---|---|
| DUSP11 | *Megadyptes antipodes* | yellow-eyed penguin | KAF1474841.1 | Host |
| DUSP11 | *Oryzias melastigma* | medakas | KAF6721144.1 | Host |
| DUSP11 | *Pleurodeles waltl* | newt | KAJ1091596.1 | Host |
| DUSP11 | *Danio rerio* | zebrafish | NP_001315299.1 | Host |
| DUSP11 | *Homo sapiens* | human | NP_003575.3 | Host |
| DUSP11 | *Mus musculus* | mouse | NP_082375.4 | Host |
| DUSP11 | *C. elegans* | nematode | NP_495959.2 | Host |
| DUSP11 | *Gymnorhina tibicen* | magpie | NXM43457.1 | Host |
| DUSP11 | *Calonectris borealis* | shearwater | NXV95132.1 | Host |
| DUSP11 | *Podilymbus podiceps* | flamingos | NXL53793.1 | Host |
| DUSP11 | *Pelecanus crispus* | pelican | XP_009482291.1 | Host |
| DUSP11 | *Struthio camelus australis* | ostrich | XP_009678555.1 | Host |
| DUSP11 | *Falco peregrinus* | falcons | XP_013152189.1 | Host |
| DUSP11 | *Gallus gallus* | Chicken | XP_015157914.2 | Host |
| DUSP11 | *bat* | microchiroptera | XP_016058068.1 | Host |
| DUSP11 | *Crocodylus porosus* | crocodiles | XP_019411446.1 | Host |
| DUSP11 | *Petromyzon marinus* | Lampreys | XP_032818491.1 | Host |
| DUSP11 | *Tyto alba* | owl | XP_032841029.2 | Host |
| DUSP11 | *Chelonia mydas* | sea turtle | XP_037742174.1 | Host |
| DUSP11 | *Xenopus laevis* | clawed frog | XP_041440763.1 | Host |
| DUSP11 | *Serinus canaria* | canary | XP_050838751.1 | Host |
| DUSP11 | *Ahaetulla prasina* | snakes | XP_058050012.1 | Host |
| DUSP11 | *Haemorhous mexicanus* | finch | XP_059726712.1 | Host |
| DUSP11 | *Anolis carolinensis* | green anole | XP_062817178.1 | Host |
| DUSP11 | *Columba livia* | rock dove | XP_064897185.1 | Host |
| vDUSP11 | Shearwaterpox virus | SWPV2 | ARE67303.1 | Viral |
| vDUSP11 | Shearwaterpox virus | SWPV1 | ARF02681.1 | Viral |
| vDUSP11 | Canarypox virus | CNPV | NP_955108.1 | Viral |
| vDUSP11 | Magpiepox virus | MPPV | QGM48715.1 | Viral |
| vDUSP11 | Cheloniid poxvirus 1 | ChePV1 | QRI42800.1 | Viral |
| vDUSP11 | Mudlarkpox virus | MLPV | QRM15361.1 | Viral |
| vDUSP11 | Penguinpox virus 2 | PEPV2 | QRM15716.1 | Viral |
| vDUSP11 | Albatrosspox virus | ALPV | QRM16049.1 | Viral |
| vDUSP11 | Finch poxvirus | FNPV | UOX38764.1 | Viral |
